# Deuteration improves small-molecule fluorophores

**DOI:** 10.1101/2020.08.17.250027

**Authors:** Jonathan B. Grimm, Liangqi Xie, Jason C. Casler, Ronak Patel, Ariana N. Tkachuk, Heejun Choi, Jennifer Lippincott-Schwartz, Timothy A. Brown, Benjamin S. Glick, Zhe Liu, Luke D. Lavis

**Affiliations:** Janelia Research Campus, The Howard Hughes Medical Institute, 19700 Helix Drive, Ashburn, VA 20147, USA; Department of Molecular Genetics and Cell Biology, University of Chicago, 920 East 58^th^ Street, Chicago, IL 60637, USA

## Abstract

Fluorescence microscopy relies on dyes that absorb short-wavelength photons and emit longer-wavelength light. In addition to this fluorescence process, dyes can undergo other photochemical reactions that result in spectral shifts and irreversible photobleaching. Increases in brightness, ‘chromostability’, and photostability of fluorescent dyes are therefore crucial for advancing the frontier of bioimaging. Here, we describe a general approach to improve small-molecule fluorophores using deuteration. Incorporating deuterium into the alkylamino substituents of rhodamines and other dyes improves fluorescence quantum yield, inhibits photochemically induced spectral shifts, and slows irreparable photobleaching. These compounds are easily synthesized and show improved performance in cellular imaging experiments.

---

Rhodamine dyes remain in wide use due to their excellent brightness, superb photostability, and tunable spectral and chemical properties.^1–4^ The photophysics of rhodamines are well understood, due to their importance as laser dyes and biological probes.^5^ Absorption of a photon excites a dye such as tetramethylrhodamine (TMR, **1**, Scheme 1) from the ground state (**1**-S_0_) ultimately to the first excited state (**1**-S_1_). After excitation, the molecule can relax back to **1**-S_0_ through different processes. Emission of a photon (fluorescence) competes with nonradiative decay pathways such as twisted internal charge transfer (TICT), where electron transfer from the aniline nitrogen to the xanthene system gives a charge-separated species (**1**-TICT) that decays back to **1**-S_0_ without emitting a photon.^6^ Alternatively, the excited dye can undergo intersystem crossing (ISC) to the first triplet excited state (**1**-T_1_) where it can sensitize singlet oxygen (^1^O_2_), returning to **1**-S_0_. The resulting ^1^O_2_ can then oxidize the aniline nitrogen to the radical cation (**1^+•^**), which can undergo deprotonation to a carbon-centered radical (**1^•^**) that ultimately dealkylates to give trimethylrhodamine (**2**).^5^ This results in a blue-shift in absorption (*λ*_bs_) and emission (*λ*_m_) maxima and is a prelude to irreversible photobleaching.

Both of these undesirable processes—TICT and dealkylation—can be mitigated through modifications in chemical structure. As both involve oxidation of the aniline nitrogen, methods to increase its ionization potential can improve both brightness and photostability. This was first demonstrated by Drexhage, a pioneer in dye chemistry, who found that replacing the *N,N*-dimethylamino groups in **1** with 5-membered pyrrolidine rings to give **3** (Figure 1a) enhanced fluorescence quantum yield (*Φ*).^7^ We advanced this idea, discovering that smaller, 4membered azetidine rings (*e.g.*, **4**) further improved the brightness and photostability of rhodamines and other fluorophores, yielding the ‘Janelia Fluor’ (JF) dyes. Additionally, the dealkylation process can be prevented by installing α-quaternary centers on the aniline nitrogens, thereby precluding deprotonation (Scheme 1). This structural motif was also introduced by Drexhage;^8^ it is found in a number of commercial fluorophores (*e.g.*, **5**-**7**, Figure 1b, Figure S1a)^9^ and this concept was revisited in the simplified di*t*-butylrhodamine (**8**).^10^

**Figure 1.**
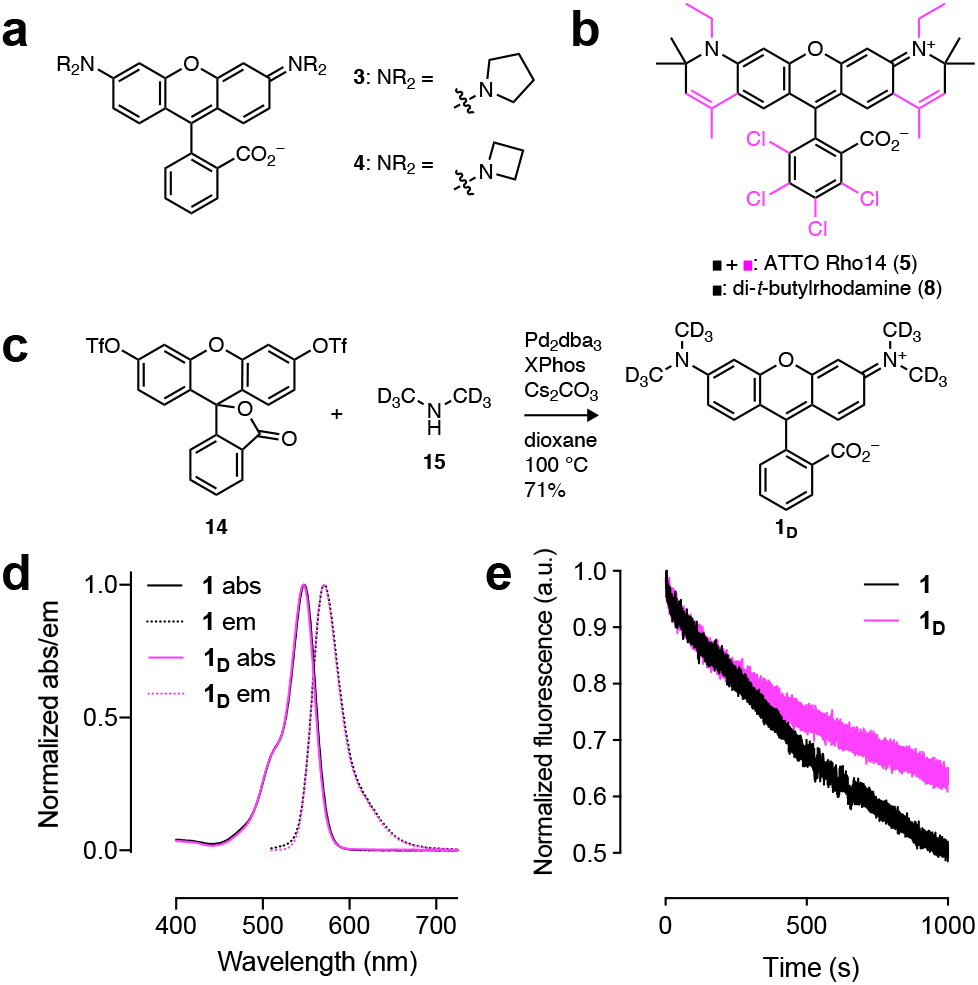
Improving rhodamine properties. (**a**) Structures of rhodamines **3** and **4**. (**b**) Structures of rhodamines **5** and **8**. (**c**) Synthesis of **1_D_**. (**d**) Normalized absorption (abs) and fluorescence emission (em) spectra of **1** and **1_D_**. (**e**) Photobleaching plot of normalized fluorescence intensity *vs.* illumination time of **1** and **1_D_**.

We envisioned an alternative strategy to increase brightness and photostability of small-molecule fluorophores such as **1** by replacing the hydrogen (H) atoms in the *N*-alkyl groups with deuterium (D). The oxidation of alkylamines shows a remarkably large secondary isotope effect,^11^ suggesting that deuteration could decrease the efficiency of the TICT process, therefore increasing *Φ*. Of course, this effect would also slow ^1^O_2_-mediated oxidation (**1**→**1**^+•^) and the stronger C.D bond could lower the rate of deprotonation (**1**+→**1**^•^, Scheme 1), together decreasing undesired dealkylation and improving both ‘chromostability’ (*i.e.*, reducing the undesirable shift in *λ*_abs_ and *λ*_em_) and photostability.

**Scheme 1.**
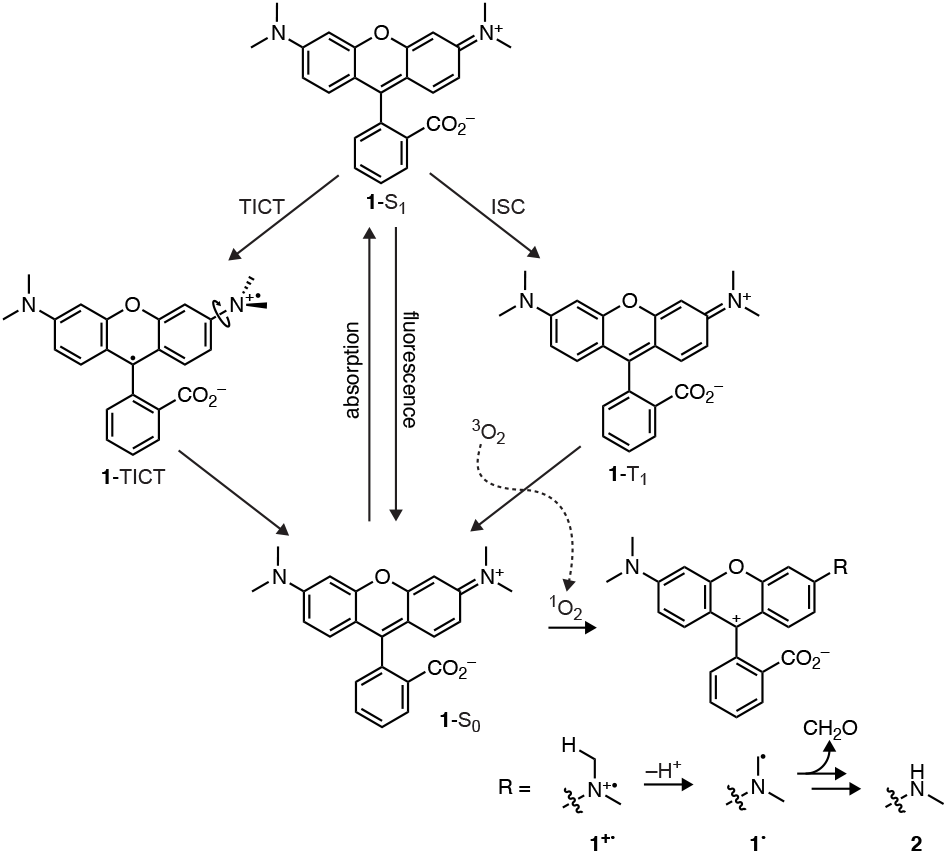
Photophysics of tetramethylrhodamine (TMR, 1)

Turro suggested deuteration could increase fluorescence quantum yield by altering vibrational modes12 and many fluorophores show improved brightness and photostability in deuterated solvents.13-14 Prior examples of deuterium-containing dyes are rare, however, due to the general difficulty of fluorophorecsynthesis.^2^ In the few extant examples, deuteration typically gives a negative or neutral effect on *Φ* as demonstrated for compounds **9**-**13** (Figure S1b).^15–17^ It was therefore unclear if this strategy of deuterating the *N*-alkyl groups would improve rhodamine properties. To test this hypothesis, the deuterated TMR analog **1_D_** was synthesized using a cross-coupling approach with fluorescein ditriflate (**14**) and dimethylamine-*d*_6_ (**15**; Figure 1c).^18^ Comparison of dyes **1** and **1_D_** revealed remarkably similar *λ*_abs_ and *λ*_em_, high extinction coefficients at *λ*_abs_ (*ε;* Table 1), and no change in the shape of the absorption or fluorescence emission spectra (Figure 1d). Deuteration did affect the brightness and photostability of the dye, however, with **1_D_** showing a ~15% increase in *Φ* compared to **1** (Table 1) and slower rate of photobleaching (Figure 1e).

**Table 1.**
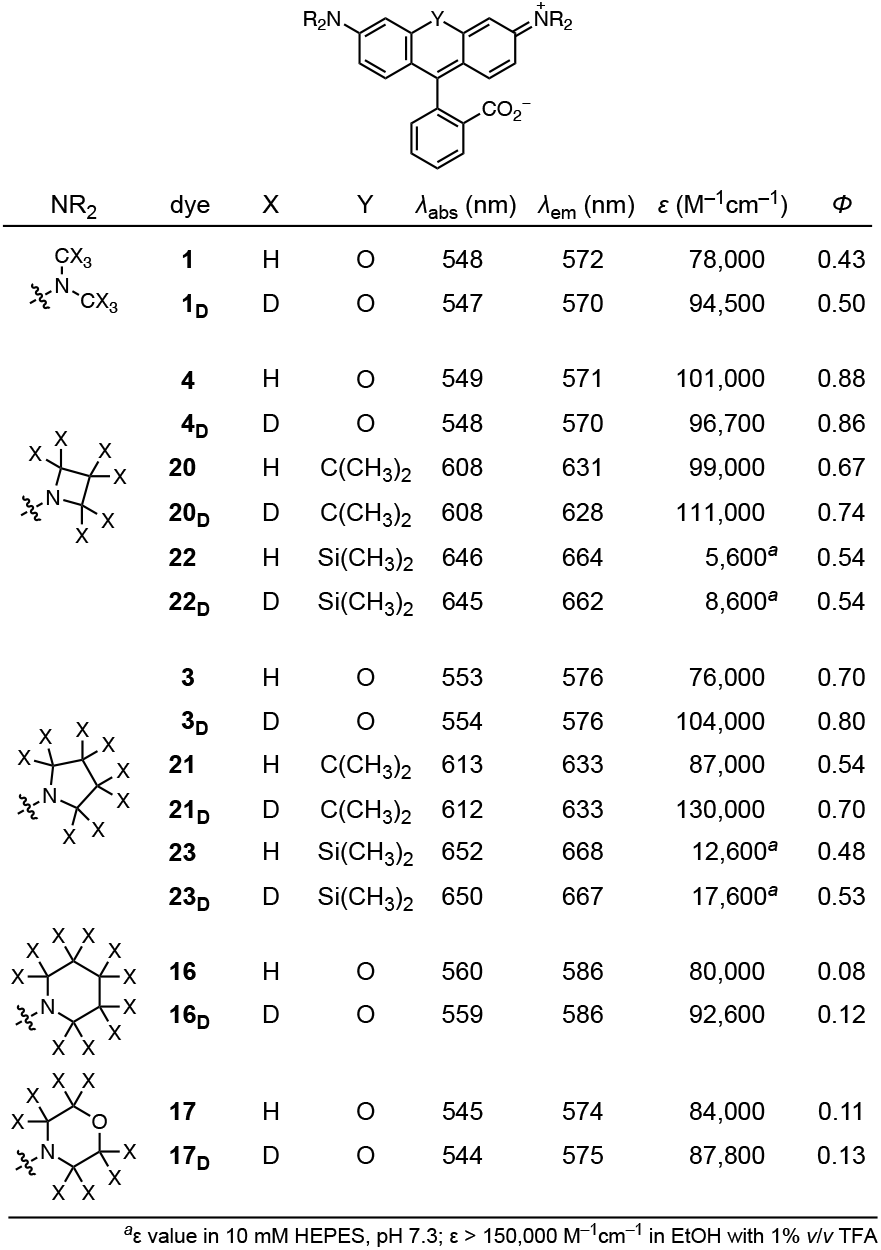
Spectral properties of rhodamines

Based on this result with TMR (**1**), other matched pairs of rhodamine dyes with H- or D-substituted cyclic *N*-alkyl groups were synthesized (Table 1, Figure S2a). Like the TMR compounds **1** and **1_D_**, increased *Φ* values were observed for the compounds containing pyrrolidine-*d*_8_ (**3_D_**) piperidine-*d_10_* (**16_D_**) and morpholine-*d*_8_ (**17_D_**) compared to the parent fluorophores **3**, **16**, and **17** (Table 1)^18^ without a change in spectral shape (Figure S2b–e). Interestingly, the deuterated azetidine compound (**4_D_**) showed no improvement in *Φ* over **4** (JF_549_; **Table 1**). This result suggests that the azetidine and deuterium substitutions suppress the same nonradiative pathways, such as TICT (Scheme 1), and are therefore not additive.

The photostability and chromostability of the brightest variants—those containing azetidine and pyrrolidine substituents— were evaluated by intermittently measuring the fluorescence emission spectra of the fluorophores during photobleaching experiments (Figure 2a). The nondeuterated JF_549_ (**4**) showed a steady rate of bleaching with a concomitant shift in *λ*_em_. Deuterated rhodamine **4_D_** bleached slower and exhibited higher resistance to undesirable dealkylation evidenced by the reduced shift in *λ*_em_ (Figure 2a). Compound **3** exhibited more complicated behavior with an initial increase in intensity along with a hypsochromic spectral shift followed by rapid bleaching (Figure 2b). This result can be explained by the lower *Φ* of **3**; dealkylation of this unoptimized dye yields a brighter trialkyl species, which then rapidly bleaches. Deuteration reduces this behavior, with **3_D_** showing higher photo- and chromo-stability. Based on these results, dyes **4_D_** and **3_D_** were named ‘JFX_549_’ and ‘JFX_554_’, respectively, to denote the <u>e</u>xtra stability afforded by deuteration.

**Figure 2.**
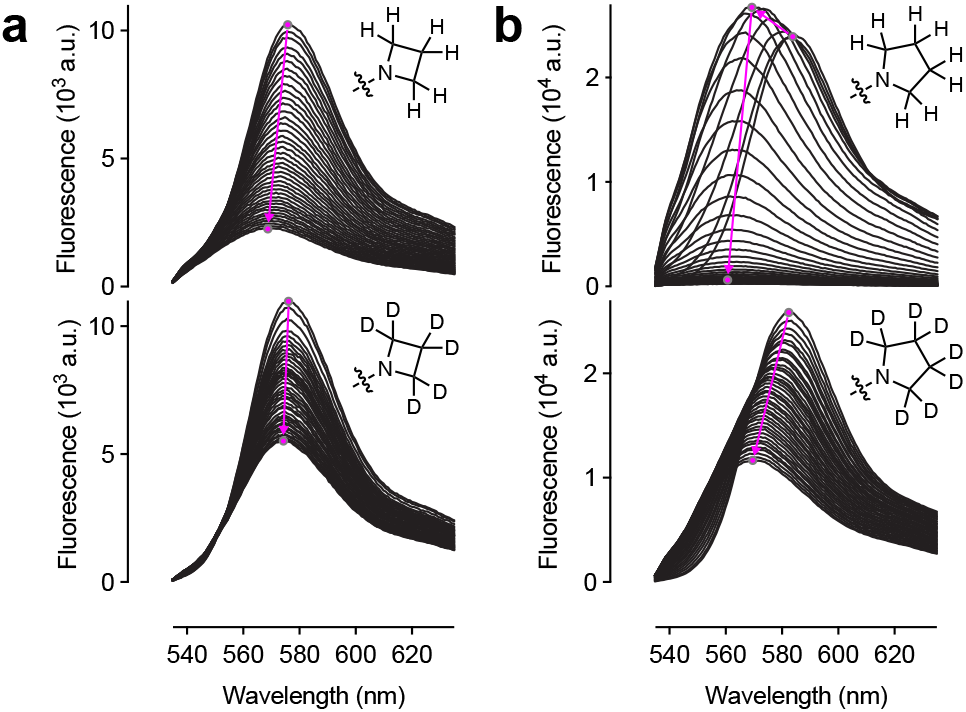
Photostability and chromostability of 3, 3_D_, 4, and 4_D_. (**a**–**b**) Sequential fluorescence emission spectra of (**a**) **4** and **4_D_** or (**b**) **3** and **3_D_** during photobleaching. The magenta arrows show the shift in *λ*_em_ and intensity over time.

The effect of deuteration on the properties of fluorophore:protein conjugates was then evaluated. The HaloTag^19^ ligand of **4** (JF_549_-HaloTag ligand, **18**) was previously synthesized starting from a 6-carboxyfluorescein derivative.^6^ This approach was used to prepare the JFX_549_-HaloTag ligand (**18_D_**) and JFX_554_-HaloTag ligand (**19_D_**), along with the HaloTag ligand of **3** (**19**; Figure 3, Scheme S1). Comparison of the HaloTag conjugates of **18** and **18_D_** *in vitro* revealed a small but significant increase in *Φ* for the **18_D_**:HaloTag conjugate compared to the nondeuterated **18**-labeled protein (Figure 3b). The free dyes **4** and **4_D_** show little difference in photophysical properties (Table 1), so this result suggests that deuteration suppresses a protein-bound-specific mode of nonradiative decay. Preparation of the HaloTag conjugates of pyrrolidinyl ligands **19** and **19_D_** (Figure 3a) allowed evaluation of the photostability and chromostability of all four labeled proteins (Figure S3a–b). Overall, the HaloTag conjugates showed substantially improved stability compared to the free dyes (Figure 2) demonstrating that the local environment around the fluorophore can substantially affect photophysics. Deuteration further improved properties, however, with the conjugates of **18_D_** and **19_D_** bleaching slower than those from **18** and **19**.

**Figure 3.**
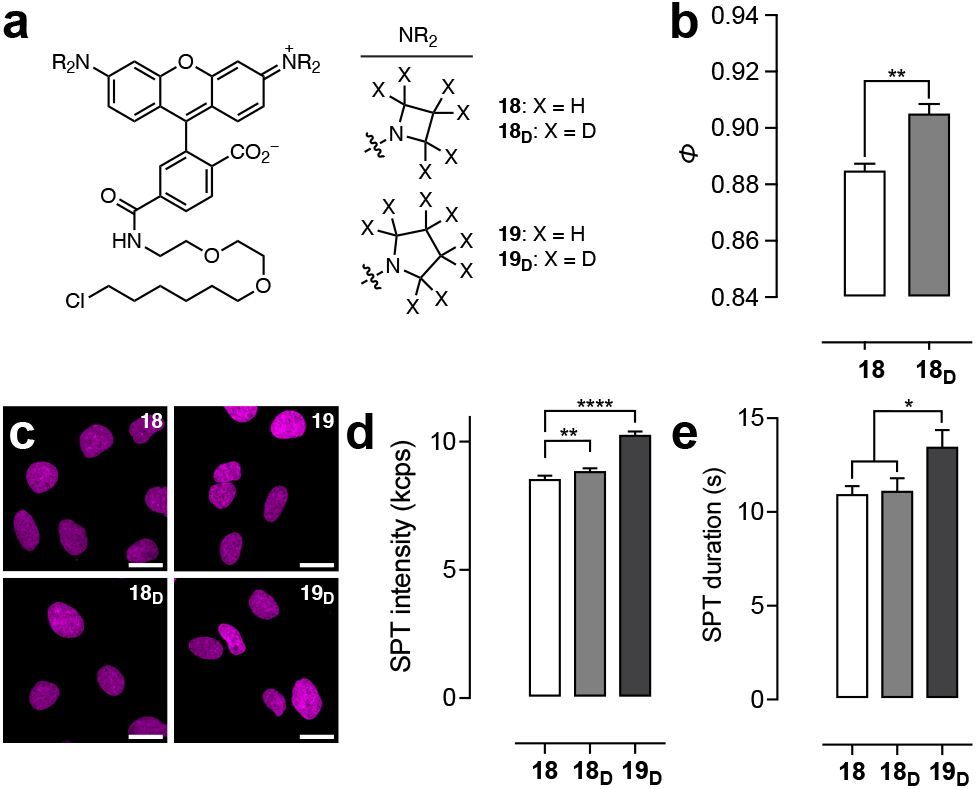
Performance of rhodamine ligands. (**a**) Structures of HaloTag ligands **18**, **18_D_, 19**, and **19_D_**. (**b**) *Φ* of HaloTag protein conjugates of **18** or **18_D_**. (**c**) Confocal microscopy images of live U2OS cells expressing HaloTag–histone H2B incubated with HaloTag ligands **18**, **18_D_, 19**, and **19_D_**; scale bars: 21 μm. (**d–e**) SPT intensity (**d**) or duration (**e**) from cells labeled with **18**, **18_D_**, or **19_D_**. All error bars: SEM.

The HaloTag ligands were then evaluated in live-cell experi-ments. All four ligands could label HaloTag–histone H2B fusions in living cells (Figure 3c) and showed similar loading kinetics (Figure S3c). **18_D_** showed a significantly longer fluorescence lifetime (*τ*) than **18** in live cells (Figure S3d), in line with the *in vitro Φ* measurements (Figure 3b). The utility in live-cell single-particle tracking (SPT) was then assessed with a focus on compounds JF_549_-HaloTag ligand (**18**), JFX_549_-HaloTag ligand (**18_D_**), and JFX_554_-HaloTag ligand (**19_D_**) since their parent fluorophores exhibit the highest molecular brightness (*ε* × *Φ*; Table 1). Pyrrolidine compound **19** showed significantly poorer performance compared to **18** in initial experiments (Figure S3e) and was not investigated further. Deuteration elicited modestly higher brightness (*i.e.,* photons/s; Figure 3d) for the azetidine **18_D_** compared to **18** under equivalent imaging conditions. Compound **18_D_** showed no improvement in photostability (*i.e.*, average duration of individual molecules; Figure 3e) over **18**. The deuterated pyrrolidine rhodamine ligand **19_D_** exhibited significantly higher brightness and photostability compared to azetidines **18** and **18_D_**, however, making it an attractive new label for cellular imaging.

This deuteration strategy was then applied to carborhodamines^20^ (**20**–**21**) and Si-rhodamines^21^ (**22**–**23**; Table 1, Scheme S2). For carborhodamines, both the deuterated azetidine (**20_D_**) and deuterated pyrrolidine (**21_D_**) showed increases of 10–20% in *Φ* compared to the parent dyes **20** and **21**. The Si-rhodamine dyes were similar to the rhodamine series, however, with azetidines **22** (JF_646_) and **22_D_** giving the same *Φ*; the deuterated pyrrolidine **23_D_** exhibited a higher *Φ* than **23**. Like the rhodamine series, fluorophores **22_D_** and **23_D_** were named ‘JFX_646_’ and ‘JFX_650_’, respectively. Deuteration also increased *Φ* in pyrrolidine-containing coumarins (**24**–**24_D_**) and phenoxazines (**25**– **25_D_**; Table S1).

The HaloTag ligands of the new Si-rhodamine compounds were then prepared (**26_D_**, **27**–**27_D_**) based on the previous synthesis of JF_646_-HaloTag ligand^6^ (**26**; Figure 4a, Scheme S3). These compounds could label HaloTag fusions in live cells (Figure 4c) with similar loading profiles (Figure S4a). Like the analogous rhodamines, the *Φ* of the deuterated azetidine JFX646-HaloTag ligand (**26_D_**) was significantly higher than the parent ligand **26** when conjugated to the HaloTag protein (Figure 4b) despite the equivalent values of the parent fluorophores (Table 1); **26_D_** also showed longer *τ* in cells (Figure S4b). Mirroring the rhodamine series (Figure 3d–e), compounds **26**, **26_D_**, and **27_D_** were tested in live-cell SPT experiments. The deuterated compounds showed marked improvements in brightness (Figure 4d) and photostability (Figure 4e) with JFX_650_-HaloTag ligand (**27_D_**) exhibiting the best overall performance. Moving beyond SPT, these Si-rhodamine ligands were evaluated in confocal imaging experiments using a strain of *S. cerevisiae* expressing a Sec7-GFP-HaloTag fusion to visualize the late Golgi apparatus. The results were consistent with the SPT experiments, with the deuterated JFX ligands **26_D_** and **27_D_** significantly brighter (Figure 4f) and more photostable (Figure 4g, Figure S4c–d, Movie S1) than **26**. Similar to the rhodamine dyes (Figure 3d–e), the deuterated pyrrolidine JFX_650_-HaloTag ligand (**27_D_**) showed the best performance.

**Figure 4.**
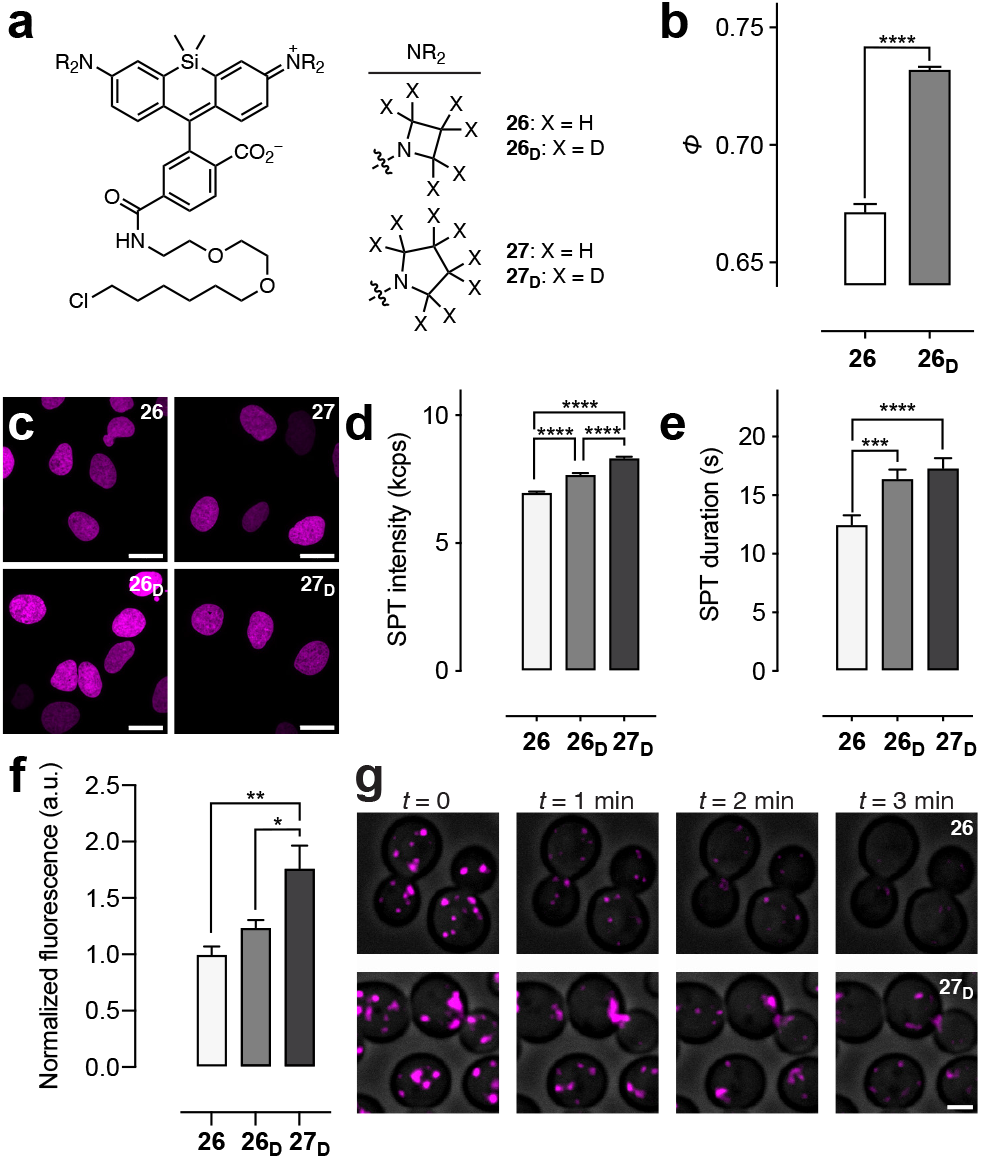
Performance of Si-rhodamine ligands. (**a**) Structures of HaloTag ligands **26**, **26_D_, 27**, and **27_D_**. (**b**) *Φ* of HaloTag protein conjugates of **26** or **26_D_**. (**c**) Confocal microscopy images of live U2OS cells expressing HaloTag–histone H2B incubated with HaloTag ligands **26**, **26_D_, 27**, and **27_D_**; scale bars: 21 μm. (**d–e**) SPT intensity (**d**) or duration (**e**) from cells labeled with **26**, **26_D_**, or **27_D_**. (**f**) Intensity from *S. cerevisiae* labeled with **26**, **26_D_**, or **27_D_**. (**g**) Image montage of *S. cerevisiae* labeled with **26** or **27_D_**; scale bar: 2 μm. All error bars: SEM.

In summary, we hypothesized that deuteration of the *N*-alkyl groups of rhodamine dyes would improve brightness, chromostability, and photostability based on known photophysics of rhodamines (Scheme 1). This hypothesis was tested first using the classic fluorophore TMR (**1**; Figure 1) and then by synthesizing other deuterated rhodamines; we discovered that deuteration maintained or improved *Φ* (Table 1) and enhanced chromo- and photostability (Figure 2). Synthesis of HaloTag ligands showed that deuterated rhodamines were superior labels for live-cell SPT experiments (Figure 3). This deuteration strategy was generalizable to other fluorophore classes (Table 1, Table S1) and the deuterated Si-rhodamines also showed improved performance in cellular imaging experiments (Figure 4). Overall, this work establishes deuteration of *N*-alkyl groups as a novel strategy for improving the properties of small-molecule fluorophores and yields new ‘JFX’ labels that exhibit superior performance in bioimaging compared to the original Janelia Fluor dyes.^6^ Future work will focus on deploying these new labels in other advanced imaging experiments, combining this strategy with complementary fluorophore tuning methods,^3–4^ and extending this approach to improve other chromophores containing *N*-alkyl groups.

## SUPPORTING INFORMATION

Supplementary figures, schemes, and table; experimental details; characterization for all new compounds; Movie S1

