## Supporting Information for "Deuteration improves small-molecule fluorophores"

### Experimental Information

| Page | Contents |
|---|---|
| S2 | Figures S1–S4 |
| S6 | Schemes S1–S3 |
| S8 | Table S1 |
| S9 | Experimental Details for Spectroscopy and Imaging |
| S14 | Experimental Details and Characterization Data for All Compounds |
| S31 | References |
| S32 | NMR and HPLC/MS |

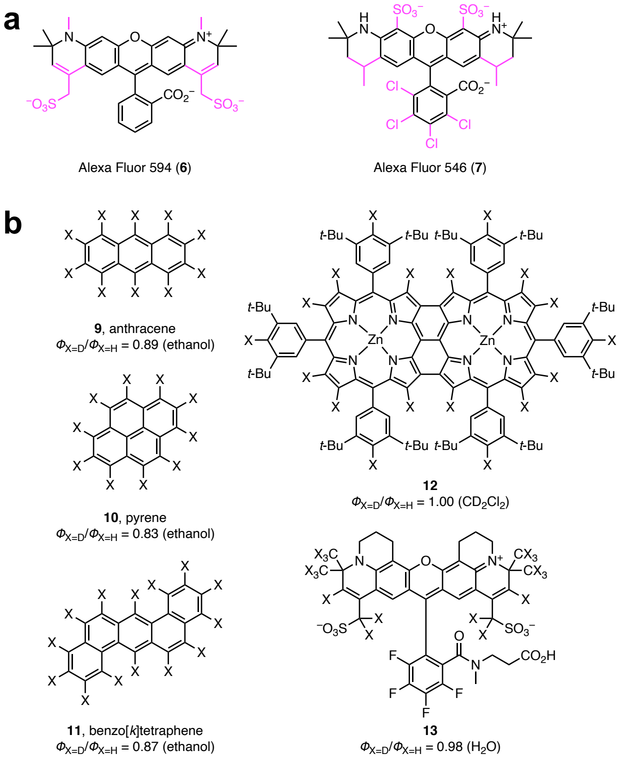


**Figure S1.** (**a**) Chemical structures of Alexa Fluor 594 (**6**) and Alexa Fluor 546 (**7**); black+magenta indicates the fluorophore structure and black-only represents the core di-*t*-butylrhodamine (**8**) structure. (**b**) Chemical structures and fluorescence quantum yields ($\Phi$) of matched pairs of nondeuterated (X = H) and deuterated (X = D) fluorophores, including polycyclic aromatic compounds (**9**–**11**), a porphyrin (**12**), and a rhodamine (**13**).

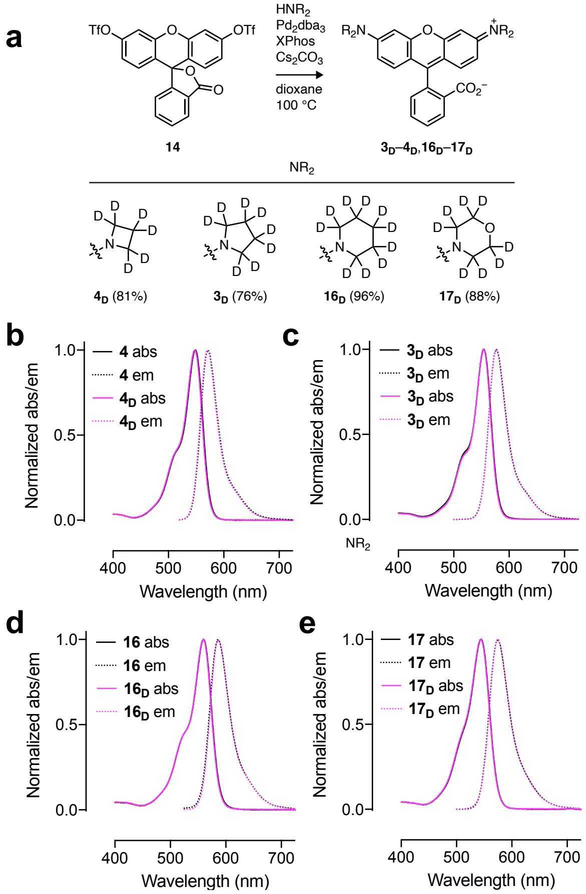


**Figure S2.** (**a**) Synthesis $\mathbf{3_D}$–$\mathbf{4_D}$ and $\mathbf{16_D}$–$\mathbf{17_D}$ from fluorescein ditriflate (**14**). (**b**–**e**) Normalized absorption (abs) and fluorescence emission (em) spectra for matched pairs of nondeuterated and deuterated rhodamine dyes: (**b**) **4** and $\mathbf{4_D}$; (**c**) **3** and $\mathbf{3_D}$; (**d**) **16** and $\mathbf{16_D}$; (**e**) **17** and $\mathbf{17_D}$.

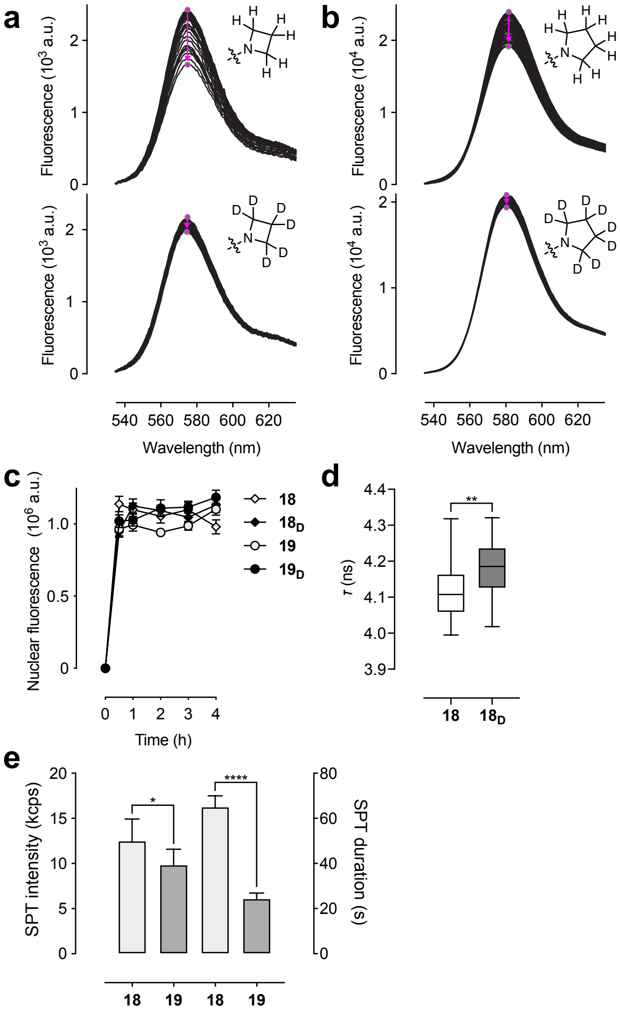


**Figure S3.** (**a**–**b**) Sequential fluorescence emission spectra of (**a**) **18**:HaloTag protein conjugate and $\mathbf{18_D}$:HaloTag protein conjugate or (**b**) **19**:HaloTag protein conjugate and $\mathbf{19_D}$:HaloTag protein conjugate during illumination; the magenta arrows show the shift in $\lambda_{em}$ over time. (**c**) Nuclear fluorescence *vs.* time upon addition of HaloTag ligands **18**, $\mathbf{18_D}$, **19**, and $\mathbf{19_D}$ (200 nM) to live U2OS cells expressing HaloTag–histone H2B; error bars indicate SEM. (**d**) Plot of fluorescence lifetime ($\tau$) of U2OS cells expressing HaloTag–histone H2B and labeled with HaloTag ligands **18** or $\mathbf{18_D}$; center line indicates median; box limits indicate upper and lower quartiles; whiskers indicate min–max. (**e**) Plot of average single-molecule intensity and trajectory length per cell in single-particle tracking experiments using JF$_{549}$-HaloTag ligand (**18**) or pyrrolidine-containing HaloTag ligand **19**; error bars indicate SEM.

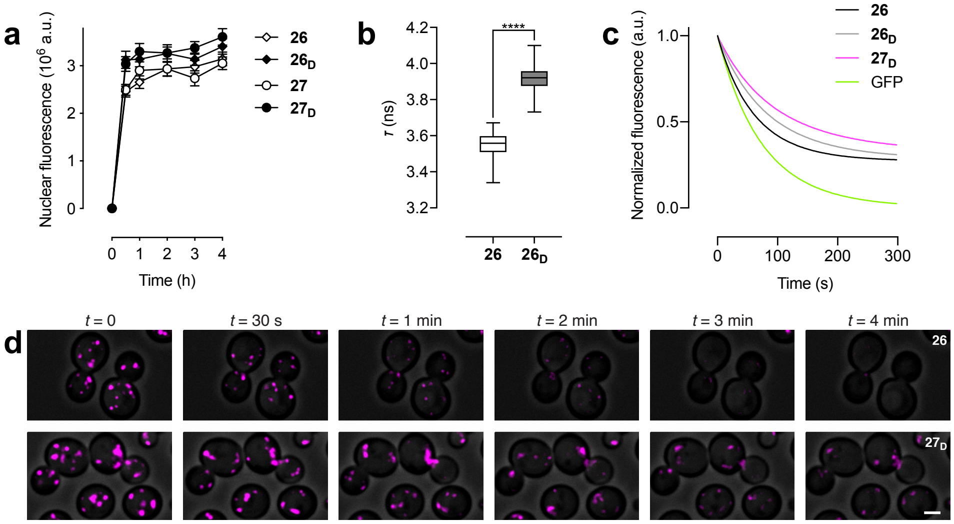


**Figure S4.** (**a**) Nuclear fluorescence *vs.* time upon addition of ligands **26**, **$26_D$**, **27**, or **$27_D$** (200 nM) to live U2OS cells expressing HaloTag–histone H2B; error bars indicate SEM. (**b**) Plot of fluorescence lifetime ($\tau$) of U2OS cells expressing HaloTag–histone H2B and labeled with HaloTag ligands **26** or **$26_D$**; center line indicates median; box limits indicate upper and lower quartiles; whiskers indicate min–max. (**c**) Photobleaching plot of normalized fluorescence intensity *vs.* time of *S. cerevisiae* cells expressing Sec7–iGFP–HaloTag fusions and labeled with ligands **26**, **$26_D$**, or **$27_D$**. (**d**) Expanded image montage from Figure 4g/Movie S1 showing *S. cerevisiae* labeled with **26** or **$27_D$**.; scale bar: 2 μm.

**Scheme S1.** Synthesis of HaloTag ligands $\mathbf{18_D}$, **19**, and $\mathbf{19_D}$.

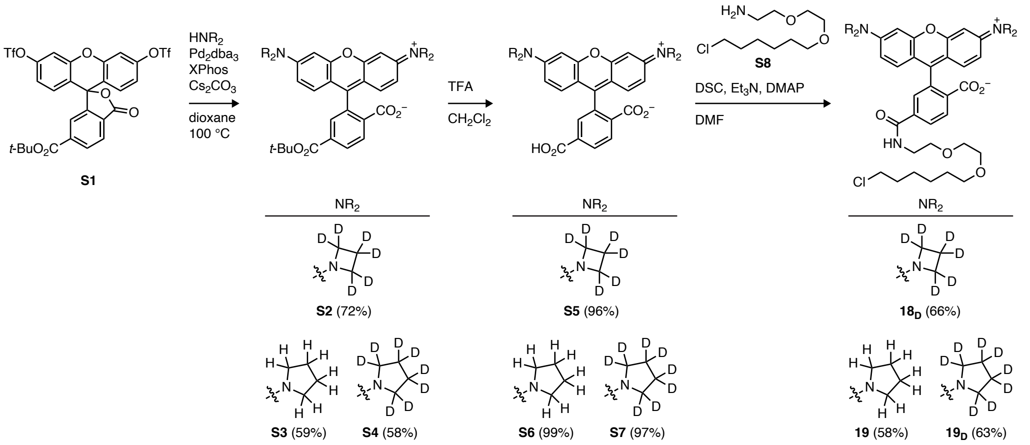


**Scheme S2.** Synthesis of nondeuterated and deuterated carborhodamines (**a**), Si-rhodamines (**b**), coumarins (**c**), and phenoxazines (**d**).

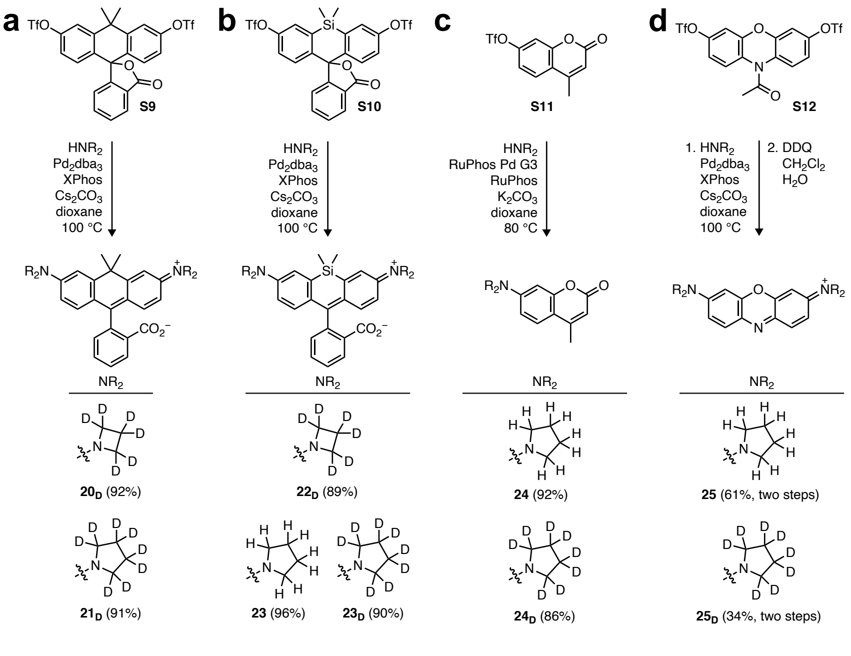

**Scheme S3.** Synthesis of HaloTag ligands $\mathbf{27_D}$, **28**, and $\mathbf{28_D}$.

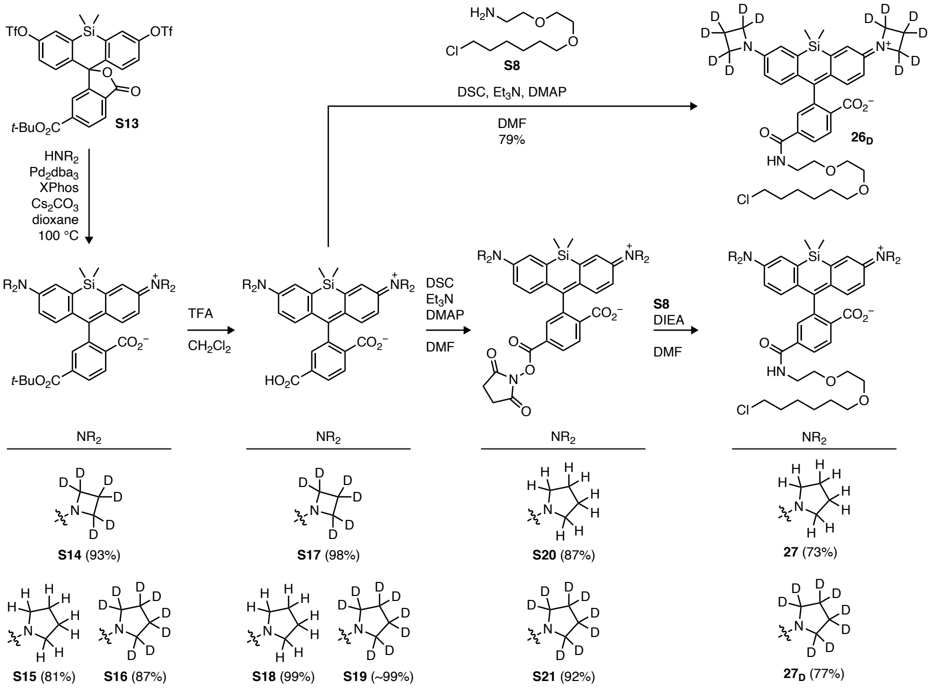

**Table S1.** Spectral properties of coumarin derivatives **24** and **$24_D$** and phenoxazine derivatives **25** and **$25_D$**.

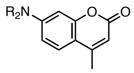


| $NR_2$ | dye | X | $\lambda_{max}$ (nm) | $\lambda_{em}$ (nm) | $\varepsilon$ ($M^{-1}cm^{-1}$) | $\Phi$ |
|---|---|---|---|---|---|---|
| 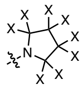 | **24** | H | 381 | 472 | 19,800 | 0.46 |
| | **$24_D$** | D | 381 | 472 | 19,700 | 0.56 |

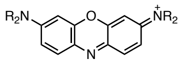


| $NR_2$ | dye | X | $\lambda_{max}$ (nm) | $\lambda_{em}$ (nm) | $\varepsilon$ ($M^{-1}cm^{-1}$) | $\Phi$ |
|---|---|---|---|---|---|---|
| 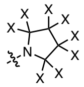 | **25** | H | 655 | 671 | 85,000 | 0.16 |
| | **$25_D$** | D | 653 | 669 | 85,000 | 0.22 |

### GENERAL EXPERIMENTAL INFORMATION FOR SPECTROSCOPY AND IMAGING

**Compound sources.** Tetramethylrhodamine (**1**; Reference Standard Grade) was purchased from AAT BioQuest. Compounds **3**, **4**, **16**–**18**, **20**–**22**, and **26** were available from previous work.[1-3]

**UV–vis and fluorescence spectroscopy (Figure 1d, Table 1, Figure S2b–e, Table S1).** Fluorescent molecules for spectroscopy were prepared as stock solutions in DMSO and diluted such that the DMSO concentration did not exceed 1% v/v. Spectroscopy was performed using 1-cm path length, 3.5-mL quartz cuvettes or 1-cm path length, 1.0-mL quartz microcuvettes from Starna Cells. All measurements were taken at ambient temperature (22 ± 2 °C). Absorption spectra were recorded on a Cary Model 100 spectrometer (Agilent). Fluorescence spectra were recorded on a Cary Eclipse fluorometer (Varian). Maximum absorption wavelength ($\lambda_{abs}$), extinction coefficient ($\varepsilon$), and maximum emission wavelength ($\lambda_{em}$) were measured in 10 mM HEPES, pH 7.3 buffer. Normalized spectra are shown for clarity.

**Quantum yield determination (Table 1, Table S1).** All reported absolute fluorescence quantum yield values ($\Phi$) were measured in our laboratory under identical conditions using a Quantaurus-QY spectrometer (model C11374, Hamamatsu). This instrument uses an integrating sphere to determine photons absorbed and emitted by a sample. Measurements were carried out using dilute samples ($A < 0.1$) and self-absorption corrections[4] were performed using the instrument software. Reported values are averages ($n = 3$).

**Photobleaching of 1 and $1_D$ (Figure 1e).** Solutions of **1** and **$1_D$** (200 nM) were prepared in 10 mM HEPES, pH 7.3. An aliquot of these solutions was added to each well of 10-well Teflon printed glass slide with 1.5 mm well diameter (Tekdon, Inc.) and sealed with a coverslip using vacuum grease. The hydrophobic coating of the slide resulted in formation of macrodroplets of aqueous dye solution in each well. Fluorophore bleaching was measured by illuminating an entire droplet using an upright microscope (Zeiss Axio Observer Z2) and a 5×/0.25 NA objective. Light illumination was provided by a mercury lamp (X-Cite Series 120-Q) with 550BP25 excitation filter at intensity 1.8 W/cm$^2$. Fluorescence emission was collected using 625BP90 filter and detected with a fiber-coupled avalanche photodiode detector (SPQM-AQRH14; Pacer); $n = 3$.

**Photostability and 'chromostability' measurements of dyes and HaloTag conjugates (Figure 2, Figure S3a–b).** To test the stability of the free dyes, solutions of **3**, **$3_D$**, **4**, and **$4_D$** (1 µM) were prepared in 10 mM HEPES, pH 7.3. To test the stability of the HaloTag conjugates, HaloTag protein was used as a 200 µM solution in PBS, pH 7.4. Solutions of HaloTag ligands **18**, **$18_D$**, **19**, or **$19_D$** (5 µM) was prepared in 10 mM HEPES, pH 7.3 containing 0.1 mg·mL$^{-1}$ CHAPS. An aliquot of HaloTag protein (2.0 equiv, 10 µM final [HaloTag]) was added and the resulting mixture was incubated at 4 °C overnight. Stability measurements were performed after the HaloTag conjugate solutions were diluted 5× (1 µM final [ligand]) into 10 mM HEPES, pH 7.3 buffer solution. These samples were prepared using the Teflon printed glass slide as described above. The fluorophore and HaloTag conjugate solutions were bleached using an upright microscope (Zeiss Axio Observer Z2) 5×/0.25 NA objective. Light illumination was provided by a 532 nm laser (Compass 215M-75; Coherent) at 18 mW power (intensity 0.96 $W/cm^2$). Fluorescence emission was collected using a 590/104 filter and emission spectra were measured using a fiber coupled spectrometer (QE 65000; Ocean Optics). Each sample was continuously bleached for 40 min with fluorescence spectra acquired every 1 min. Spectra are averages ($n$ = 2).

**Quantum yield determination of HaloTag protein conjugates (Figure 3b, Figure 4b).** HaloTag protein was used as a 100 µM solution in 75 mM NaCl, 50 mM TRIS·HCl, pH 7.4 with 50% v/v glycerol (TBS–glycerol). A solution of HaloTag ligands **18**, **$18_D$**, **27**, or **$27_D$** (5 µM) was prepared in 10 mM HEPES, pH 7.3 containing 0.1 $mg{\cdot}mL^{-1}$ CHAPS. An aliquot of HaloTag protein (1.5 equiv, 7.5 µM final [HaloTag]) was added and the resulting mixture was incubated until consistent absorption signal was observed (60–120 min). After incubation, this solution was diluted 5–10-fold in 10 mM HEPES, pH 7.3 containing 0.1 $mg{\cdot}mL^{-1}$ CHAPS to give 500 nm–1 µM dye concentration, which is optimal for quantum yield determination ($n$ = 14).

**Confocal fluorescence microscopy (Figure 3c, Figure 4c).** U2OS cells (ATCC) with an integrated histone H2B–HaloTag fusion protein expressing plasmid via the piggyBac transposon system were cultured in Dulbecco's modified Eagle medium (DMEM, phenol red-free; Life Technologies) supplemented with 10% v/v fetal bovine serum (FBS, Life Technologies), 1 mM GlutaMAX (Life Technologies) and maintained at 37 °C in a humidified 5% (v/v) $CO_2$ environment. These cell lines were kept under the selection of 500 µg/mL Geneticin (Life Technologies) and undergo regular mycoplasma testing by the Janelia Cell Culture Facility. Cells were incubated with HaloTag ligands **18**, **$18_D$**, **19**, **$19_D$**, **26**, **$26_D$**, **27**, or **$27_D$**. (200 nM, 2 h) and washed 3× with dye-free media. Cells were imaged on a Zeiss LSM 800 confocal microscope with a Plan APO 63×/1.4 NA oil DIC M27 objective. The confocal images were processed using FIJI[5] and displayed as maximum intensity image projections.

**Dye loading kinetics (Figure Sc3, Figure S4a).** Live U2OS cells stably expressing histone H2B–HaloTag fusion proteins were labeled over a time course of 0–4 h with 200 nM of HaloTag ligands **18**, **$18_D$**, **19**, **$19_D$**, **26**, **$26_D$**, **27**, or **$27_D$**. Cells were briefly washed 3× with dye-free media and immediately imaged live using widefield microscopy on a Nikon Eclipse Ti, Plan APO λ 20×/0.75 air objective. Fluorescence was quantified from the average of the summed intensity of nuclear signals in single-plane widefield images analyzed using Nikon NIS-Elements AR software. Fields of view were chosen randomly; $n$ = 100 nuclear signals per compound.

**In-cell fluorescence lifetime ($\tau$) measurements (Figure S3d, Figure S4b).** U2-OS cells stably expressing histone H2B–HaloTag protein fusions were plated on a coverslip pre-coated with human fibronectin (Millipore). After 24 h, the cells were incubated with 100 nM of dyes **18**, **$18_D$**, **26**, or **$26_D$** for 30 min in DMEM, (Corning) supplemented with 10% v/v FBS (Corning), 2 mM L-Glutamine, 100 U/mL penicillin and 100 µg/mL streptomycin at 37 °C and 5% $CO_2$. Prior to imaging the cells were briefly washed 3× with dye-free media. These cells were imaged using a Zeiss 880 laser scanning confocal microscope equipped with Ti:Sapphire laser (Coherent Chameleon). The labeled U2-OS cell samples were excited with 850 nm (**18** and **$18_D$**) or 830 nm (**26** and **$26_D$**). The excitation laser was filtered through a short-pass filter (675LP). The emitted light passed through a dichroic (DIC560) and collected by the PMA hybrid detector (PicoQuant) with bandpass filters (BPF600/50 for **18** and **18$_D$**; no filter for **26** and **26$_D$**) and fed to the single-photon counting board (TimeHarp 260, PicoQuant) and monitored at 80 MHz. The resulting histogram was fit to a single exponential decay function using SymPhoTime 64 (PicoQuant); $n$ = 30 for **18** and **18$_D$**, $n$ = 27 for **26**, and $n$ = 29 for **26$_D$**.

**Initial comparison of HaloTag ligands 18 and 19 in single-particle tracking (SPT) experiments (Figure S3e).** U2-OS cells stably expressing histone H2B–HaloTag protein fusions were plated on a coverslip pre-coated with human fibronectin (Millipore). After 24 h, the cells were incubated with 1 nM of either **18** or **19** for 30 min DMEM (Corning) supplemented with 10% v/v FBS (Corning), 2 mM L-Glutamine, 100 U/mL penicillin and 100 µg/mL streptomycin at 37 °C and 5% $CO_2$. Prior to imaging the cells were briefly washed 3× with dye-free media. These cells were imaged using a Zeiss Elyra microscope equipped with 561 nm laser excitation and a 100× Plan-Apochromat 1.45 NA objective (Zeiss) and EMCCD (Andor). The cells were illuminated using oblique illumination and spt-*d*STORM was performed as previously described.[6] The emitted light was collected with a multi-bandpass filter (BP420-480 + BP570-640 + LP740). The single particles are imaged at 20 Hz and trajectories were mapped using TrackMate. The average trajectory length (*i.e.*, duration time) per cell was quantified from all the trajectories with the maximum displacement r < 400 nm within a single step. SPT intensity was quantified using Zeiss Zen Black for the average intensity of all single-particle emitters inside the cells. For SPT intensity, $n$ = 7 cells using **18** and $n$ = 8 cells using **19**. For SPT duration time, $n$ = 7 cells using **18** and $n$ = 16 cells using **19**.

**Single-particle tracking (SPT) experiments (Figure 3d–e, Figure 4d–e).** SPT experiments were carried out as previously described.[7] JM8.N4 mouse embryonic stem cells (ESCs) stably expressing histone H2B–HaloTag fusion proteins were generated by coexpressing the PiggyBac EF1a-H2B-HaloTag-IRES-Neo vector with the super piggyBac transposase as described previously,[8] followed by G418 selection (500 µg/mL) for 2 weeks and verified by FACS sorting/confocal imaging staining with $JF_{549}$-HaloTag ligand (**18**; 100 nM, 30 min). Cells were seeded on 25 mm #1.5 coverglass pre-cleaned with KOH and ethanol and coated with Matrigel according to the manufacturer's instruction. All live cell imaging experiments were conducted using an ESC imaging medium composed of FluoroBrite DMEM (Thermo Fisher Scientific) plus 15% v/v ESC-qualified FBS, 1× GlutaMax, 1× NEAA, 0.1 mM 2-mercaptoethanol, and LIF. To achieve sparse labeling, the histone H2B–HaloTag ESCs were stained with 2 nM of HaloTag ligands **18**, **18$_D$**, **19$_D$**, **26**, **26$_D$**, or **27$_D$**. for 15 min at 37 °C with 5% v/v $CO_2$ and then washed with dye-free media 3× for 30 min each wash. To quantify the brightness and photostability of the HaloTag ligands, labeled cells were mounted onto a high speed motorized Nikon Eclipse Ti-E inverted microscope equipped with the following: 100× Apo TIRF 1.49 NA objective with a correction collar; four excitation laser lines (405/488/561/642 nm) and matching TIRF quad cube (405/488/561/640 nm reflection bands); automatic TIRF illuminator with motorized X axis and manual Y axis for beam positioning and focus: perfect focus 3 system; Triple DU-897 iXon Ultra EMCCD cameras on a Cairn Tri-cam emission splitter with filters 525/50 (GFP) 600/50 (RFP) and 705/72 (Cy5); humidified incubation chamber maintained at 37 °C with 5% v/v $CO_2$ (Tokai Hit). SPT was performed at 2 Hz for ~500 frames using `10% of maximal excitation power. The excitation laser was controlled by an acousto-optic tunable filter (AOTF) and reflected into the objective by a multi-band dichroic (405/488/561/633 BrightLine quad-band bandpass filter; Semrock). HaloTag ligands **18**, **$18_D$**, and **$19_D$** were illuminated with 561 nm laser, whereas **26**, **$26_D$**, and **$27_D$** were excited with a 641 nm laser. To minimize drift during imaging, the incubation chamber was fully thermally equilibrated and all imaging was conducted in an isolated ultra-clean room with minimal mechanical vibrations. The microscope, laser lines, and camera integration were controlled using the Nikon NIS-Elements software. We performed two independent biological replicates each with at least eight cells. Cells with obvious movement during acquisition were removed from further analysis. SMT analysis was based on multiple target tracing[9] (MTT) with tracking parameters similar to previous work.[7] For SPT brightness (kilocounts per second, kcps), intensity values from the first frame of the imaging experiment were analyzed: $n$ = 3351 single-molecule events using **18;** $n$ = 3838 single-molecule events using **$18_D$**; $n$ = 4724 single-molecule events using **$19_D$**; $n$ = 5523 single-molecule events using **26**; $n$ = 3295 single-molecule events using **$26_D$**; ; $n$ = 7125 single-molecule events using **$27_D$**. For SPT duration (s), the single-molecule trajectories of each cell were combined and a survival curve was generated based on the 1-cumulative distribution function (1-CDF) histogram and mathematical fitting (MATLAB) and previously described.[7] Histone H2B molecules generally considered immobile under the imaging conditions and so disappear from the detection plane primarily by photobleaching. Thus, the dissociation rate from this analysis equals the inverse of the fluorophore duration time, allowing comparison of photobleaching; $n$ = 15 cells using **18**; $n$ = 10 cells using **$18_D$**; $n$ = 11 cells using **$19_D$**; $n$ = 14 cells using **26**; $n$ = 13 cells using **$26_D$**; n = 12 cells using **$27_D$**.

**Imaging in Yeast (Figure 4f–g, Figure S4c–d).** A *Saccharomyces cerevisiae* strain expressing the late Golgi marker Sec7-iGFP-HaloTag fusion from the native locus was grown shaking at room temperature to mid-log phase in NSD. Cells were incubated with HaloTag ligands **26**, **$26_D$**, or **$27_D$**. (1 µM diluted from a 1mM stock in DMSO; 30 min), filter washed to remove excess dye, and adhered to a concanavalin A coated confocal dish. The cells were then imaged on a Leica SP8 laser scanning confocal microscope under identical conditions. Movie S1 is a combination of representative average projected z-stacks of cells labeled with the indicated dye. Deconvolution using Huygens Essential X11 software was performed to smooth the noisy confocal data. Average brightness per cell was calculated by measuring the total fluorescence intensity of the first frame of the movies and dividing by the total number of cells. The GFP fluorescence intensity was used as an internal control to correct for variation in expression between cells; $n$ = 25 for **26**; ; $n$ = 35 for **$26_D$**; $n$ = 37 for **$27_D$**. Photobleaching curves were generated via the EMBL bleach correction plug-in on ImageJ. The graph consists of the average photobleaching traces of movies ($n$ = 5) representing 25–40 cells per condition.

**Statistics and Reproducibility.** For spectroscopy measurements, photobleaching experiments, or photo- and chromo-stability evaluations, reported $n$ values represent measurements of different samples prepared from the same dye DMSO stock solution or HaloTag conjugate stock solution. For cell loading experiments, $n$ represents the number of intensity values from individual nuclei extracted from three random fields of view at the indicated time points. For initial comparison of HaloTag ligands **18** and **19** in SPT, $n$ indicates separate cells. For the comparison of the SPT intensity of HaloTag ligands **18**, **18**$_{D}$, and **19**$_{D}$, or **26**, **26**$_{D}$, and **27**$_{D}$, *n* indicates separate events extracted by the MTT algorithm. For full comparison of the SPT duration time of HaloTag ligands **18**, **18**$_{D}$, and **19**$_{D}$, or **26**, **26**$_{D}$, and **27**$_{D}$, *n* indicates values extracted from separate cells. For cell-loading experiments, *n* represents the number of intensity values from individual cells extracted from three fields of view at the indicated time points. For fluorescence microscopy experiments, all procedures were repeated at least once on a separate biological sample to ensure results were similar. Statistical analyses were performed using GraphPad Prism. P values were indicated as follows: 0.1234 (ns), 0.0332 (*), 0.0021 (**), 0.0002 (***), <0.0001 (****).

### General Experimental Information for Synthesis

Commercial reagents were obtained from reputable suppliers and used as received. All solvents were purchased in septum-sealed bottles stored under an inert atmosphere. Fluorescein ditriflate[1] (**14**), 6-*tert*-butoxycarbonylfluorescein ditriflate[3] (**S1**), carbofluorescein ditriflate[2] (**S9**), Si-fluorescein ditriflate[3] (**S10**), 4-methylumbelliferone triflate[10] (**S11**), 10-acetyl-10*H*-phenoxazine-3,7-diyl bis(trifluoromethanesulfonate)[3] (**S12**), and 6-*tert*-butoxycarbonyl-Si-fluorescein ditriflate[3] (**S13**) were synthesized as previously described. Azetidine-2,2,3,3,4,4-$d_6$ was used as either the free base (Cambridge Isotopes) or the hydrochloride salt (synthesized as previously described[11]). All reactions were sealed with septa through which a nitrogen atmosphere was introduced unless otherwise noted. Reactions were conducted in round-bottomed flasks or septum-capped crimp-top vials containing Teflon-coated magnetic stir bars. Heating of reactions was accomplished with a silicon oil bath or an aluminum reaction block on top of a stirring hotplate equipped with an electronic contact thermometer to maintain the indicated temperatures.

Reactions were monitored by thin layer chromatography (TLC) on precoated TLC glass plates (silica gel 60 $F_{254}$, 250 µm thickness) or by LC/MS (Phenomenex Kinetex 2.1 mm × 30 mm 2.6 µm C18 column; 5 µL injection; 5–98% MeCN/$H_2O$, linear gradient, with constant 0.1% v/v $HCO_2H$ additive; 6 min run; 0.5 mL/min flow; ESI; positive ion mode). TLC chromatograms were visualized by UV illumination or developed with *p*-anisaldehyde, ceric ammonium molybdate, or $KMnO_4$ stain. Reaction products were purified by flash chromatography on an automated purification system using pre-packed silica gel columns or by preparative HPLC (Phenomenex Gemini–NX 30 × 150 mm 5 µm C18 column). Analytical HPLC analysis was performed with an Agilent Eclipse XDB 4.6 × 150 mm 5 µm C18 column under the indicated conditions. High-resolution mass spectrometry was performed by the High Resolution Mass Spectrometry Facility at the University of Iowa.

NMR spectra were recorded on a 400 MHz spectrometer. $^{1}H$ and $^{13}C$ chemical shifts were referenced to TMS or residual solvent peaks. Data for $^{1}H$ NMR spectra are reported as follows: chemical shift (δ ppm), multiplicity (s = singlet, d = doublet, t = triplet, q = quartet, dd = doublet of doublets, m = multiplet), coupling constant (Hz), integration. Data for $^{13}C$ NMR spectra are reported by chemical shift (δ ppm) with hydrogen multiplicity (C, CH, $CH_2$, $CH_3$) information obtained from DEPT spectra.

### EXPERIMENTALS

#### Pd-Catalyzed Cross-Coupling Reactions

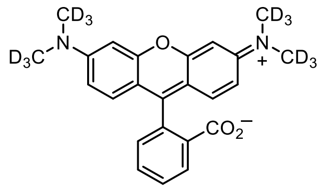

**TMR-$d_{12}$ ($1_D$).** A vial was charged with fluorescein ditriflate (**14**, 150 mg, 0.251 mmol), dimethyl-$d_6$-amine hydrochloride (**15**, 52.9 mg, 0.604 mmol, 2.4 eq), $Pd_2dba_3$ (23.0 mg, 25.1 μmol, 0.1 eq), XPhos (36.0 mg, 75.4 μmol, 0.3 eq), and $Cs_2CO_3$ (393 mg, 1.21 mmol, 4.8 eq). The vial was sealed and evacuated/backfilled with nitrogen (3×). Dioxane (1.5 mL) was added, and the reaction was flushed again with nitrogen (3×). The reaction was then stirred at 100 °C for 4 h. It was subsequently cooled to room temperature, diluted with MeOH, deposited onto Celite, and concentrated to dryness. Purification by silica gel chromatography (0–10% MeOH (2 M $NH_3$)/$CH_2Cl_2$, linear gradient; dry load on Celite) followed by reverse phase HPLC (10–50% MeCN/$H_2O$, linear gradient, with constant 0.1% v/v TFA additive) afforded $\mathbf{1_D}$ (92 mg, 71%, TFA salt) as a dark red solid. $^1H$ NMR ($CD_3OD$, 400 MHz) δ 8.37 – 8.32 (m, 1H), 7.86 (td, $J$ = 7.5, 1.5 Hz, 1H), 7.80 (td, $J$ = 7.6, 1.5 Hz, 1H), 7.44 – 7.38 (m, 1H), 7.15 (d, $J$ = 9.5 Hz, 2H), 7.05 (dd, $J$ = 9.5, 2.5 Hz, 2H), 6.97 (d, $J$ = 2.5 Hz, 2H); $^{13}C$ NMR ($CD_3OD$, 101 MHz) δ 168.0 (C), 161.8 (C), 159.1 (C), 159.0 (C), 135.3 (C), 133.8 (CH), 132.5 (CH), 132.3 (C), 132.1 (CH), 131.5 (CH), 131.4 (CH), 115.4 (CH), 115.0 (C), 97.3 (CH); Analytical HPLC: $t_R$ = 10.5 min, >99% purity (10–95% MeCN/$H_2O$, linear gradient, with constant 0.1% v/v TFA additive; 20 min run; 1 mL/min flow; ESI; positive ion mode; detection at 550 nm); HRMS (ESI) calcd for $C_{24}H_{11}D_{12}N_2O_3$ $[M+H]^+$ 399.2456, found 399.2454.

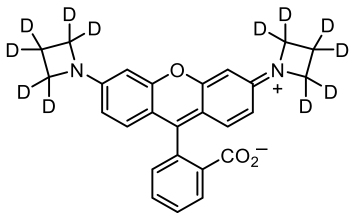

**JFX$_{549}$** ($\mathbf{4_D}$). A vial was charged with fluorescein ditriflate (**14**, 200 mg, 0.335 mmol), azetidine-2,2,3,3,4,4-$d_6$ hydrochloride[11] (167 mg, 1.68 mmol, 5 eq), $Pd_2dba_3$ (30.7 mg, 33.5 μmol, 0.1 eq), XPhos (48.0 mg, 0.101 mmol, 0.3 eq), and $Cs_2CO_3$ (874 mg, 2.68 mmol, 8 eq). The vial was sealed and evacuated/backfilled with nitrogen (3×). Dioxane (2 mL) was added, and the reaction was flushed again with nitrogen (3×). The reaction was then stirred at 100 °C for 2 h. It was subsequently cooled to room temperature, diluted with MeOH, deposited onto Celite, and concentrated to dryness. Purification by silica gel chromatography (0–10% MeOH (2 M $NH_3$)/$CH_2Cl_2$, linear gradient; dry load on Celite) afforded $\mathbf{4_D}$ (115 mg, 81%) as a dark red solid. $^1H$ NMR ($CDCl_3$, 400 MHz) δ 8.02 – 7.95 (m, 1H), 7.63 (td, $J$ = 7.4, 1.3 Hz, 1H), 7.57 (td, $J$ = 7.4, 1.1 Hz, 1H), 7.17 (dt, $J$ = 7.5, 1.0 Hz, 1H), 6.56 (d, $J$ = 8.6 Hz, 2H), 6.20 (d, $J$ = 2.3 Hz, 2H), 6.08 (dd, $J$ = 8.6, 2.3 Hz, 2H); $^{13}C$ NMR ($CDCl_3$, 101 MHz) δ 169.9 (C), 153.8 (C), 153.1 (C), 153.0 (C), 134.6 (CH), 129.4 (CH), 129.0 (CH), 127.8 (C), 125.0 (CH), 124.3 (CH), 107.9 (C), 107.8 (CH), 97.7 (CH); Analytical

HPLC: $t_R$ = 11.3 min, >99% purity (10–95% $MeCN/H_2O$, linear gradient, with constant 0.1% v/v TFA additive; 20 min run; 1 mL/min flow; ESI; positive ion mode; detection at 550 nm); HRMS (ESI) calcd for $C_{26}H_{11}D_{12}N_2O_3$ $[M+H]^+$ 423.2456, found 423.2454.

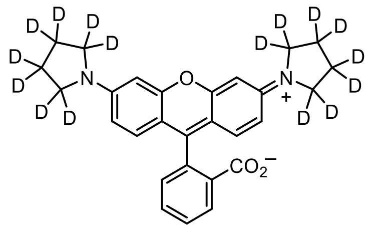

**JFX$_{554}$ (3$_D$).** A vial was charged with fluorescein ditriflate (**14**, 200 mg, 0.335 mmol), $Pd_2dba_3$ (30.7 mg, 33.5 µmol, 0.1 eq), XPhos (48.0 mg, 0.101 mmol, 0.3 eq), and $Cs_2CO_3$ (306 mg, 0.939 mmol, 2.8 eq). The vial was sealed and evacuated/backfilled with nitrogen (3×). Dioxane (2 mL) was added, and the reaction was flushed again with nitrogen (3×). Following the addition of pyrrolidine-2,2,3,3,4,4,5,5-$d_8$ (64 mg, 0.805 mmol, 2.4 eq), the reaction was stirred at 100 °C for 3 h. It was subsequently cooled to room temperature, diluted with MeOH, deposited onto Celite, and concentrated to dryness. Purification by silica gel chromatography (0–10% MeOH (2 M $NH_3$)/$CH_2Cl_2$, linear gradient; dry load on Celite) afforded **3$_D$** (116 mg, 76%) as a dark red-purple solid. $^1H$ NMR ($CD_3OD$, 400 MHz) δ 8.11 – 8.07 (m, 1H), 7.65 (td, *J* = 7.5, 1.6 Hz, 1H), 7.60 (td, *J* = 7.4, 1.6 Hz, 1H), 7.26 (d, *J* = 9.3 Hz, 2H), 7.25 – 7.22 (m, 1H), 6.85 (dd, *J* = 9.3, 2.4 Hz, 2H), 6.75 (d, *J* = 2.3 Hz, 2H); $^{13}C$ NMR ($CD_3OD$, 101 MHz) δ 163.4 (C), 158.9 (C), 156.1 (C), 141.8 (C), 133.9 (C), 132.9 (CH), 131.0 (CH), 130.7 (CH), 130.5 (CH), 130.4 (CH), 115.6 (CH), 115.1 (C), 97.5 (CH); Analytical HPLC: $t_R$ = 12.4 min, >99% purity (10–95% $MeCN/H_2O$, linear gradient, with constant 0.1% v/v TFA additive; 20 min run; 1 mL/min flow; ESI; positive ion mode; detection at 550 nm); HRMS (ESI) calcd for $C_{28}H_{11}D_{16}N_2O_3$ $[M+H]^+$ 455.3020, found 455.3018.

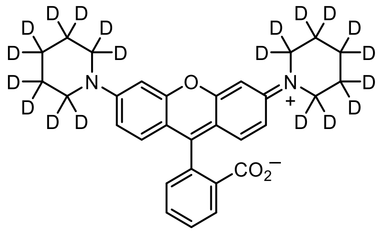

**2-(3,6-Bis(piperidin-1-yl-$d_{10}$)xanthylium-9-yl)benzoate (16$_D$)**. The title compound (96%, dark red-purple solid) was prepared from fluorescein ditriflate (**14**) and piperidine-2,2,3,3,4,4,5,5,6,6-$d_{10}$ according to the procedure described for **3$_D$**. $^1H$ NMR ($CD_3OD$, 400 MHz) δ 8.11 – 8.05 (m, 1H), 7.66 (td, *J* = 7.4, 1.8 Hz, 1H), 7.62 (td, *J* = 7.3, 1.7 Hz, 1H), 7.26 – 7.21 (m, 1H), 7.17 (d, *J* = 9.4 Hz, 2H), 7.06 (dd, *J* = 9.4, 2.6 Hz, 2H), 7.01 (d, *J* = 2.5 Hz, 2H); $^{13}C$ NMR ($CD_3OD$, 101 MHz) δ 158.9 (C), 157.6 (C), 152.1 (C), 139.7 (C), 136.7 (C), 132.6 (CH), 131.4 (CH), 130.8 (CH), 130.3 (CH), 129.7 (CH), 115.1 (CH), 114.5 (C), 98.6 (CH); Analytical HPLC: $t_R$ =12.7 min, >99% purity (10–95% $MeCN/H_2O$, linear gradient, with constant 0.1% v/v TFA additive; 20 min run; 1 mL/min flow; ESI; positive ion mode; detection at 550 nm); HRMS (ESI) calcd for $C_{30}H_{11}D_{20}N_2O_3$ $[M+H]^+$ 487.3585, found 487.3588.

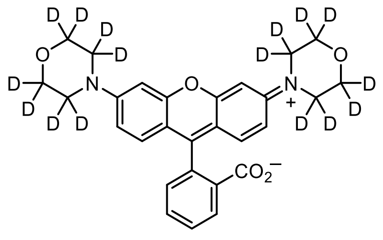

**2-(3,6-Bis(morpholino-*d*$_8$)xanthylium-9-yl)benzoate (17$_D$).** The title compound (88%, pink solid) was prepared from fluorescein ditriflate (**14**) and morpholine-2,2,3,3,5,5,6,6-*d*$_8$ according to the procedure described for **3$_D$**. $^1$H NMR ($CDCl_3$, 400 MHz) δ 8.04 – 7.98 (m, 1H), 7.64 (td, *J* = 7.4, 1.4 Hz, 1H), 7.59 (td, *J* = 7.4, 1.2 Hz, 1H), 7.18 – 7.13 (m, 1H), 6.67 (d, *J* = 2.5 Hz, 2H), 6.65 (d, *J* = 8.8 Hz, 2H), 6.57 (dd, *J* = 8.8, 2.5 Hz, 2H); $^{13}$C NMR ($CDCl_3$, 101 MHz) δ 169.8 (C), 153.4 (C), 153.1 (C), 152.8 (C), 134.9 (CH), 129.6 (CH), 128.9 (CH), 127.2 (C), 125.0 (CH), 124.1 (CH), 111.4 (CH), 109.9 (C), 101.8 (CH), 83.9 (C); Analytical HPLC: $t_R$ = 10.2 min, >99% purity (10–95% MeCN/$H_2O$, linear gradient, with constant 0.1% v/v TFA additive; 20 min run; 1 mL/min flow; ESI; positive ion mode; detection at 550 nm); HRMS (ESI) calcd for $C_{28}H_{11}D_{16}N_2O_5$ $[M+H]^+$ 487.2919, found 487.2926.

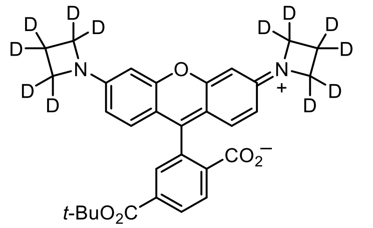

**6-*tert*-Butoxycarbonyl-JFX$_{549}$ (S2).** The title compound (72%, dark red-purple solid) was prepared from 6-*tert*-butoxycarbonylfluorescein ditriflate (**S1**) and azetidine-2,2,3,3,4,4-*d*$_6$ hydrochloride[11] according to the procedure described for **4$_D$**. $^1$H NMR ($CDCl_3$, 400 MHz) δ 8.19 (dd, *J* = 8.0, 1.3 Hz, 1H), 8.01 (dd, *J* = 8.0, 0.5 Hz, 1H), 7.77 – 7.70 (m, 1H), 6.53 (d, *J* = 8.6 Hz, 2H), 6.21 (d, *J* = 2.3 Hz, 2H), 6.09 (dd, *J* = 8.6, 2.3 Hz, 2H), 1.54 (s, 9H); $^{13}$C NMR ($CDCl_3$, 101 MHz) δ 169.1 (C), 164.5 (C), 153.8 (C), 153.3 (C), 152.9 (C), 138.1 (C), 130.8 (C), 130.6 (CH), 128.9 (CH), 125.3 (CH), 124.8 (CH), 107.8 (CH), 107.2 (C), 97.7 (CH), 86.5 (C), 82.5 (C), 28.2 ($CH_3$); Analytical HPLC: $t_R$ = 12.9 min, >99% purity (10–95% MeCN/$H_2O$, linear gradient, with constant 0.1% v/v TFA additive; 20 min run; 1 mL/min flow; ESI; positive ion mode; detection at 550 nm); HRMS (ESI) calcd for $C_{31}H_{19}D_{12}N_2O_5$ $[M+H]^+$ 523.2981, found 523.2982.

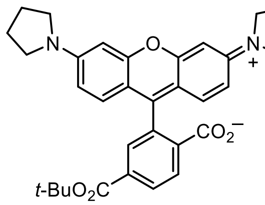

**6-*tert*-Butoxycarbonyl-Rh$_P$ (S3).** The title compound (59%, dark red-purple solid) was prepared from 6-*tert*-butoxycarbonylfluorescein ditriflate (**S1**) and pyrrolidine according to the procedure described for **3$_D$**. $^1$H NMR

($CD_3OD$, 400 MHz) δ 8.20 (dd, *J* = 8.1, 1.7 Hz, 1H), 8.11 (d, *J* = 8.1 Hz, 1H), 7.78 (d, *J* = 1.7 Hz, 1H), 7.22 (d, *J* = 9.4 Hz, 2H), 6.87 (dd, *J* = 9.4, 2.3 Hz, 2H), 6.77 (d, *J* = 2.3 Hz, 2H), 3.65 – 3.52 (m, 8H), 2.20 – 2.06 (m, 8H), 1.59 (s, 9H); $^{13}C$ NMR ($CD_3OD$, 101 MHz) δ 172.4 (C), 166.1 (C), 162.2 (C), 158.9 (C), 156.1 (C), 145.9 (C), 133.8 (C), 133.6 (C), 132.7 (CH), 131.5 (CH), 131.2 (CH), 131.0 (CH), 115.9 (CH), 115.1 (C), 97.6 (CH), 83.1 (C), 49.9 ($CH_2$), 28.4 ($CH_3$), 26.2 ($CH_2$); Analytical HPLC: $t_R$ = 13.7 min, >99% purity (10–95% MeCN/$H_2O$, linear gradient, with constant 0.1% v/v TFA additive; 20 min run; 1 mL/min flow; ESI; positive ion mode; detection at 550 nm); HRMS (ESI) calcd for $C_{33}H_{35}N_2O_5$ $[M+H]^+$ 539.2540, found 539.2544.

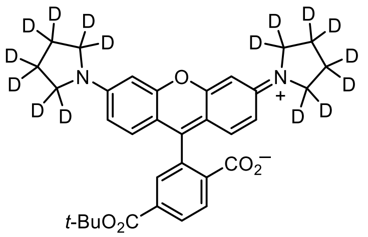


**6-*tert*-Butoxycarbonyl-JFX$_{554}$ (S4).** The title compound (58%, dark red-purple solid) was prepared from 6-*tert*-butoxycarbonylfluorescein ditriflate (**S1**) and pyrrolidine-2,2,3,3,4,4,5,5-$d_8$ according to the procedure described for **3$_D$**. $^1H$ NMR ($CD_3OD$, 400 MHz) δ 8.20 (dd, *J* = 8.2, 1.7 Hz, 1H), 8.11 (d, *J* = 8.1 Hz, 1H), 7.78 (d, *J* = 1.6 Hz, 1H), 7.21 (d, *J* = 9.3 Hz, 2H), 6.87 (dd, *J* = 9.3, 2.4 Hz, 2H), 6.76 (d, *J* = 2.3 Hz, 2H), 1.59 (s, 9H); $^{13}C$ NMR ($CD_3OD$, 101 MHz) δ 172.5 (C), 166.1 (C), 162.2 (C), 158.9 (C), 156.2 (C), 145.9 (C), 133.7 (C), 133.6 (C), 132.7 (CH), 131.5 (CH), 131.2 (CH), 131.0 (CH), 115.9 (CH), 115.0 (C), 97.6 (CH), 83.1 (C), 28.4 ($CH_3$); Analytical HPLC: $t_R$ = 13.6 min, >99% purity (10–95% MeCN/$H_2O$, linear gradient, with constant 0.1% v/v TFA additive; 20 min run; 1 mL/min flow; ESI; positive ion mode; detection at 550 nm); HRMS (ESI) calcd for $C_{33}H_{19}D_{16}N_2O_5$ $[M+H]^+$ 555.3545, found 555.3544.

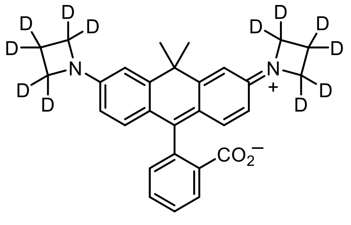


**JFX$_{608}$ (20$_D$).** The title compound (92%, pale blue solid) was prepared from carbofluorescein ditriflate (**S9**) and azetidine-2,2,3,3,4,4-$d_6$ according to the procedure described for **3$_D$**. $^1H$ NMR ($CDCl_3$, 400 MHz) δ 8.00 – 7.95 (m, 1H), 7.58 (td, *J* = 7.4, 1.4 Hz, 1H), 7.53 (td, *J* = 7.4, 1.2 Hz, 1H), 7.08 – 7.04 (m, 1H), 6.58 (d, *J* = 2.4 Hz, 2H), 6.55 (d, *J* = 8.6 Hz, 2H), 6.20 (dd, *J* = 8.6, 2.4 Hz, 2H), 1.82 (s, 3H), 1.72 (s, 3H); $^{13}C$ NMR ($CDCl_3$, 101 MHz) δ 170.9 (C), 155.5 (C), 152.5 (C), 146.9 (C), 134.5 (CH), 129.0 (CH), 128.9 (CH), 127.4 (C), 125.0 (CH), 124.1 (CH), 120.6 (C), 110.4 (CH), 107.9 (CH), 88.5 (C), 38.5 (C), 35.7 ($CH_3$), 32.3 ($CH_3$); Analytical HPLC: $t_R$ = 11.9 min, >99% purity (10–95% MeCN/$H_2O$, linear gradient, with constant 0.1% v/v TFA additive; 20 min run; 1 mL/min flow; ESI; positive ion mode; detection at 600 nm); HRMS (ESI) calcd for $C_{29}H_{17}D_{12}N_2O_2$ $[M+H]^+$ 449.2977, found 449.2980.

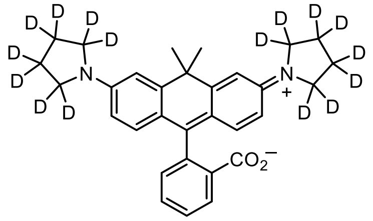


**JFX$_{612}$ (21$_D$).** The title compound (91%, pale blue solid) was prepared from carbofluorescein ditriflate (**S9**) and pyrrolidine-2,2,3,3,4,4,5,5-$d_8$ according to the procedure described for **3$_D$**. $^1$H NMR ($CDCl_3$, 400 MHz) δ 8.00 – 7.96 (m, 1H), 7.57 (td, $J$ = 7.3, 1.5 Hz, 1H), 7.52 (td, $J$ = 7.4, 1.3 Hz, 1H), 7.08 – 7.04 (m, 1H), 6.72 (d, $J$ = 2.5 Hz, 2H), 6.58 (d, $J$ = 8.7 Hz, 2H), 6.35 (dd, $J$ = 8.7, 2.5 Hz, 2H), 1.88 (s, 3H), 1.77 (s, 3H); $^{13}$C NMR ($CDCl_3$, 101 MHz) δ 171.1 (C), 155.7 (C), 148.3 (C), 147.2 (C), 134.4 (CH), 129.2 (CH), 128.7 (CH), 127.6 (C), 124.9 (CH), 124.1 (CH), 118.8 (C), 111.0 (CH), 108.3 (CH), 38.6 (C), 36.0 ($CH_3$), 32.5 ($CH_3$); Analytical HPLC: $t_R$ = 9.8 min, >99% purity (30–95% MeCN/$H_2O$, linear gradient, with constant 0.1% v/v TFA additive; 20 min run; 1 mL/min flow; ESI; positive ion mode; detection at 600 nm); HRMS (ESI) calcd for $C_{31}H_{17}D_{16}N_2O_2$ [M+H]$^+$ 481.3541, found 481.3545.

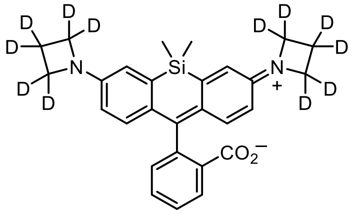


**JFX$_{646}$ (22$_D$)**. The title compound (89%, light blue solid) was prepared from Si-fluorescein ditriflate (**S10**) and azetidine-2,2,3,3,4,4-$d_6$ according to the procedure described for **3$_D$**. $^1$H NMR ($CDCl_3$, 400 MHz) δ 7.96 (dt, $J$ = 7.6, 1.0 Hz, 1H), 7.64 (td, $J$ = 7.5, 1.1 Hz, 1H), 7.54 (td, $J$ = 7.5, 0.8 Hz, 1H), 7.31 (dt, $J$ = 7.7, 0.9 Hz, 1H), 6.75 (d, $J$ = 8.7 Hz, 2H), 6.67 (d, $J$ = 2.6 Hz, 2H), 6.25 (dd, $J$ = 8.6, 2.7 Hz, 2H), 0.61 (s, 3H), 0.59 (s, 3H); $^{13}$C NMR ($CDCl_3$, 101 MHz) δ 170.8 (C), 154.3 (C), 151.1 (C), 137.1 (C), 133.7 (CH), 132.9 (C), 128.8 (CH), 128.0 (CH), 127.2 (C), 125.8 (CH), 124.8 (CH), 115.7 (CH), 112.3 (CH), 92.1 (C), 0.5 ($CH_3$), -1.5 ($CH_3$); Analytical HPLC: $t_R$ = 12.5 min, >99% purity (10–95% MeCN/$H_2O$, linear gradient, with constant 0.1% v/v TFA additive; 20 min run; 1 mL/min flow; ESI; positive ion mode; detection at 650 nm); HRMS (ESI) calcd for $C_{28}H_{17}D_{12}N_2O_2Si$ [M+H]$^+$ 465.2746, found 465.2749.

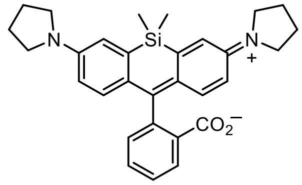


**SiRh$_P$ (23).** The title compound (96%, pale blue solid) was prepared from Si-fluorescein ditriflate (**S10**) and pyrrolidine according to the procedure described for **3$_D$**. $^1$H NMR ($CDCl_3$, 400 MHz) δ 7.95 (dt, $J$ = 7.6, 0.9 Hz, 1H), 7.62 (td, $J$ = 7.5, 1.1 Hz, 1H), 7.52 (td, $J$ = 7.5, 0.9 Hz, 1H), 7.31 – 7.27 (m, 1H), 6.79 (d, $J$ = 2.7 Hz, 2H), 6.76 (d, $J$ = 8.8 Hz, 2H), 6.38 (dd, $J$ = 8.8, 2.8 Hz, 2H), 3.34 – 3.24 (m, 8H), 2.05 – 1.93 (m, 8H), 0.63 (s, 3H), 0.60 (s, 3H); $^{13}$C NMR ($CDCl_3$, 101 MHz) δ 171.0 (C), 154.9 (C), 146.8 (C), 137.2 (C), 133.7 (CH), 131.0 (C), 128.6 (CH), 128.4

(CH), 127.2 (C), 125.6 (CH), 124.7 (CH), 115.9 (CH), 112.7 (CH), 92.5 (C), 47.6 ($CH_2$), 25.6 ($CH_2$), 0.6 ($CH_3$), -1.3 ($CH_3$); Analytical HPLC: >99% purity (30–95% MeCN/$H_2O$, linear gradient, with constant 0.1% v/v TFA additive; 20 min run; 1 mL/min flow; ESI; positive ion mode; detection at 650 nm); HRMS (ESI) calcd for $C_{30}H_{33}N_2O_2Si$ $[M+H]^+$ 481.2306, found 481.2317.

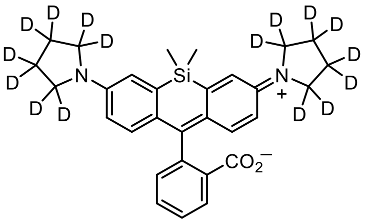


**JFX$_{650}$** (**23$_D$**). The title compound (90%, off-white solid) was prepared from Si-fluorescein ditriflate (**S10**) and pyrrolidine-2,2,3,3,4,4,5,5-*d*$_8$ according to the procedure described for **3$_D$**. $^1$H NMR ($CDCl_3$, 400 MHz) δ 7.95 (dt, *J* = 7.7, 1.0 Hz, 1H), 7.61 (td, *J* = 7.5, 1.1 Hz, 1H), 7.52 (td, *J* = 7.5, 0.9 Hz, 1H), 7.28 (dt, *J* = 7.8, 1.0 Hz, 1H), 6.79 (d, *J* = 2.8 Hz, 2H), 6.76 (d, *J* = 8.8 Hz, 2H), 6.38 (dd, *J* = 8.8, 2.8 Hz, 2H), 0.63 (s, 3H), 0.59 (s, 3H); $^{13}$C NMR ($CDCl_3$, 101 MHz) δ 171.0 (C), 154.9 (C), 147.0 (C), 137.2 (C), 133.7 (CH), 131.0 (C), 128.6 (CH), 128.5 (CH), 127.2 (C), 125.7 (CH), 124.7 (CH), 115.9 (CH), 112.8 (CH), 92.5 (C), 0.6 ($CH_3$), -1.3 ($CH_3$); Analytical HPLC: $t_R$ = 13.0 min, >99% purity (10–95% MeCN/$H_2O$, linear gradient, with constant 0.1% v/v TFA additive; 20 min run; 1 mL/min flow; ESI; positive ion mode; detection at 650 nm); HRMS (ESI) calcd for $C_{30}H_{17}D_{16}N_2O_2Si$ $[M+H]^+$ 497.3310, found 497.3312.

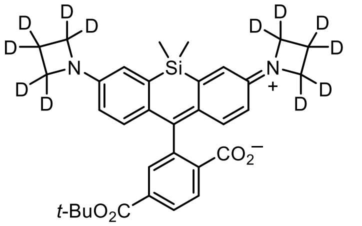


**6-*tert*-Butoxycarbonyl-JFX$_{646}$ (S14).** The title compound (93%, off-white solid) was prepared from 6-*tert*-butoxycarbonyl-Si-fluorescein ditriflate (**S13**) and azetidine-2,2,3,3,4,4-*d*$_6$ according to the procedure described for **3$_D$**. $^1$H NMR ($CDCl_3$, 400 MHz) δ 8.11 (dd, *J* = 8.0, 1.3 Hz, 1H), 7.95 (dd, *J* = 8.0, 0.6 Hz, 1H), 7.83 – 7.80 (m, 1H), 6.82 (d, *J* = 8.7 Hz, 2H), 6.66 (d, *J* = 2.7 Hz, 2H), 6.29 (dd, *J* = 8.7, 2.7 Hz, 2H), 1.54 (s, 9H), 0.64 (s, 3H), 0.58 (s, 3H); $^{13}$C NMR ($CDCl_3$, 101 MHz) δ 170.3 (C), 164.5 (C), 155.4 (C), 151.1 (C), 137.2 (C), 136.2 (C), 132.4 (C), 129.9 (CH), 129.2 (C), 127.7 (CH), 125.6 (CH), 125.2 (CH), 115.7 (CH), 112.6 (CH), 91.9 (C), 82.3 (C), 28.2 ($CH_3$), 0.2 ($CH_3$), -0.7 ($CH_3$); Analytical HPLC: $t_R$ = 14.1 min, 98.8% purity (10–95% MeCN/$H_2O$, linear gradient, with constant 0.1% v/v TFA additive; 20 min run; 1 mL/min flow; ESI; positive ion mode; detection at 650 nm); HRMS (ESI) calcd for $C_{33}H_{25}D_{12}N_2O_4Si$ $[M+H]^+$ 565.3270, found 565.3277.

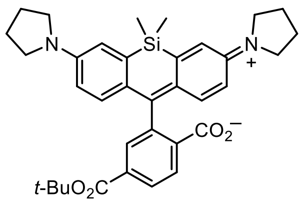

**6-*tert*-Butoxycarbonyl-SiRh$_P$ (S15).** The title compound (81%, off-white solid) was prepared from 6-*tert*-butoxycarbonyl-Si-fluorescein ditriflate (**S13**) and pyrrolidine according to the procedure described for **3$_D$**. $^1$H NMR ($CDCl_3$, 400 MHz) δ 8.10 (dd, *J* = 8.0, 1.3 Hz, 1H), 7.96 (dd, *J* = 8.0, 0.5 Hz, 1H), 7.83 – 7.79 (m, 1H), 6.84 (d, *J* = 8.8 Hz, 2H), 6.79 (d, *J* = 2.7 Hz, 2H), 6.44 (dd, *J* = 8.8, 2.8 Hz, 2H), 3.35 – 3.25 (m, 8H), 2.04 – 1.95 (m, 8H), 1.53 (s, 9H), 0.67 (s, 3H), 0.60 (s, 3H); $^{13}$C NMR ($CDCl_3$, 101 MHz) δ 170.6 (C), 164.5 (C), 155.9 (C), 146.8 (C), 137.2 (C), 136.3 (C), 130.5 (C), 129.8 (CH), 129.3 (C), 128.1 (CH), 125.5 (CH), 125.1 (CH), 115.9 (CH), 113.2 (CH), 92.2 (C), 82.2 (C), 47.6 ($CH_2$), 28.2 ($CH_3$), 25.6 ($CH_2$), 0.2 ($CH_3$), -0.5 ($CH_3$); Analytical HPLC: $t_R$ = 14.6 min, >99% purity (10–95% MeCN/$H_2O$, linear gradient, with constant 0.1% v/v TFA additive; 20 min run; 1 mL/min flow; ESI; positive ion mode; detection at 650 nm); HRMS (ESI) calcd for $C_{35}H_{41}N_2O_4Si$ $[M+H]^+$ 581.2830, found 581.2839.

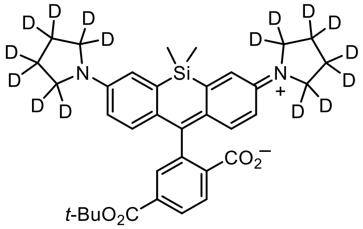

**6-*tert*-Butoxycarbonyl-JFX$_{650}$ (S16).** The title compound (87%, off-white solid) was prepared from 6-*tert*-butoxycarbonyl-Si-fluorescein ditriflate (**S13**) and pyrrolidine-2,2,3,3,4,4,5,5-$d_8$ according to the procedure described for **3$_D$**. $^1$H NMR ($CDCl_3$, 400 MHz) δ 8.10 (dd, *J* = 8.0, 1.3 Hz, 1H), 7.95 (dd, *J* = 8.0, 0.6 Hz, 1H), 7.83 – 7.78 (m, 1H), 6.84 (d, *J* = 8.8 Hz, 2H), 6.79 (d, *J* = 2.7 Hz, 2H), 6.43 (dd, *J* = 8.8, 2.8 Hz, 2H), 1.53 (s, 9H), 0.67 (s, 3H), 0.59 (s, 3H); $^{13}$C NMR ($CDCl_3$, 101 MHz) δ 170.6 (C), 164.5 (C), 155.9 (C), 146.9 (C), 137.2 (C), 136.3 (C), 130.5 (C), 129.8 (CH), 129.3 (C), 128.1 (CH), 125.5 (CH), 125.1 (CH), 115.9 (CH), 113.1 (CH), 92.3 (C), 82.2 (C), 28.2 ($CH_3$), 0.2 ($CH_3$), -0.5 ($CH_3$); Analytical HPLC: $t_R$ = 14.4 min, >99% purity (10–95% MeCN/$H_2O$, linear gradient, with constant 0.1% v/v TFA additive; 20 min run; 1 mL/min flow; ESI; positive ion mode; detection at 650 nm); HRMS (ESI) calcd for $C_{35}H_{25}D_{16}N_2O_4Si$ $[M+H]^+$ 597.3834, found 597.3835.

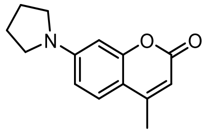

**4-Methyl-7-(pyrrolidin-1-yl)-2*H*-chromen-2-one (24).** A vial was charged with 4-methylumbelliferone triflate (**S11**; 250 mg, 0.811 mmol), RuPhos-G3-palladacycle (33.9 mg, 40.6 µmol, 0.05 eq), RuPhos (18.9 mg, 40.6 µmol, 0.05 eq), and $K_2CO_3$ (157 mg, 1.14 mmol, 1.4 eq). The vial was sealed and evacuated/backfilled with nitrogen (3×). Dioxane (4 mL) was added, and the reaction was flushed again with nitrogen (3×). Following the addition of pyrrolidine (81.2 $\mu$L, 0.973 mmol, 1.2 eq), the reaction was stirred at 80 °C for 6 h. It was subsequently cooled to room temperature, filtered through Celite with $CH_2Cl_2$, and concentrated *in vacuo*. Purification by silica gel chromatography (0–50% EtOAc/hexanes, linear gradient) afforded 171 mg (92%) of **24** as a yellow solid. $^1$H NMR ($CDCl_3$, 400 MHz) δ 7.39 (d, $J$ = 8.8 Hz, 1H), 6.48 (dd, $J$ = 8.8, 2.4 Hz, 1H), 6.38 (d, $J$ = 2.4 Hz, 1H), 5.94 (q, $J$ = 1.1 Hz, 1H), 3.40 – 3.30 (m, 4H), 2.34 (d, $J$ = 1.1 Hz, 3H), 2.10 – 1.99 (m, 4H); $^{13}$C NMR ($CDCl_3$, 101 MHz) δ 162.3 (C), 155.9 (C), 153.2 (C), 150.5 (C), 125.5 (CH), 109.4 (C), 109.1 (CH), 108.8 (CH), 98.0 (CH), 47.8 ($CH_2$), 25.6 ($CH_2$), 18.6 ($CH_3$); Analytical HPLC: $t_R$ = 13.2 min, >99% purity (10–95% MeCN/$H_2O$, linear gradient, with constant 0.1% v/v TFA additive; 20 min run; 1 mL/min flow; ESI; positive ion mode; detection at 375 nm); HRMS (ESI) calcd for $C_{14}H_{16}NO_2$ $[M+H]^+$ 230.1176, found 230.1180.

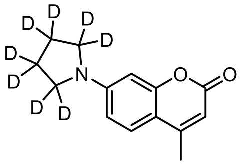

**4-Methyl-7-(pyrrolidin-1-yl-*d*$_8$)-2*H*-chromen-2-one (24$_D$).** The title compound (86%, yellow solid) was prepared from 4-methylumbelliferone triflate (**S11**) and pyrrolidine-2,2,3,3,4,4,5,5-*d*$_8$ according to the procedure described for **24**. $^1$H NMR ($CDCl_3$, 400 MHz) δ 7.38 (d, $J$ = 8.8 Hz, 1H), 6.48 (dd, $J$ = 8.8, 2.4 Hz, 1H), 6.38 (d, $J$ = 2.4 Hz, 1H), 5.94 (q, $J$ = 1.1 Hz, 1H), 2.34 (d, $J$ = 1.1 Hz, 3H); $^{13}$C NMR ($CDCl_3$, 101 MHz) δ 162.3 (C), 155.9 (C), 153.2 (C), 150.6 (C), 125.5 (CH), 109.3 (C), 109.1 (CH), 108.7 (CH), 98.0 (CH), 18.6 ($CH_3$); Analytical HPLC: $t_R$ = 13.1 min, >99% purity (10–95% MeCN/$H_2O$, linear gradient, with constant 0.1% v/v TFA additive; 20 min run; 1 mL/min flow; ESI; positive ion mode; detection at 375 nm); HRMS (ESI) calcd for $C_{14}H_8D_8NO_2$ $[M+H]^+$ 238.1678, found 238.1682.

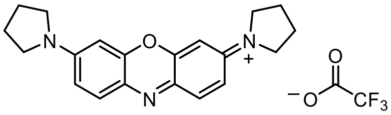

**3,7-Di(pyrrolidin-1-yl)phenoxazin-5-ium trifluoroacetate (25).** A vial was charged with 10-acetyl-10*H*-phenoxazine-3,7-diyl bis(trifluoromethanesulfonate) (**S12**; 250 mg, 0.480 mmol), $Pd_2dba_3$ (43.9 mg, 48.0 µmol, 0.1 eq), XPhos (68.6 mg, 0.144 mmol, 0.3 eq), and $Cs_2CO_3$ (437 mg, 1.34 mmol, 2.8 eq). The vial was sealed and evacuated/backfilled with nitrogen (3×). Dioxane (2.5 mL) was added, and the reaction was flushed again with nitrogen (3×). Following the addition of pyrrolidine (96.1 µL, 1.15 mmol, 2.4 eq), the reaction was stirred at 80 °C for 4 h. It was then cooled to room temperature, filtered through Celite with $CH_2Cl_2$, and concentrated *in vacuo*. Purification by silica gel chromatography (0–40% EtOAc/toluene, linear gradient) afforded the *N*-acetyl leuco-dye (112 mg, 64%) as an off-white solid.

The intermediate leuco-dye (112 mg, 0.308 mmol) was taken up in a mixture of $CH_2Cl_2$ (9 mL) and water (1 mL). DDQ (105 mg, 0.462 mmol, 1.5 eq) was added, and the reaction was stirred at room temperature for 2 h. The crude reaction mixture was then deposited onto Celite and concentrated to dryness. Silica gel chromatography (0–20% MeOH/$CH_2Cl_2$, linear gradient, with constant 1% v/v AcOH additive; dry load with Celite) followed by reverse phase

HPLC (10–50% MeCN/$H_2O$, linear gradient, with constant 0.1% v/v TFA additive) afforded 127 mg (95%) of **25** as a dark blue solid. $^1$H NMR ($CD_3OD$, 400 MHz) δ 7.77 (d, *J* = 9.5 Hz, 2H), 7.25 (dd, *J* = 9.4, 2.5 Hz, 2H), 6.81 (d, *J* = 2.5 Hz, 2H), 3.82 – 3.66 (m, 8H), 2.22 – 2.13 (m, 8H); $^{13}$C NMR ($CD_3OD$, 101 MHz) δ 156.6 (C), 150.4 (C), 135.5 (C), 135.3 (CH), 119.4 (CH), 98.0 (CH), 50.9 (bs, $CH_2$), 26.2 (bs, $CH_2$); Analytical HPLC: $t_R$ = 10.4 min, >99% purity (10–95% MeCN/$H_2O$, linear gradient, with constant 0.1% v/v TFA additive; 20 min run; 1 mL/min flow; ESI; positive ion mode; detection at 650 nm); HRMS (ESI) calcd for $C_{20}H_{22}N_3O$ $[M]^+$ 320.1757, found 320.1763.

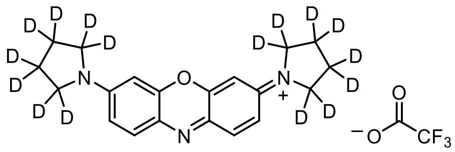

**3,7-Bis(pyrrolidin-1-yl-*d*$_8$)phenoxazin-5-ium trifluoroacetate (25$_D$).** The title compound (34% 2-step yield, dark blue solid) was prepared from 10-acetyl-10*H*-phenoxazine-3,7-diyl bis(trifluoromethanesulfonate)[3] (**S12**) and pyrrolidine-2,2,3,3,4,4,5,5-*d*$_8$ according to the procedure described for **25**. $^1$H NMR ($CD_3OD$, 400 MHz) δ 7.77 (d, *J* = 9.4 Hz, 2H), 7.25 (dd, *J* = 9.5, 2.5 Hz, 2H), 6.80 (d, *J* = 2.5 Hz, 2H); $^{13}$C NMR ($CD_3OD$, 101 MHz) δ 156.7 (C), 150.4 (C), 135.5 (C), 135.3 (CH), 119.4 (CH), 97.9 (CH); Analytical HPLC: $t_R$ = 10.3 min, >99% purity (10–95% MeCN/$H_2O$, linear gradient, with constant 0.1% v/v TFA additive; 20 min run; 1 mL/min flow; ESI; positive ion mode; detection at 650 nm); HRMS (ESI) calcd for $C_{20}H_6D_{16}N_3O$ $[M]^+$ 336.2762, found 336.2765.

### *t*-Butyl Ester Deprotection

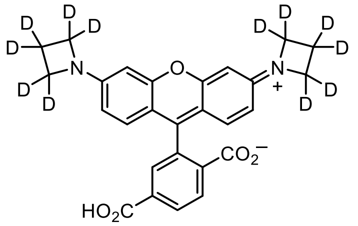


**6-Carboxy-JFX$_{549}$ (S5).** 6-*tert*-Butoxycarbonyl-JFX$_{549}$ (**S2**; 102 mg, 0.195 mmol) was taken up in $CH_2Cl_2$ (2.5 mL), and trifluoroacetic acid (0.5 mL) was added. The reaction was stirred at room temperature for 6 h. Toluene (3 mL) was added; the reaction mixture was concentrated to dryness and then azeotroped with MeOH three times to provide **S5** as a red-brown solid (109 mg, 96%, TFA salt). Analytical HPLC and NMR indicated that the material was >95% pure and did not require further purification prior to amide coupling. $^1$H NMR ($CD_3OD$, 400 MHz) δ 8.40 (d, *J* = 8.1 Hz, 1H), 8.37 (dd, *J* = 8.2, 1.5 Hz, 1H), 7.95 – 7.93 (m, 1H), 7.06 (d, *J* = 9.2 Hz, 2H), 6.60 (dd, *J* = 9.2, 2.2 Hz, 2H), 6.54 (d, *J* = 2.2 Hz, 2H); $^{13}$C NMR ($CD_3OD$, 101 MHz) δ 167.7 (C), 167.3 (C), 160.0 (C), 158.8 (C), 158.2 (C), 136.0 (C), 135.9 (C), 135.5 (C), 132.8 (CH), 132.3 (CH), 132.2 (CH), 114.8 (C), 113.6 (CH), 95.2 (CH); Analytical HPLC: $t_R$ = 9.9 min, >99% purity (10–95% MeCN/$H_2O$, linear gradient, with constant 0.1% v/v TFA additive; 20 min run; 1 mL/min flow; ESI; positive ion mode; detection at 550 nm); HRMS (ESI) calcd for $C_{27}H_{11}D_{12}N_2O_5$ [M+H]$^+$ 467.2355, found 467.2354.

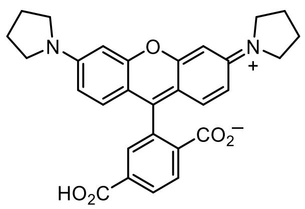


**6-Carboxy-Rh$_P$ (S6).** The title compound (99%, dark red-purple solid, TFA salt) was prepared from 6-*tert*-butoxycarbonyl-Rh$_P$ (**S3**) according to the procedure described for **S5**. $^1$H NMR ($CD_3OD$, 400 MHz) δ 8.42 (d, *J* = 8.2 Hz, 1H), 8.39 (dd, *J* = 8.2, 1.5 Hz, 1H), 7.99 – 7.96 (m, 1H), 7.11 (d, *J* = 9.4 Hz, 2H), 6.92 (dd, *J* = 9.4, 2.3 Hz, 2H), 6.84 (d, *J* = 2.3 Hz, 2H), 3.68 – 3.56 (m, 8H), 2.21 – 2.07 (m, 8H); $^{13}$C NMR ($CD_3OD$, 101 MHz) δ 167.7 (C), 167.4 (C), 160.0 (C), 158.9 (C), 156.2 (C), 136.1 (C), 135.9 (C), 135.5 (C), 132.8 (CH), 132.31 (CH), 132.28 (CH), 132.0 (CH), 116.4 (CH), 114.9 (C), 97.8 (CH), 50.0 ($CH_2$), 26.2 ($CH_2$); Analytical HPLC: $t_R$ = 10.8 min, >99% purity (5 µL injection; 10–95% MeCN/$H_2O$, linear gradient, with constant 0.1% v/v TFA additive; 20 min run; 1 mL/min flow; ESI; positive ion mode; detection at 550 nm); HRMS (ESI) calcd for $C_{29}H_{27}N_2O_5$ [M+H]$^+$ 483.1914, found 483.1919.

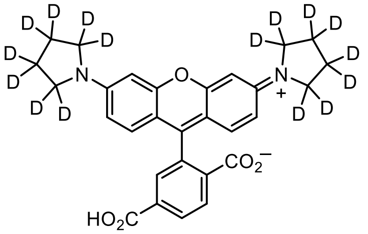

**6-Carboxy-JFX$_{554}$ (S7).** The title compound (97%, dark red-purple solid, TFA salt) was prepared from 6-*tert*-butoxycarbonyl-JFX$_{554}$ (**S4**) according to the procedure described for **S5**. $^1$H NMR ($CD_3OD$, 400 MHz) δ 8.41 (d, *J* = 8.2 Hz, 1H), 8.38 (dd, *J* = 8.2, 1.5 Hz, 1H), 7.99 – 7.96 (m, 1H), 7.11 (d, *J* = 9.4 Hz, 2H), 6.91 (dd, *J* = 9.3, 2.3 Hz, 2H), 6.83 (d, *J* = 2.3 Hz, 2H); $^{13}$C NMR ($CD_3OD$, 101 MHz) δ 167.7 (C), 167.4 (C), 159.9 (C), 158.9 (C), 156.4 (C), 136.1 (C), 135.9 (C), 135.5 (C), 132.8 (CH), 132.32 (CH), 132.27 (CH), 132.0 (CH), 116.4 (CH), 114.8 (C), 97.8 (CH); Analytical HPLC: $t_R$ = 10.8 min, >99% purity (10–95% MeCN/$H_2O$, linear gradient, with constant 0.1% v/v TFA additive; 20 min run; 1 mL/min flow; ESI; positive ion mode; detection at 550 nm); HRMS (ESI) calcd for $C_{29}H_{11}D_{16}N_2O_5$ $[M+H]^+$ 499.2919, found 499.2922.

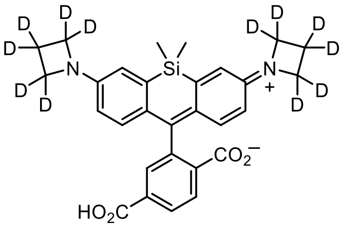

**6-Carboxy-JFX$_{646}$ (S17).** The title compound (98%, green solid, TFA salt) was prepared from 6-*tert*-butoxycarbonyl-JFX$_{646}$ (**S14**) according to the procedure described for **S5**. $^1$H NMR ($CD_3OD$, 400 MHz) δ 8.29 – 8.25 (m, 2H), 7.80 (t, *J* = 1.0 Hz, 1H), 6.90 (d, *J* = 2.6 Hz, 2H), 6.87 (d, *J* = 9.2 Hz, 2H), 6.33 (dd, *J* = 9.2, 2.6 Hz, 2H), 0.60 (s, 3H), 0.53 (s, 3H); Analytical HPLC: $t_R$ = 10.9 min, 97.8% purity (10–95% MeCN/$H_2O$, linear gradient, with constant 0.1% v/v TFA additive; 20 min run; 1 mL/min flow; ESI; positive ion mode; detection at 650 nm); HRMS (ESI) calcd for $C_{29}H_{17}D_{12}N_2O_4Si$ $[M+H]^+$ 509.2644, found 509.2648.

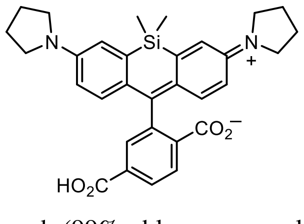

**6-Carboxy-SiRh$_P$ (S18).** The title compound (99%, blue-green solid, TFA salt) was prepared from 6-*tert*-butoxycarbonyl-SiRh$_P$ (**S15**) according to the procedure described for **S5**. $^1$H NMR ($CD_3OD$, 400 MHz) δ 8.34 (d, *J* = 8.2 Hz, 1H), 8.29 (dd, *J* = 8.2, 1.7 Hz, 1H), 7.82 (d, *J* = 1.6 Hz, 1H), 7.20 (d, *J* = 2.7 Hz, 2H), 6.94 (d, *J* = 9.4 Hz, 2H), 6.61 (dd, *J* = 9.5, 2.7 Hz, 2H), 3.73 – 3.58 (m, 8H), 2.16 – 2.05 (m, 8H), 0.64 (s, 3H), 0.57 (s, 3H); $^{13}$C NMR ($CD_3OD$, 101 MHz) δ 170.3 (C), 168.0 (C), 167.5 (C), 152.9 (C), 149.3 (C), 142.5 (C), 141.6 (CH), 136.1 (C), 135.2 (C), 132.5 (CH), 132.4 (CH), 131.0 (CH), 129.1 (C), 122.6 (CH), 115.7 (CH), 49.9 ($CH_2$), 26.1 ($CH_2$), -0.8 ($CH_3$), -

1.8 ($CH_3$); Analytical HPLC: $t_R$ = 11.5 min, >99% purity (10–95% $MeCN/H_2O$, linear gradient, with constant 0.1% v/v TFA additive; 20 min run; 1 mL/min flow; ESI; positive ion mode; detection at 650 nm); HRMS (ESI) calcd for $C_{31}H_{33}N_2O_4Si$ $[M+H]^+$ 525.2204, found 525.2214.

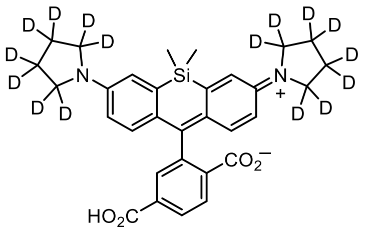

**6-Carboxy-JFX$_{650}$ (S19).** The title compound (~99%, blue-green solid, TFA salt) was prepared from 6-*tert*-butoxycarbonyl-JFX$_{650}$ (**S16**) according to the procedure described for **S5**. $^1$H NMR ($CD_3OD$, 400 MHz) δ 8.34 (d, *J* = 8.1 Hz, 1H), 8.29 (dd, *J* = 8.2, 1.6 Hz, 1H), 7.83 – 7.81 (m, 1H), 7.19 (d, *J* = 2.7 Hz, 2H), 6.94 (d, *J* = 9.5 Hz, 2H), 6.60 (dd, *J* = 9.5, 2.7 Hz, 2H), 0.64 (s, 3H), 0.57 (s, 3H); $^{13}$C NMR ($CD_3OD$, 101 MHz) δ 168.0 (C), 167.5 (C), 153.0 (C), 149.3 (C), 142.5 (C), 141.6 (CH), 136.1 (C), 135.2 (C), 132.5 (CH), 132.4 (CH), 131.0 (CH), 129.0 (C), 122.6 (CH), 115.7 (CH), -0.8 ($CH_3$), -1.8 ($CH_3$); Analytical HPLC: $t_R$ = 11.5 min, >99% purity (10–95% $MeCN/H_2O$, linear gradient, with constant 0.1% v/v TFA additive; 20 min run; 1 mL/min flow; ESI; positive ion mode; detection at 650 nm); HRMS (ESI) calcd for $C_{31}H_{17}D_{16}N_2O_4Si$ $[M+H]^+$ 541.3208, found 541.3213.

### *N*-Hydroxysuccimidyl Ester Formation

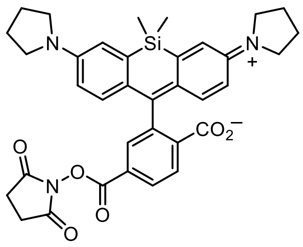

**SiRh$_P$-NHS (S20).** 6-Carboxy-SiRh$_P$ (**S18**; 244 mg, 0.382 mmol, TFA salt) was combined with DSC (235 mg, 0.917 mmol, 2.4 eq) in DMF (5 mL). After adding $Et_3N$ (319 µL, 2.29 mmol, 6 eq) and DMAP (4.7 mg, 38.2 µmol, 0.1 eq), the reaction was stirred at room temperature for 30 min. It was subsequently diluted with 10% w/v citric acid and extracted with EtOAc (2×). The combined organic extracts were washed with brine, dried over anhydrous $MgSO_4$, filtered, and concentrated *in vacuo*. Silica gel chromatography (0–100% EtOAc/toluene, linear gradient) yielded 206 mg (87%) of **S20** as a blue-green solid. $^1H$ NMR ($CDCl_3$, 400 MHz) δ 8.27 (dd, *J* = 8.0, 1.4 Hz, 1H), 8.07 (dd, *J* = 8.0, 0.8 Hz, 1H), 8.00 (dd, *J* = 1.3, 0.8 Hz, 1H), 6.79 (d, *J* = 2.7 Hz, 2H), 6.74 (d, *J* = 8.8 Hz, 2H), 6.43 (dd, *J* = 8.9, 2.7 Hz, 2H), 3.35 – 3.25 (m, 8H), 2.88 (s, 4H), 2.04 – 1.94 (m, 8H), 0.64 (s, 3H), 0.58 (s, 3H); $^{13}C$ NMR ($CDCl_3$, 101 MHz) δ 169.6 (C), 168.9 (C), 161.2 (C), 155.8 (C), 147.0 (C), 136.9 (C), 132.0 (C), 130.6 (CH), 130.1 (C), 129.7 (C), 128.3 (CH), 126.8 (CH), 126.2 (CH), 116.1 (CH), 113.2 (CH), 92.9 (C), 47.6 ($CH_2$), 25.8 ($CH_2$), 25.6 ($CH_2$), 0.4 ($CH_3$), -1.1 ($CH_3$); Analytical HPLC: $t_R$ = 12.2 min, 98.6% purity (10–95% MeCN/$H_2O$, linear gradient, with constant 0.1% v/v TFA additive; 20 min run; 1 mL/min flow; ESI; positive ion mode; detection at 650 nm); HRMS (ESI) calcd for $C_{35}H_{36}N_3O_6Si$ $[M+H]^+$ 622.2368, found 622.2369.

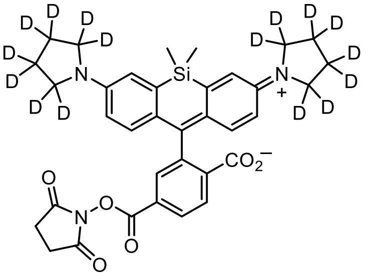

**JFX$_{650}$-NHS (S21).** The title compound (92%, blue-green solid) was prepared from 6-carboxy-JFX$_{650}$ (**S19**) according to the procedure described for **S20**. $^1H$ NMR ($CDCl_3$, 400 MHz) δ 8.27 (dd, *J* = 8.0, 1.4 Hz, 1H), 8.07 (dd, *J* = 8.0, 0.6 Hz, 1H), 8.01 (dd, *J* = 1.4, 0.8 Hz, 1H), 6.78 (d, *J* = 2.8 Hz, 2H), 6.73 (d, *J* = 8.8 Hz, 2H), 6.42 (dd, *J* = 8.8, 2.8 Hz, 2H), 2.89 (s, 4H), 0.64 (s, 3H), 0.58 (s, 3H); Analytical HPLC: $t_R$ = 12.1 min, 98.8% purity (10–95% MeCN/$H_2O$, linear gradient, with constant 0.1% v/v TFA additive; 20 min run; 1 mL/min flow; ESI; positive ion mode; detection at 650 nm); HRMS (ESI) calcd for $C_{35}H_{20}D_{16}N_3O_6Si$ $[M+H]^+$ 638.3372, found 638.3380.

### HaloTag Ligand Formation

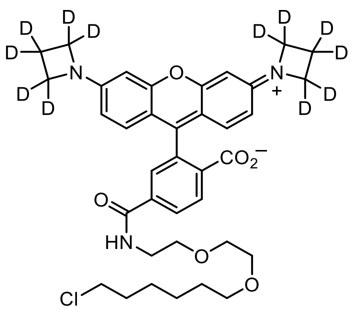

**JFX$_{549}$–HaloTag ligand (18$_D$).** 6-Carboxy-JFX$_{549}$ (**S5**; 25 mg, 43.1 μmol, TFA salt) was combined with DSC (26.5 mg, 0.103 mmol, 2.4 eq) in DMF (1 mL). After adding $Et_3N$ (36.0 μL, 0.258 mmol, 6 eq) and DMAP (0.5 mg, 4.3 μmol, 0.1 eq), the reaction was stirred at room temperature for 30 min. A solution of 2-(2-((6-chlorohexyl)oxy)ethoxy)-ethanamine (**S8**, 'HaloTag(O2)amine'; 43.6 mg, 0.129 mmol, 3 eq) in DMF (500 μL) was then added. The reaction was stirred an additional 2 h at room temperature. Purification of the crude reaction mixture by reverse phase HPLC (20–60% MeCN/$H_2O$, linear gradient, with constant 0.1% v/v TFA additive) afforded 22.4 mg (66%, TFA salt) of **18$_D$** as a dark red solid. $^1H$ NMR ($CD_3OD$, 400 MHz) δ 8.78 (t, $J$ = 5.5 Hz, 1H), 8.39 (d, $J$ = 8.2 Hz, 1H), 8.20 (dd, $J$ = 8.2, 1.8 Hz, 1H), 7.80 (d, $J$ = 1.7 Hz, 1H), 7.06 (d, $J$ = 9.2 Hz, 2H), 6.60 (dd, $J$ = 9.2, 2.2 Hz, 2H), 6.55 (d, $J$ = 2.2 Hz, 2H), 3.69 – 3.55 (m, 8H), 3.53 (t, $J$ = 6.6 Hz, 2H), 3.43 (t, $J$ = 6.5 Hz, 2H), 1.76 – 1.67 (m, 2H), 1.55 – 1.46 (m, 2H), 1.45 – 1.28 (m, 4H); Analytical HPLC: $t_R$ = 12.3 min, >99% purity (10–95% MeCN/$H_2O$, linear gradient, with constant 0.1% v/v TFA additive; 20 min run; 1 mL/min flow; ESI; positive ion mode; detection at 550 nm); HRMS (ESI) calcd for $C_{37}H_{31}D_{12}ClN_3O_6$ $[M+H]^+$ 672.3588, found 672.3590.

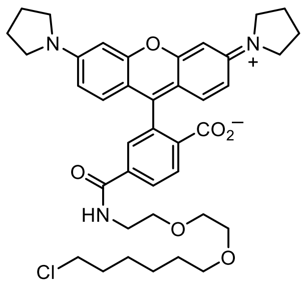

**Rh$_P$–HaloTag ligand (19).** The title compound (58%, dark red-purple solid, TFA salt) was prepared from 6-carboxy-Rh$_P$ (**S6**) and 2-(2-((6-chlorohexyl)oxy)ethoxy)ethanamine (**S8**) according to the procedure described for **18$_D$**. $^1H$ NMR ($CD_3OD$, 400 MHz) δ 8.76 (t, $J$ = 5.3 Hz, 1H), 8.40 (d, $J$ = 8.2 Hz, 1H), 8.21 (dd, $J$ = 8.3, 1.8 Hz, 1H), 7.83 (d, $J$ = 1.7 Hz, 1H), 7.12 (d, $J$ = 9.3 Hz, 2H), 6.91 (dd, $J$ = 9.3, 2.3 Hz, 2H), 6.84 (d, $J$ = 2.3 Hz, 2H), 3.71 – 3.54 (m, 16H), 3.51 (d, $J$ = 6.6 Hz, 2H), 3.43 (t, $J$ = 6.5 Hz, 2H), 2.20 – 2.05 (m, 8H), 1.76 – 1.66 (m, 2H), 1.54 – 1.45 (m, 2H), 1.45 – 1.26 (m, 4H); Analytical HPLC: $t_R$ = 13.2 min, 98.4% purity (10–95% MeCN/$H_2O$, linear gradient, with constant 0.1% v/v TFA additive; 20 min run; 1 mL/min flow; ESI; positive ion mode; detection at 550 nm); HRMS (ESI) calcd for $C_{39}H_{47}ClN_3O_6$ $[M+H]^+$ 688.3148, found 688.3156.

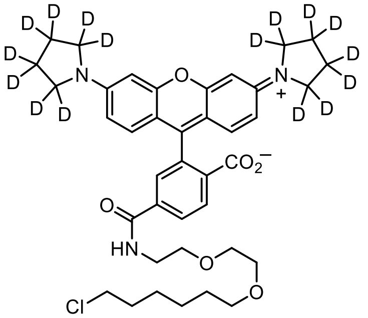

**JFX$_{554}$–HaloTag ligand (19$_D$).** The title compound (63%, dark red-purple solid, TFA salt) was prepared from 6-carboxy-JFX$_{554}$ (**S7**) and 2-(2-((6-chlorohexyl)oxy)ethoxy)ethanamine (**S8**) according to the procedure described for **18$_D$**. $^1$H NMR ($CD_3OD$, 400 MHz) δ 8.76 (t, *J* = 5.2 Hz, 1H), 8.40 (d, *J* = 8.2 Hz, 1H), 8.21 (dd, *J* = 8.3, 1.8 Hz, 1H), 7.83 (d, *J* = 1.7 Hz, 1H), 7.12 (d, *J* = 9.3 Hz, 2H), 6.91 (dd, *J* = 9.3, 2.3 Hz, 2H), 6.84 (d, *J* = 2.3 Hz, 2H), 3.70 – 3.54 (m, 8H), 3.52 (t, *J* = 6.6 Hz, 2H), 3.43 (t, *J* = 6.5 Hz, 2H), 1.76 – 1.66 (m, 2H), 1.55 – 1.45 (m, 2H), 1.45 – 1.27 (m, 4H); Analytical HPLC: $t_R$ = 13.1 min, 98.4% purity (10–95% MeCN/$H_2O$, linear gradient, with constant 0.1% v/v TFA additive; 20 min run; 1 mL/min flow; ESI; positive ion mode; detection at 550 nm); HRMS (ESI) calcd for $C_{39}H_{31}D_{16}ClN_3O_6$ $[M+H]^+$ 704.4152, found 704.4155.

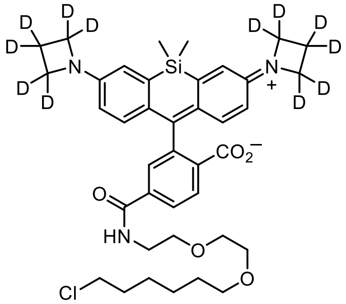

**JFX$_{646}$–HaloTag ligand (26$_D$).** The title compound (79%, pale blue-green solid) was prepared from 6-carboxy-JFX$_{646}$ (**S17**) and 2-(2-((6-chlorohexyl)oxy)ethoxy)ethanamine (**S8**) according to the procedure described for **18$_D$**. $^1$H NMR ($CDCl_3$, 400 MHz) δ 7.98 (d, *J* = 7.9 Hz, 1H), 7.90 (dd, *J* = 8.0, 1.4 Hz, 1H), 7.70 – 7.66 (m, 1H), 6.76 (s, 1H), 6.75 (d, *J* = 8.7 Hz, 2H), 6.66 (d, *J* = 2.7 Hz, 2H), 6.26 (dd, *J* = 8.6, 2.7 Hz, 2H), 3.67 – 3.59 (m, 6H), 3.56 – 3.53 (m, 2H), 3.50 (t, *J* = 6.6 Hz, 2H), 3.39 (t, *J* = 6.7 Hz, 2H), 1.76 – 1.69 (m, 2H), 1.54 – 1.48 (m, 2H), 1.44 – 1.26 (m, 4H), 0.63 (s, 3H), 0.57 (s, 3H); Analytical HPLC: $t_R$ = 13.2 min, 98.7% purity (10–95% MeCN/$H_2O$, linear gradient, with constant 0.1% v/v TFA additive; 20 min run; 1 mL/min flow; ESI; positive ion mode; detection at 650 nm); HRMS (ESI) calcd for $C_{39}H_{37}D_{12}ClN_3O_5Si$ $[M+H]^+$ 714.3878, found 714.3885.

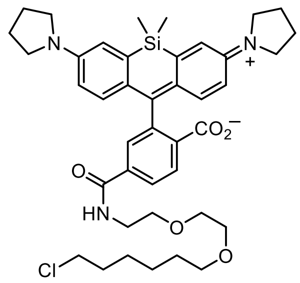


**SiRh$_P$–HaloTag ligand (27).** SiRh$_P$-NHS (**S20**; 75 mg, 0.121 mmol) and 2-(2-((6-chlorohexyl)oxy)ethoxy)ethanamine (**S8**; 61.1 mg, 0.181 mmol, 1.5 eq) were combined in DMF (3 mL), and DIEA (63.0 µL, 0.362 mmol, 3 eq) was added. After stirring the reaction at room temperature for 1 h, it was diluted with saturated $NaHCO_3$ and extracted with EtOAc (2×). The combined organic extracts were washed with water and brine, dried over anhydrous $MgSO_4$, filtered, and evaporated. Purification of the crude product by silica gel chromatography (10–100% EtOAc/toluene, linear gradient) provided **27** as a pale blue-green solid (64.4 mg, 73%). $^1$H NMR ($CDCl_3$, 400 MHz) δ 7.99 (d, *J* = 7.9 Hz, 1H), 7.91 (dd, *J* = 7.9, 1.4 Hz, 1H), 7.67 – 7.64 (m, 1H), 6.79 (d, *J* = 2.7 Hz, 2H), 6.75 (s, 1H), 6.75 (d, *J* = 8.8 Hz, 2H), 6.39 (dd, *J* = 8.9, 2.7 Hz, 2H), 3.67 – 3.58 (m, 6H), 3.57 – 3.52 (m, 2H), 3.50 (t, *J* = 6.7 Hz, 2H), 3.39 (t, *J* = 6.7 Hz, 2H), 3.35 – 3.24 (m, 8H), 2.05 – 1.93 (m, 8H), 1.77 – 1.68 (m, 2H), 1.55 – 1.47 (m, 2H), 1.44 – 1.26 (m, 4H), 0.65 (s, 3H), 0.59 (s, 3H); Analytical HPLC: $t_R$ = 13.5 min, >99% purity (10–95% MeCN/$H_2O$, linear gradient, with constant 0.1% v/v TFA additive; 20 min run; 1 mL/min flow; ESI; positive ion mode; detection at 650 nm); HRMS (ESI) calcd for $C_{41}H_{53}ClN_3O_5Si$ [M+H]$^+$ 730.3438, found 730.3442.

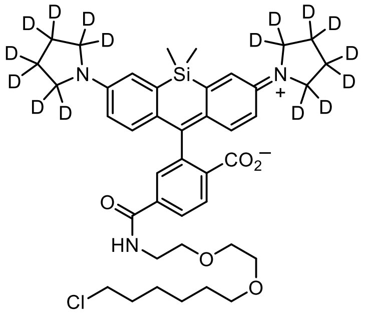


**JFX$_{650}$–HaloTag ligand (27$_D$).** The title compound (77%, pale blue-green solid) was prepared from JFX$_{650}$-NHS (**S21**) and 2-(2-((6-chlorohexyl)oxy)ethoxy)ethanamine (**S8**) according to the procedure described for **27**. $^1$H NMR ($CDCl_3$, 400 MHz) δ 7.99 (dd, *J* = 8.0, 0.5 Hz, 1H), 7.91 (dd, *J* = 8.0, 1.4 Hz, 1H), 7.67 – 7.63 (m, 1H), 6.78 (d, *J* = 2.8 Hz, 2H), 6.747 (d, *J* = 8.8 Hz, 2H), 6.744 (s, 1H), 6.39 (dd, *J* = 8.8, 2.8 Hz, 2H), 3.66 – 3.58 (m, 6H), 3.57 – 3.52 (m, 2H), 3.50 (t, *J* = 6.7 Hz, 2H), 3.39 (t, *J* = 6.7 Hz, 2H), 1.77 – 1.68 (m, 2H), 1.55 – 1.47 (m, 2H), 1.44 – 1.35 (m, 2H), 1.34 – 1.25 (m, 2H), 0.65 (s, 3H), 0.58 (s, 3H); Analytical HPLC: $t_R$ = 13.4 min, >99% purity (10–95% MeCN/$H_2O$, linear gradient, with constant 0.1% v/v TFA additive; 20 min run; 1 mL/min flow; ESI; positive ion mode; detection at 650 nm); HRMS (ESI) calcd for $C_{41}H_{37}D_{16}ClN_3O_5Si$ [M+H]$^+$ 746.4442, found 746.4449.

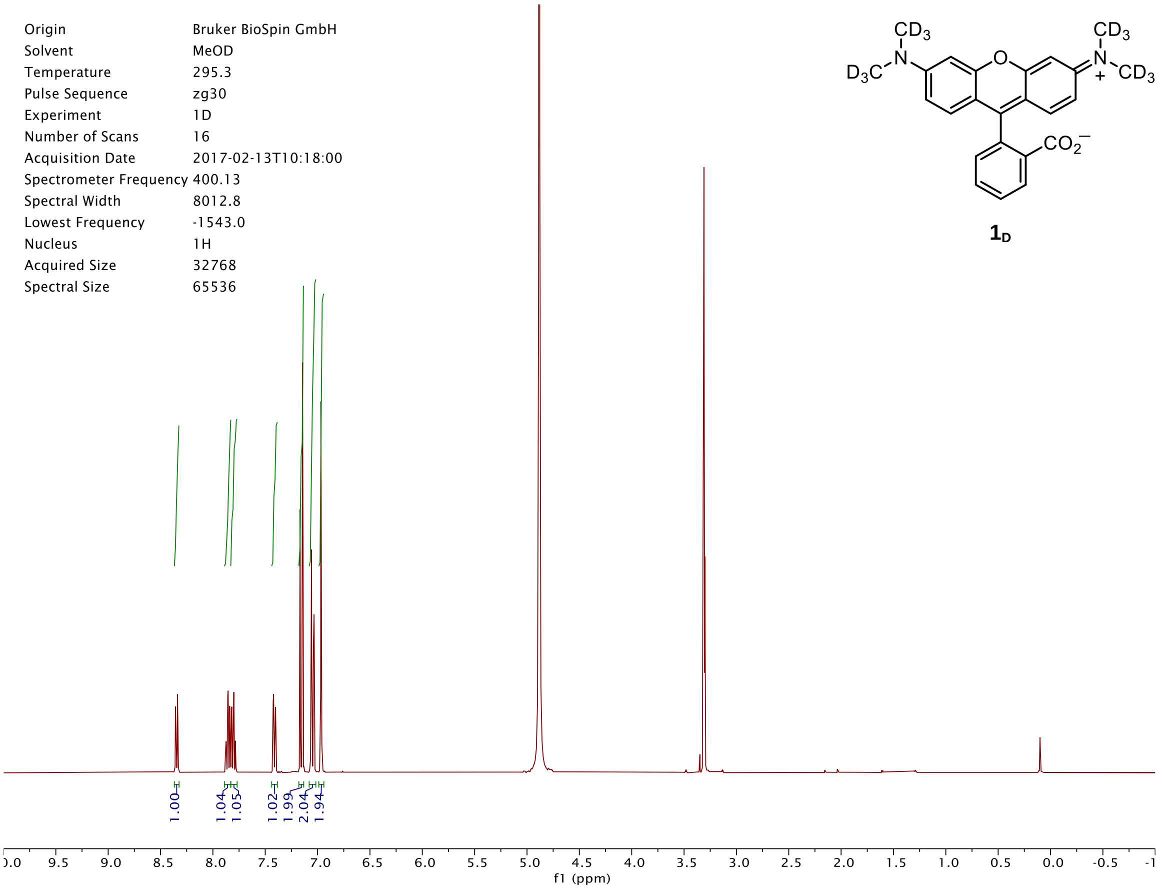


| | |
|---|---|
| Origin | Bruker BioSpin GmbH |
| Solvent | MeOD |
| Temperature | 295.3 |
| Pulse Sequence | zg30 |
| Experiment | 1D |
| Number of Scans | 16 |
| Acquisition Date | 2017-02-13T10:18:00 |
| Spectrometer Frequency | 400.13 |
| Spectral Width | 8012.8 |
| Lowest Frequency | -1543.0 |
| Nucleus | 1H |
| Acquired Size | 32768 |
| Spectral Size | 65536 |

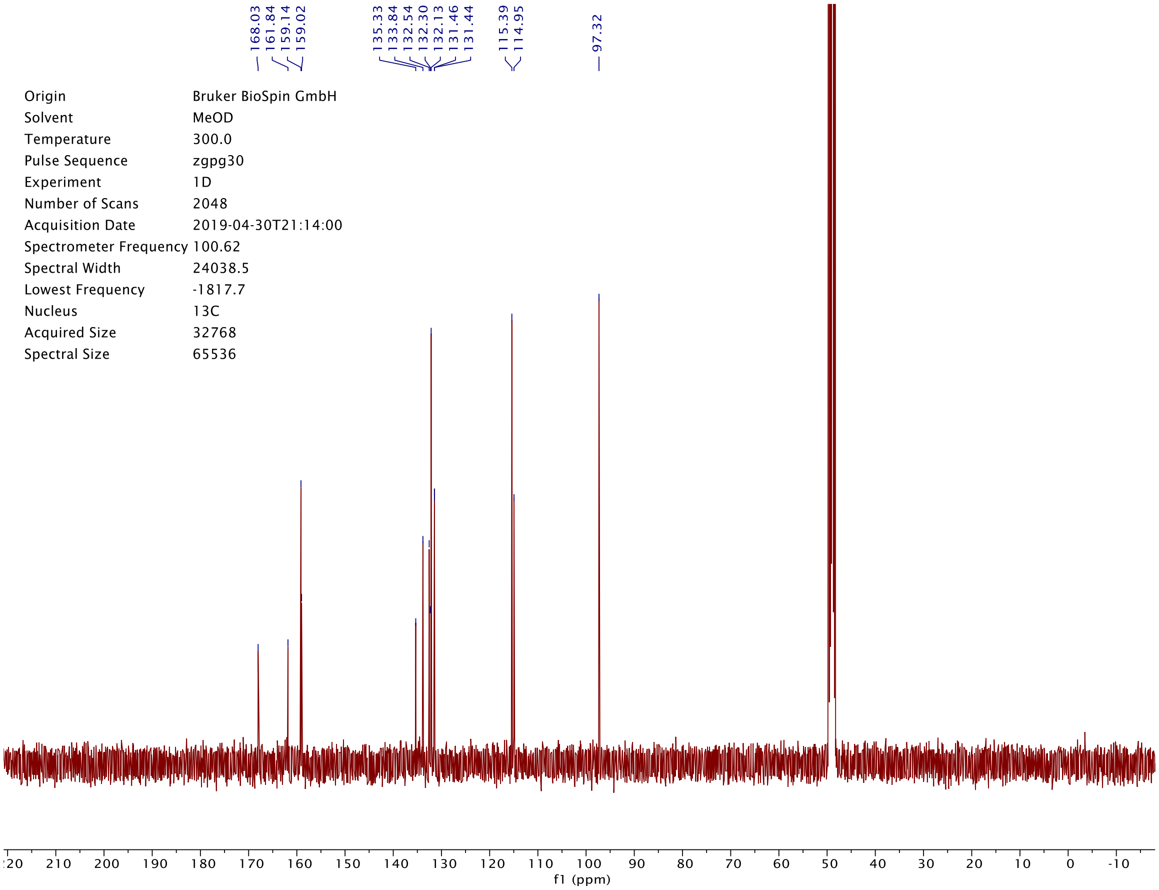


| | |
|---|---|
| Origin | Bruker BioSpin GmbH |
| Solvent | MeOD |
| Temperature | 300.0 |
| Pulse Sequence | zgpg30 |
| Experiment | 1D |
| Number of Scans | 2048 |
| Acquisition Date | 2019-04-30T21:14:00 |
| Spectrometer Frequency | 100.62 |
| Spectral Width | 24038.5 |
| Lowest Frequency | -1817.7 |
| Nucleus | 13C |
| Acquired Size | 32768 |
| Spectral Size | 65536 |

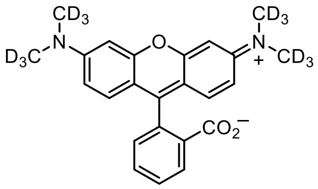


**$1_D$**

DAD1 C, Sig=550,4 Ref=off (2019_03\DAILY_SEQUENCE_LC 2019-03-26 11-59-39\2019_030000002.D)

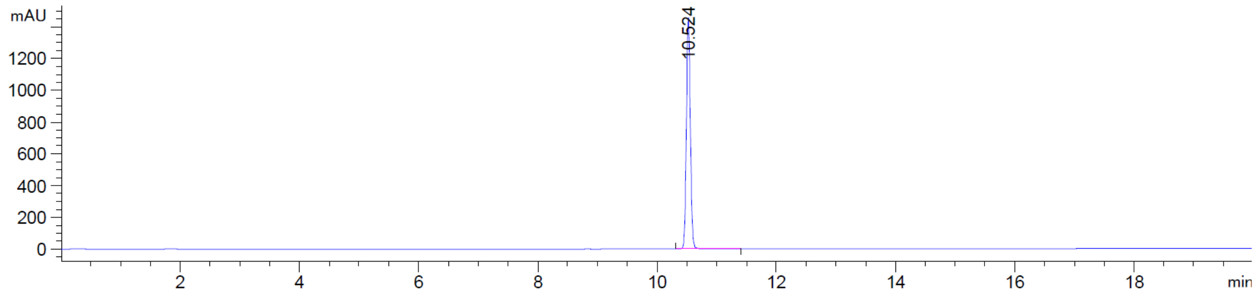


*MSD2 SPC, time=10.535:10.607 of C:\CHEM32\1\DATA\2019_03\DAILY_SEQUENCE_LC 2019-03-26 11-59-39\2019_030000002.D ES-

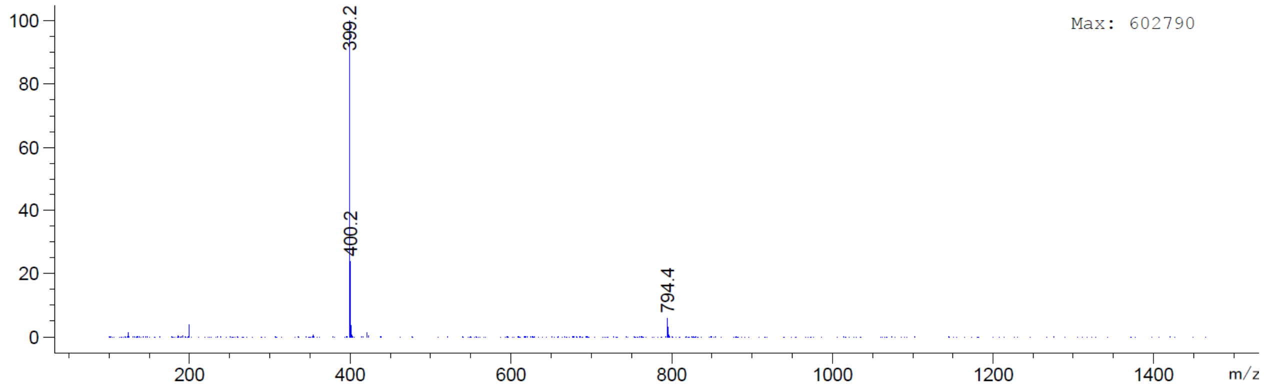

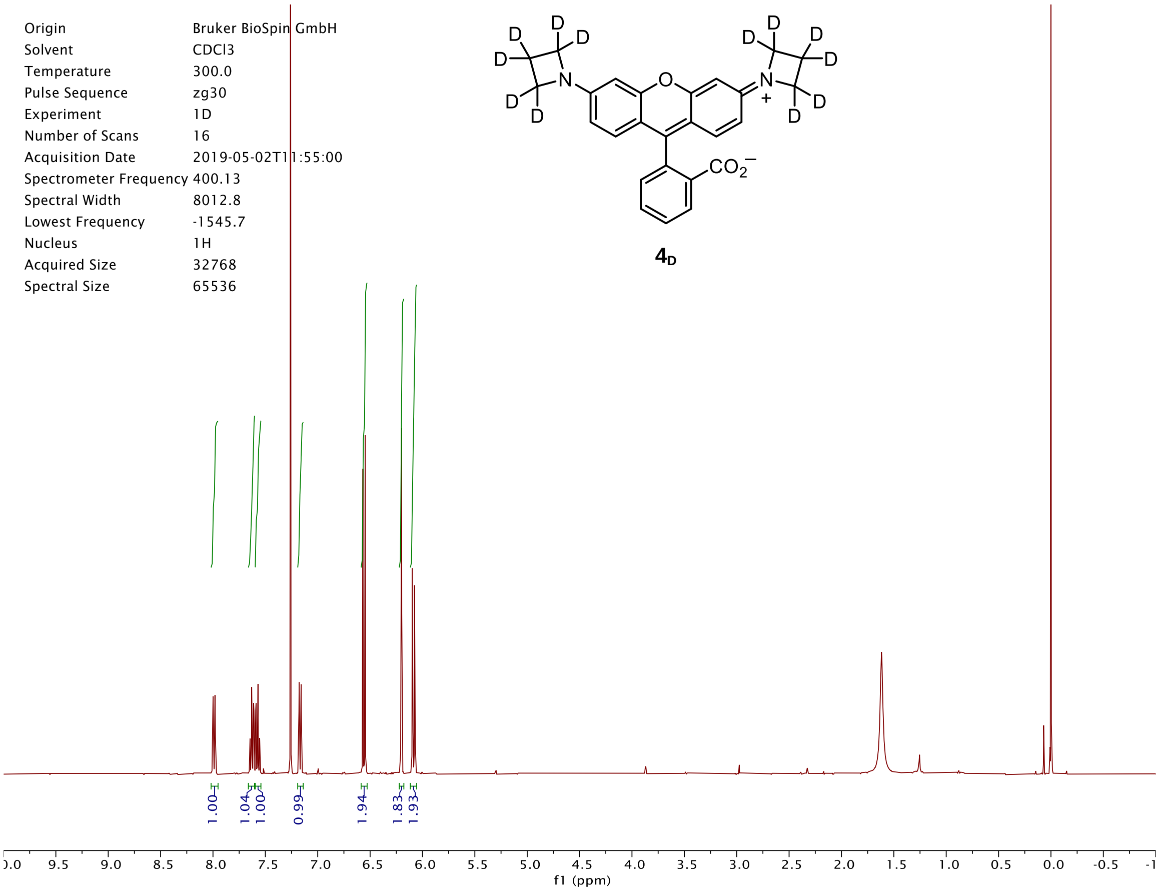

Origin Bruker BioSpin GmbH
Solvent CDCl3
Temperature 300.0
Pulse Sequence zg30
Experiment 1D
Number of Scans 16
Acquisition Date 2019-05-02T11:55:00
Spectrometer Frequency 400.13
Spectral Width 8012.8
Lowest Frequency -1545.7
Nucleus 1H
Acquired Size 32768
Spectral Size 65536
4D
1.00
1.04
1.00
0.99
1.94
1.83
1.93
f1 (ppm)


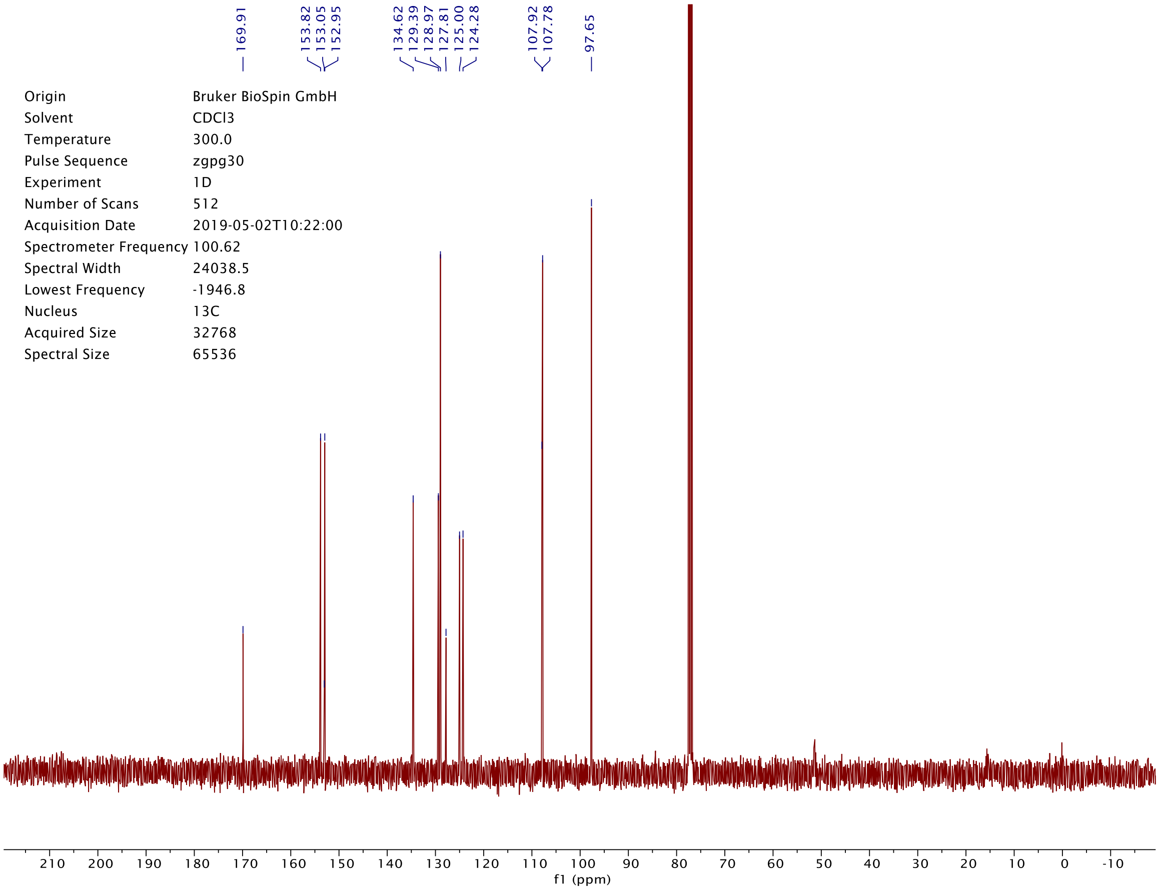

169.91
153.82
153.05
152.95
134.62
129.39
128.97
127.81
125.00
124.28
107.92
107.78
97.65
Origin Bruker BioSpin GmbH
Solvent CDCl3
Temperature 300.0
Pulse Sequence zgpg30
Experiment 1D
Number of Scans 512
Acquisition Date 2019-05-02T10:22:00
Spectrometer Frequency 100.62
Spectral Width 24038.5
Lowest Frequency -1946.8
Nucleus 13C
Acquired Size 32768
Spectral Size 65536
f1 (ppm)

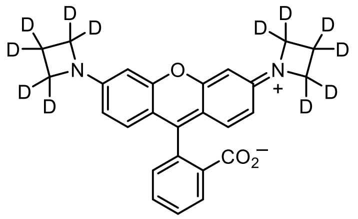

**4**$_D$

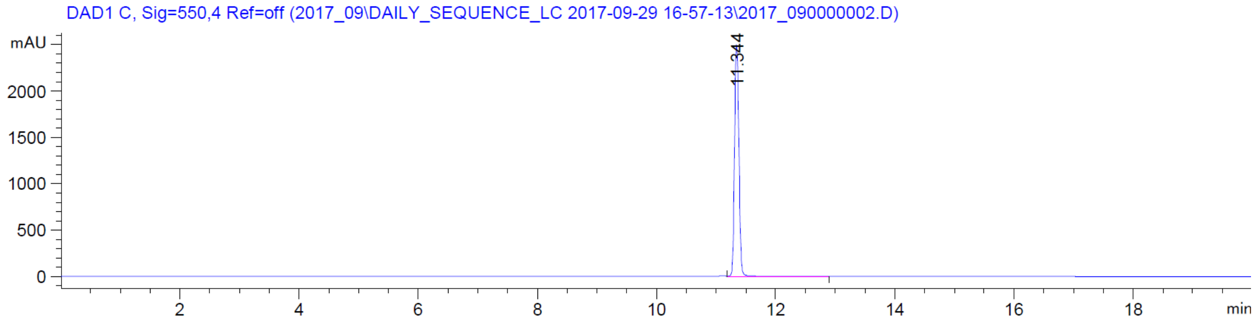

DAD1 C, Sig=550,4 Ref=off (2017_09\DAILY_SEQUENCE_LC 2017-09-29 16-57-13\2017_090000002.D)
mAU
2000
1500
1000
500
0
2
4
6
8
10
12
14
16
18
min
11.344


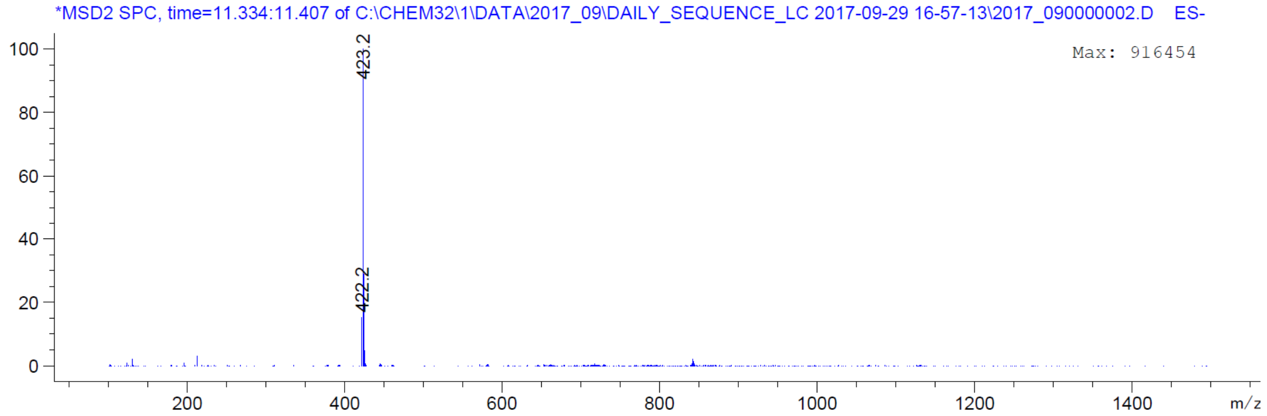

*MSD2 SPC, time=11.334:11.407 of C:\CHEM32\1\DATA\2017_09\DAILY_SEQUENCE_LC 2017-09-29 16-57-13\2017_090000002.D ES-
Max: 916454
100
80
60
40
20
0
200
400
600
800
1000
1200
1400
m/z
423.2
422.2

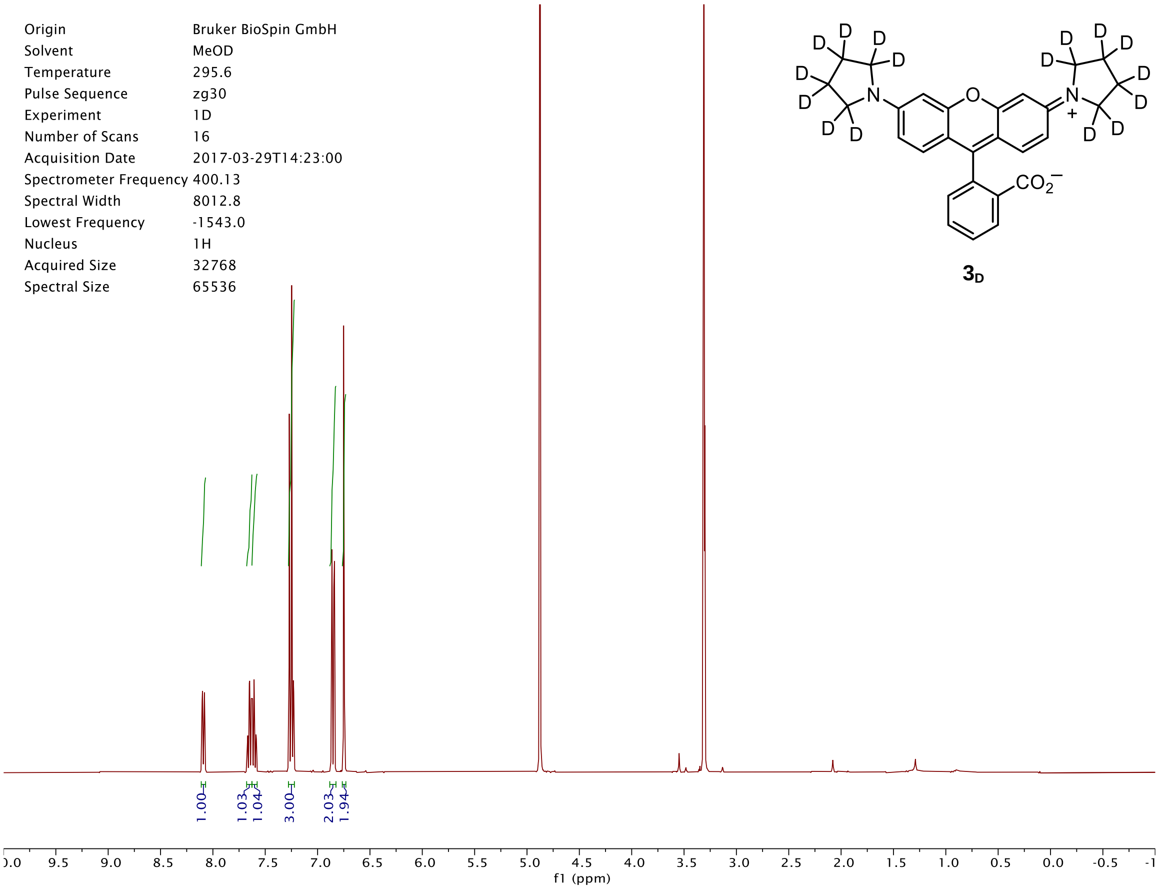
Origin Bruker BioSpin GmbH
Solvent MeOD
Temperature 295.6
Pulse Sequence zg30
Experiment 1D
Number of Scans 16
Acquisition Date 2017-03-29T14:23:00
Spectrometer Frequency 400.13
Spectral Width 8012.8
Lowest Frequency -1543.0
Nucleus 1H
Acquired Size 32768
Spectral Size 65536
CO2−
3D
1.00
1.03
1.04
3.00
2.03
1.94
f1 (ppm)

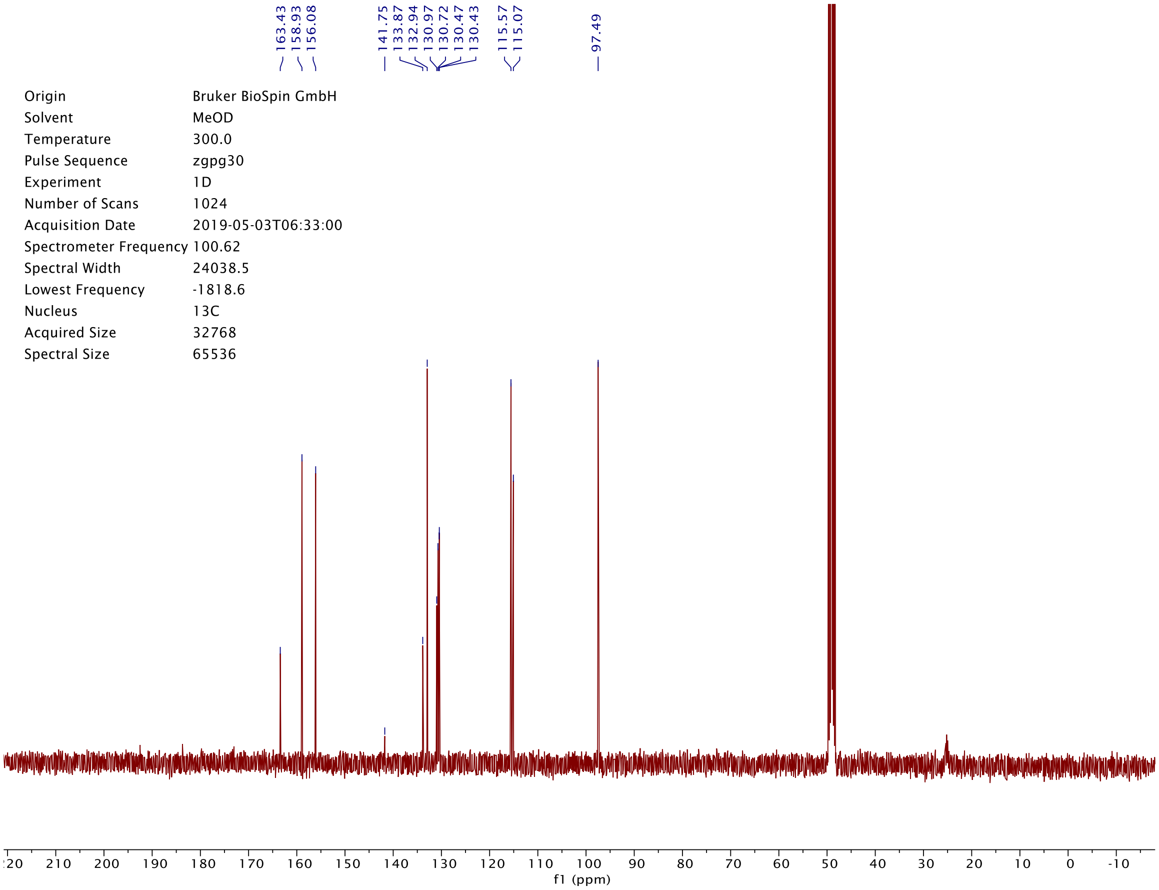
163.43
158.93
156.08
141.75
133.87
132.94
130.97
130.72
130.47
130.43
115.57
115.07
97.49
Origin Bruker BioSpin GmbH
Solvent MeOD
Temperature 300.0
Pulse Sequence zgpg30
Experiment 1D
Number of Scans 1024
Acquisition Date 2019-05-03T06:33:00
Spectrometer Frequency 100.62
Spectral Width 24038.5
Lowest Frequency -1818.6
Nucleus 13C
Acquired Size 32768
Spectral Size 65536
f1 (ppm)

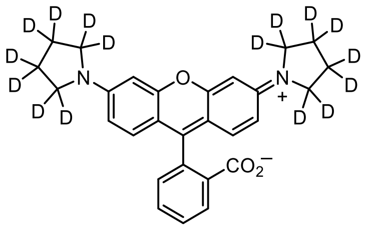

**$3_D$**

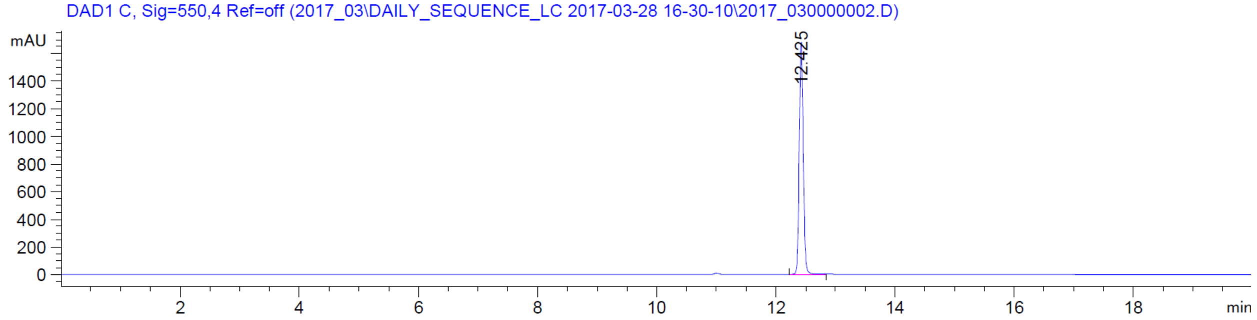

DAD1 C, Sig=550,4 Ref=off (2017_03\DAILY_SEQUENCE_LC 2017-03-28 16-30-10\2017_030000002.D)
mAU
1400
1200
1000
800
600
400
200
0
12.425
2
4
6
8
10
12
14
16
18
min


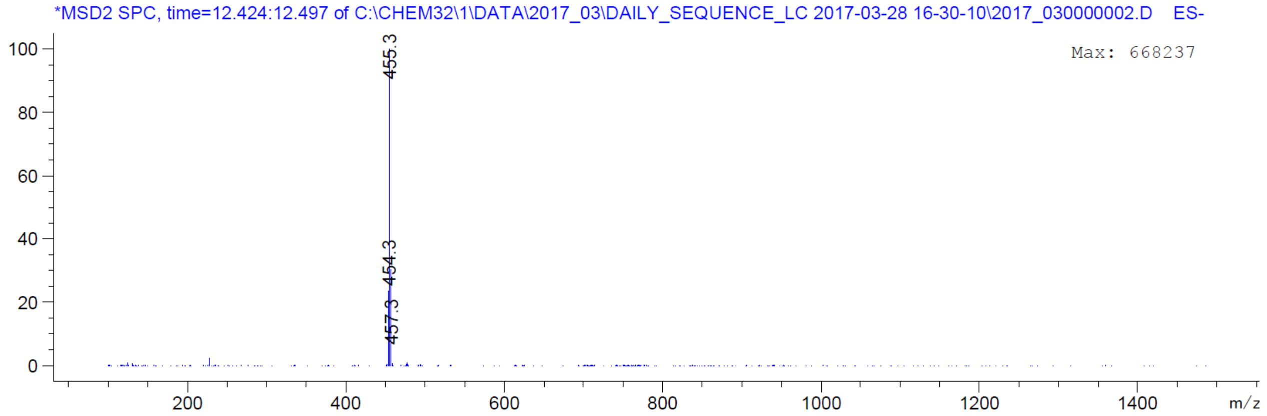

*MSD2 SPC, time=12.424:12.497 of C:\CHEM32\1\DATA\2017_03\DAILY_SEQUENCE_LC 2017-03-28 16-30-10\2017_030000002.D ES-
Max: 668237
100
80
60
40
20
0
455.3
454.3
457.3
200
400
600
800
1000
1200
1400
m/z

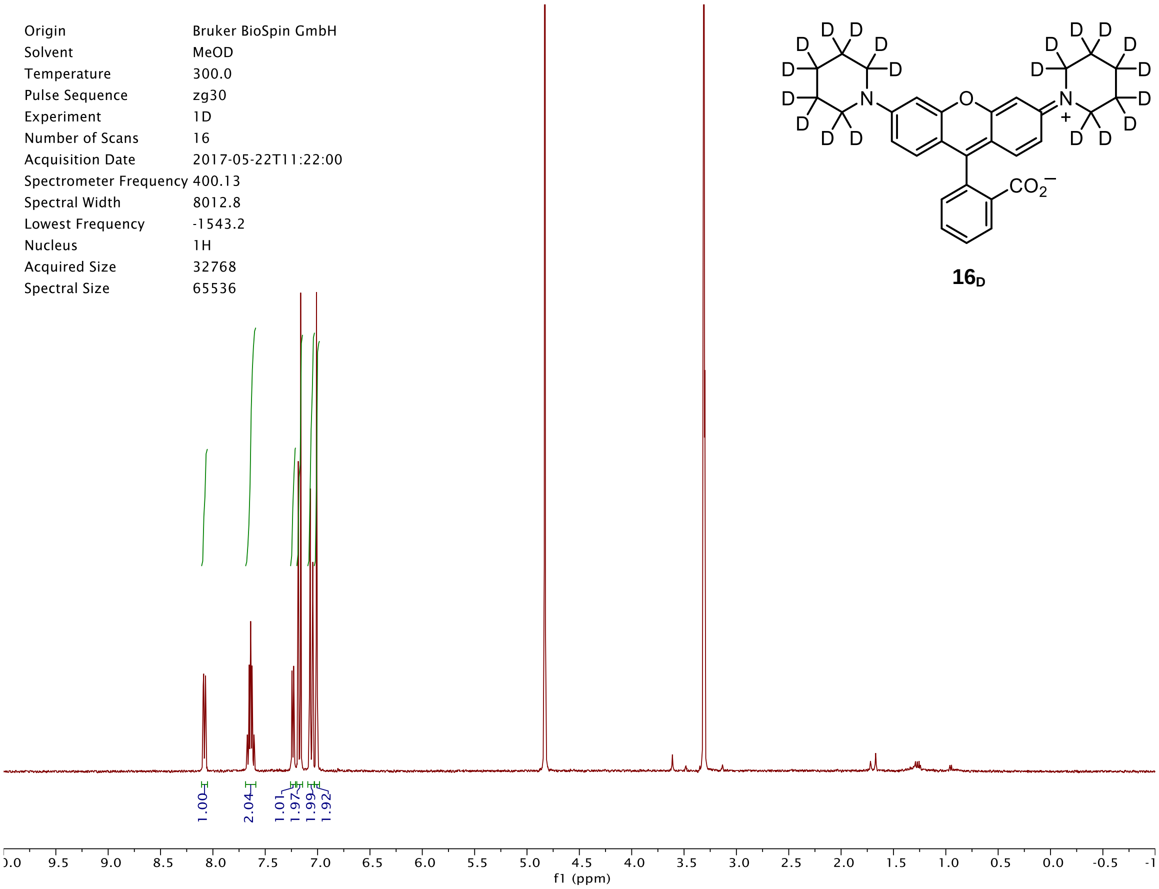
D
N
O
N
+
$CO_2^-$
Origin Bruker BioSpin GmbH
Solvent MeOD
Temperature 300.0
Pulse Sequence zg30
Experiment 1D
Number of Scans 16
Acquisition Date 2017-05-22T11:22:00
Spectrometer Frequency 400.13
Spectral Width 8012.8
Lowest Frequency -1543.2
Nucleus 1H
Acquired Size 32768
Spectral Size 65536
$16_D$
1.00
2.04
1.01
1.97
1.99
1.92
f1 (ppm)

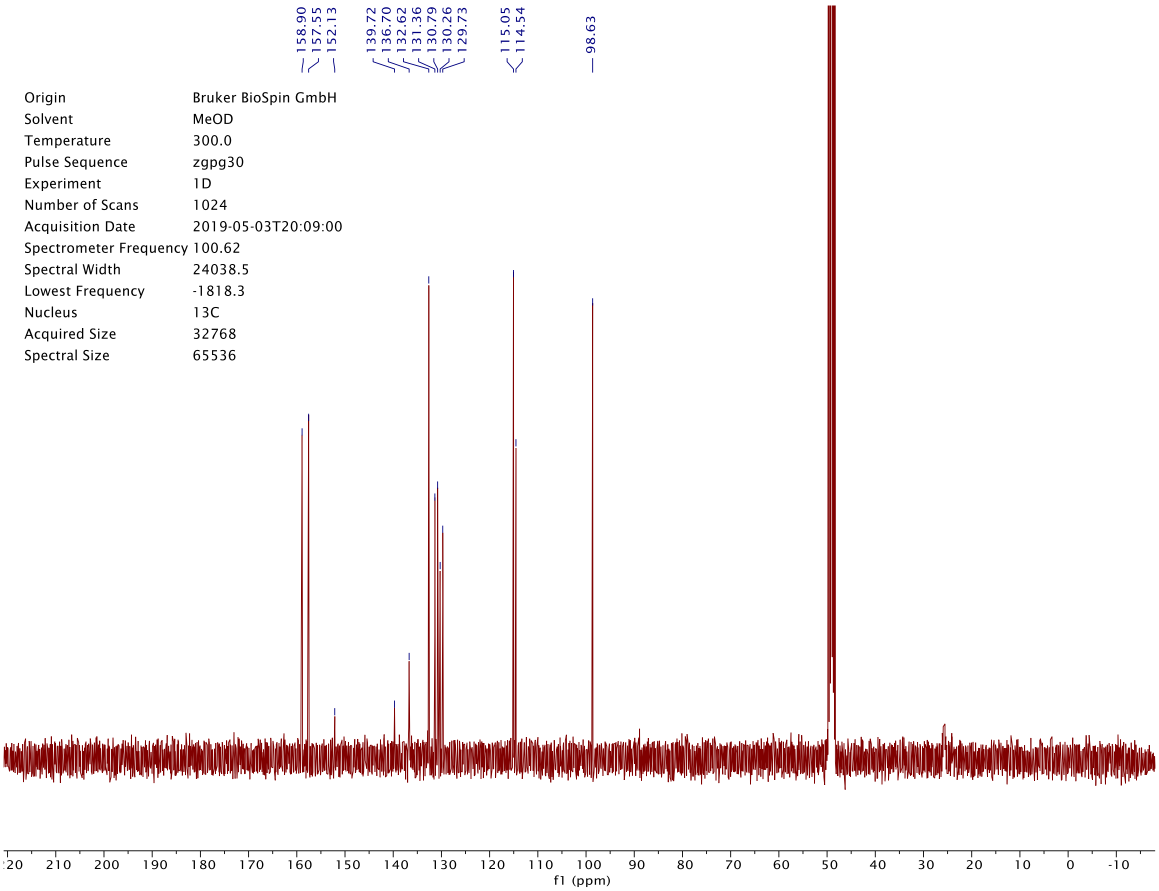
158.90
157.55
152.13
139.72
136.70
132.62
131.36
130.79
130.26
129.73
115.05
114.54
98.63
Origin Bruker BioSpin GmbH
Solvent MeOD
Temperature 300.0
Pulse Sequence zgpg30
Experiment 1D
Number of Scans 1024
Acquisition Date 2019-05-03T20:09:00
Spectrometer Frequency 100.62
Spectral Width 24038.5
Lowest Frequency -1818.3
Nucleus 13C
Acquired Size 32768
Spectral Size 65536
f1 (ppm)

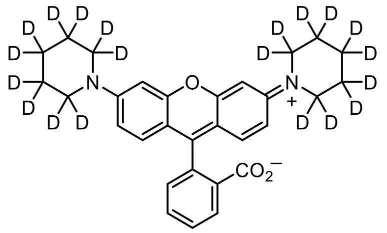

**$16_D$**

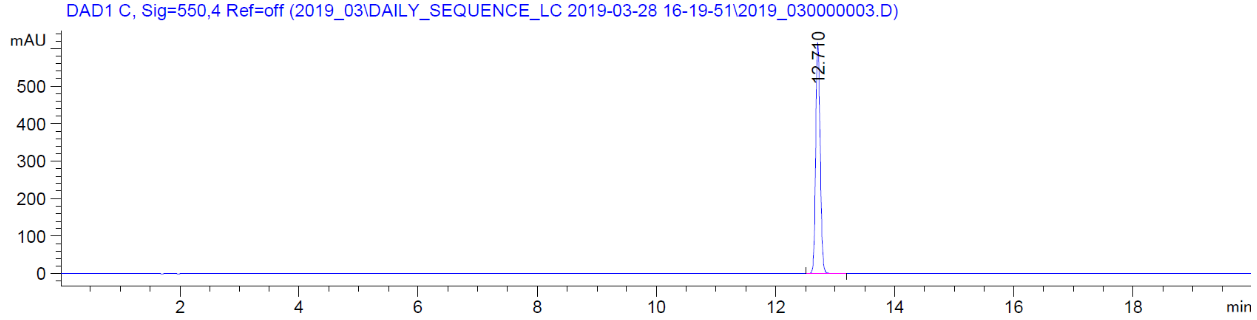

DAD1 C, Sig=550,4 Ref=off (2019_03\DAILY_SEQUENCE_LC 2019-03-28 16-19-51\2019_030000003.D)
mAU
12.710
min


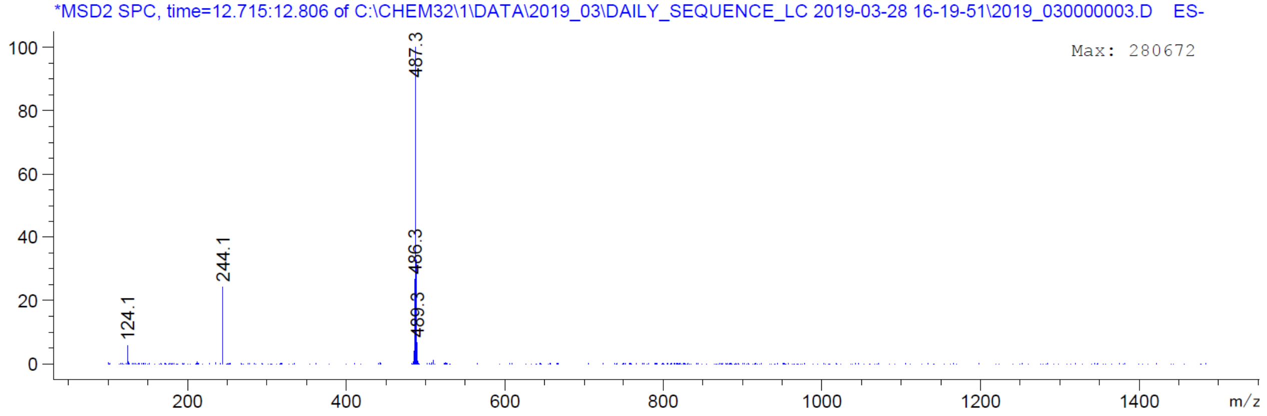

*MSD2 SPC, time=12.715:12.806 of C:\CHEM32\1\DATA\2019_03\DAILY_SEQUENCE_LC 2019-03-28 16-19-51\2019_030000003.D ES-
Max: 280672
487.3
486.3
489.3
244.1
124.1
m/z

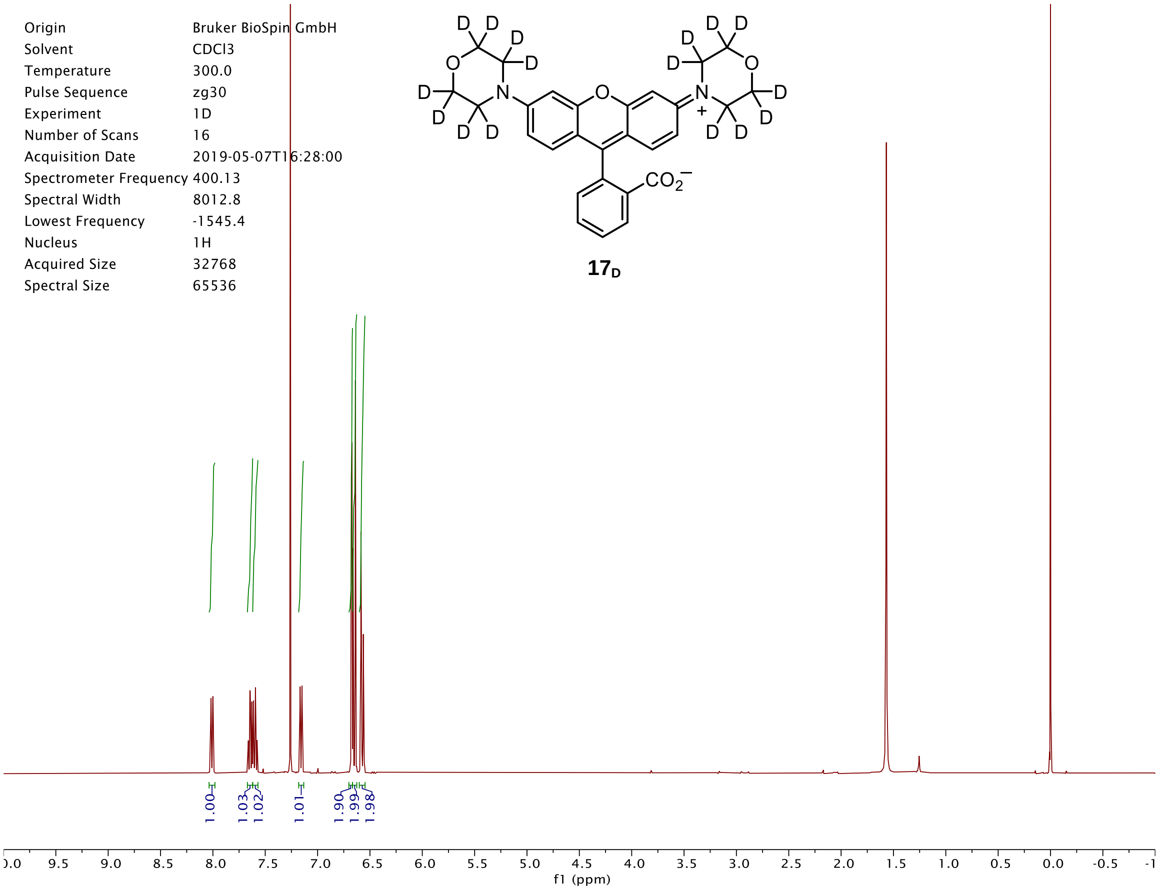
Origin Bruker BioSpin GmbH
Solvent CDCl3
Temperature 300.0
Pulse Sequence zg30
Experiment 1D
Number of Scans 16
Acquisition Date 2019-05-07T16:28:00
Spectrometer Frequency 400.13
Spectral Width 8012.8
Lowest Frequency -1545.4
Nucleus 1H
Acquired Size 32768
Spectral Size 65536
1.00
1.03
1.02
1.01
1.90
1.99
1.98
f1 (ppm)

**17$_D$**

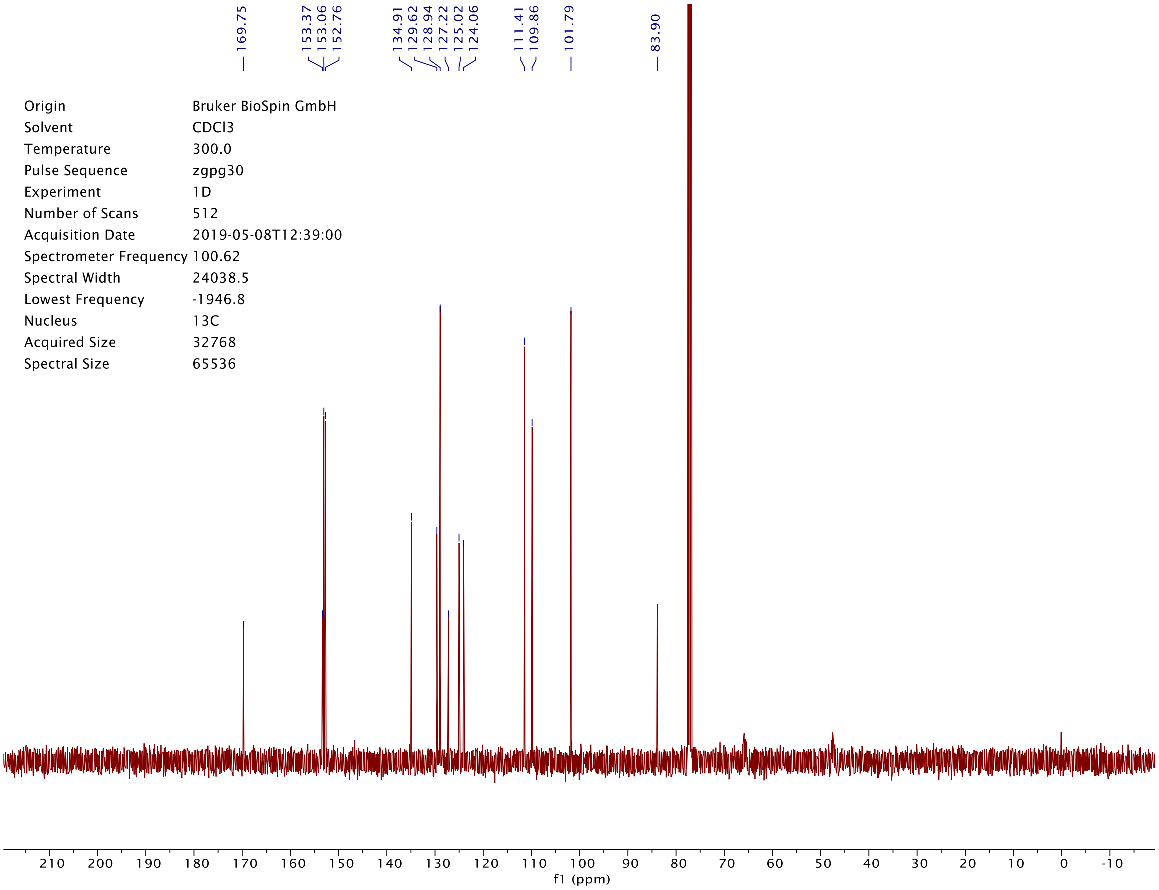
169.75
153.37
153.06
152.76
134.91
129.62
128.94
127.22
125.02
124.06
111.41
109.86
101.79
83.90
Origin Bruker BioSpin GmbH
Solvent CDCl3
Temperature 300.0
Pulse Sequence zgpg30
Experiment 1D
Number of Scans 512
Acquisition Date 2019-05-08T12:39:00
Spectrometer Frequency 100.62
Spectral Width 24038.5
Lowest Frequency -1946.8
Nucleus 13C
Acquired Size 32768
Spectral Size 65536
f1 (ppm)

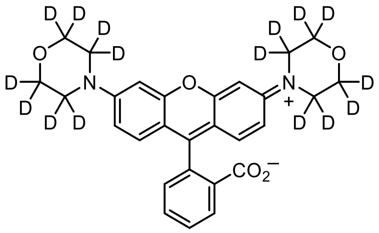

**17**$_D$

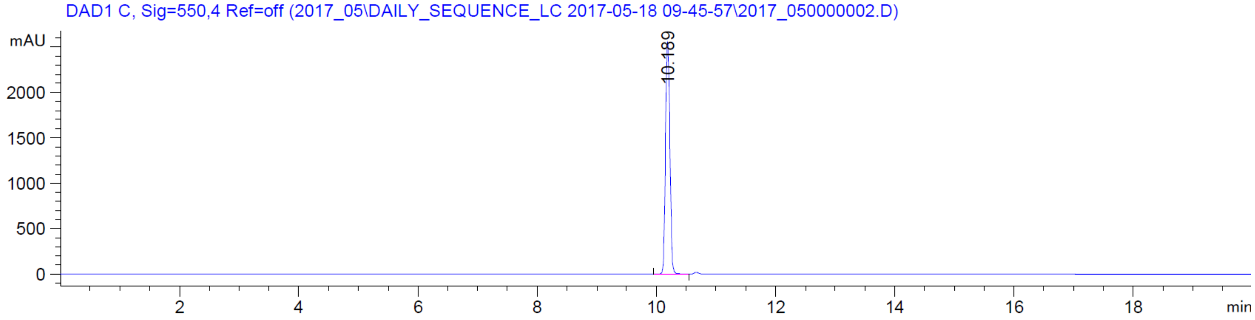

DAD1 C, Sig=550,4 Ref=off (2017_05\DAILY_SEQUENCE_LC 2017-05-18 09-45-57\2017_050000002.D)
mAU
2000
1500
1000
500
0
10.189
2
4
6
8
10
12
14
16
18
min


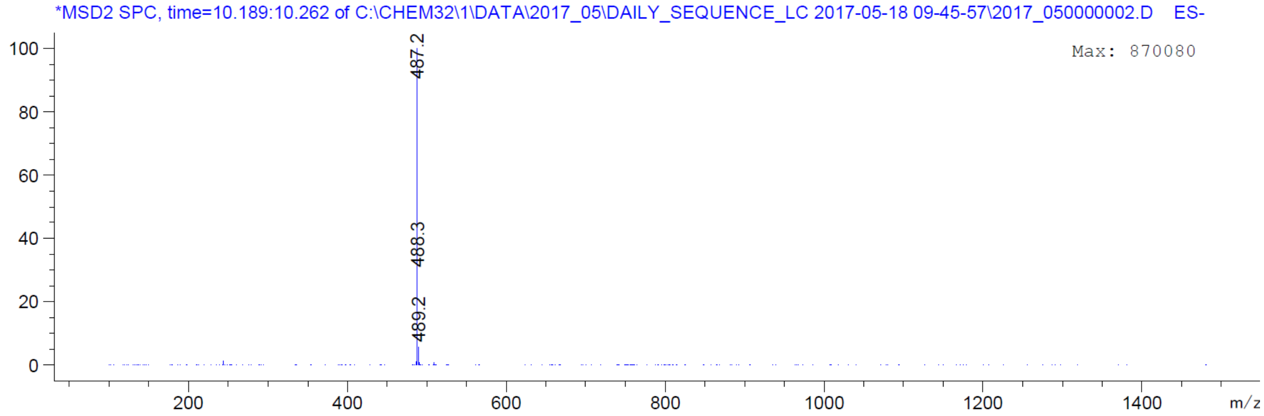

*MSD2 SPC, time=10.189:10.262 of C:\CHEM32\1\DATA\2017_05\DAILY_SEQUENCE_LC 2017-05-18 09-45-57\2017_050000002.D ES-
Max: 870080
100
80
60
40
20
0
487.2
488.3
489.2
200
400
600
800
1000
1200
1400
m/z

| | |
|---|---|
| Origin | Bruker BioSpin GmbH |
| Solvent | CDCl3 |
| Temperature | 295.5 |
| Pulse Sequence | zg30 |
| Experiment | 1D |
| Number of Scans | 16 |
| Acquisition Date | 2017-10-13T12:19:00 |
| Spectrometer Frequency | 400.13 |
| Spectral Width | 8012.8 |
| Lowest Frequency | -1544.9 |
| Nucleus | 1H |
| Acquired Size | 32768 |
| Spectral Size | 65536 |

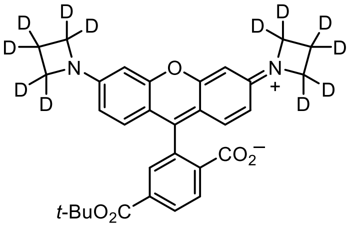


**S2**

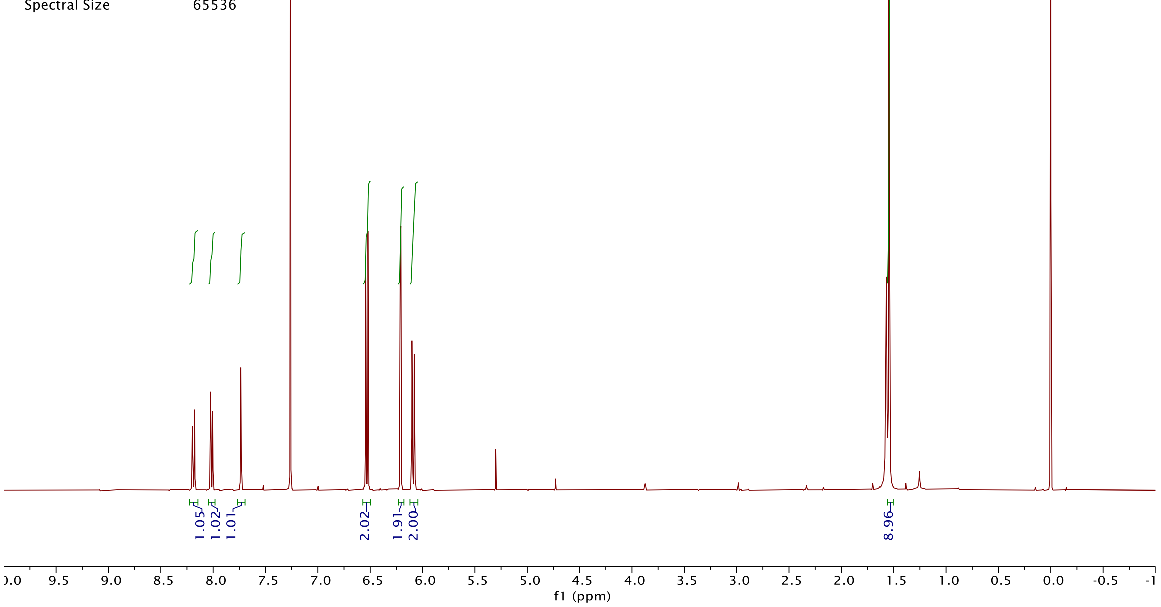


| | |
|---|---|
| Origin | Bruker BioSpin GmbH |
| Solvent | CDCl3 |
| Temperature | 300.0 |
| Pulse Sequence | zgpg30 |
| Experiment | 1D |
| Number of Scans | 512 |
| Acquisition Date | 2019-05-15T01:58:00 |
| Spectrometer Frequency | 100.62 |
| Spectral Width | 24038.5 |
| Lowest Frequency | -1946.1 |
| Nucleus | 13C |
| Acquired Size | 32768 |
| Spectral Size | 65536 |

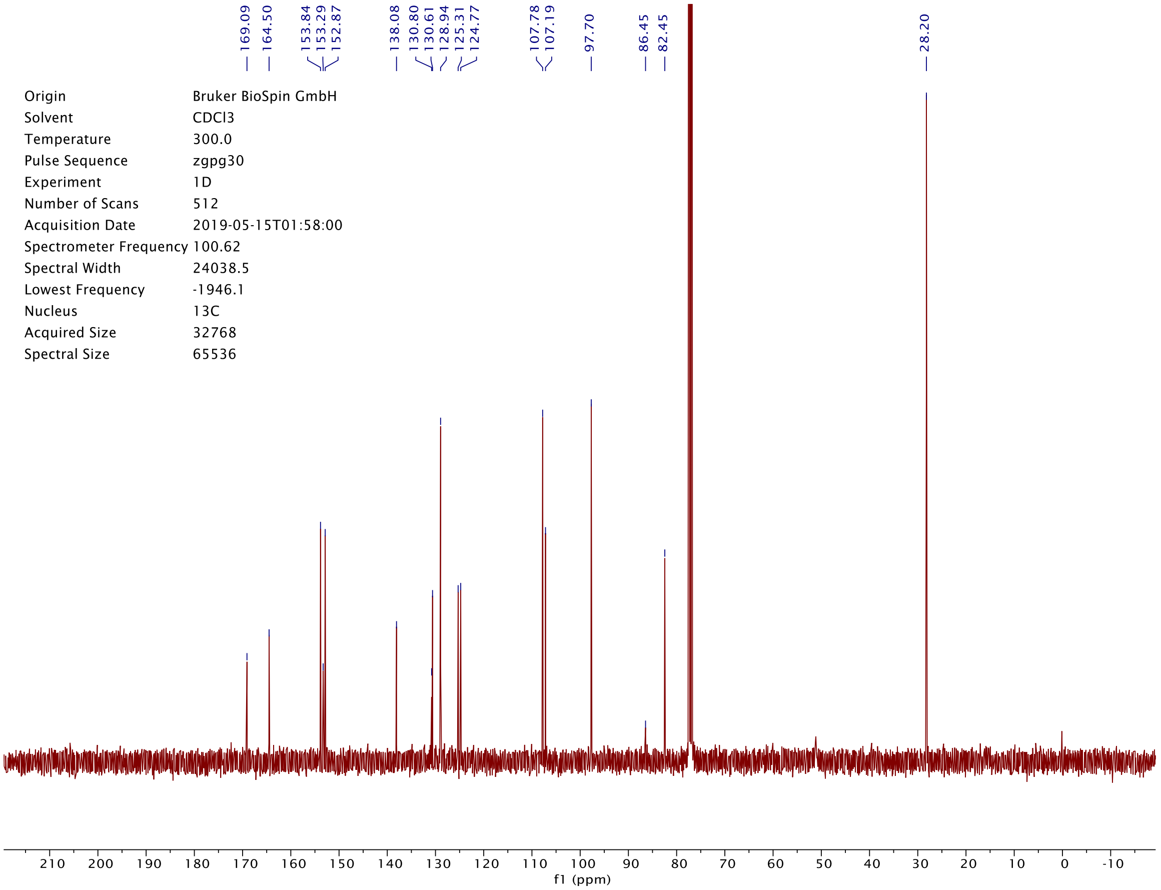

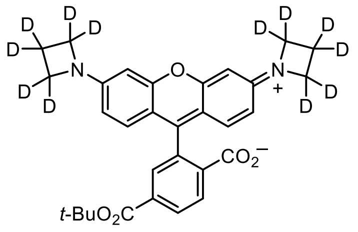

D
D
D
D
D
D
D
D
D
D
D
D
N
O
N
+
$CO_2^-$
t-Bu$O_2$C


**S2**

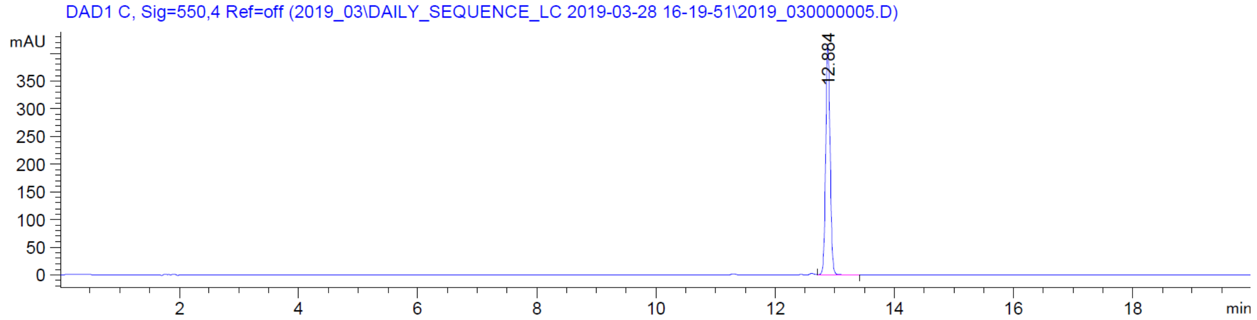

DAD1 C, Sig=550,4 Ref=off (2019_03\DAILY_SEQUENCE_LC 2019-03-28 16-19-51\2019_030000005.D)
mAU
350
300
250
200
150
100
50
0
12.884
2
4
6
8
10
12
14
16
18
min


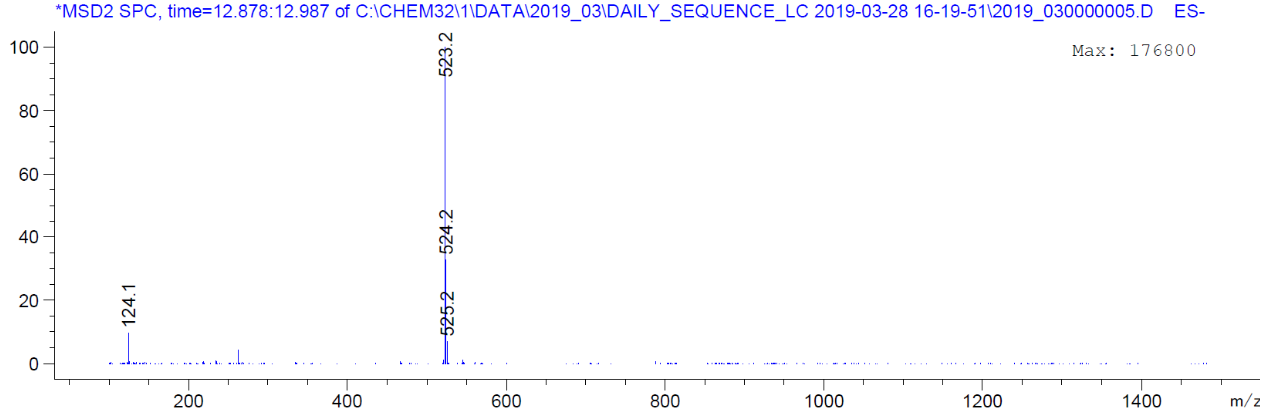

*MSD2 SPC, time=12.878:12.987 of C:\CHEM32\1\DATA\2019_03\DAILY_SEQUENCE_LC 2019-03-28 16-19-51\2019_030000005.D ES-
Max: 176800
100
80
60
40
20
0
523.2
524.2
525.2
124.1
200
400
600
800
1000
1200
1400
m/z

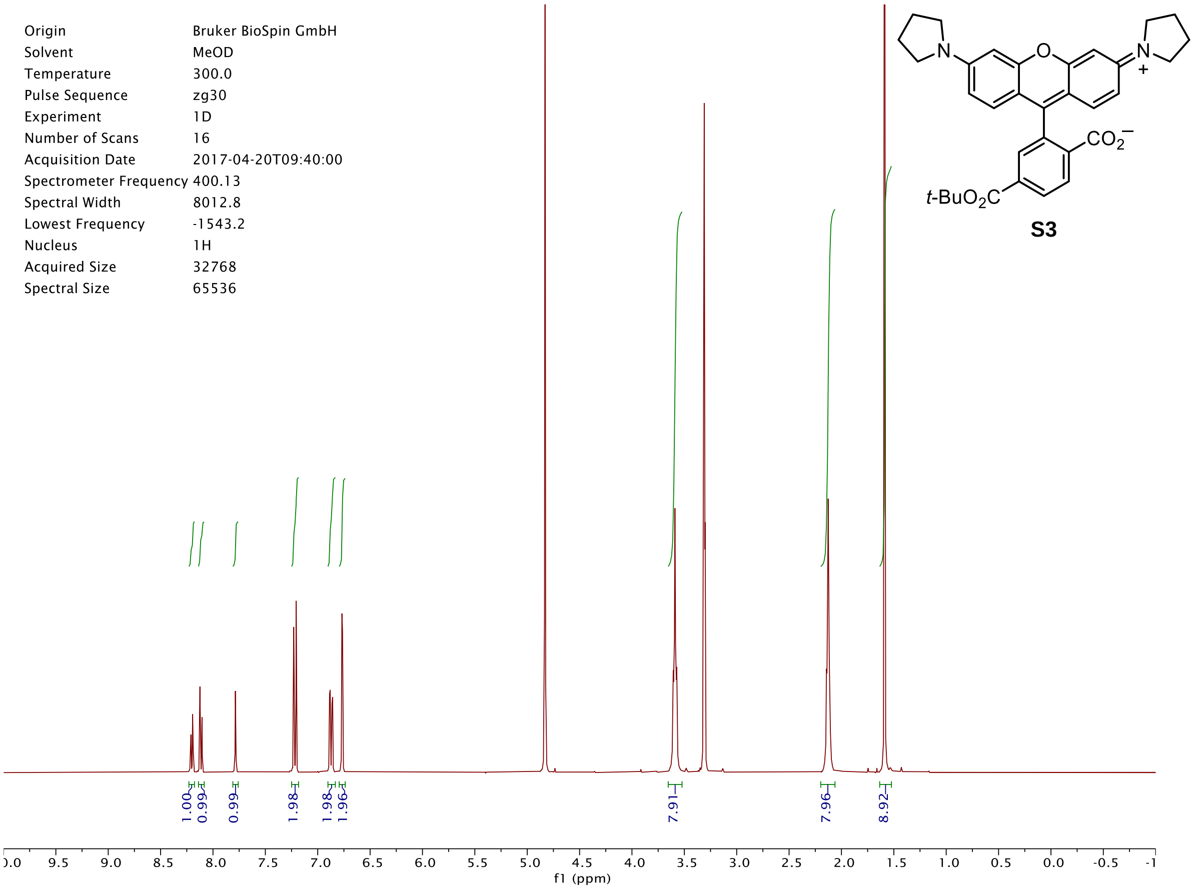
t-BuO2C
CO2−
N
O
N
+
Origin Bruker BioSpin GmbH
Solvent MeOD
Temperature 300.0
Pulse Sequence zg30
Experiment 1D
Number of Scans 16
Acquisition Date 2017-04-20T09:40:00
Spectrometer Frequency 400.13
Spectral Width 8012.8
Lowest Frequency -1543.2
Nucleus 1H
Acquired Size 32768
Spectral Size 65536
1.00
0.99
0.99
1.98
1.98
1.96
7.91
7.96
8.92
f1 (ppm)

**S3**

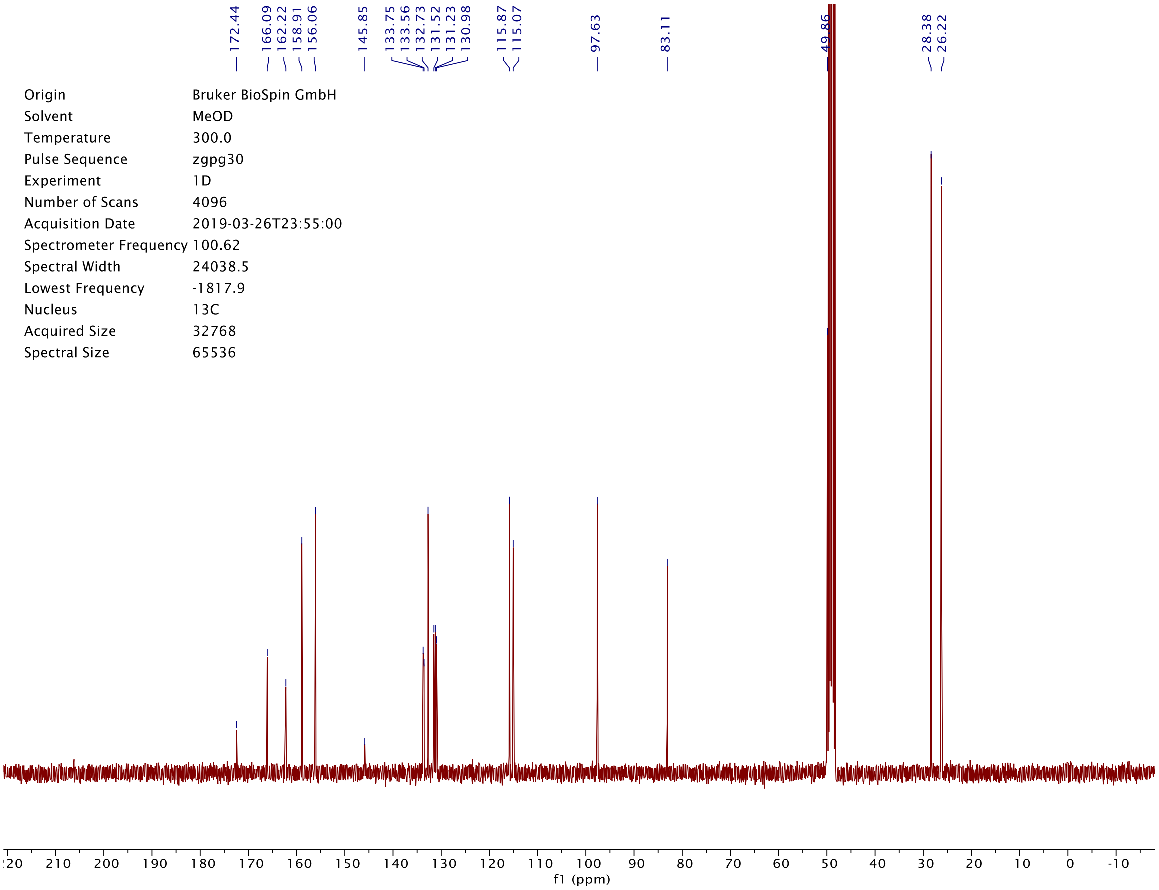
172.44
166.09
162.22
158.91
156.06
145.85
133.75
133.56
132.73
131.52
131.23
130.98
115.87
115.07
97.63
83.11
49.86
28.38
26.22
Origin Bruker BioSpin GmbH
Solvent MeOD
Temperature 300.0
Pulse Sequence zgpg30
Experiment 1D
Number of Scans 4096
Acquisition Date 2019-03-26T23:55:00
Spectrometer Frequency 100.62
Spectral Width 24038.5
Lowest Frequency -1817.9
Nucleus 13C
Acquired Size 32768
Spectral Size 65536
f1 (ppm)

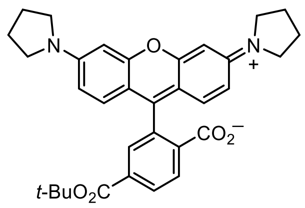
N
O
N
+
CO2−
t-BuO2C

**S3**

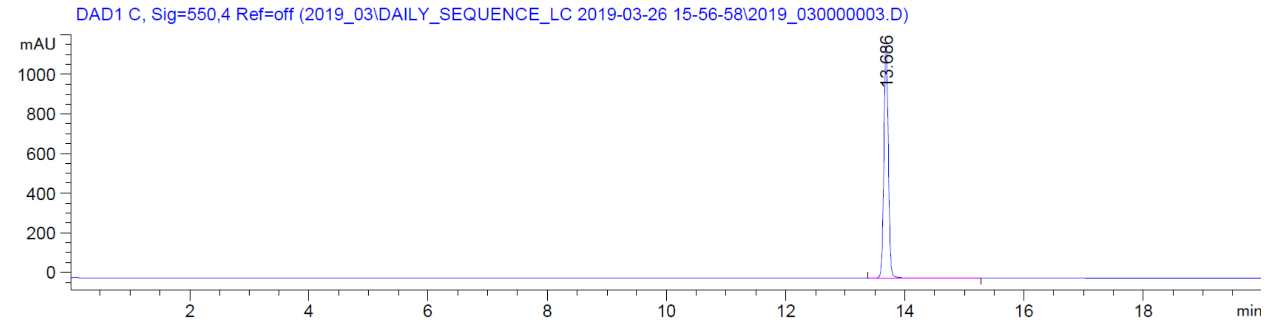
DAD1 C, Sig=550,4 Ref=off (2019_03\DAILY_SEQUENCE_LC 2019-03-26 15-56-58\2019_030000003.D)
mAU
1000
800
600
400
200
0
13.686
2
4
6
8
10
12
14
16
18
min

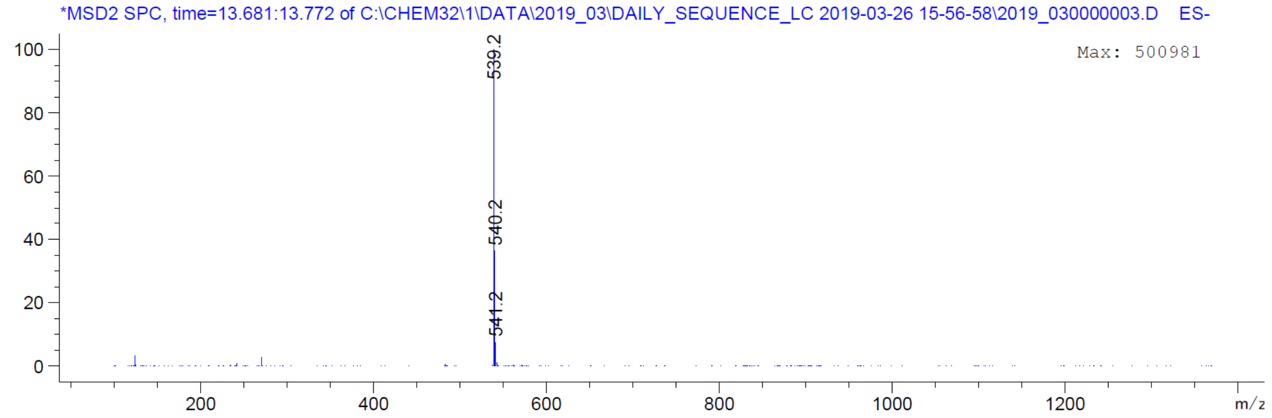
*MSD2 SPC, time=13.681:13.772 of C:\CHEM32\1\DATA\2019_03\DAILY_SEQUENCE_LC 2019-03-26 15-56-58\2019_030000003.D ES-
Max: 500981
100
80
60
40
20
0
539.2
540.2
541.2
200
400
600
800
1000
1200
m/z

| | |
|---|---|
| Origin | Bruker BioSpin GmbH |
| Solvent | MeOD |
| Temperature | 300.0 |
| Pulse Sequence | zg30 |
| Experiment | 1D |
| Number of Scans | 16 |
| Acquisition Date | 2017-04-20T09:45:00 |
| Spectrometer Frequency | 400.13 |
| Spectral Width | 8012.8 |
| Lowest Frequency | -1543.2 |
| Nucleus | 1H |
| Acquired Size | 32768 |
| Spectral Size | 65536 |

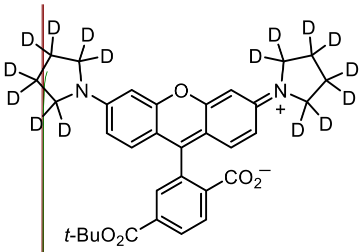


**S4**

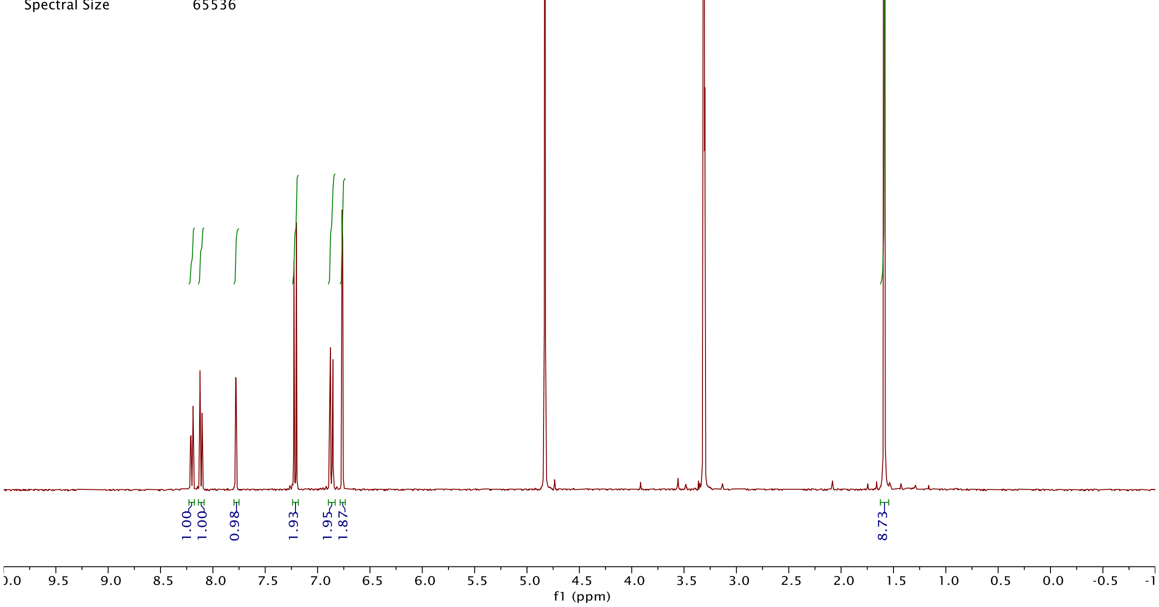


| | |
|---|---|
| Origin | Bruker BioSpin GmbH |
| Solvent | MeOD |
| Temperature | 300.0 |
| Pulse Sequence | zgpg30 |
| Experiment | 1D |
| Number of Scans | 4096 |
| Acquisition Date | 2019-05-15T00:11:00 |
| Spectrometer Frequency | 100.62 |
| Spectral Width | 24038.5 |
| Lowest Frequency | -1817.7 |
| Nucleus | 13C |
| Acquired Size | 32768 |
| Spectral Size | 65536 |

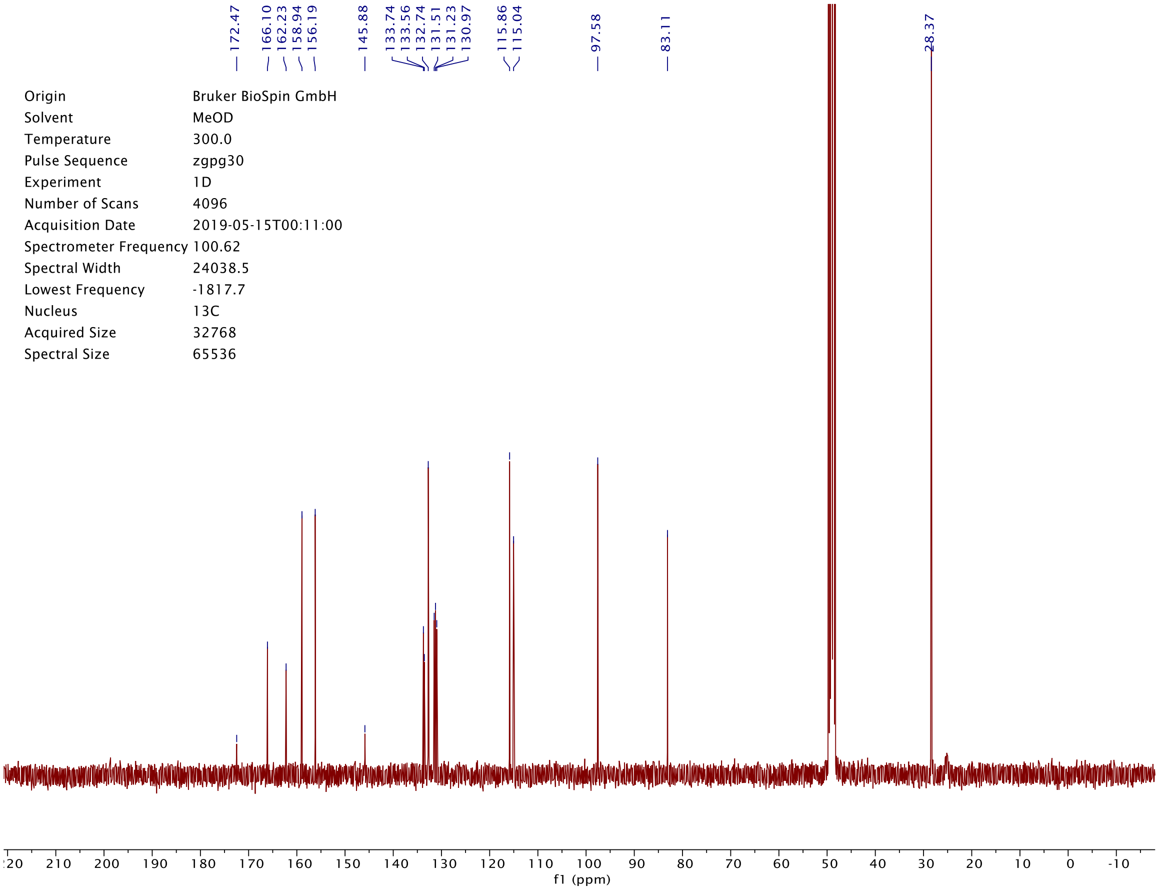

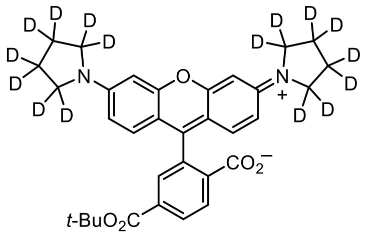

**S4**

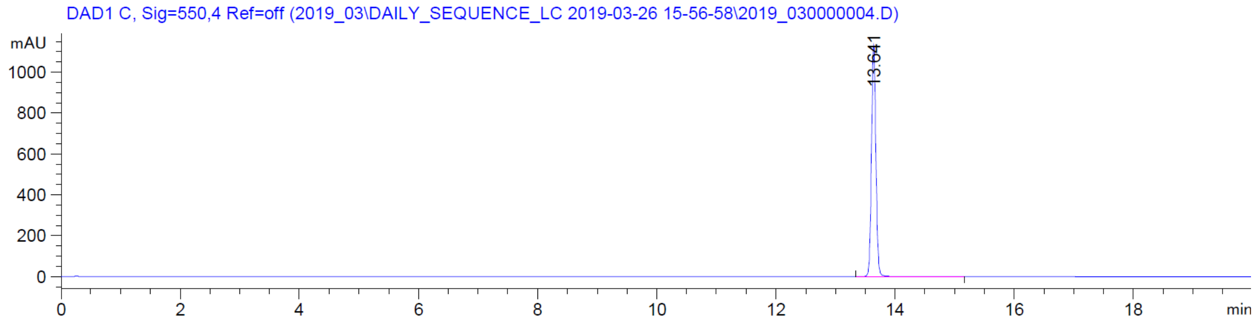

DAD1 C, Sig=550,4 Ref=off (2019_03\DAILY_SEQUENCE_LC 2019-03-26 15-56-58\2019_030000004.D)
mAU
13.641
min


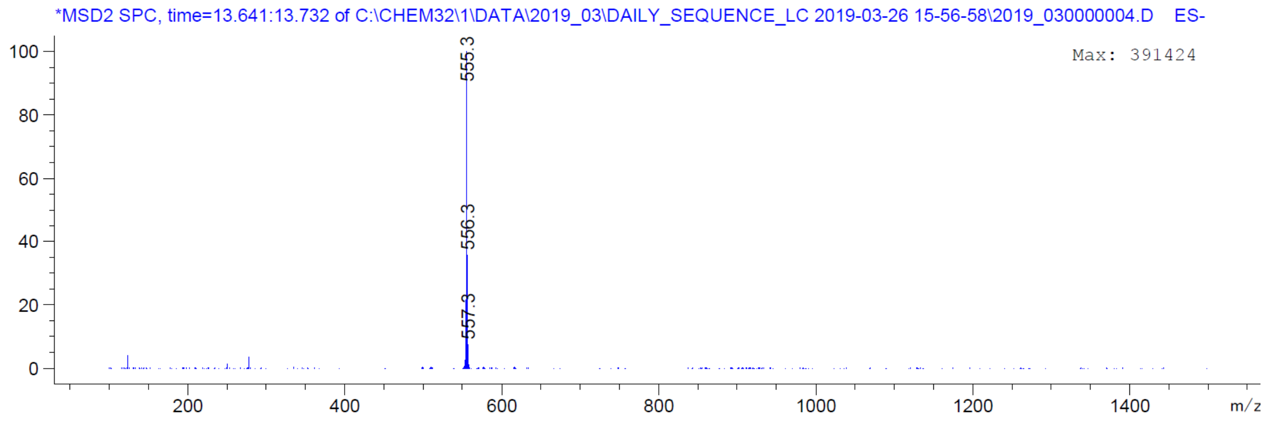

*MSD2 SPC, time=13.641:13.732 of C:\CHEM32\1\DATA\2019_03\DAILY_SEQUENCE_LC 2019-03-26 15-56-58\2019_030000004.D ES-
Max: 391424
555.3
556.3
557.3
m/z

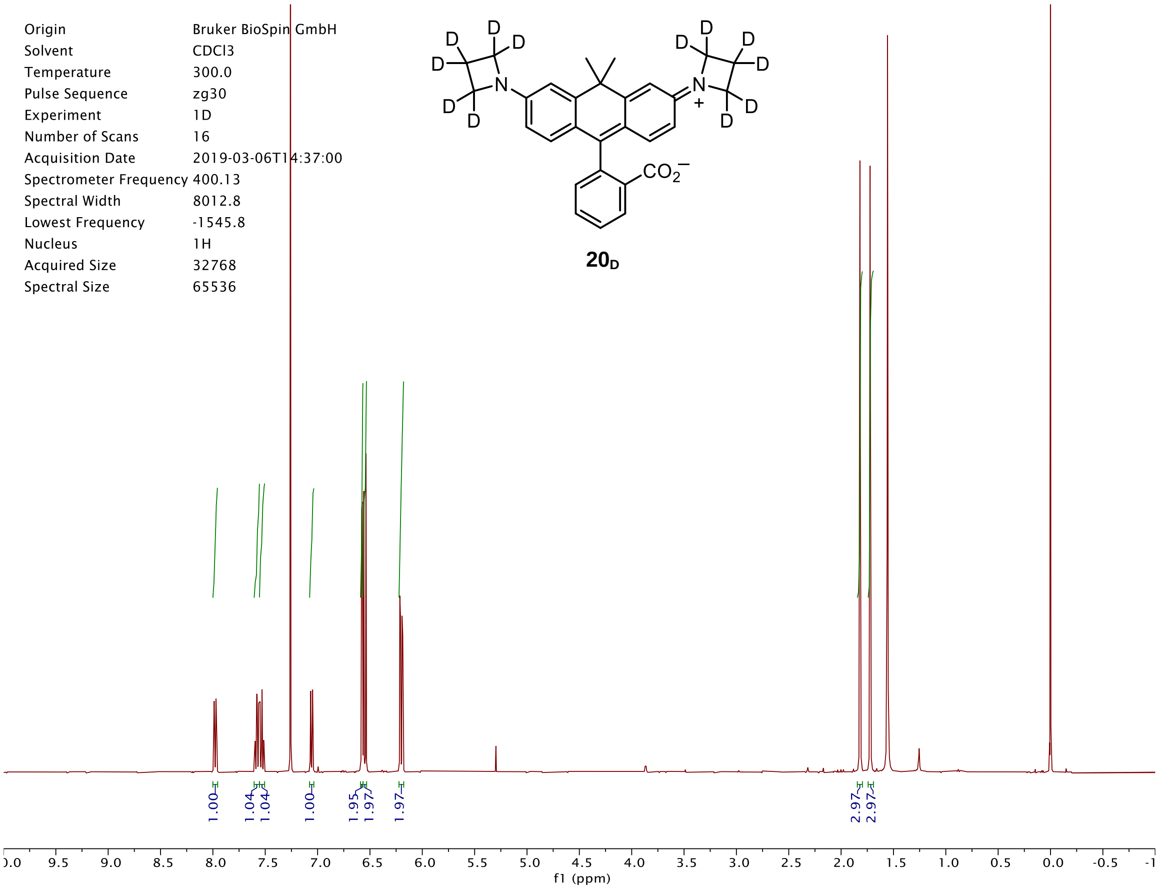
Origin Bruker BioSpin GmbH
Solvent CDCl3
Temperature 300.0
Pulse Sequence zg30
Experiment 1D
Number of Scans 16
Acquisition Date 2019-03-06T14:37:00
Spectrometer Frequency 400.13
Spectral Width 8012.8
Lowest Frequency -1545.8
Nucleus 1H
Acquired Size 32768
Spectral Size 65536
20D
1.00 1.04 1.04 1.00 1.95 1.97 1.97 2.97 2.97
f1 (ppm)

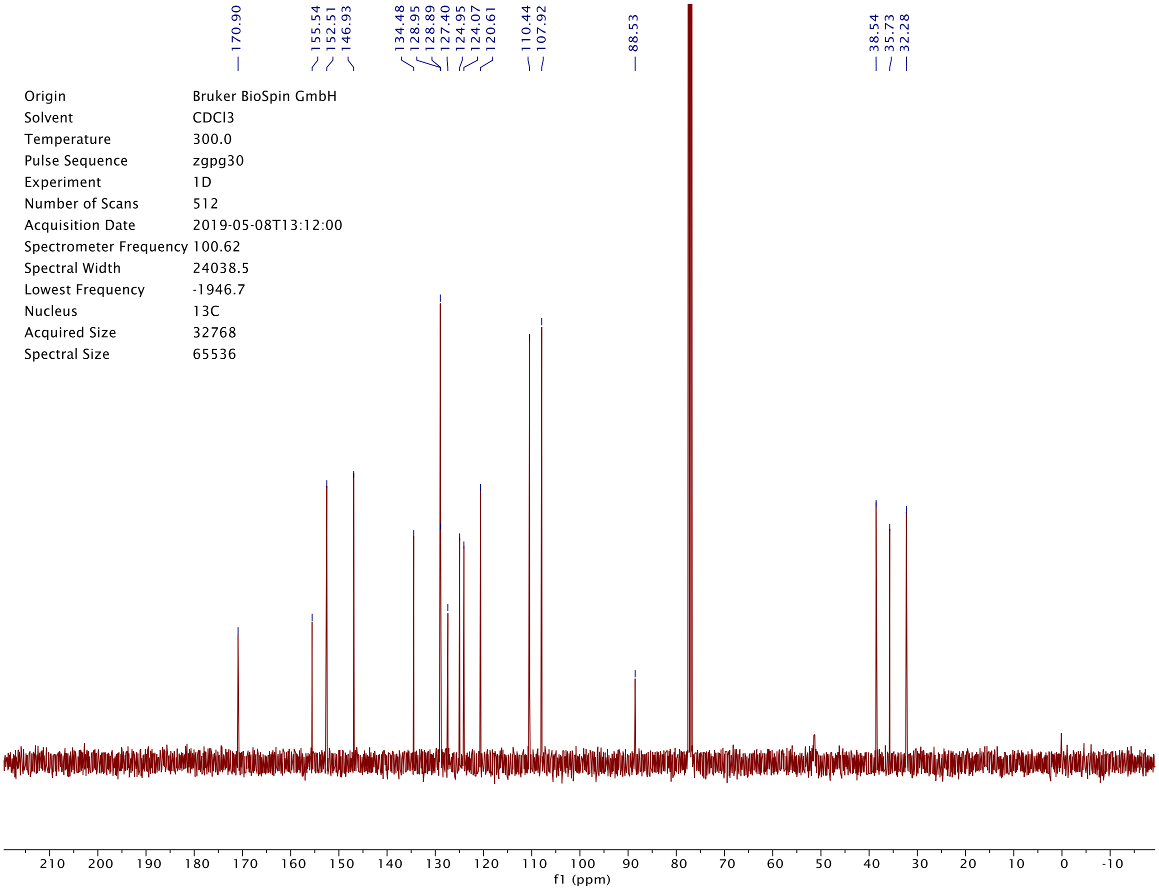
170.90 155.54 152.51 146.93 134.48 128.95 128.89 127.40 124.95 124.07 120.61 110.44 107.92 88.53 38.54 35.73 32.28
Origin Bruker BioSpin GmbH
Solvent CDCl3
Temperature 300.0
Pulse Sequence zgpg30
Experiment 1D
Number of Scans 512
Acquisition Date 2019-05-08T13:12:00
Spectrometer Frequency 100.62
Spectral Width 24038.5
Lowest Frequency -1946.7
Nucleus 13C
Acquired Size 32768
Spectral Size 65536
f1 (ppm)

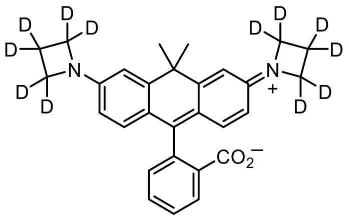
D
D
D
D
N
D
D
D
D
N
+
D
D
D
D
$CO_2^-$

**$20_D$**

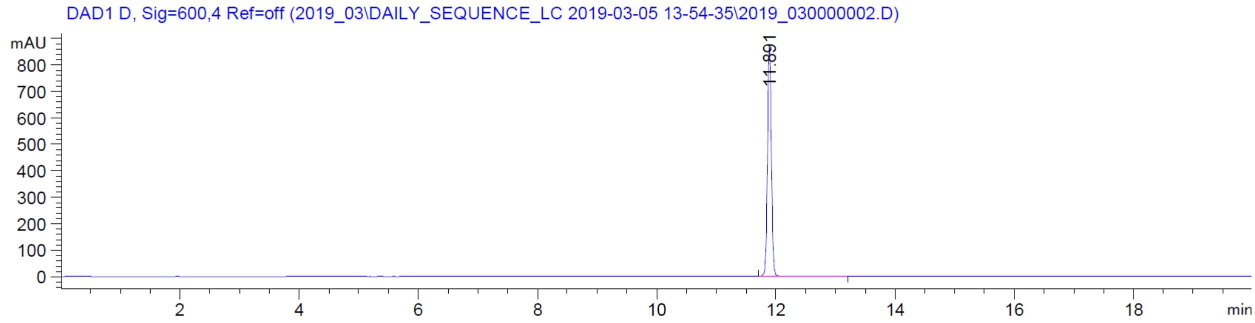
DAD1 D, Sig=600,4 Ref=off (2019_03\DAILY_SEQUENCE_LC 2019-03-05 13-54-35\2019_030000002.D)
mAU
800
700
600
500
400
300
200
100
0
11.891
2
4
6
8
10
12
14
16
18
min

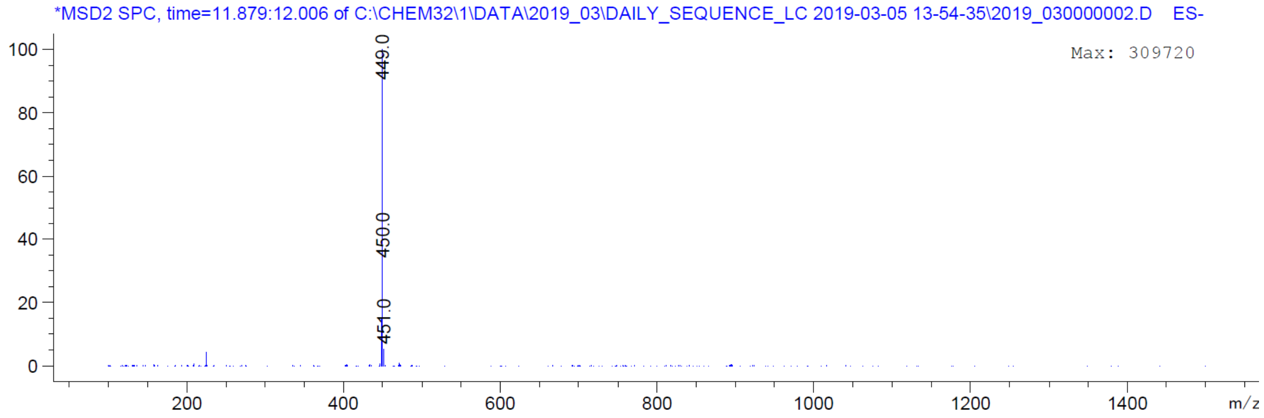
*MSD2 SPC, time=11.879:12.006 of C:\CHEM32\1\DATA\2019_03\DAILY_SEQUENCE_LC 2019-03-05 13-54-35\2019_030000002.D ES-
Max: 309720
100
80
60
40
20
0
449.0
450.0
451.0
200
400
600
800
1000
1200
1400
m/z

| | |
|---|---|
| Origin | Bruker BioSpin GmbH |
| Solvent | CDCl3 |
| Temperature | 300.0 |
| Pulse Sequence | zg30 |
| Experiment | 1D |
| Number of Scans | 16 |
| Acquisition Date | 2019-04-02T10:33:00 |
| Spectrometer Frequency | 400.13 |
| Spectral Width | 8012.8 |
| Lowest Frequency | -1546.4 |
| Nucleus | 1H |
| Acquired Size | 32768 |
| Spectral Size | 65536 |

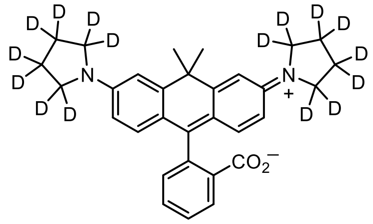


**$21_D$**

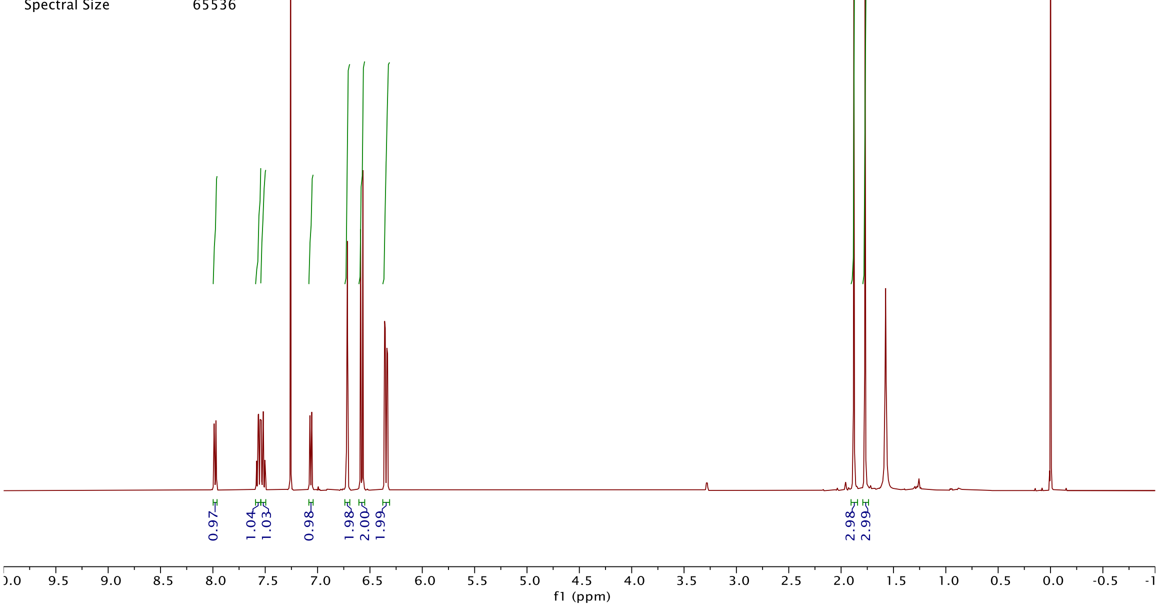


| | |
|---|---|
| Origin | Bruker BioSpin GmbH |
| Solvent | CDCl3 |
| Temperature | 300.0 |
| Pulse Sequence | zgpg30 |
| Experiment | 1D |
| Number of Scans | 256 |
| Acquisition Date | 2019-05-14T12:23:00 |
| Spectrometer Frequency | 100.62 |
| Spectral Width | 24038.5 |
| Lowest Frequency | -1946.3 |
| Nucleus | 13C |
| Acquired Size | 32768 |
| Spectral Size | 65536 |

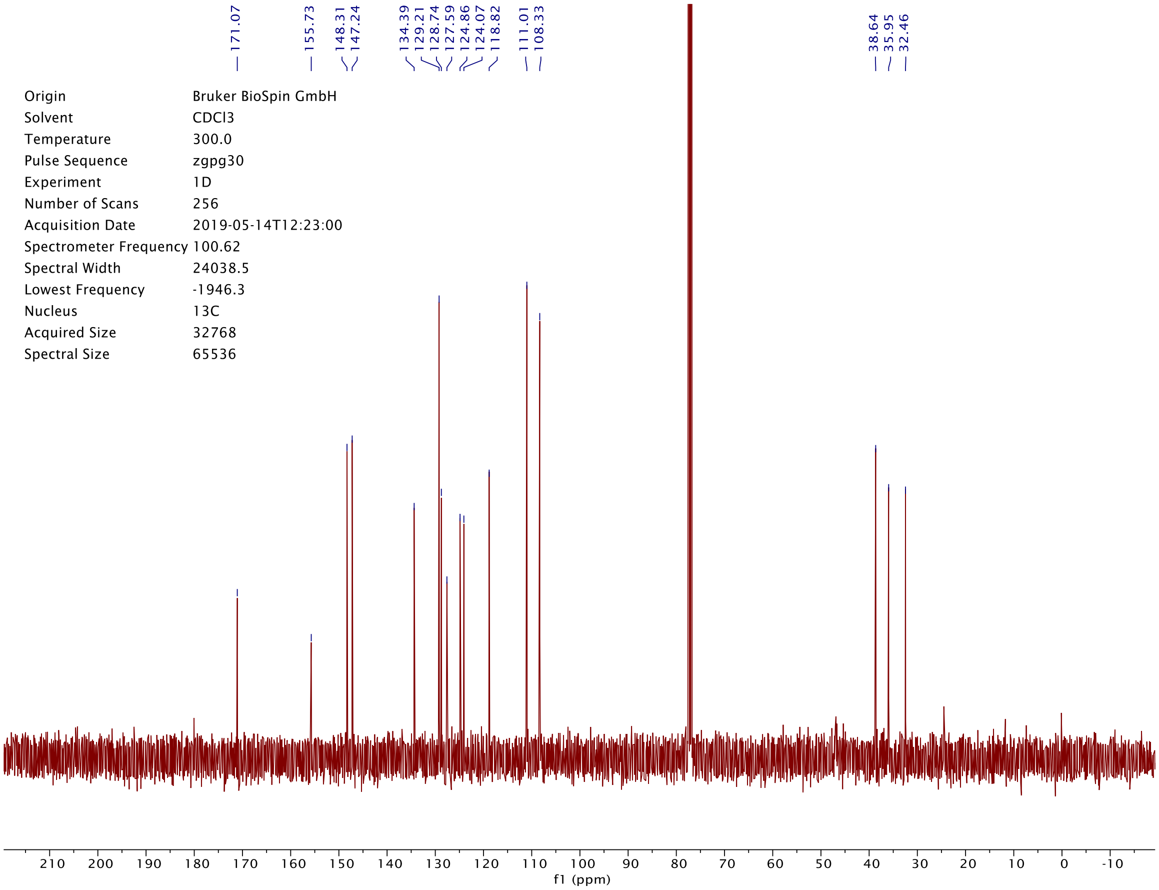

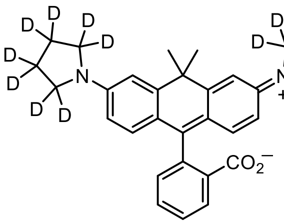

**21$_D$**

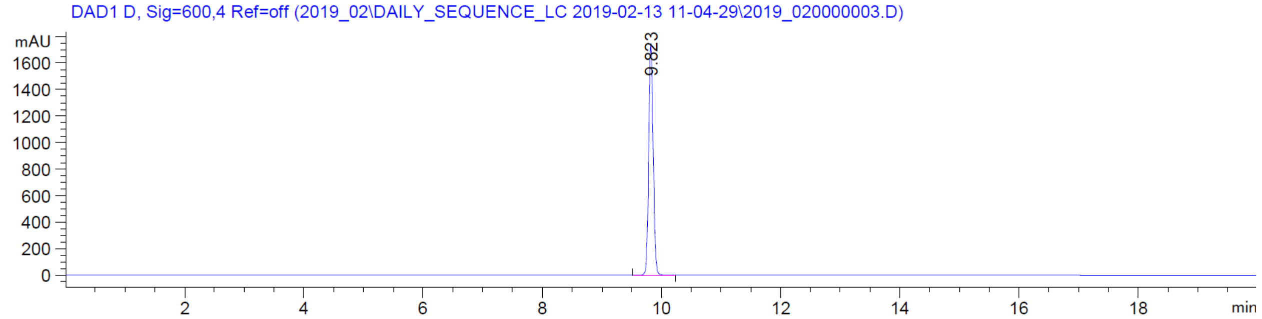

DAD1 D, Sig=600,4 Ref=off (2019_02\DAILY_SEQUENCE_LC 2019-02-13 11-04-29\2019_020000003.D)
mAU
1600
1400
1200
1000
800
600
400
200
0
9.823
2
4
6
8
10
12
14
16
18
min


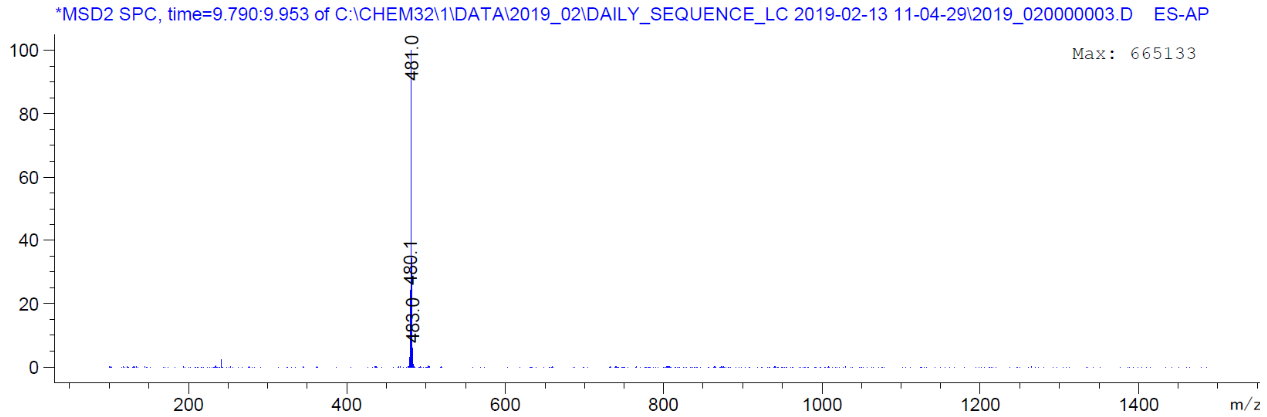

*MSD2 SPC, time=9.790:9.953 of C:\CHEM32\1\DATA\2019_02\DAILY_SEQUENCE_LC 2019-02-13 11-04-29\2019_020000003.D ES-AP
Max: 665133
100
80
60
40
20
0
481.0
480.1
483.0
200
400
600
800
1000
1200
1400
m/z

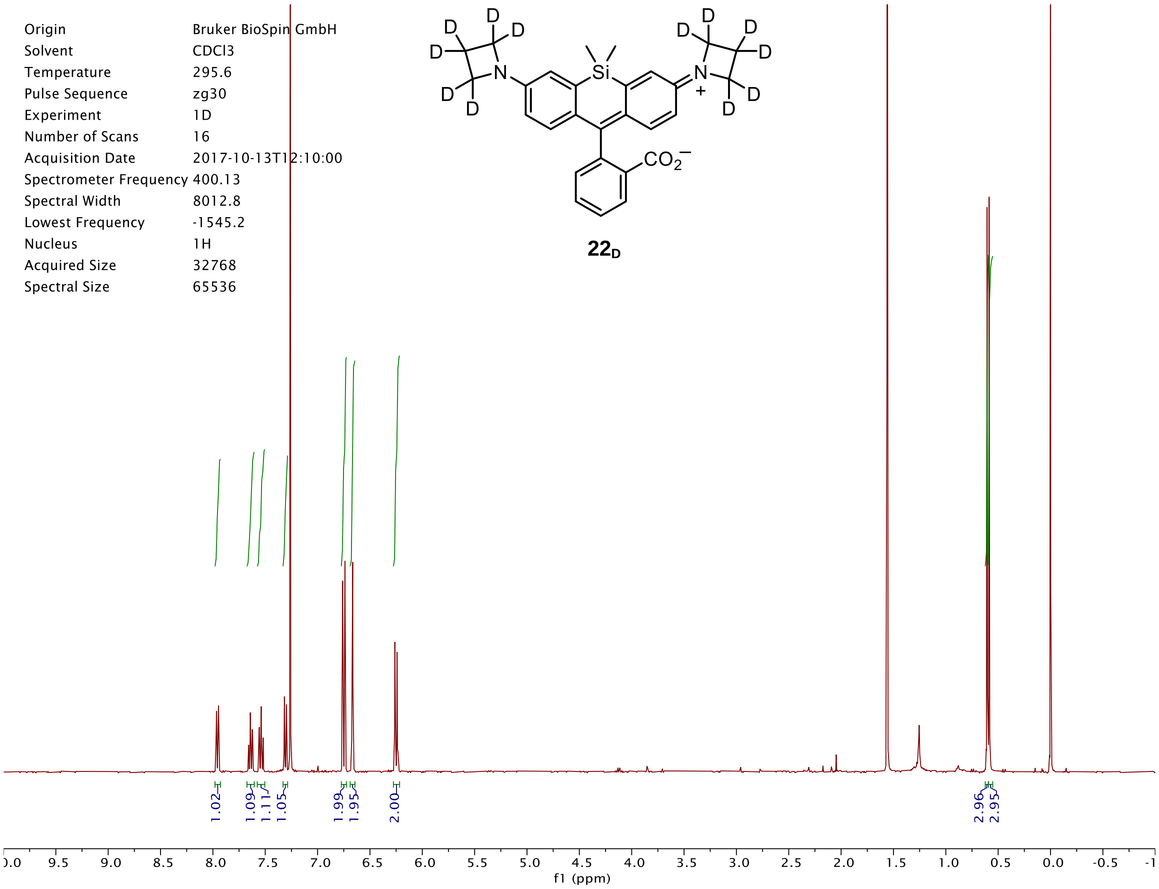

Origin Bruker BioSpin GmbH
Solvent CDCl3
Temperature 295.6
Pulse Sequence zg30
Experiment 1D
Number of Scans 16
Acquisition Date 2017-10-13T12:10:00
Spectrometer Frequency 400.13
Spectral Width 8012.8
Lowest Frequency -1545.2
Nucleus 1H
Acquired Size 32768
Spectral Size 65536
1.02
1.09
1.11
1.05
1.99
1.95
2.00
2.96
2.95
f1 (ppm)
D
Si
N
CO2−
+


**$22_D$**

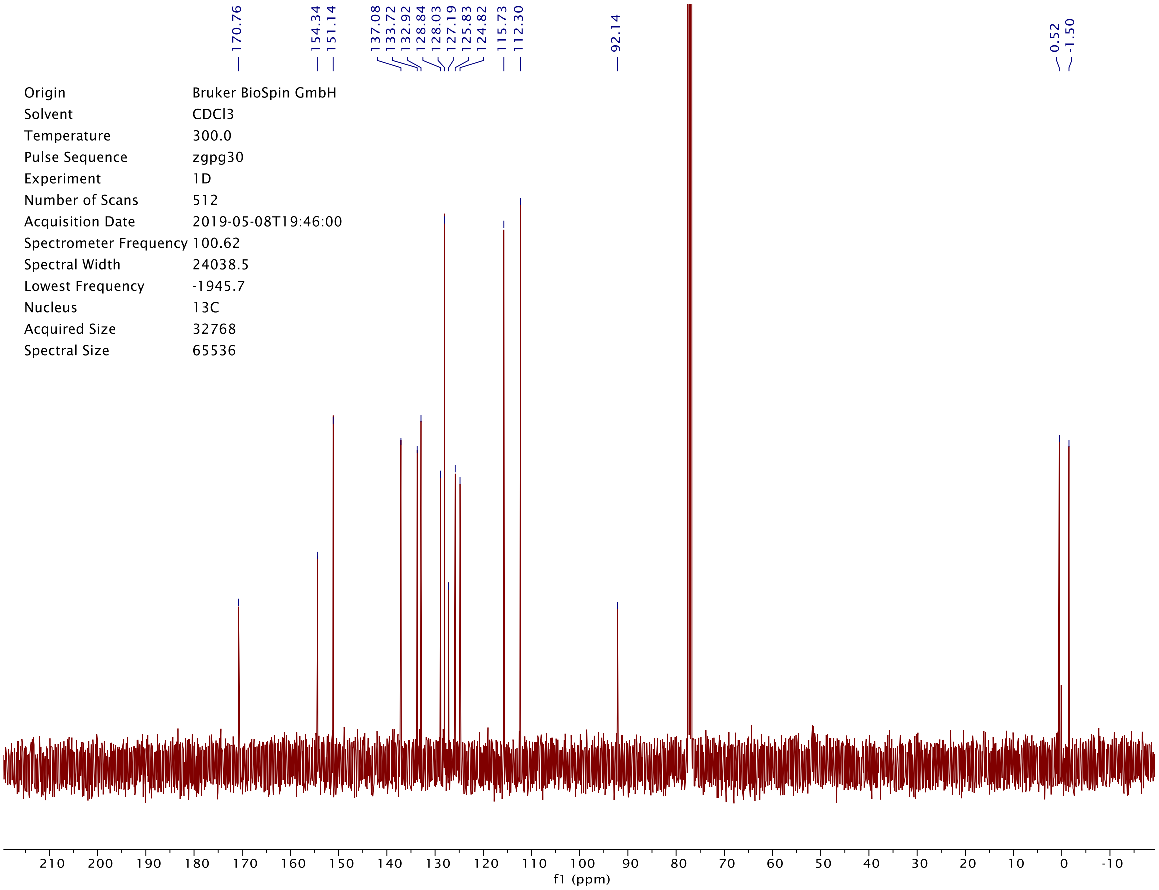

170.76
154.34
151.14
137.08
133.72
132.92
128.84
128.03
127.19
125.83
124.82
115.73
112.30
92.14
0.52
-1.50
Origin Bruker BioSpin GmbH
Solvent CDCl3
Temperature 300.0
Pulse Sequence zgpg30
Experiment 1D
Number of Scans 512
Acquisition Date 2019-05-08T19:46:00
Spectrometer Frequency 100.62
Spectral Width 24038.5
Lowest Frequency -1945.7
Nucleus 13C
Acquired Size 32768
Spectral Size 65536
f1 (ppm)

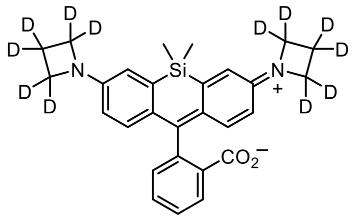

**$22_D$**

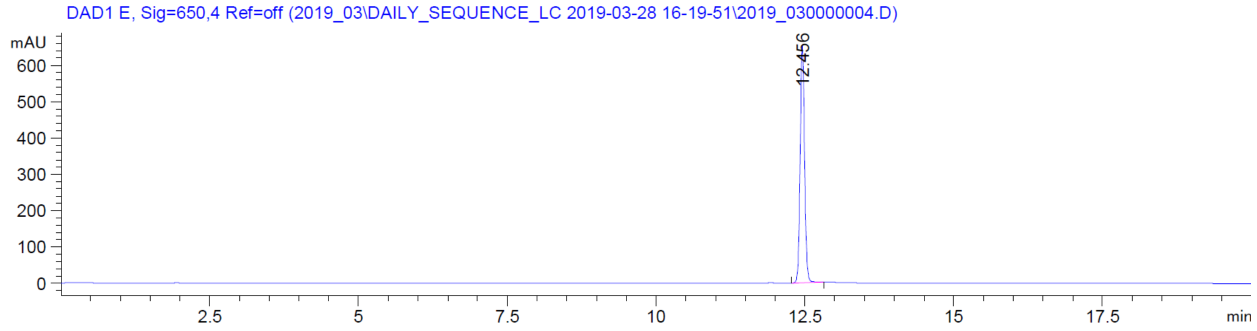
DAD1 E, Sig=650,4 Ref=off (2019_03\DAILY_SEQUENCE_LC 2019-03-28 16-19-51\2019_030000004.D)
mAU
12.456
min

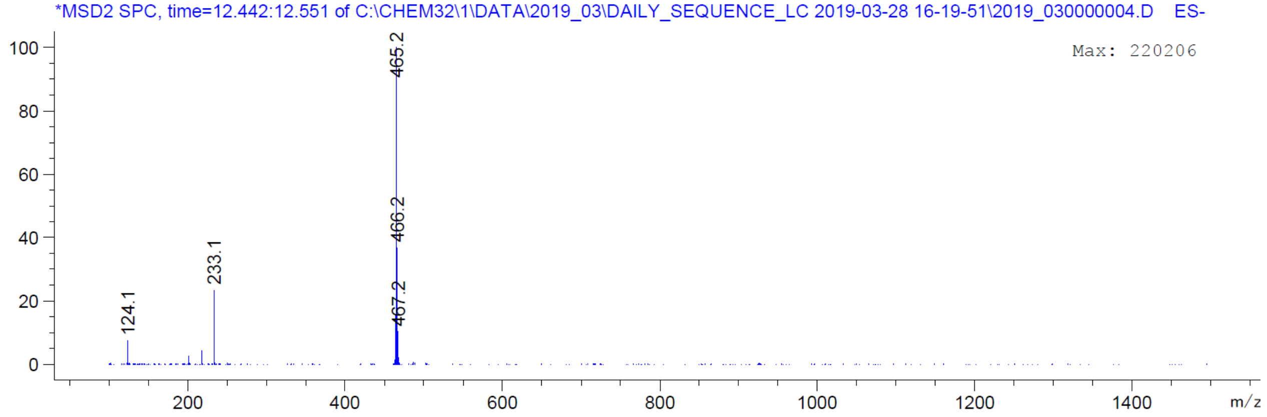
*MSD2 SPC, time=12.442:12.551 of C:\CHEM32\1\DATA\2019_03\DAILY_SEQUENCE_LC 2019-03-28 16-19-51\2019_030000004.D ES-
Max: 220206
465.2
466.2
467.2
233.1
124.1
m/z

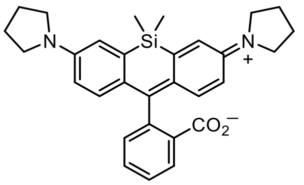


**23**

| | |
|---|---|
| Origin | Bruker BioSpin GmbH |
| Solvent | CDCl3 |
| Temperature | 295.5 |
| Pulse Sequence | zg30 |
| Experiment | 1D |
| Number of Scans | 16 |
| Acquisition Date | 2013-05-24T09:18:00 |
| Spectrometer Frequency | 400.13 |
| Spectral Width | 8223.7 |
| Lowest Frequency | -1647.3 |
| Nucleus | 1H |
| Acquired Size | 32768 |
| Spectral Size | 65536 |

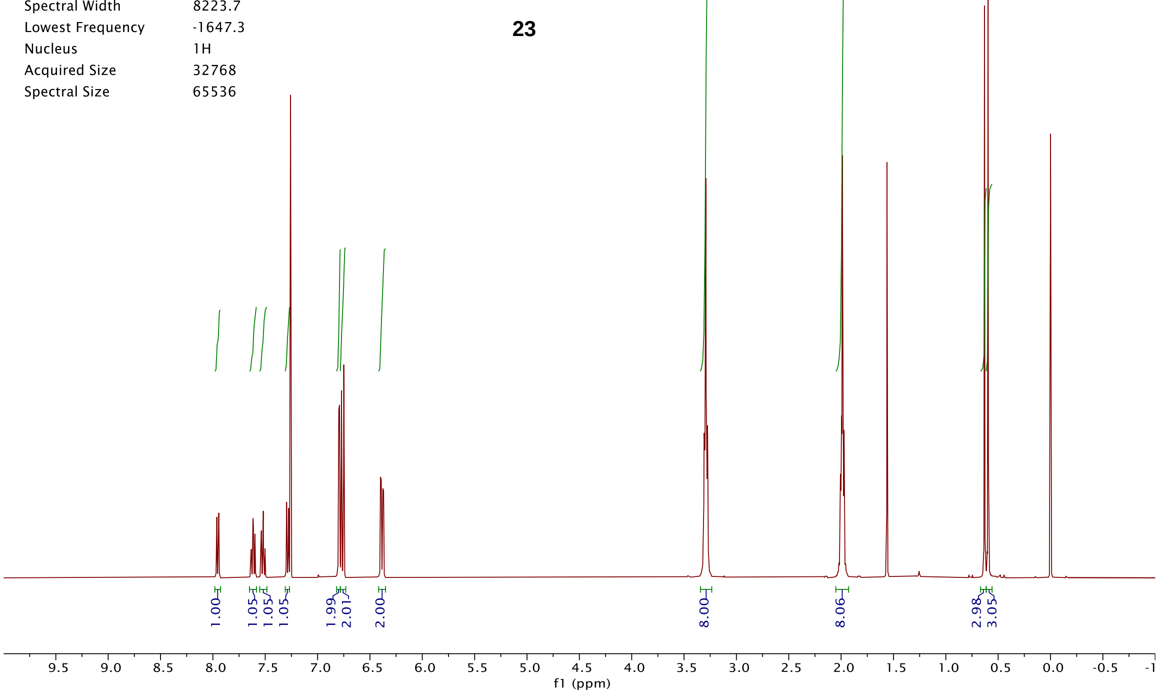


| | |
|---|---|
| Origin | Bruker BioSpin GmbH |
| Solvent | CDCl3 |
| Temperature | 296.8 |
| Pulse Sequence | zgpg30 |
| Experiment | 1D |
| Number of Scans | 2048 |
| Acquisition Date | 2013-05-25T00:01:00 |
| Spectrometer Frequency | 100.62 |
| Spectral Width | 24038.5 |
| Lowest Frequency | -1946.3 |
| Nucleus | 13C |
| Acquired Size | 32768 |
| Spectral Size | 65536 |

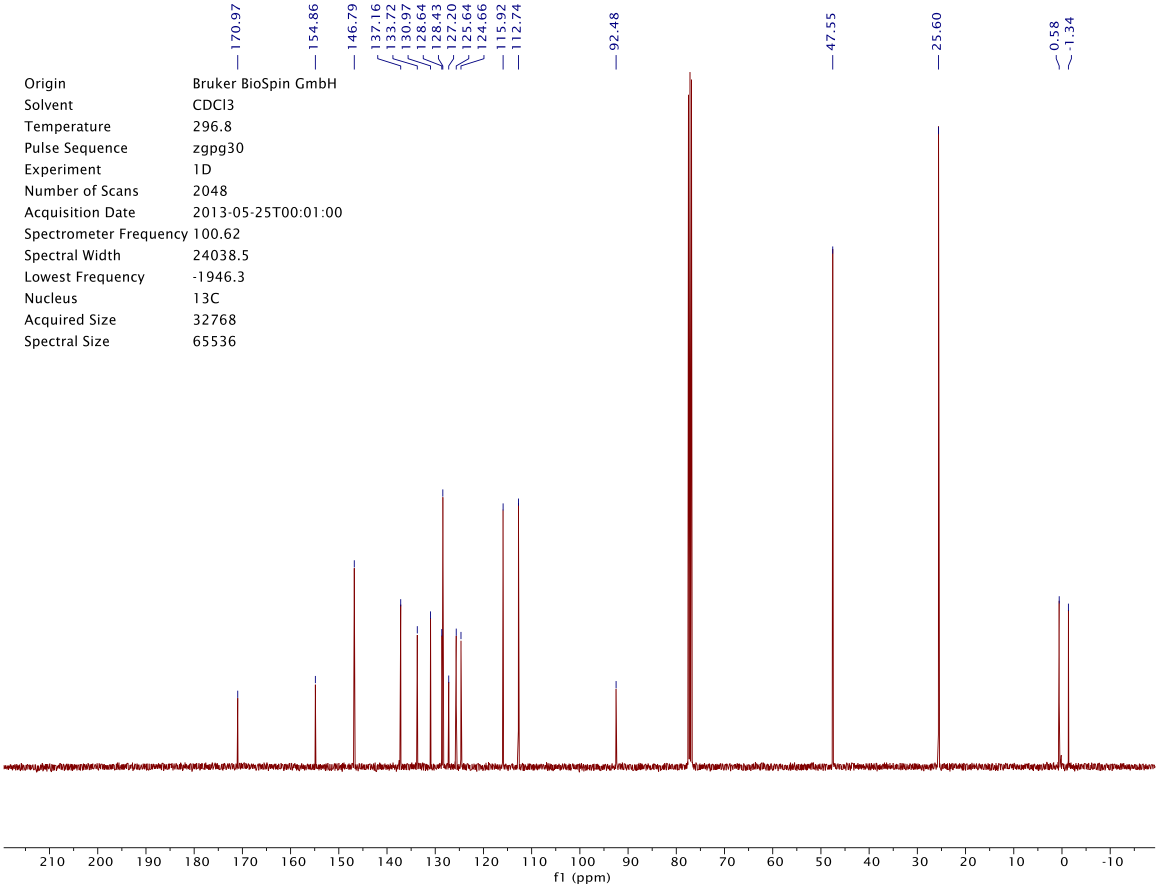

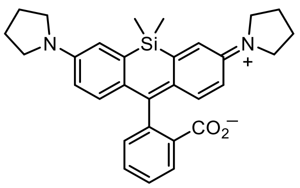

**23**

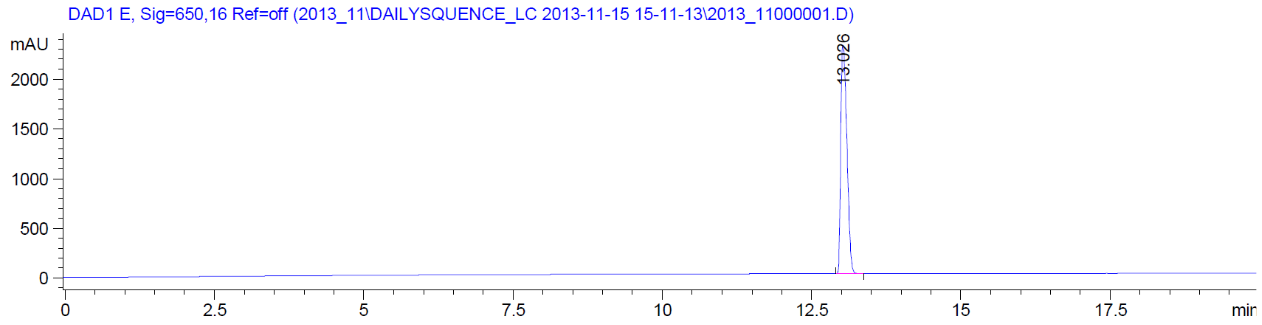

DAD1 E, Sig=650,16 Ref=off (2013_11\DAILYSQUENCE_LC 2013-11-15 15-11-13\2013_11000001.D)
mAU
2000
1500
1000
500
0
13.026
0
2.5
5
7.5
10
12.5
15
17.5
min


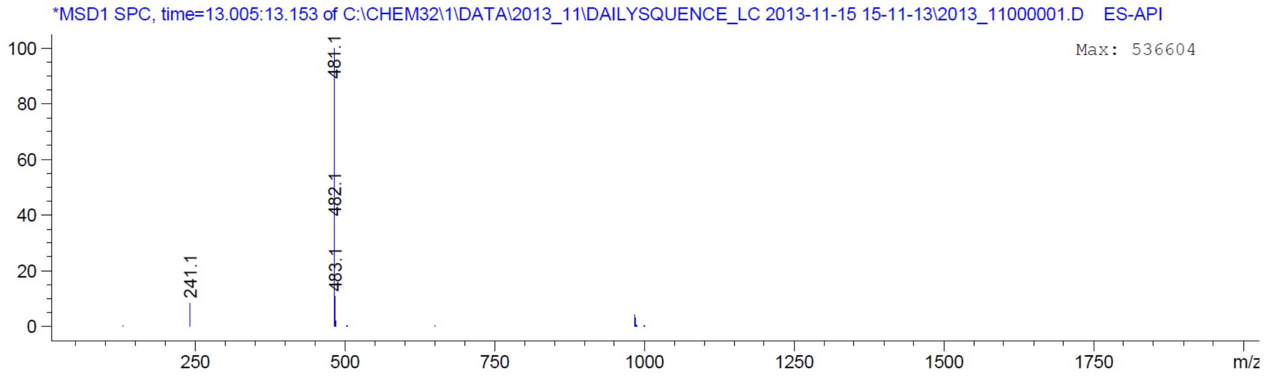

*MSD1 SPC, time=13.005:13.153 of C:\CHEM32\1\DATA\2013_11\DAILYSQUENCE_LC 2013-11-15 15-11-13\2013_11000001.D ES-API
Max: 536604
100
80
60
40
20
0
481.1
482.1
483.1
241.1
250
500
750
1000
1250
1500
1750
m/z

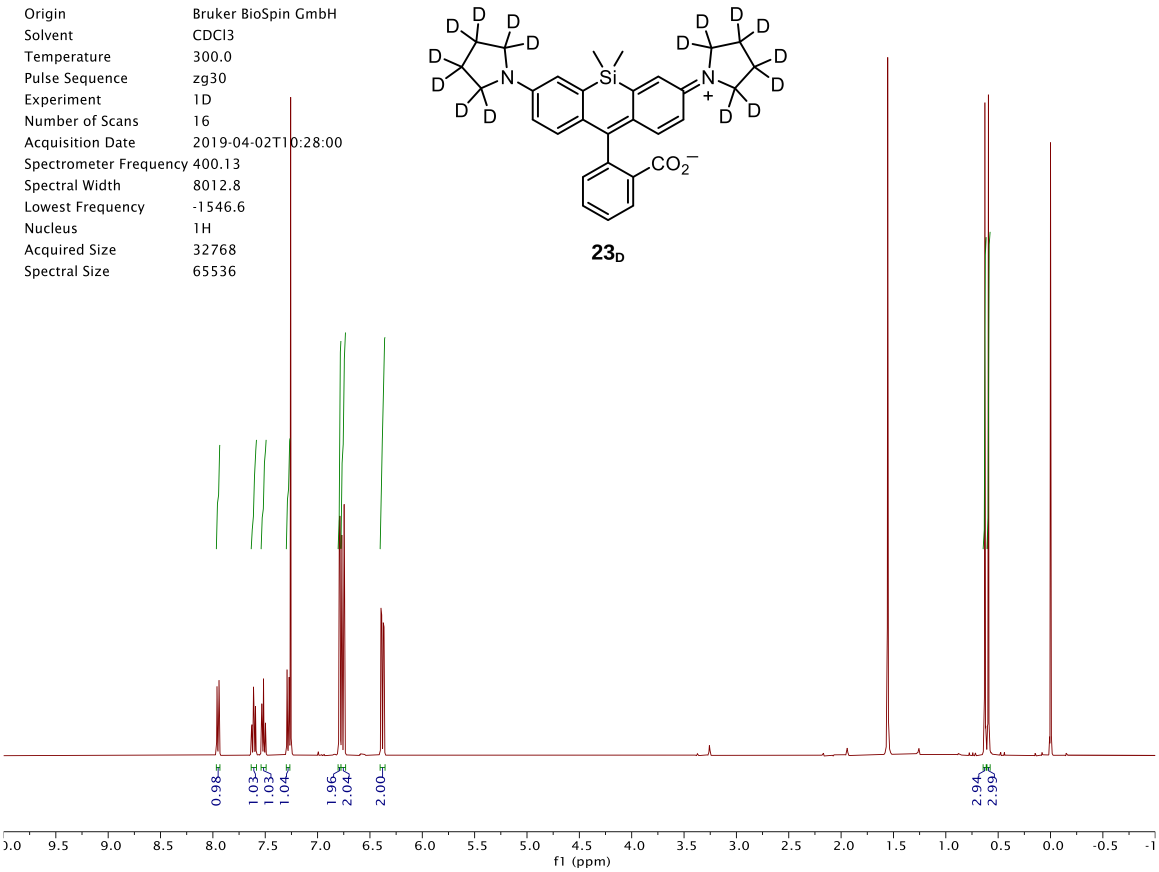
D
D
D
D
D
D
D
D
N
Si
N
+
D
D
D
D
D
D
D
D
CO2−
23D
Origin Bruker BioSpin GmbH
Solvent CDCl3
Temperature 300.0
Pulse Sequence zg30
Experiment 1D
Number of Scans 16
Acquisition Date 2019-04-02T10:28:00
Spectrometer Frequency 400.13
Spectral Width 8012.8
Lowest Frequency -1546.6
Nucleus 1H
Acquired Size 32768
Spectral Size 65536
0.98
1.03
1.03
1.04
1.96
2.04
2.00
2.94
2.99
f1 (ppm)

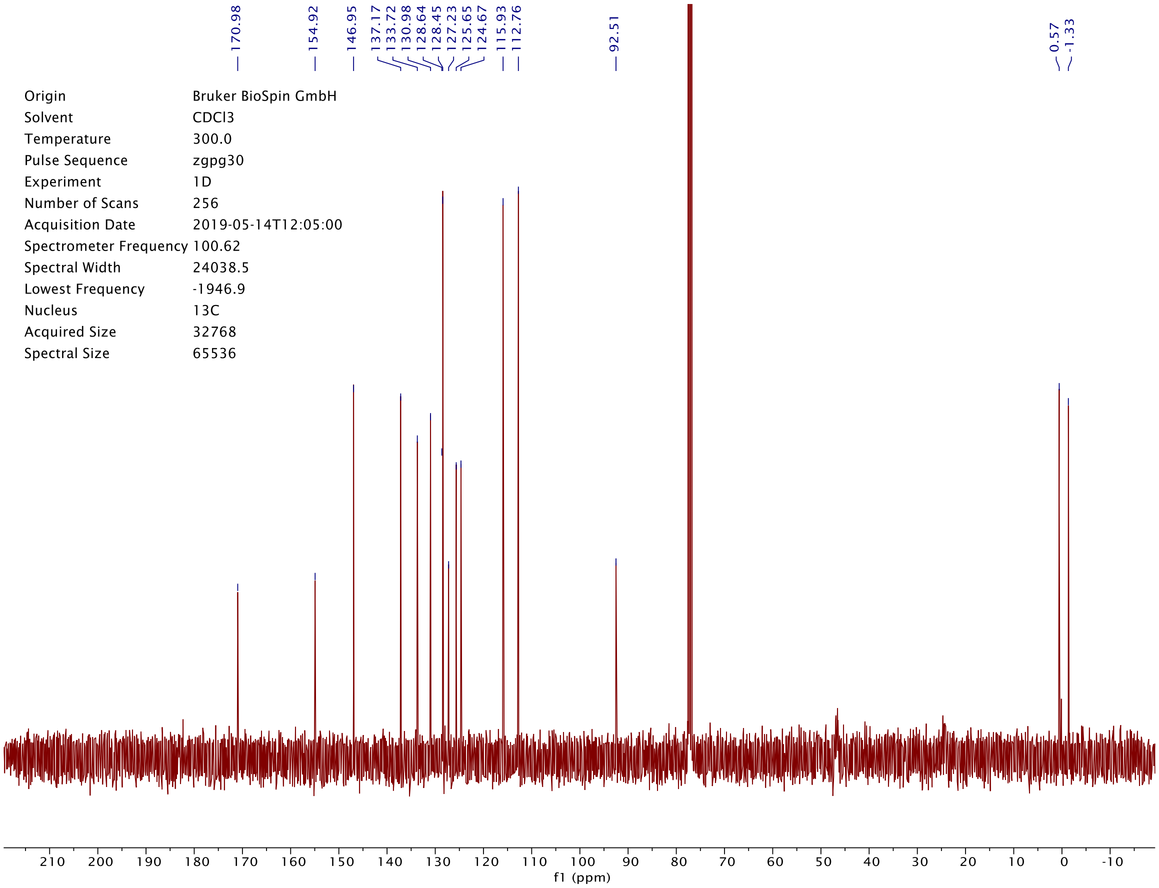
170.98
154.92
146.95
137.17
133.72
130.98
128.64
128.45
127.23
125.65
124.67
115.93
112.76
92.51
0.57
-1.33
Origin Bruker BioSpin GmbH
Solvent CDCl3
Temperature 300.0
Pulse Sequence zgpg30
Experiment 1D
Number of Scans 256
Acquisition Date 2019-05-14T12:05:00
Spectrometer Frequency 100.62
Spectral Width 24038.5
Lowest Frequency -1946.9
Nucleus 13C
Acquired Size 32768
Spectral Size 65536
f1 (ppm)

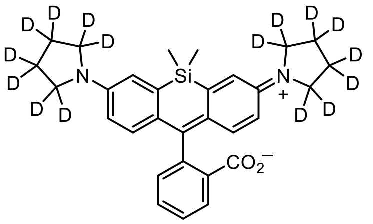

**$23_D$**

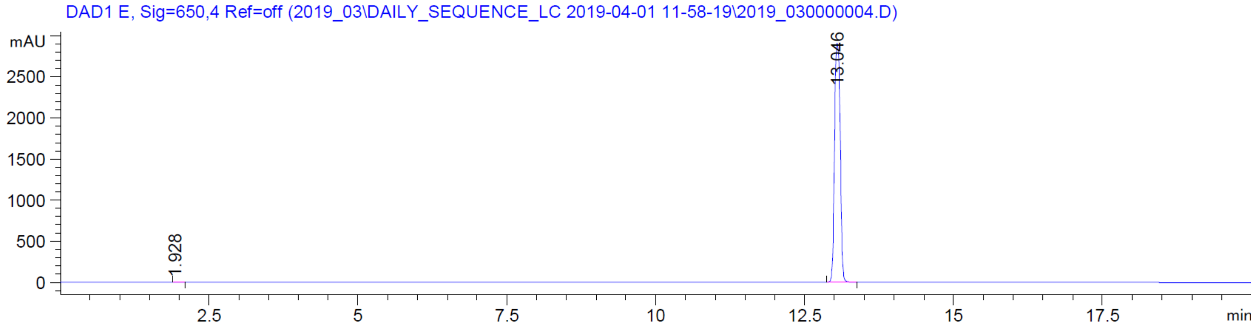
DAD1 E, Sig=650,4 Ref=off (2019_03\DAILY_SEQUENCE_LC 2019-04-01 11-58-19\2019_030000004.D)
mAU
13.046
1.928
min

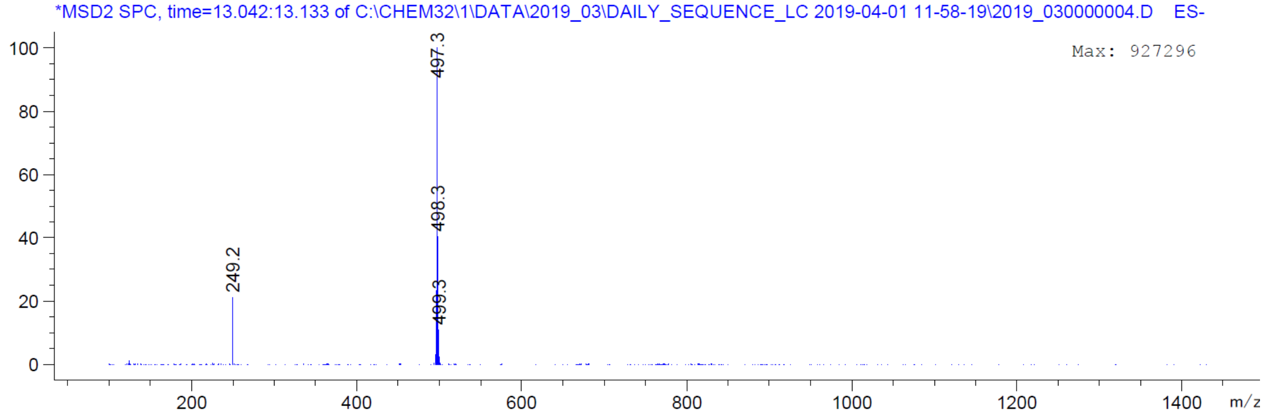
*MSD2 SPC, time=13.042:13.133 of C:\CHEM32\1\DATA\2019_03\DAILY_SEQUENCE_LC 2019-04-01 11-58-19\2019_030000004.D ES-
Max: 927296
497.3
498.3
499.3
249.2
m/z

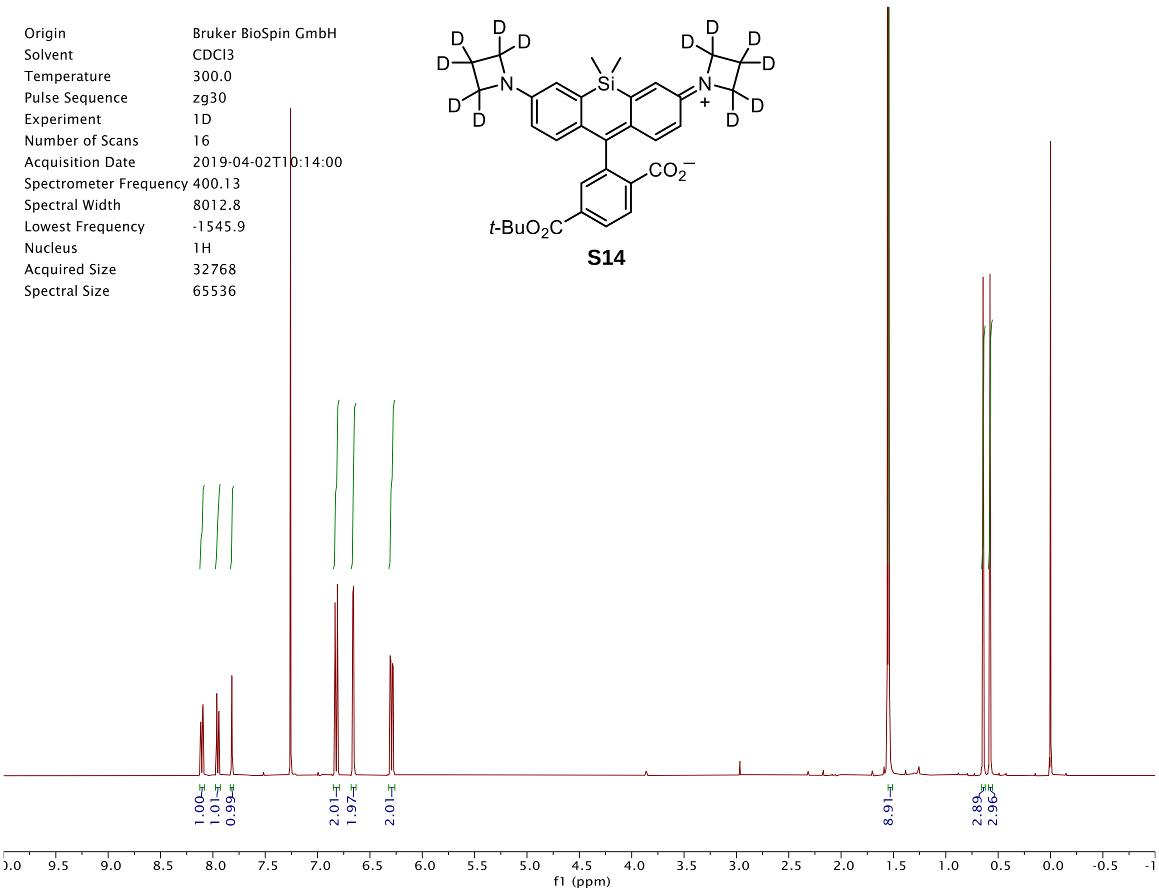
Origin Bruker BioSpin GmbH
Solvent CDCl3
Temperature 300.0
Pulse Sequence zg30
Experiment 1D
Number of Scans 16
Acquisition Date 2019-04-02T10:14:00
Spectrometer Frequency 400.13
Spectral Width 8012.8
Lowest Frequency -1545.9
Nucleus 1H
Acquired Size 32768
Spectral Size 65536
t-BuO2C
CO2−
S14
1.00
1.01
0.99
2.01
1.97
2.01
8.91
2.89
2.96
f1 (ppm)

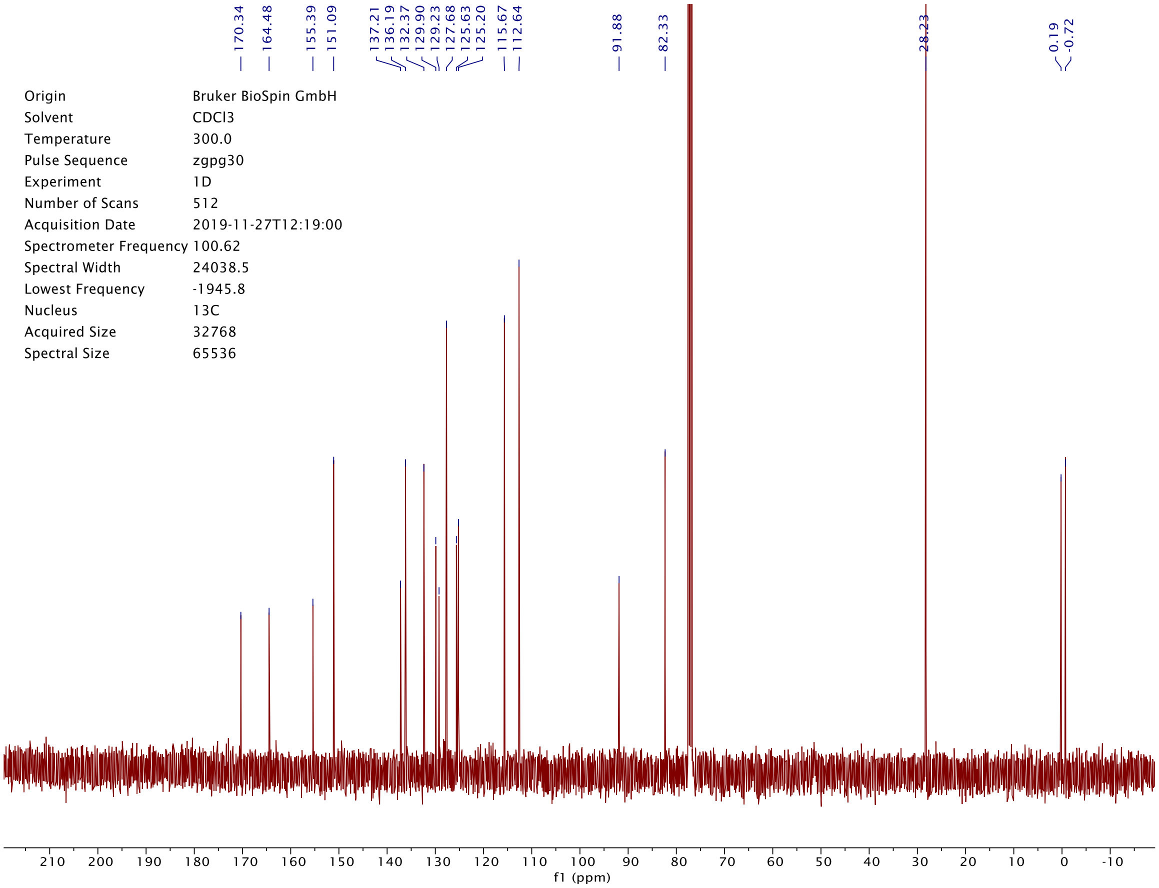
170.34
164.48
155.39
151.09
137.21
136.19
132.37
129.90
129.23
127.68
125.63
125.20
115.67
112.64
91.88
82.33
28.23
0.19
-0.72
Origin Bruker BioSpin GmbH
Solvent CDCl3
Temperature 300.0
Pulse Sequence zgpg30
Experiment 1D
Number of Scans 512
Acquisition Date 2019-11-27T12:19:00
Spectrometer Frequency 100.62
Spectral Width 24038.5
Lowest Frequency -1945.8
Nucleus 13C
Acquired Size 32768
Spectral Size 65536
f1 (ppm)

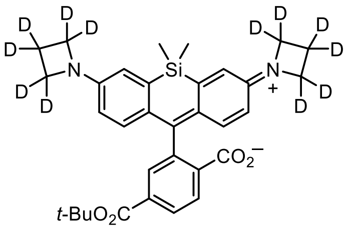
D
D
D
D
D
D
D
D
D
D
D
D
N
Si
N
+
$CO_2^-$
t-BuO2C

**S14**

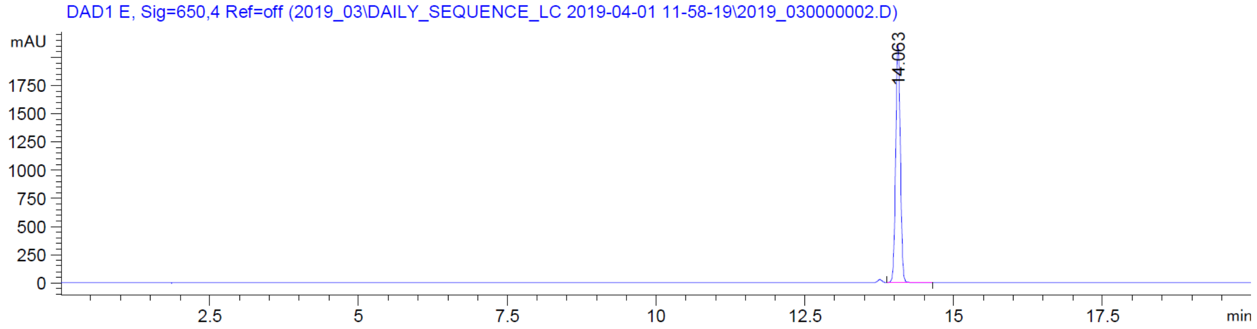
DAD1 E, Sig=650,4 Ref=off (2019_03\DAILY_SEQUENCE_LC 2019-04-01 11-58-19\2019_030000002.D)
mAU
1750
1500
1250
1000
750
500
250
0
14.063
2.5
5
7.5
10
12.5
15
17.5
min

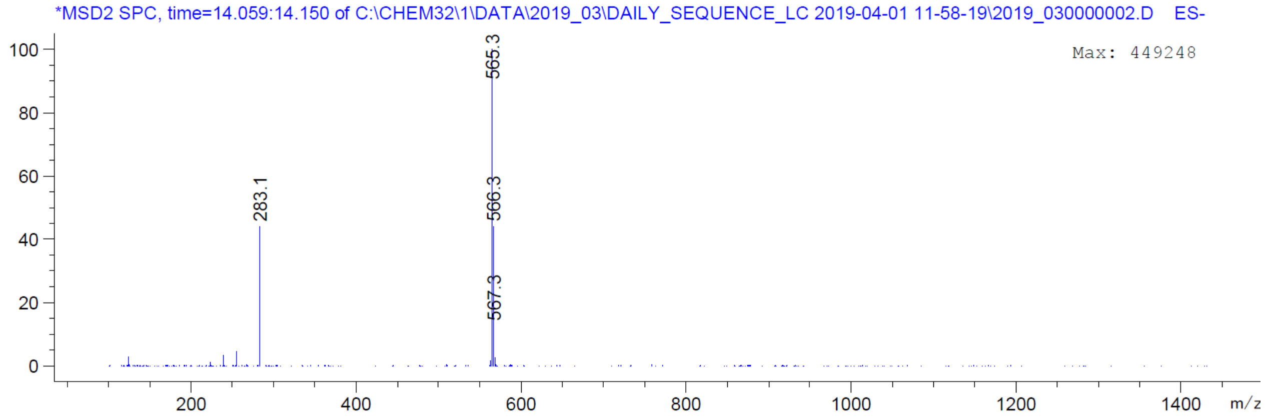
*MSD2 SPC, time=14.059:14.150 of C:\CHEM32\1\DATA\2019_03\DAILY_SEQUENCE_LC 2019-04-01 11-58-19\2019_030000002.D ES-
Max: 449248
100
80
60
40
20
0
565.3
283.1
566.3
567.3
200
400
600
800
1000
1200
1400
m/z

**S15**

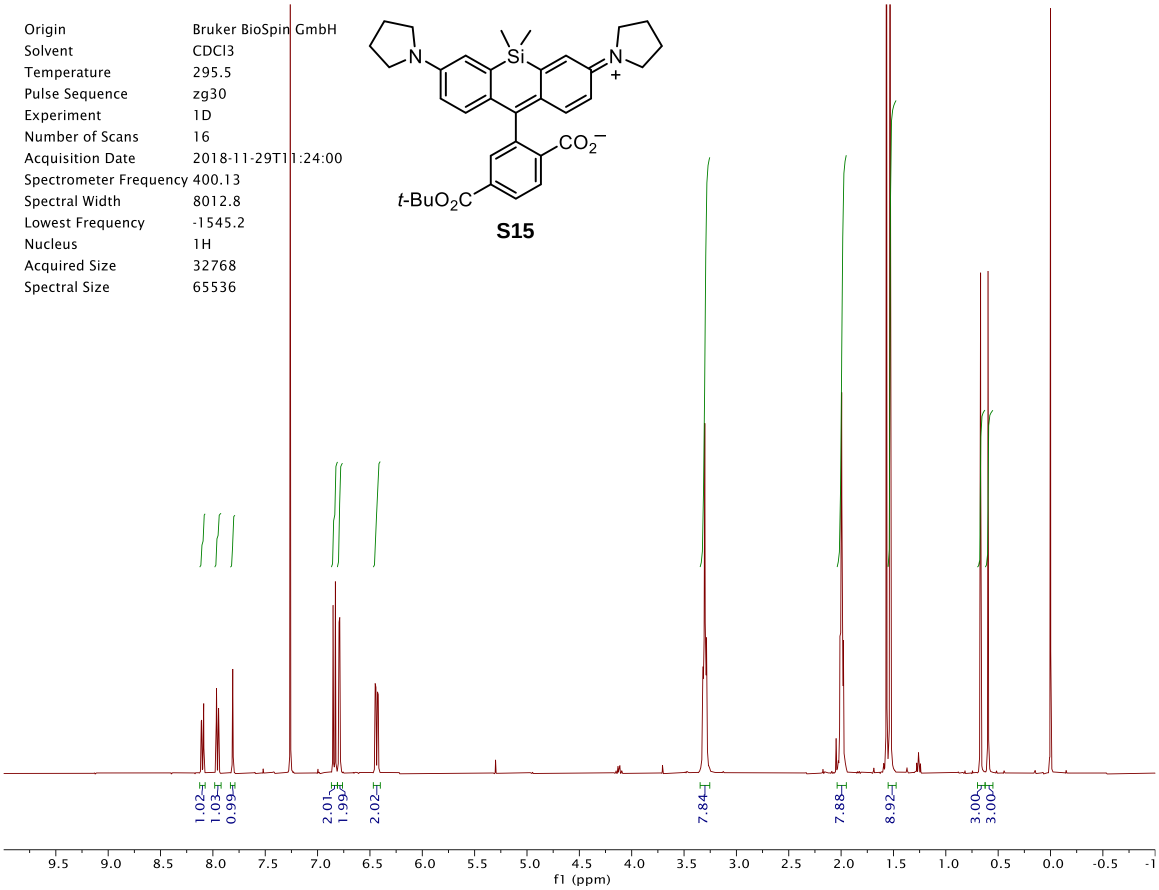

Si
N
N
+
$CO_2^-$
$t$-$BuO_2C$
Origin Bruker BioSpin GmbH
Solvent CDCl3
Temperature 295.5
Pulse Sequence zg30
Experiment 1D
Number of Scans 16
Acquisition Date 2018-11-29T11:24:00
Spectrometer Frequency 400.13
Spectral Width 8012.8
Lowest Frequency -1545.2
Nucleus 1H
Acquired Size 32768
Spectral Size 65536
1.02
1.03
0.99
2.01
1.99
2.02
7.84
7.88
8.92
3.00
3.00
f1 (ppm)


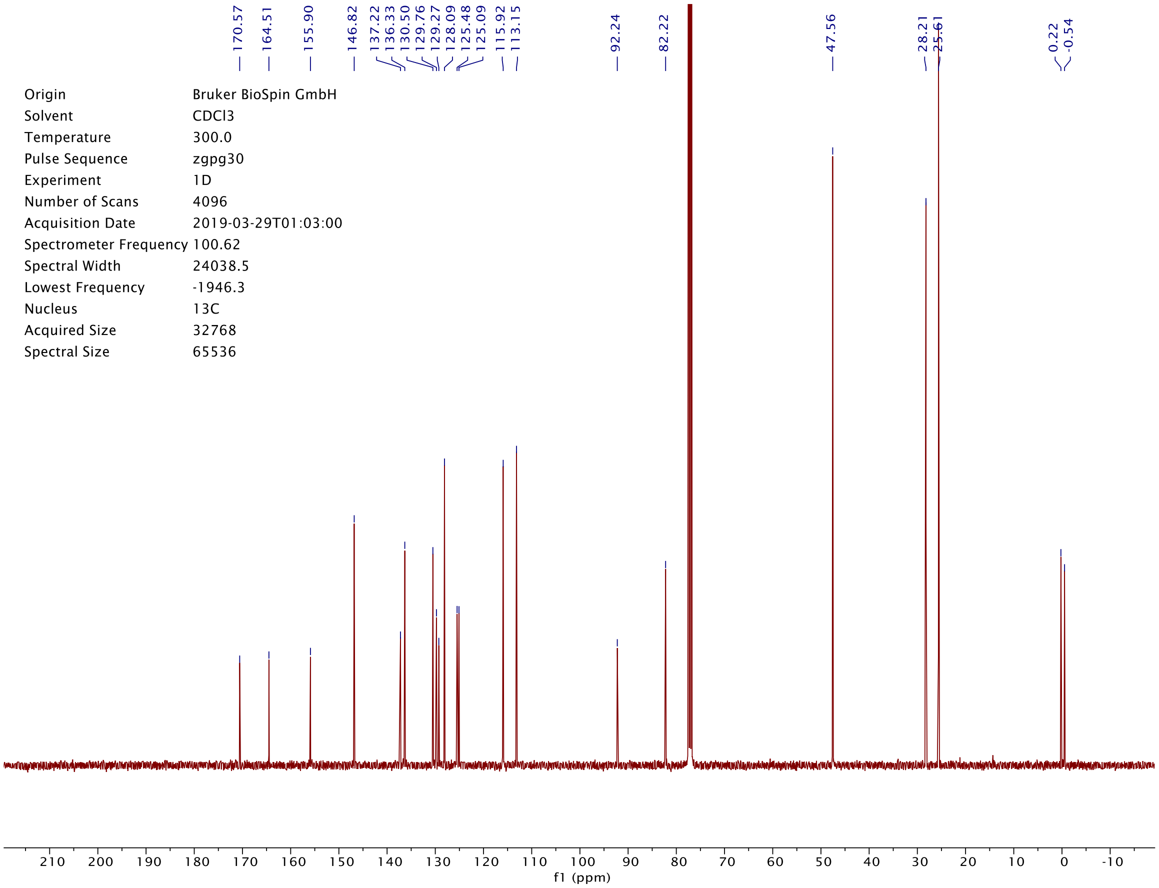

170.57
164.51
155.90
146.82
137.22
136.33
130.50
129.76
129.27
128.09
125.48
125.09
115.92
113.15
92.24
82.22
47.56
28.21
25.61
0.22
-0.54
Origin Bruker BioSpin GmbH
Solvent CDCl3
Temperature 300.0
Pulse Sequence zgpg30
Experiment 1D
Number of Scans 4096
Acquisition Date 2019-03-29T01:03:00
Spectrometer Frequency 100.62
Spectral Width 24038.5
Lowest Frequency -1946.3
Nucleus 13C
Acquired Size 32768
Spectral Size 65536
f1 (ppm)

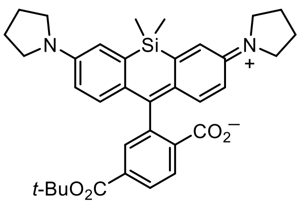
Si
N
N
+
$CO_2^-$
$t$-$BuO_2C$

**S15**

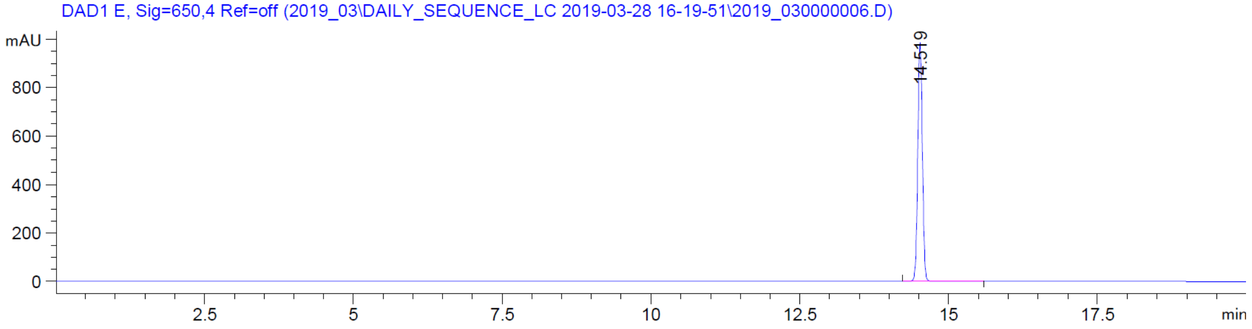
DAD1 E, Sig=650,4 Ref=off (2019_03\DAILY_SEQUENCE_LC 2019-03-28 16-19-51\2019_030000006.D)
mAU
800
600
400
200
0
14.519
2.5
5
7.5
10
12.5
15
17.5
min

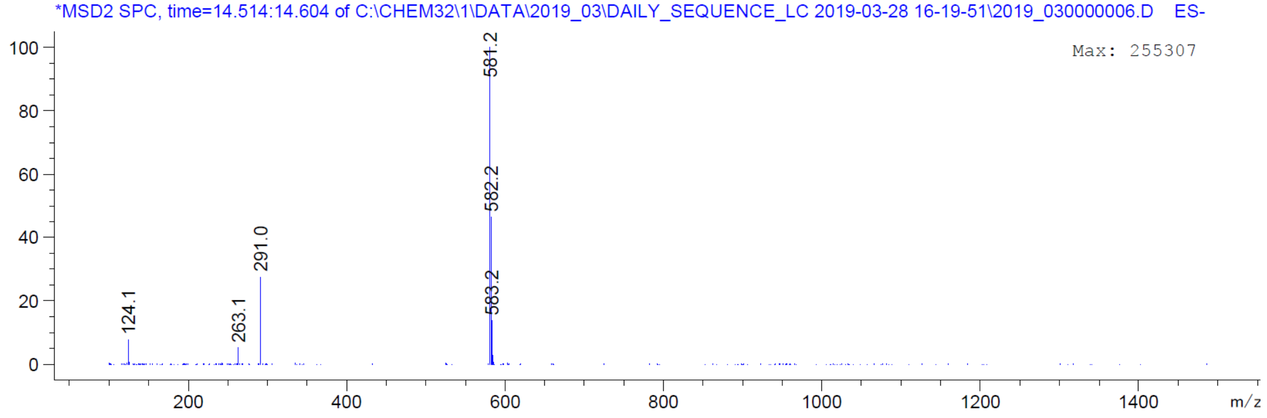
*MSD2 SPC, time=14.514:14.604 of C:\CHEM32\1\DATA\2019_03\DAILY_SEQUENCE_LC 2019-03-28 16-19-51\2019_030000006.D ES-
Max: 255307
100
80
60
40
20
0
124.1
263.1
291.0
581.2
582.2
583.2
200
400
600
800
1000
1200
1400
m/z

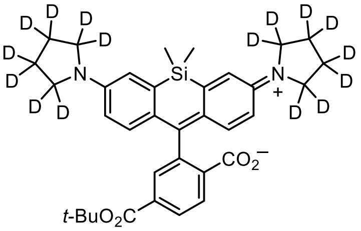
D
D
D
D
D
D
D
D
N
Si
N
+
D
D
D
D
D
D
D
D
$CO_2^-$
$t$-$BuO_2C$

**S16**

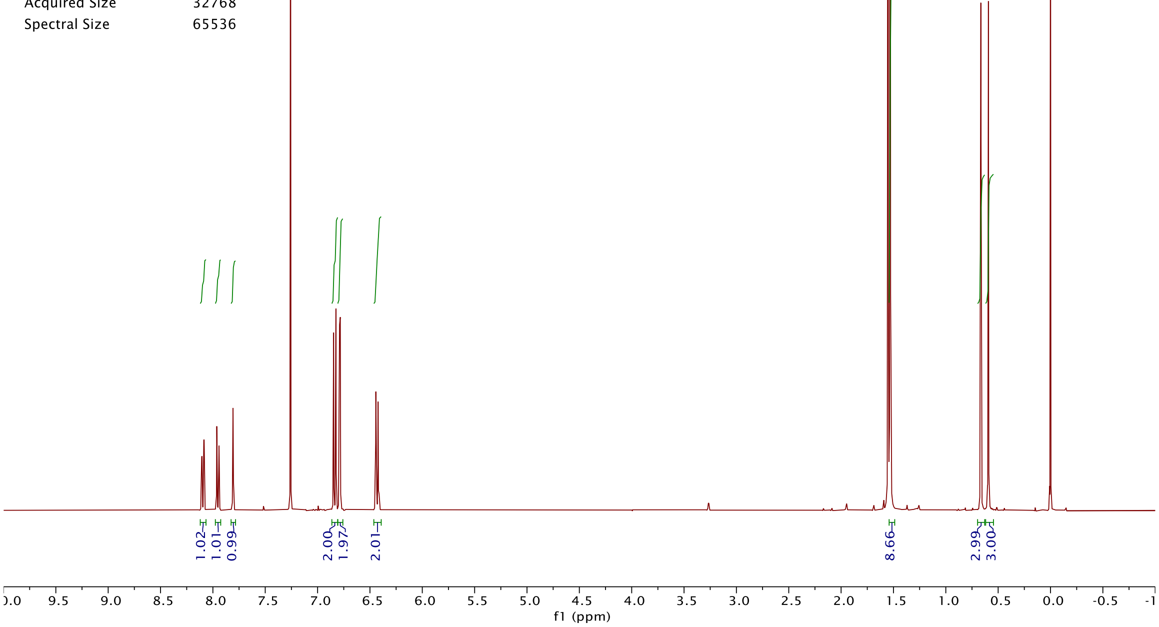
Origin Bruker BioSpin GmbH
Solvent CDCl3
Temperature 300.0
Pulse Sequence zg30
Experiment 1D
Number of Scans 16
Acquisition Date 2019-04-02T10:23:00
Spectrometer Frequency 400.13
Spectral Width 8012.8
Lowest Frequency -1546.1
Nucleus 1H
Acquired Size 32768
Spectral Size 65536
1.02
1.01
0.99
2.00
1.97
2.01
8.66
2.99
3.00
f1 (ppm)

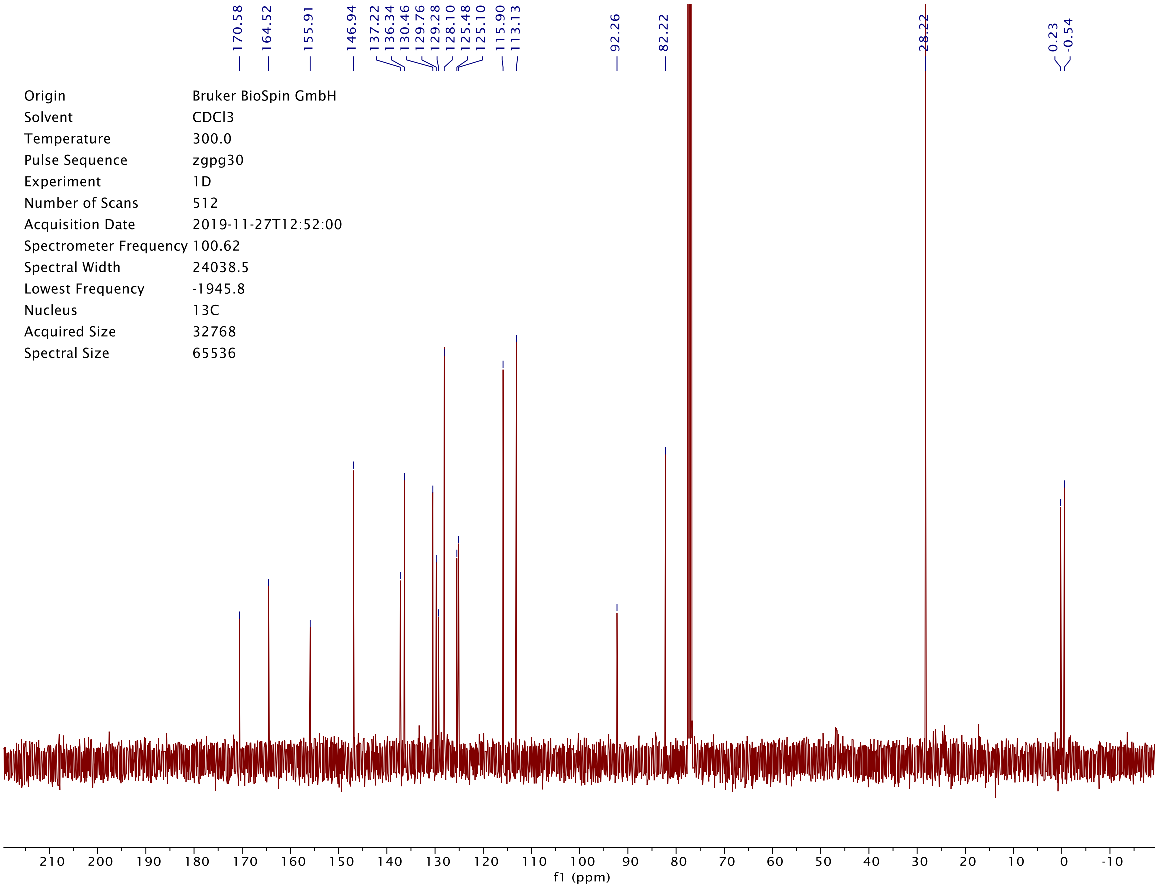
170.58
164.52
155.91
146.94
137.22
136.34
130.46
129.76
129.28
128.10
125.48
125.10
115.90
113.13
92.26
82.22
28.22
0.23
-0.54
Origin Bruker BioSpin GmbH
Solvent CDCl3
Temperature 300.0
Pulse Sequence zgpg30
Experiment 1D
Number of Scans 512
Acquisition Date 2019-11-27T12:52:00
Spectrometer Frequency 100.62
Spectral Width 24038.5
Lowest Frequency -1945.8
Nucleus 13C
Acquired Size 32768
Spectral Size 65536
f1 (ppm)

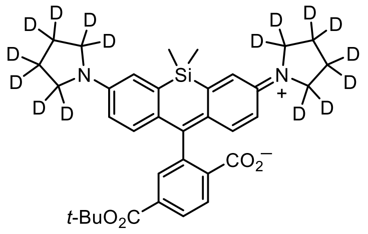

**S16**

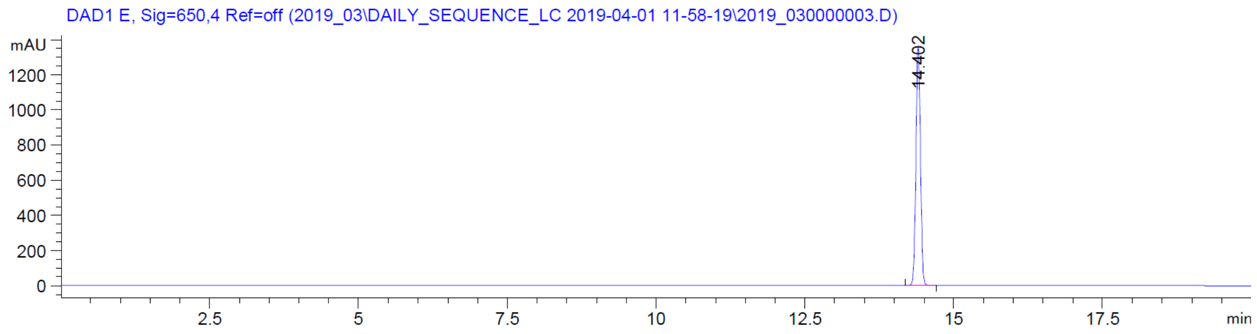
DAD1 E, Sig=650,4 Ref=off (2019_03\DAILY_SEQUENCE_LC 2019-04-01 11-58-19\2019_030000003.D)
mAU
14.402
min

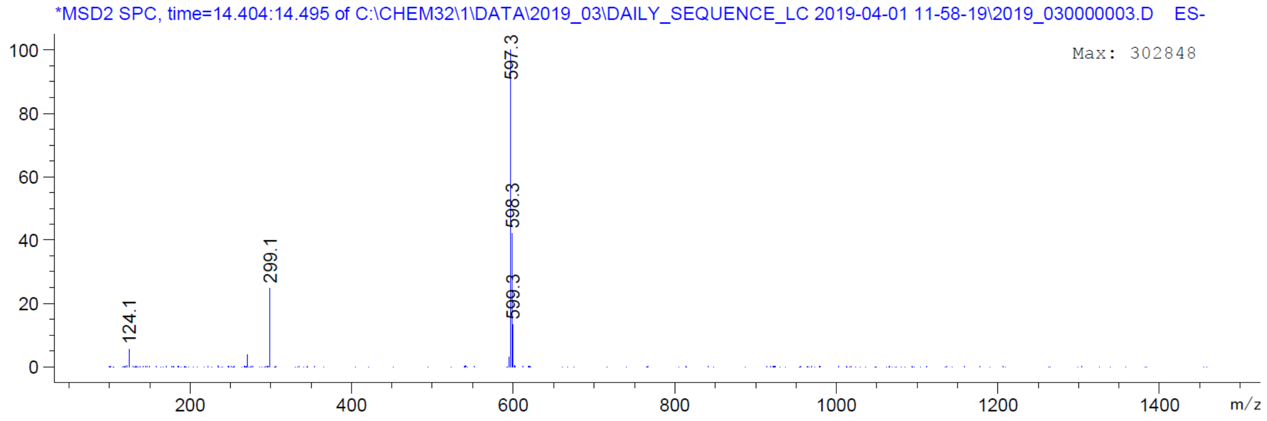
*MSD2 SPC, time=14.404:14.495 of C:\CHEM32\1\DATA\2019_03\DAILY_SEQUENCE_LC 2019-04-01 11-58-19\2019_030000003.D ES-
Max: 302848
597.3
598.3
599.3
299.1
124.1
m/z

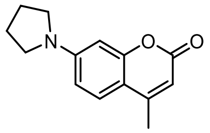

**24**

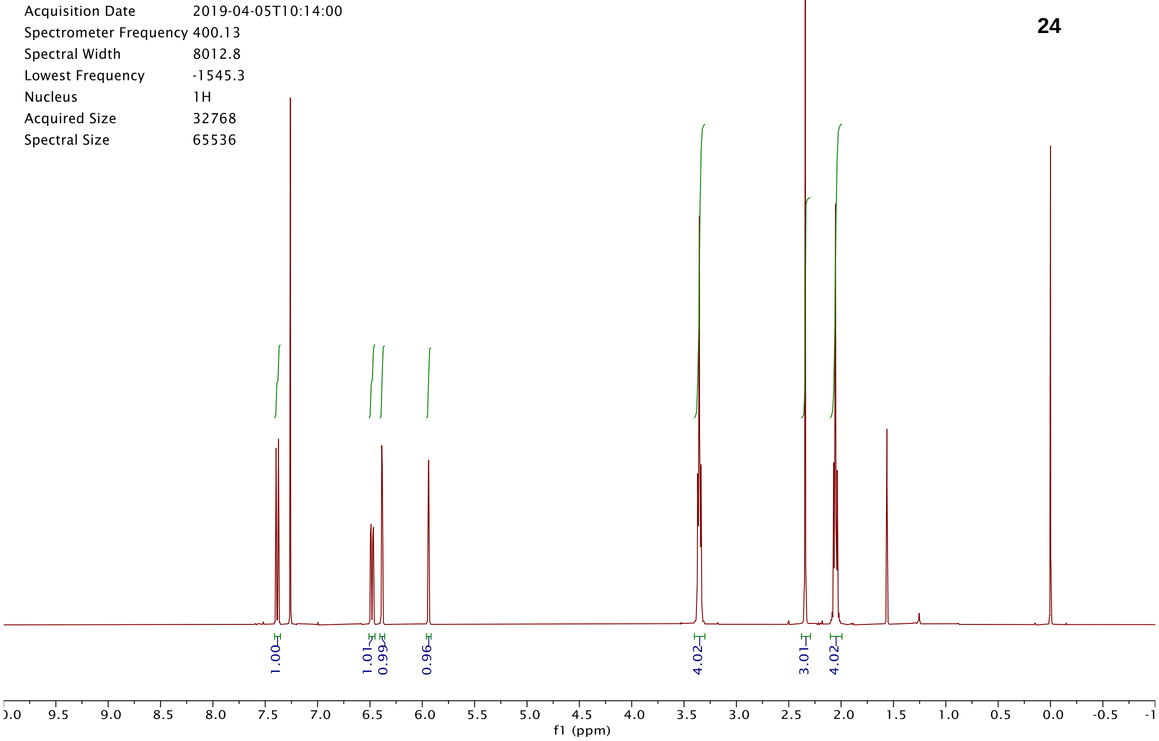

Origin Bruker BioSpin GmbH
Solvent CDCl3
Temperature 300.0
Pulse Sequence zg30
Experiment 1D
Number of Scans 16
Acquisition Date 2019-04-05T10:14:00
Spectrometer Frequency 400.13
Spectral Width 8012.8
Lowest Frequency -1545.3
Nucleus 1H
Acquired Size 32768
Spectral Size 65536
1.00
1.01
0.99
0.96
4.02
3.01
4.02
f1 (ppm)


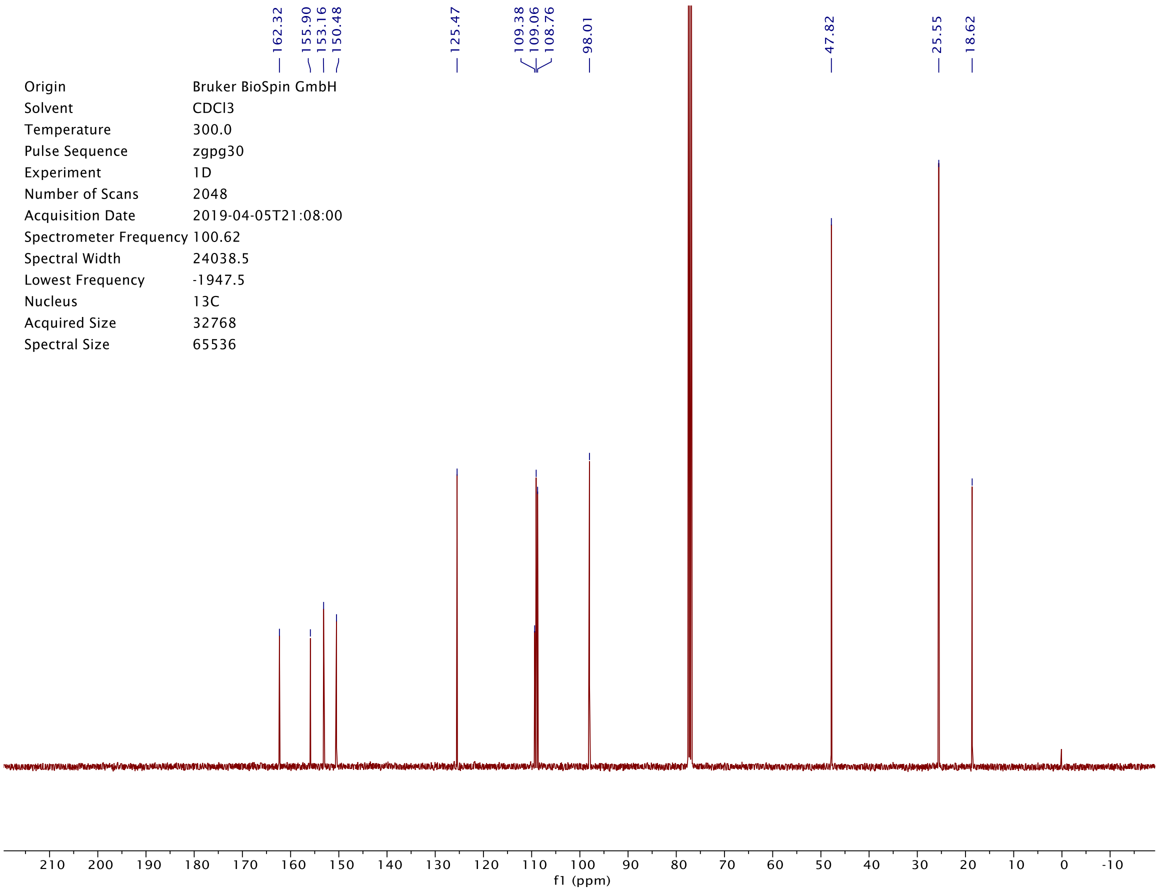

162.32
155.90
153.16
150.48
125.47
109.38
109.06
108.76
98.01
47.82
25.55
18.62
Origin Bruker BioSpin GmbH
Solvent CDCl3
Temperature 300.0
Pulse Sequence zgpg30
Experiment 1D
Number of Scans 2048
Acquisition Date 2019-04-05T21:08:00
Spectrometer Frequency 100.62
Spectral Width 24038.5
Lowest Frequency -1947.5
Nucleus 13C
Acquired Size 32768
Spectral Size 65536
f1 (ppm)

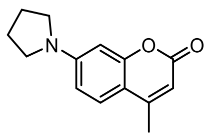

**24**

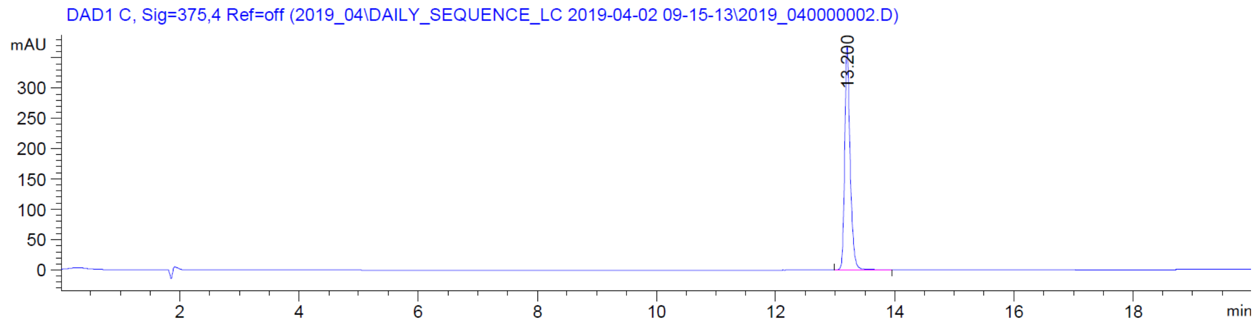

DAD1 C, Sig=375,4 Ref=off (2019_04\DAILY_SEQUENCE_LC 2019-04-02 09-15-13\2019_040000002.D)
mAU
13.200
min


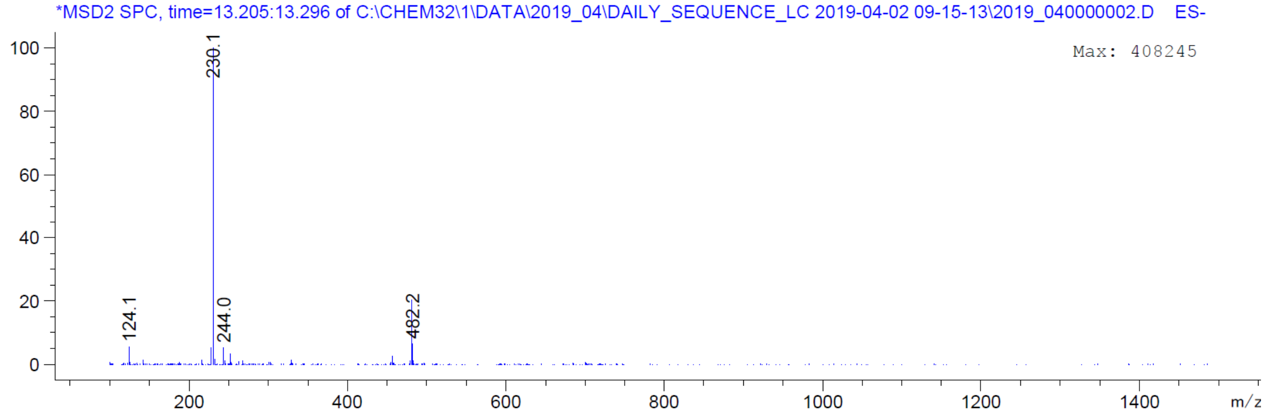

*MSD2 SPC, time=13.205:13.296 of C:\CHEM32\1\DATA\2019_04\DAILY_SEQUENCE_LC 2019-04-02 09-15-13\2019_040000002.D ES-
Max: 408245
230.1
124.1
244.0
482.2
m/z

| | |
|---|---|
| Origin | Bruker BioSpin GmbH |
| Solvent | CDCl3 |
| Temperature | 300.0 |
| Pulse Sequence | zg30 |
| Experiment | 1D |
| Number of Scans | 16 |
| Acquisition Date | 2019-04-05T10:19:00 |
| Spectrometer Frequency | 400.13 |
| Spectral Width | 8012.8 |
| Lowest Frequency | -1545.2 |
| Nucleus | 1H |
| Acquired Size | 32768 |
| Spectral Size | 65536 |

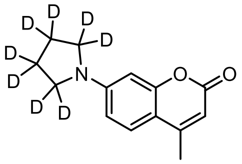


**$24_D$**

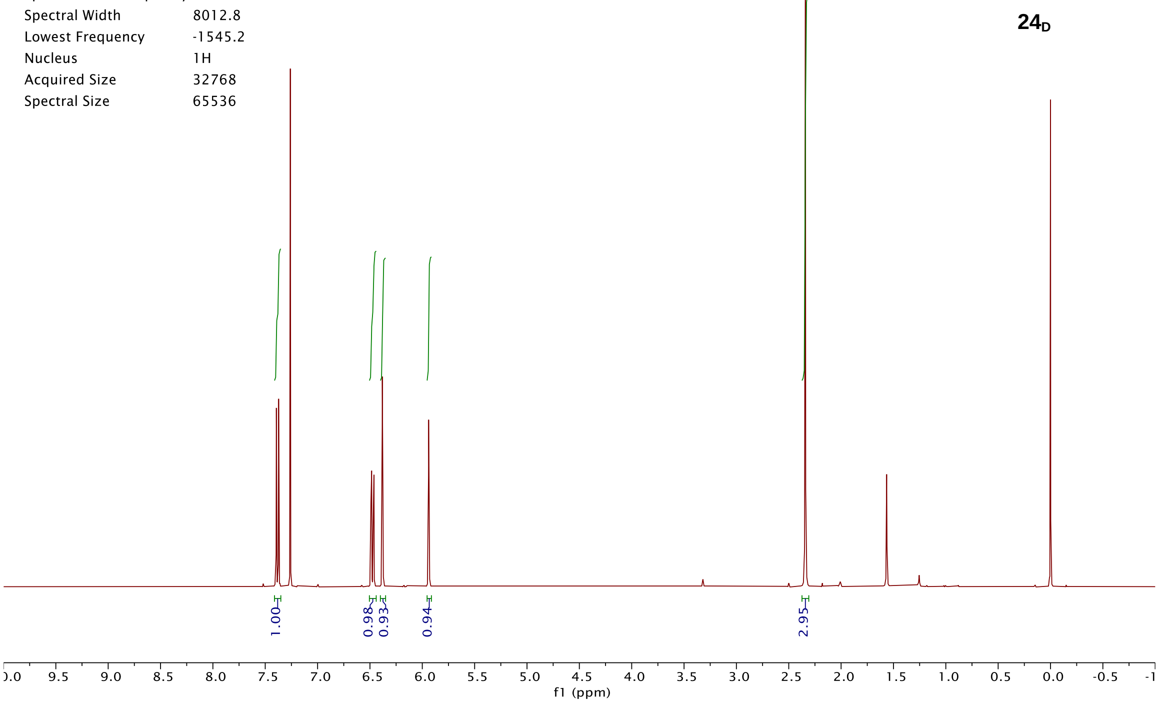


| | |
|---|---|
| Origin | Bruker BioSpin GmbH |
| Solvent | CDCl3 |
| Temperature | 300.0 |
| Pulse Sequence | zgpg30 |
| Experiment | 1D |
| Number of Scans | 256 |
| Acquisition Date | 2019-05-10T14:56:00 |
| Spectrometer Frequency | 100.62 |
| Spectral Width | 24038.5 |
| Lowest Frequency | -1947.2 |
| Nucleus | 13C |
| Acquired Size | 32768 |
| Spectral Size | 65536 |

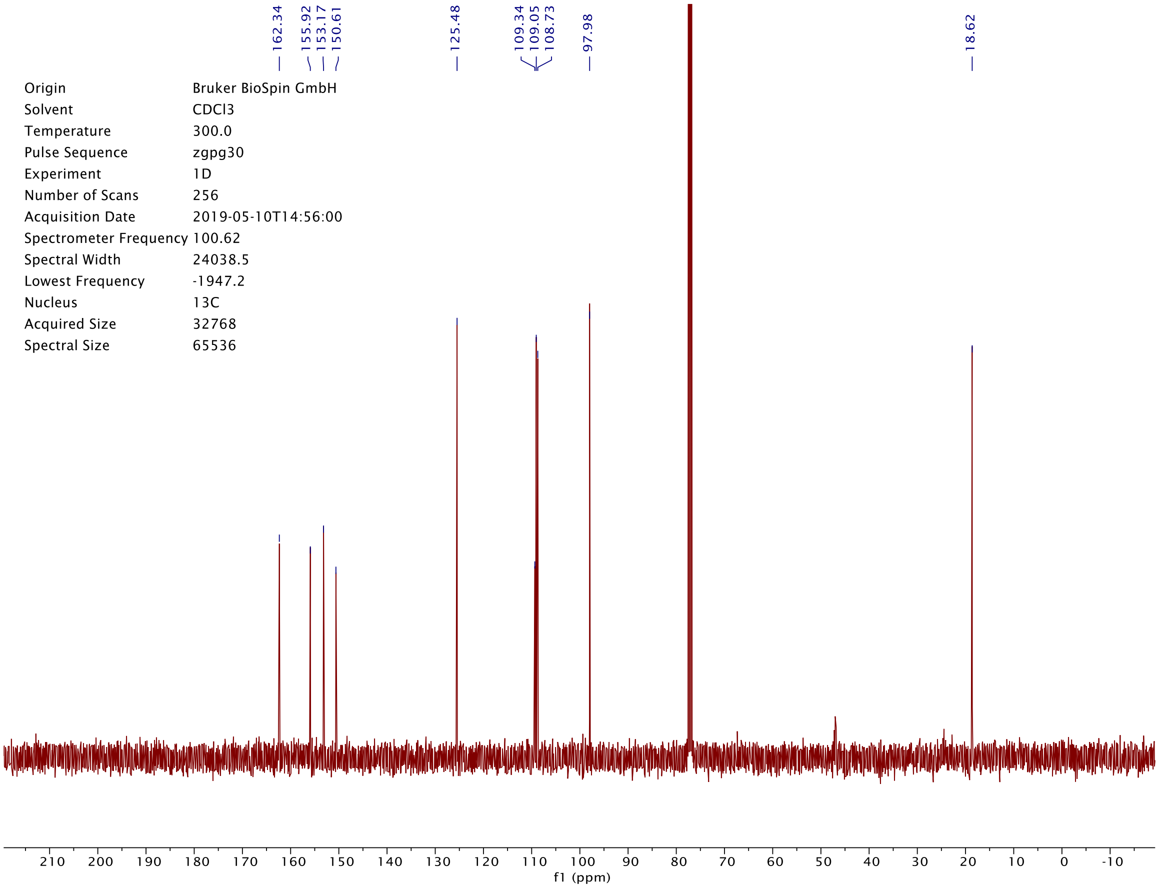

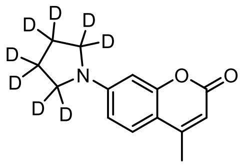

**24**$_D$

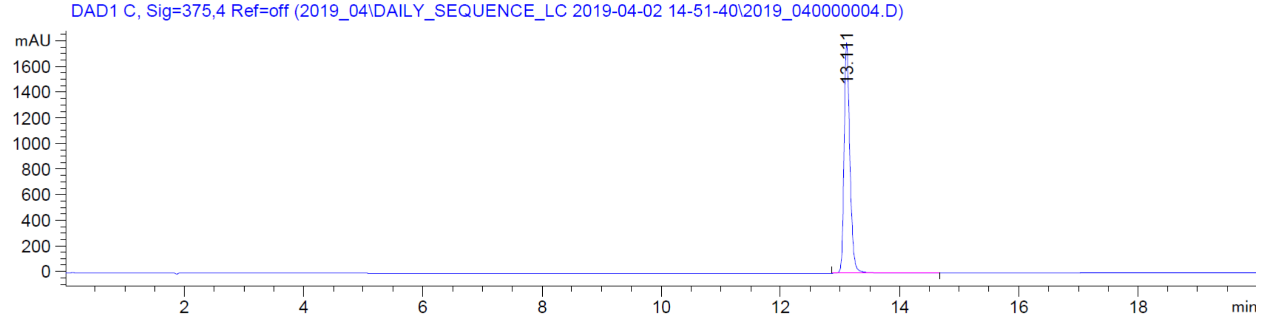

DAD1 C, Sig=375,4 Ref=off (2019_04\DAILY_SEQUENCE_LC 2019-04-02 14-51-40\2019_040000004.D)
mAU
1600
1400
1200
1000
800
600
400
200
0
13.111
2
4
6
8
10
12
14
16
18
min


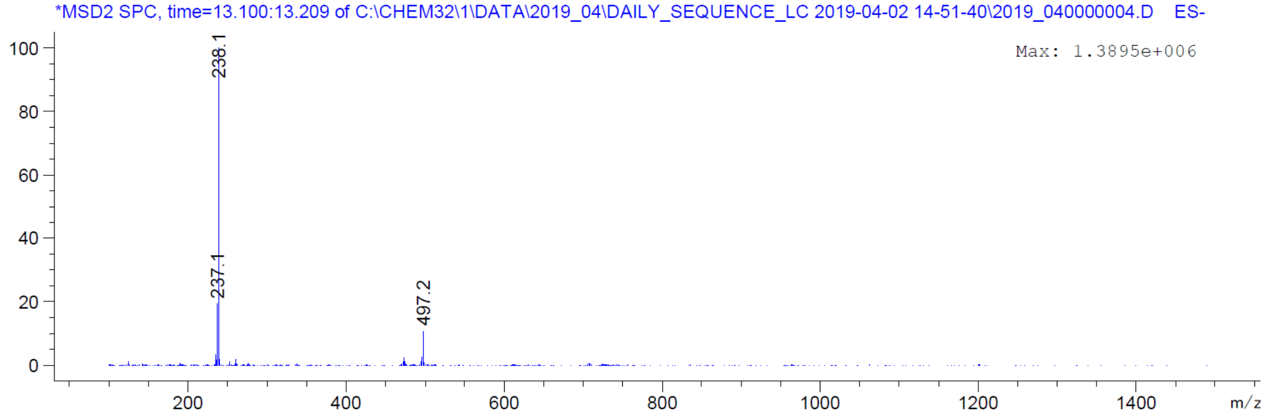

*MSD2 SPC, time=13.100:13.209 of C:\CHEM32\1\DATA\2019_04\DAILY_SEQUENCE_LC 2019-04-02 14-51-40\2019_040000004.D ES-
Max: 1.3895e+006
100
80
60
40
20
0
238.1
237.1
497.2
200
400
600
800
1000
1200
1400
m/z

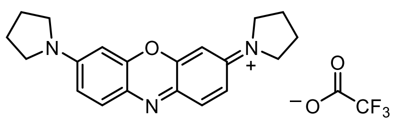

**25**

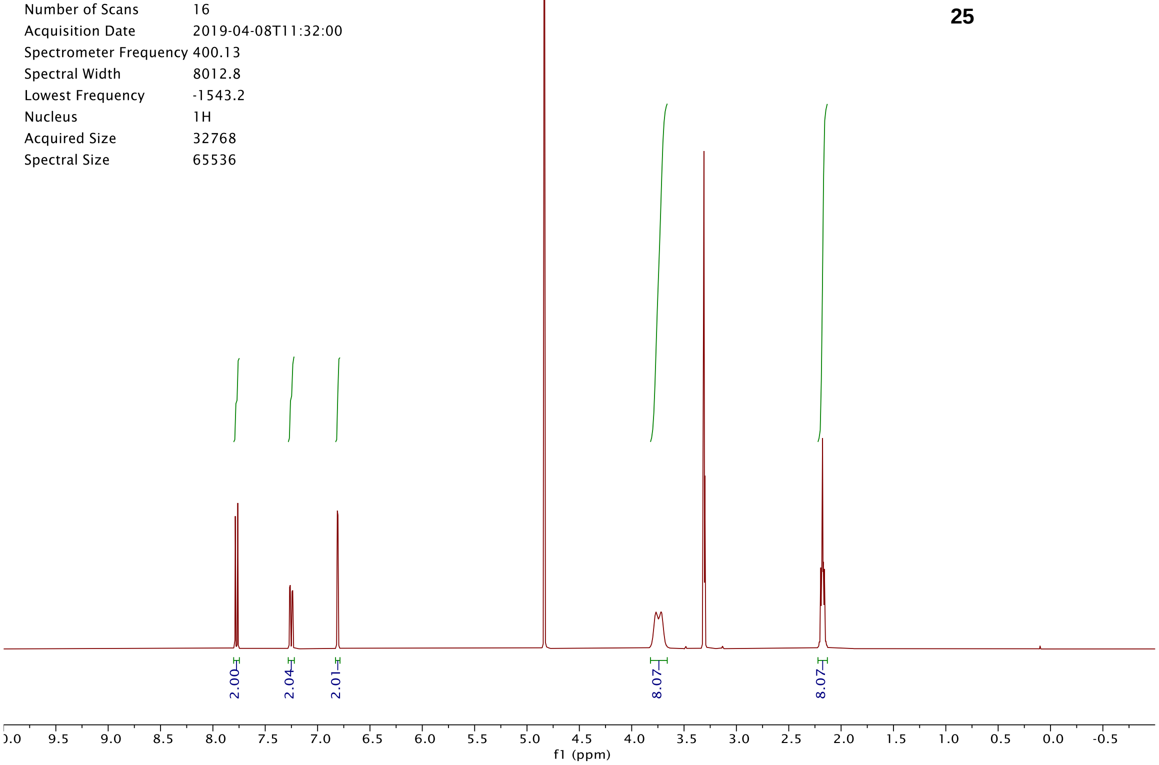

Origin Bruker BioSpin GmbH
Solvent MeOD
Temperature 300.0
Pulse Sequence zg30
Experiment 1D
Number of Scans 16
Acquisition Date 2019-04-08T11:32:00
Spectrometer Frequency 400.13
Spectral Width 8012.8
Lowest Frequency -1543.2
Nucleus 1H
Acquired Size 32768
Spectral Size 65536
2.00
2.04
2.01
8.07
8.07
f1 (ppm)


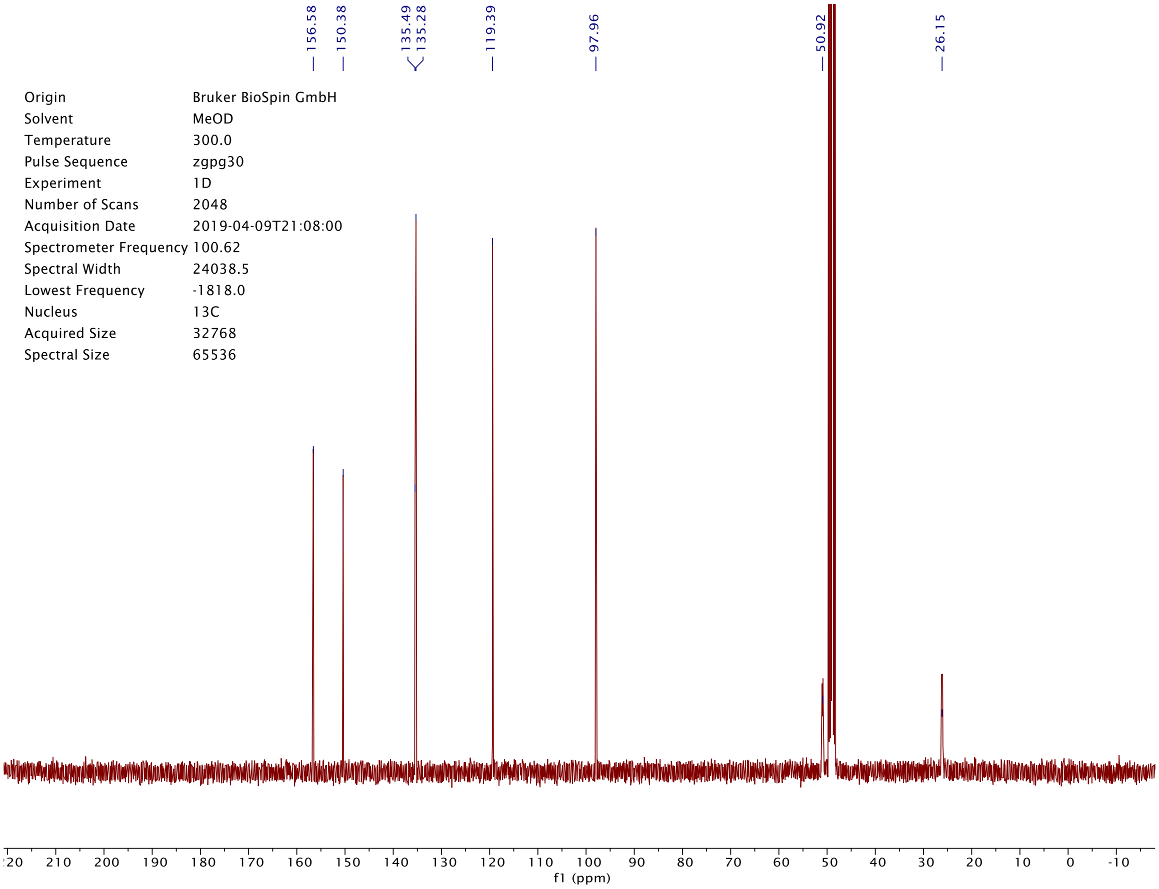

156.58
150.38
135.49
135.28
119.39
97.96
50.92
26.15
Origin Bruker BioSpin GmbH
Solvent MeOD
Temperature 300.0
Pulse Sequence zgpg30
Experiment 1D
Number of Scans 2048
Acquisition Date 2019-04-09T21:08:00
Spectrometer Frequency 100.62
Spectral Width 24038.5
Lowest Frequency -1818.0
Nucleus 13C
Acquired Size 32768
Spectral Size 65536
f1 (ppm)

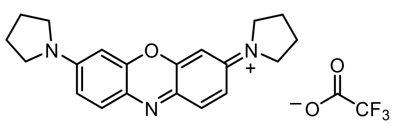

**25**

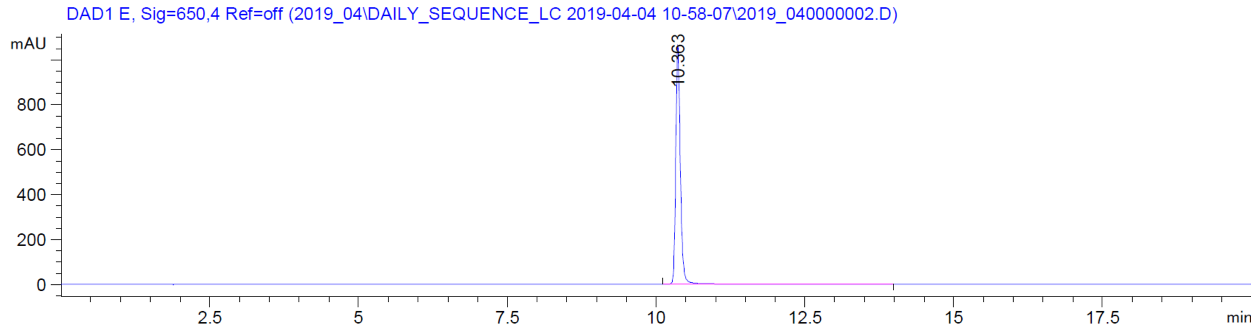

DAD1 E, Sig=650,4 Ref=off (2019_04\DAILY_SEQUENCE_LC 2019-04-04 10-58-07\2019_040000002.D)
mAU
800
600
400
200
0
10.363
2.5
5
7.5
10
12.5
15
17.5
min


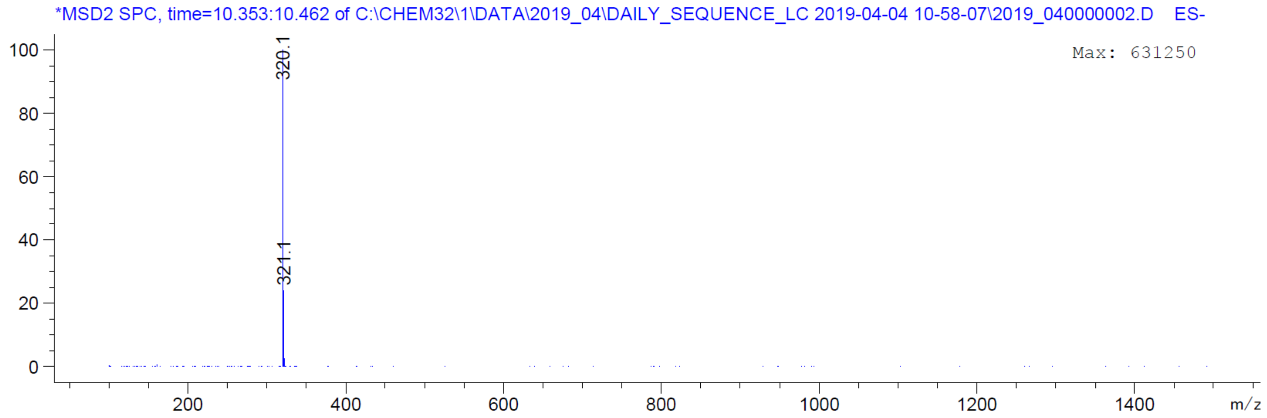

*MSD2 SPC, time=10.353:10.462 of C:\CHEM32\1\DATA\2019_04\DAILY_SEQUENCE_LC 2019-04-04 10-58-07\2019_040000002.D ES-
Max: 631250
100
80
60
40
20
0
320.1
321.1
200
400
600
800
1000
1200
1400
m/z

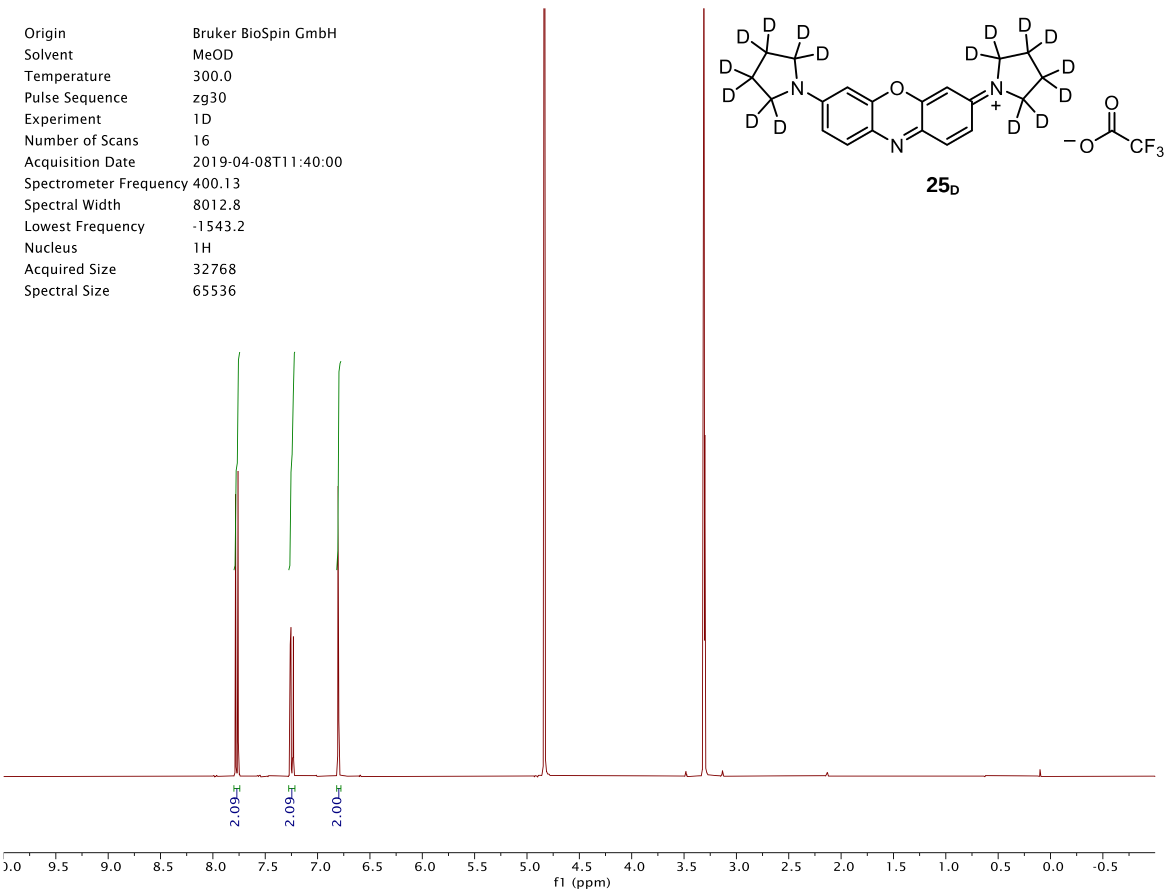
Origin Bruker BioSpin GmbH
Solvent MeOD
Temperature 300.0
Pulse Sequence zg30
Experiment 1D
Number of Scans 16
Acquisition Date 2019-04-08T11:40:00
Spectrometer Frequency 400.13
Spectral Width 8012.8
Lowest Frequency -1543.2
Nucleus 1H
Acquired Size 32768
Spectral Size 65536
25D
2.09
2.09
2.00
f1 (ppm)

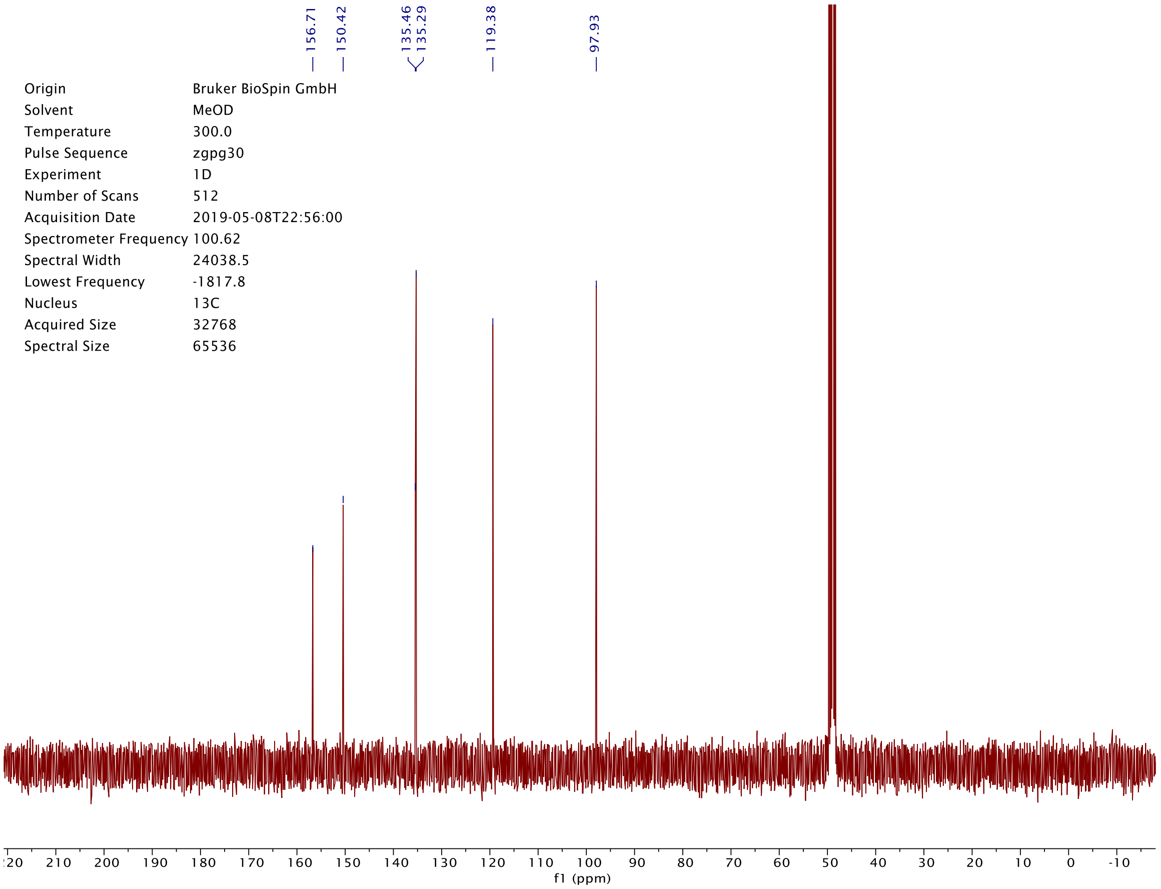
156.71
150.42
135.46
135.29
119.38
97.93
Origin Bruker BioSpin GmbH
Solvent MeOD
Temperature 300.0
Pulse Sequence zgpg30
Experiment 1D
Number of Scans 512
Acquisition Date 2019-05-08T22:56:00
Spectrometer Frequency 100.62
Spectral Width 24038.5
Lowest Frequency -1817.8
Nucleus 13C
Acquired Size 32768
Spectral Size 65536
f1 (ppm)

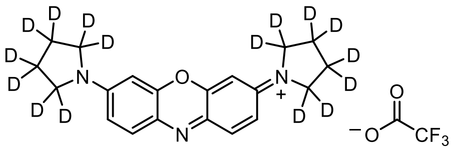

**25**$_D$

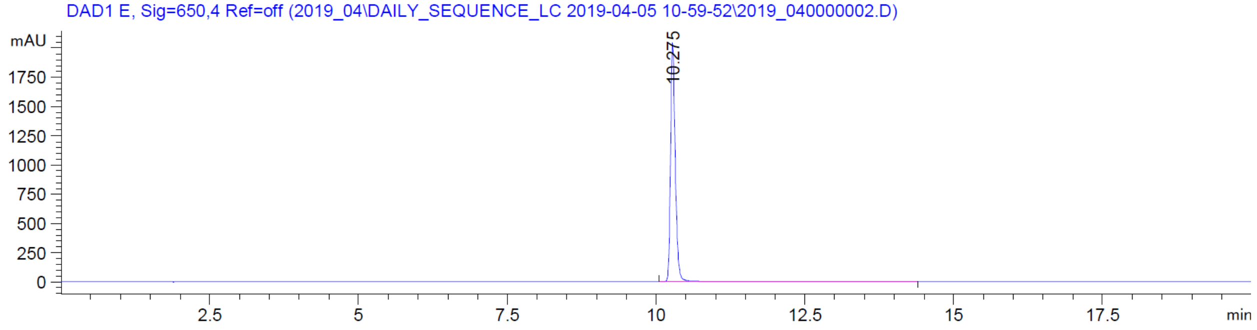
DAD1 E, Sig=650,4 Ref=off (2019_04\DAILY_SEQUENCE_LC 2019-04-05 10-59-52\2019_040000002.D)
mAU
10.275
min

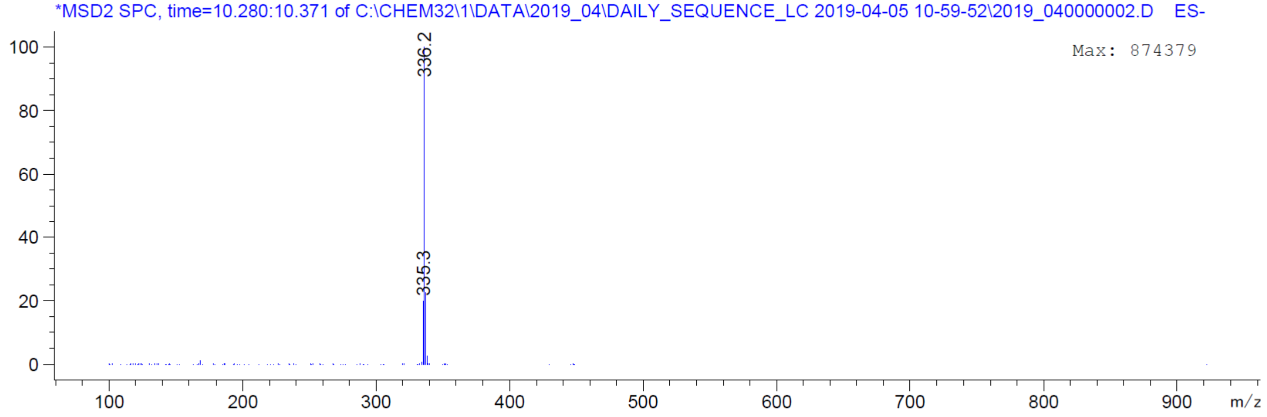
*MSD2 SPC, time=10.280:10.371 of C:\CHEM32\1\DATA\2019_04\DAILY_SEQUENCE_LC 2019-04-05 10-59-52\2019_040000002.D ES-
Max: 874379
336.2
335.3
m/z

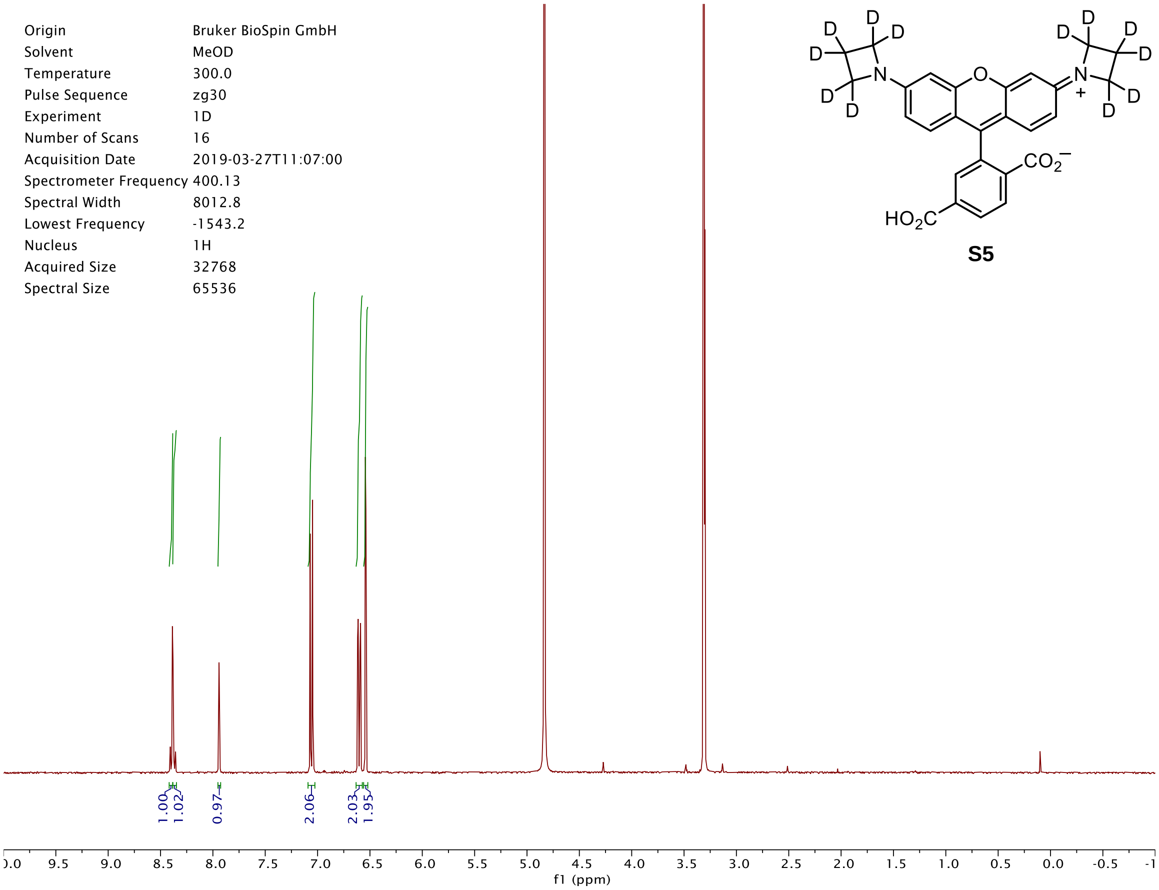
Origin Bruker BioSpin GmbH
Solvent MeOD
Temperature 300.0
Pulse Sequence zg30
Experiment 1D
Number of Scans 16
Acquisition Date 2019-03-27T11:07:00
Spectrometer Frequency 400.13
Spectral Width 8012.8
Lowest Frequency -1543.2
Nucleus 1H
Acquired Size 32768
Spectral Size 65536
$CO_2^-$
$HO_2C$
1.00
1.02
0.97
2.06
2.03
1.95
f1 (ppm)

**S5**

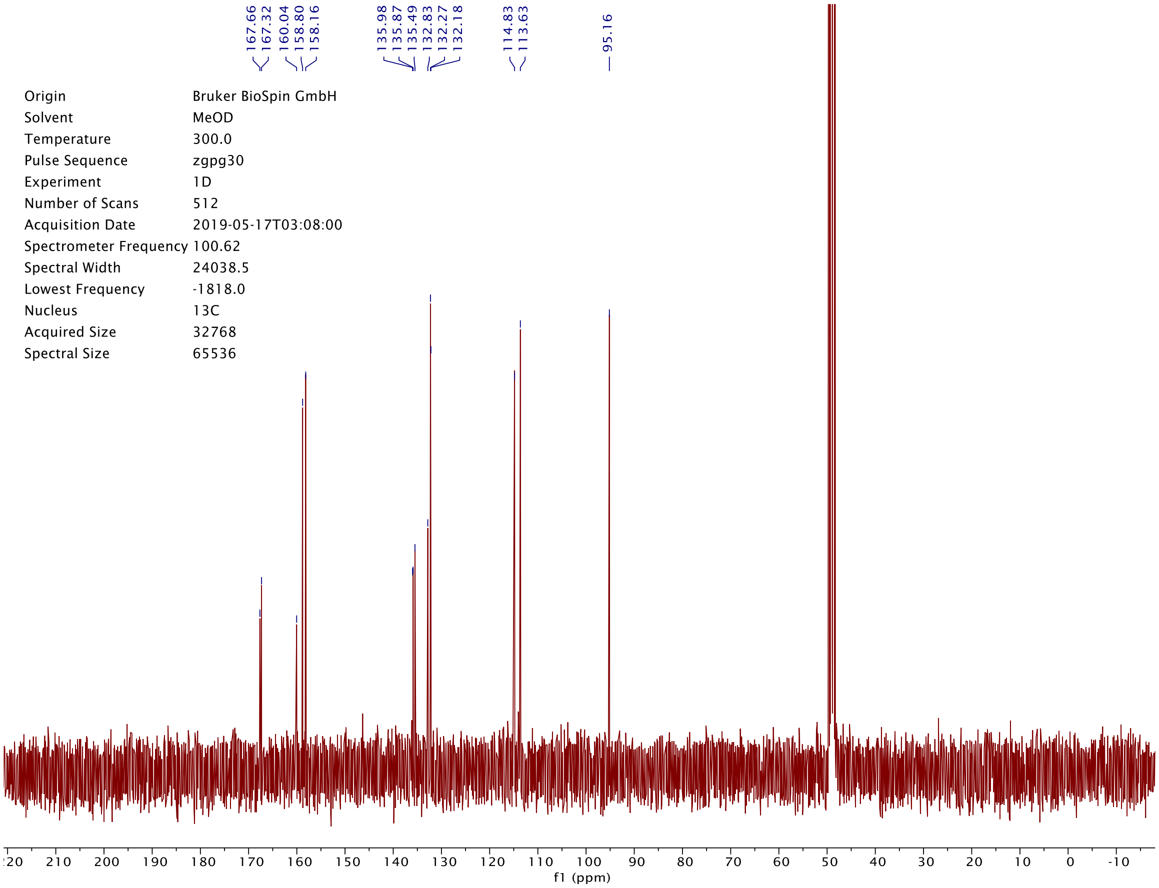
167.66
167.32
160.04
158.80
158.16
135.98
135.87
135.49
132.83
132.27
132.18
114.83
113.63
95.16
Origin Bruker BioSpin GmbH
Solvent MeOD
Temperature 300.0
Pulse Sequence zgpg30
Experiment 1D
Number of Scans 512
Acquisition Date 2019-05-17T03:08:00
Spectrometer Frequency 100.62
Spectral Width 24038.5
Lowest Frequency -1818.0
Nucleus 13C
Acquired Size 32768
Spectral Size 65536
f1 (ppm)

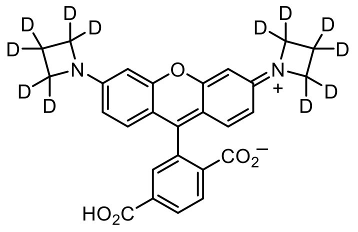
HO2C
CO2
N
O
D

**S5**

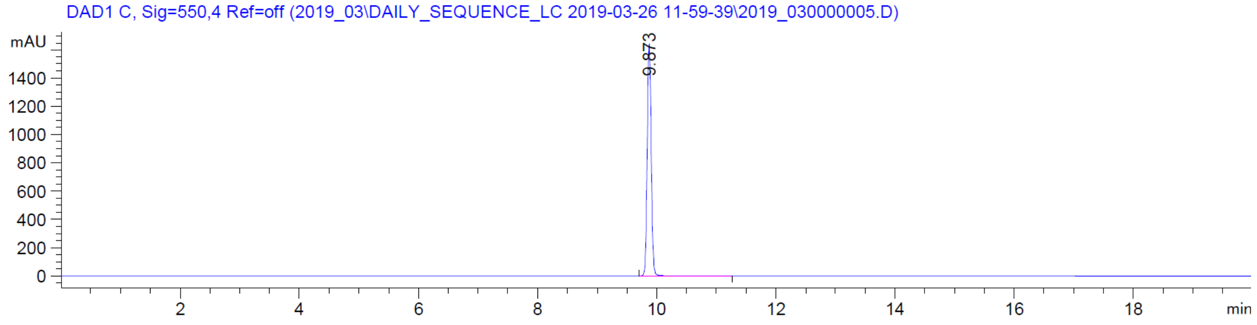
DAD1 C, Sig=550,4 Ref=off (2019_03\DAILY_SEQUENCE_LC 2019-03-26 11-59-39\2019_030000005.D)
mAU
9.873
min

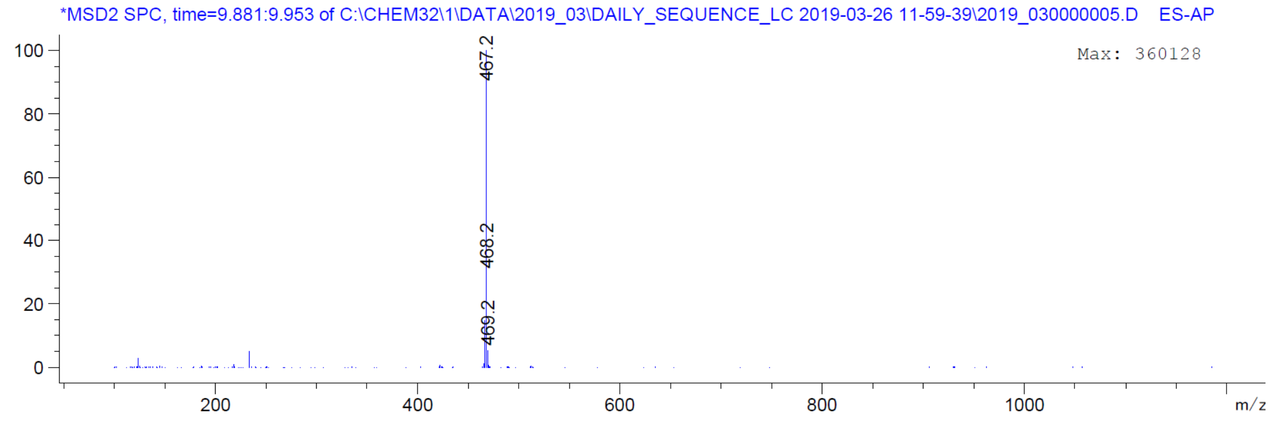
*MSD2 SPC, time=9.881:9.953 of C:\CHEM32\1\DATA\2019_03\DAILY_SEQUENCE_LC 2019-03-26 11-59-39\2019_030000005.D ES-AP
Max: 360128
467.2
468.2
469.2
m/z

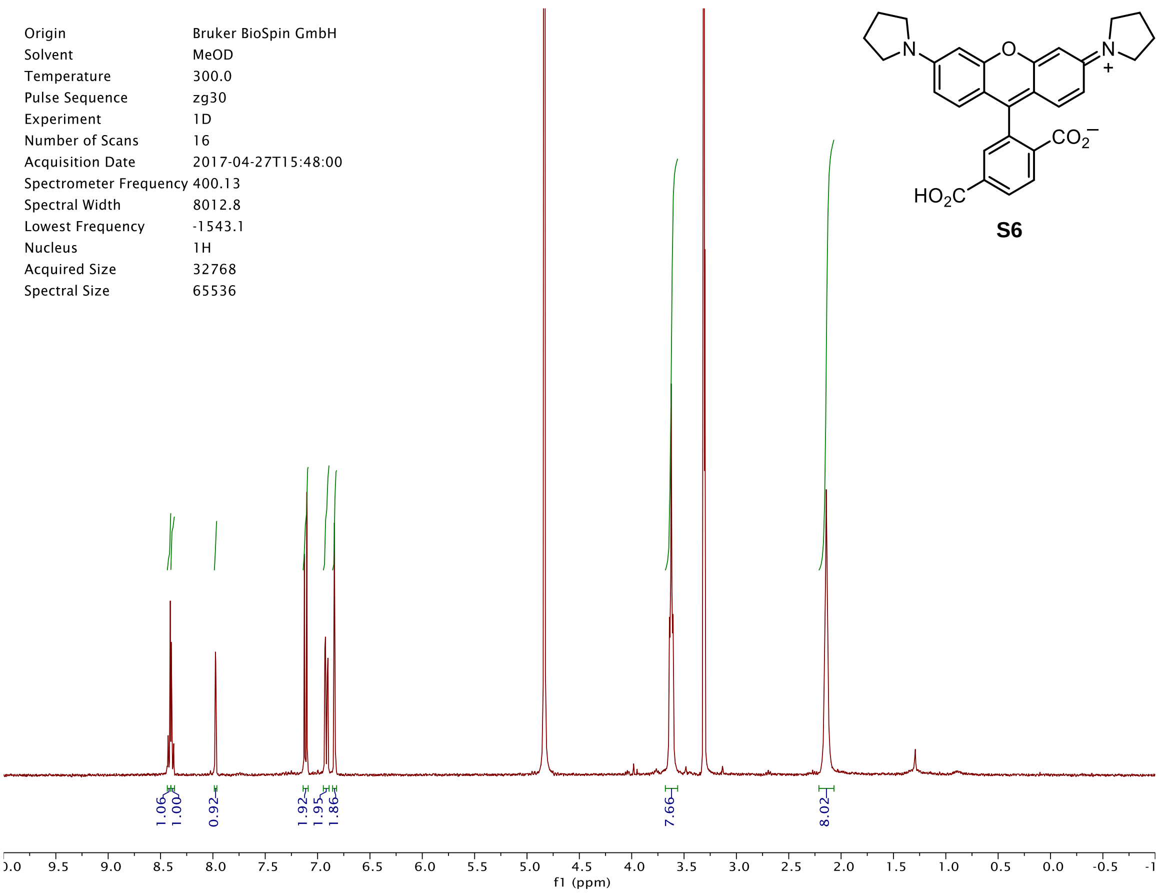

Origin Bruker BioSpin GmbH
Solvent MeOD
Temperature 300.0
Pulse Sequence zg30
Experiment 1D
Number of Scans 16
Acquisition Date 2017-04-27T15:48:00
Spectrometer Frequency 400.13
Spectral Width 8012.8
Lowest Frequency -1543.1
Nucleus 1H
Acquired Size 32768
Spectral Size 65536
N
O
N
+
$CO_2^-$
$HO_2C$
S6
1.06
1.00
0.92
1.92
1.95
1.86
7.66
8.02
f1 (ppm)


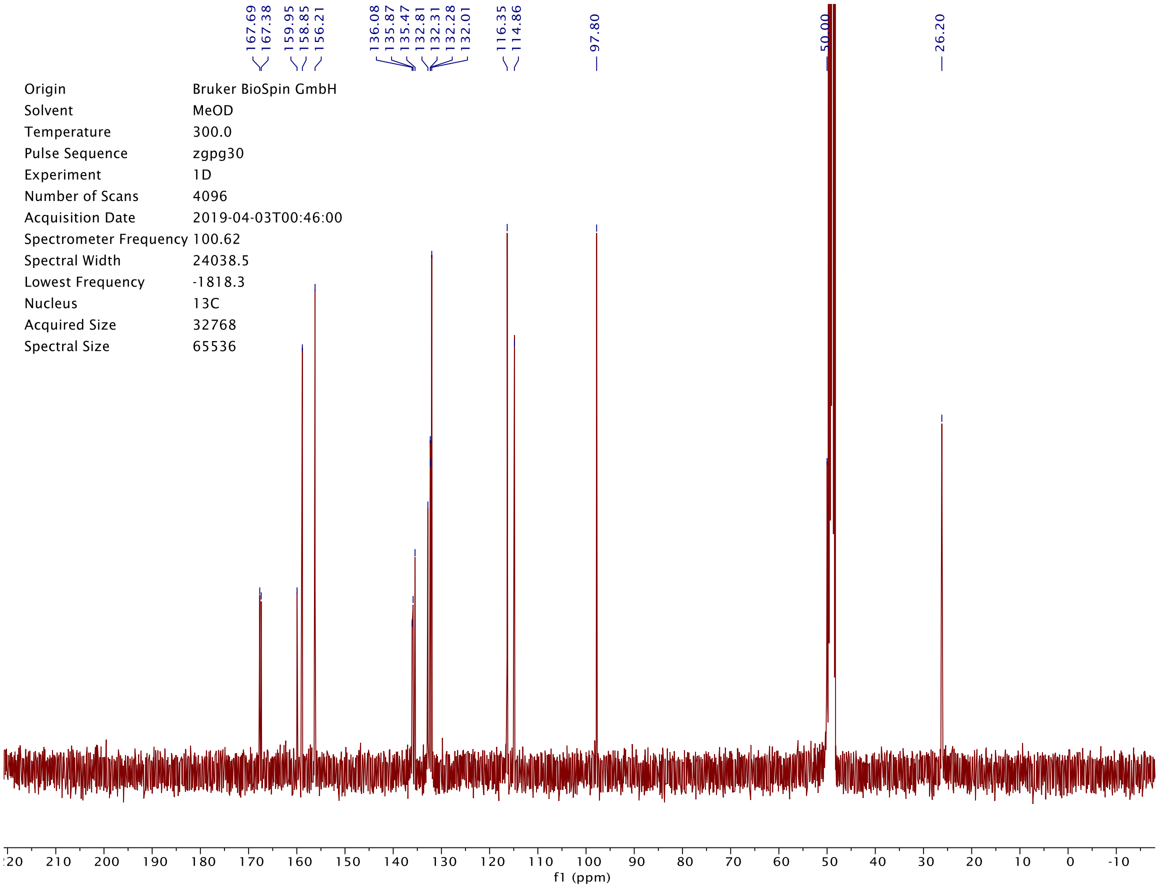

167.69
167.38
159.95
158.85
156.21
136.08
135.87
135.47
132.81
132.31
132.28
132.01
116.35
114.86
97.80
50.00
26.20
Origin Bruker BioSpin GmbH
Solvent MeOD
Temperature 300.0
Pulse Sequence zgpg30
Experiment 1D
Number of Scans 4096
Acquisition Date 2019-04-03T00:46:00
Spectrometer Frequency 100.62
Spectral Width 24038.5
Lowest Frequency -1818.3
Nucleus 13C
Acquired Size 32768
Spectral Size 65536
f1 (ppm)

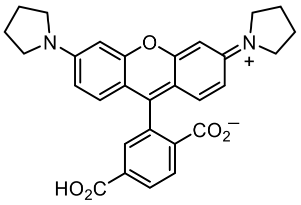
N
O
N
+
$CO_2^-$
$HO_2C$

**S6**

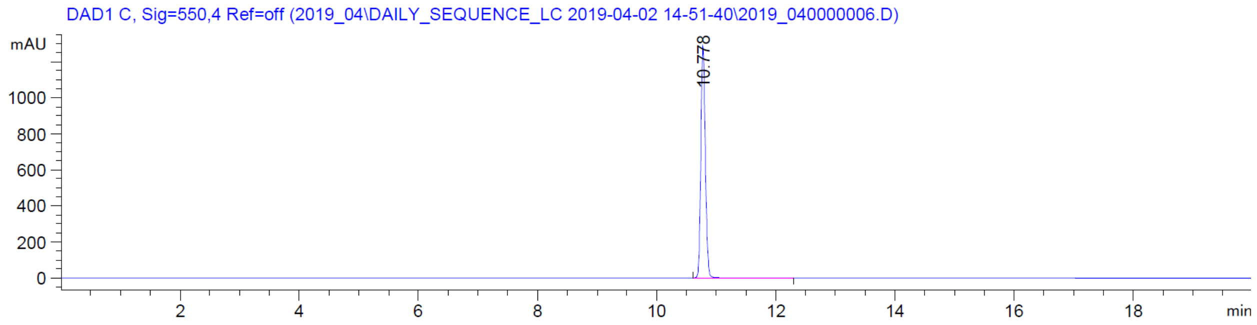
DAD1 C, Sig=550,4 Ref=off (2019_04\DAILY_SEQUENCE_LC 2019-04-02 14-51-40\2019_040000006.D)
mAU
1000
800
600
400
200
0
10.778
2
4
6
8
10
12
14
16
18
min

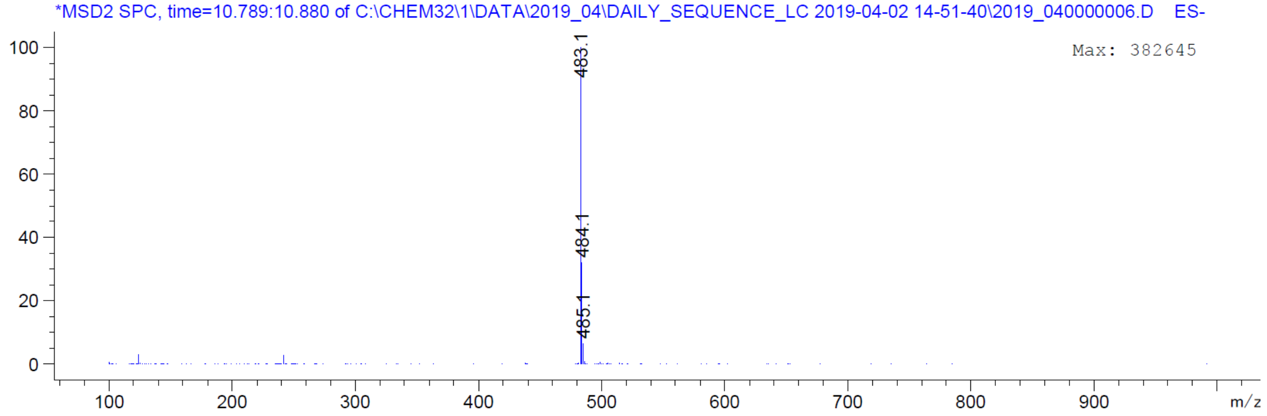
*MSD2 SPC, time=10.789:10.880 of C:\CHEM32\1\DATA\2019_04\DAILY_SEQUENCE_LC 2019-04-02 14-51-40\2019_040000006.D ES-
Max: 382645
100
80
60
40
20
0
483.1
484.1
485.1
100
200
300
400
500
600
700
800
900
m/z

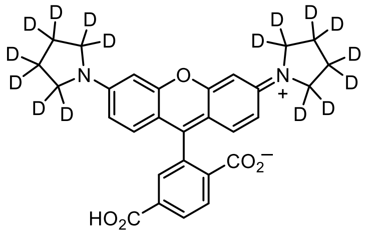
D
D
D
D
D
D
D
D
N
O
N
+
D
D
D
D
D
D
D
D
$CO_2^-$
$HO_2C$

**S7**

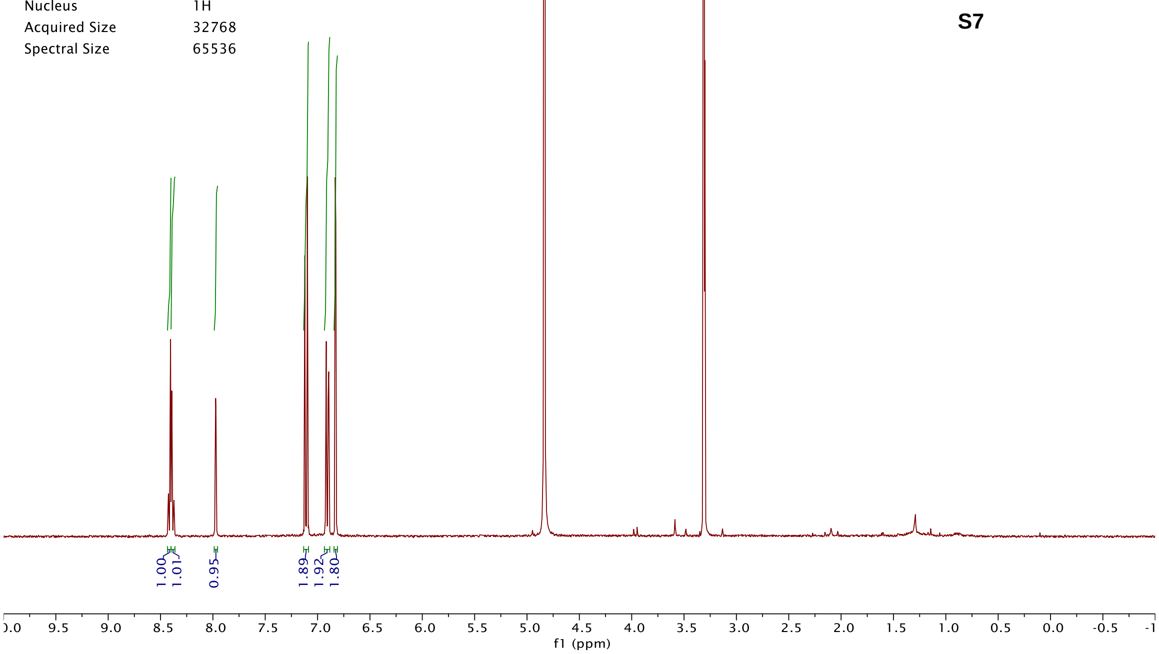
Origin Bruker BioSpin GmbH
Solvent MeOD
Temperature 300.0
Pulse Sequence zg30
Experiment 1D
Number of Scans 16
Acquisition Date 2017-04-27T15:52:00
Spectrometer Frequency 400.13
Spectral Width 8012.8
Lowest Frequency -1543.2
Nucleus 1H
Acquired Size 32768
Spectral Size 65536
1.00
1.01
0.95
1.89
1.92
1.80
f1 (ppm)

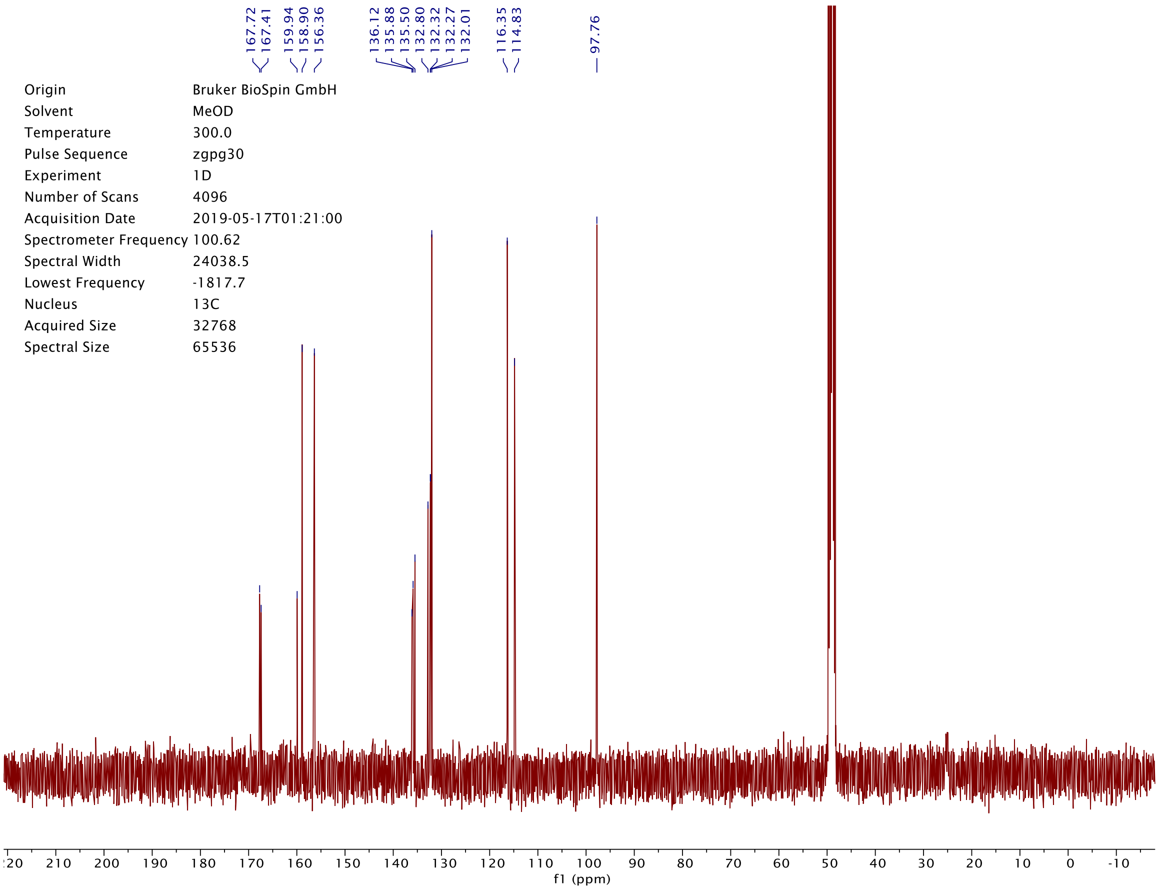
167.72
167.41
159.94
158.90
156.36
136.12
135.88
135.50
132.80
132.32
132.27
132.01
116.35
114.83
97.76
Origin Bruker BioSpin GmbH
Solvent MeOD
Temperature 300.0
Pulse Sequence zgpg30
Experiment 1D
Number of Scans 4096
Acquisition Date 2019-05-17T01:21:00
Spectrometer Frequency 100.62
Spectral Width 24038.5
Lowest Frequency -1817.7
Nucleus 13C
Acquired Size 32768
Spectral Size 65536
f1 (ppm)

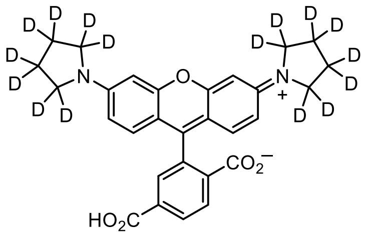

**S7**

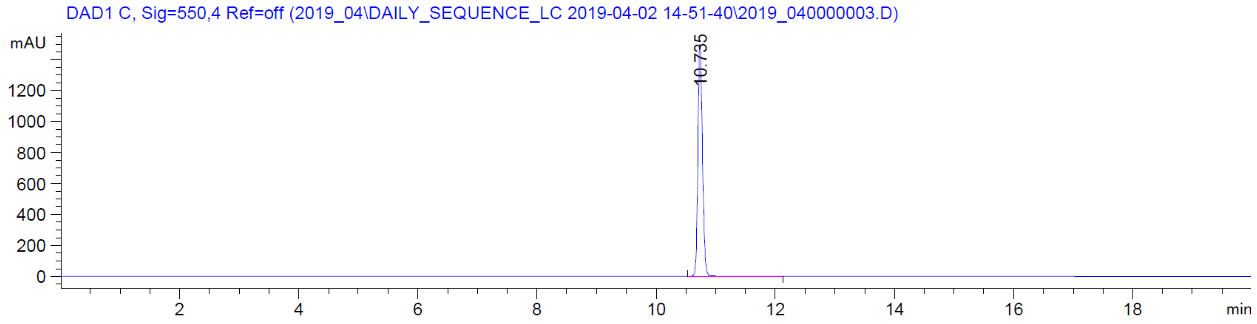
DAD1 C, Sig=550,4 Ref=off (2019_04\DAILY_SEQUENCE_LC 2019-04-02 14-51-40\2019_040000003.D)
mAU
10.735
min

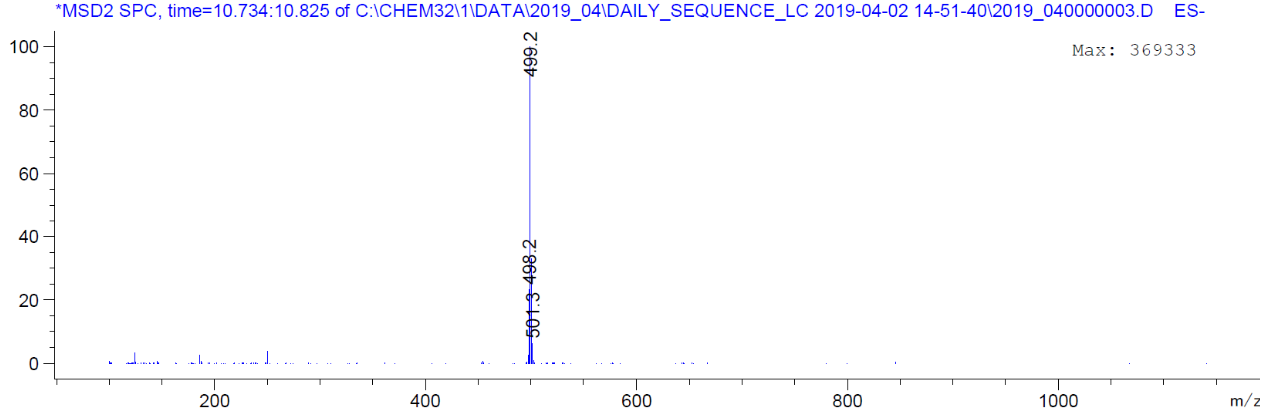
*MSD2 SPC, time=10.734:10.825 of C:\CHEM32\1\DATA\2019_04\DAILY_SEQUENCE_LC 2019-04-02 14-51-40\2019_040000003.D ES-
Max: 369333
499.2
498.2
501.3
m/z

| | |
|---|---|
| Origin | Bruker BioSpin GmbH |
| Solvent | MeOD |
| Temperature | 300.0 |
| Pulse Sequence | zg30 |
| Experiment | 1D |
| Number of Scans | 16 |
| Acquisition Date | 2019-03-25T15:36:00 |
| Spectrometer Frequency | 400.13 |
| Spectral Width | 8012.8 |
| Lowest Frequency | -1543.2 |
| Nucleus | 1H |
| Acquired Size | 32768 |
| Spectral Size | 65536 |

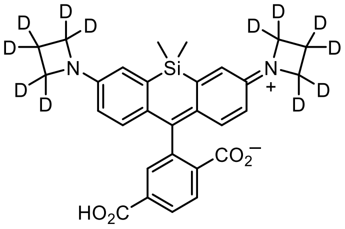


**S17**

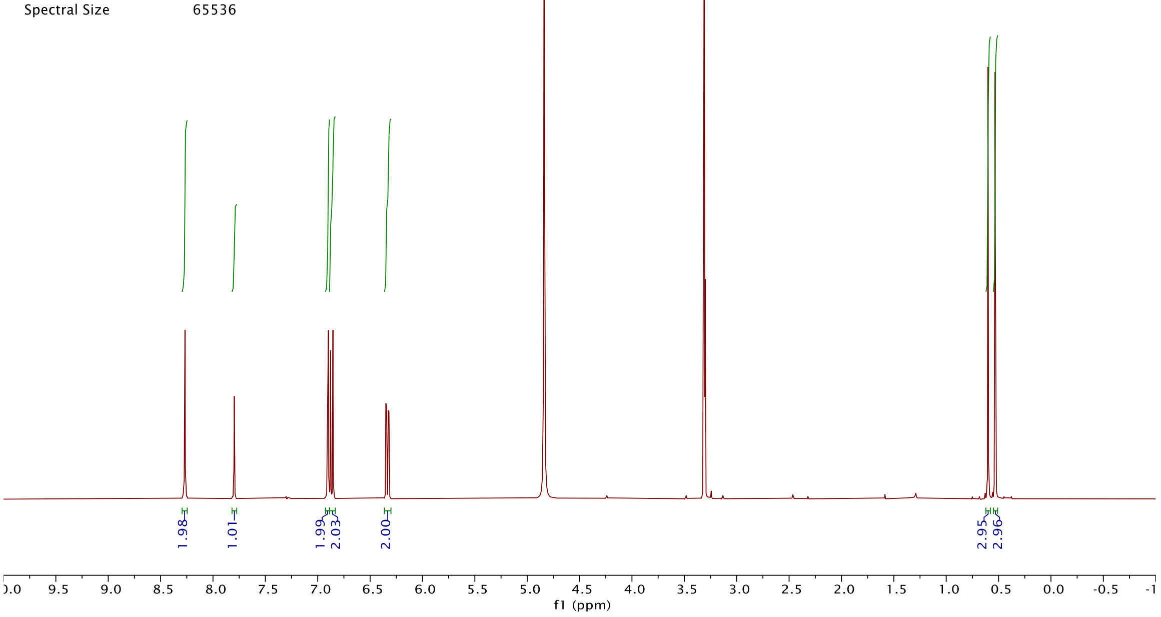


DAD1 E, Sig=650,4 Ref=off (2019_03\DAILY_SEQUENCE_LC 2019-03-25 15-17-17\2019_030000003.D)

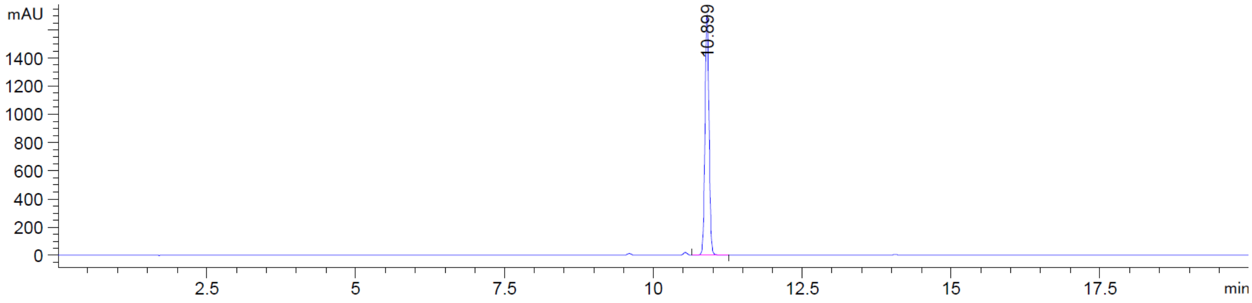


*MSD2 SPC, time=10.898:10.989 of C:\CHEM32\1\DATA\2019_03\DAILY_SEQUENCE_LC 2019-03-25 15-17-17\2019_030000003.D ES-

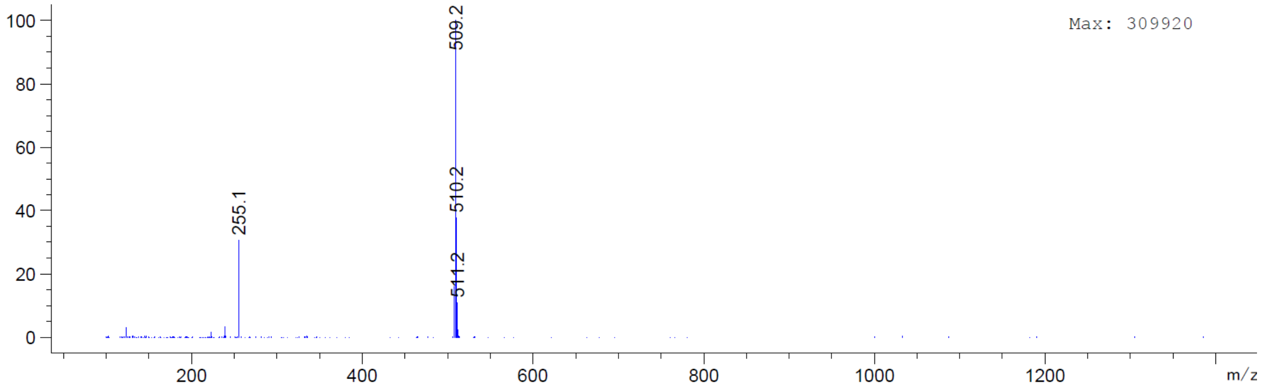

| | |
|---|---|
| Origin | Bruker BioSpin GmbH |
| Solvent | MeOD |
| Temperature | 295.6 |
| Pulse Sequence | zg30 |
| Experiment | 1D |
| Number of Scans | 16 |
| Acquisition Date | 2018-12-03T10:38:00 |
| Spectrometer Frequency | 400.13 |
| Spectral Width | 8012.8 |
| Lowest Frequency | -1543.3 |
| Nucleus | 1H |
| Acquired Size | 32768 |
| Spectral Size | 65536 |

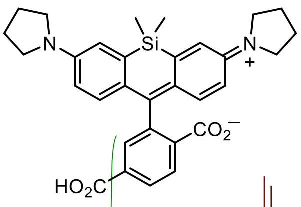


**S18**

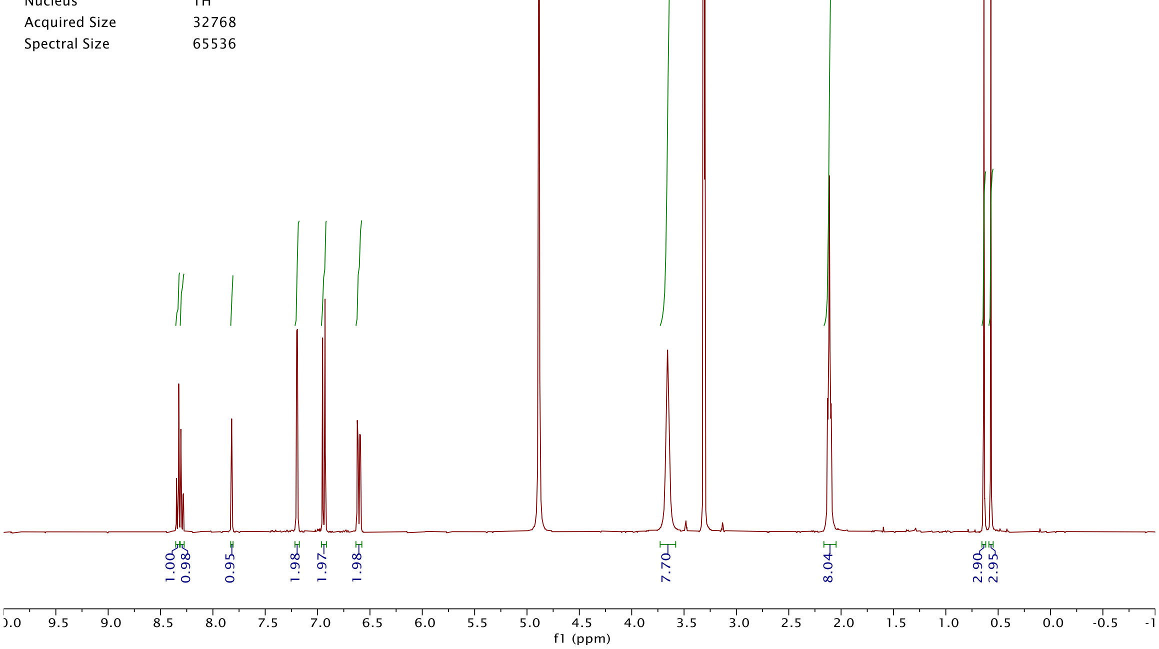


| | |
|---|---|
| Origin | Bruker BioSpin GmbH |
| Solvent | MeOD |
| Temperature | 300.0 |
| Pulse Sequence | zgpg30 |
| Experiment | 1D |
| Number of Scans | 2048 |
| Acquisition Date | 2019-03-29T21:07:00 |
| Spectrometer Frequency | 100.62 |
| Spectral Width | 24038.5 |
| Lowest Frequency | -1818.3 |
| Nucleus | 13C |
| Acquired Size | 32768 |
| Spectral Size | 65536 |

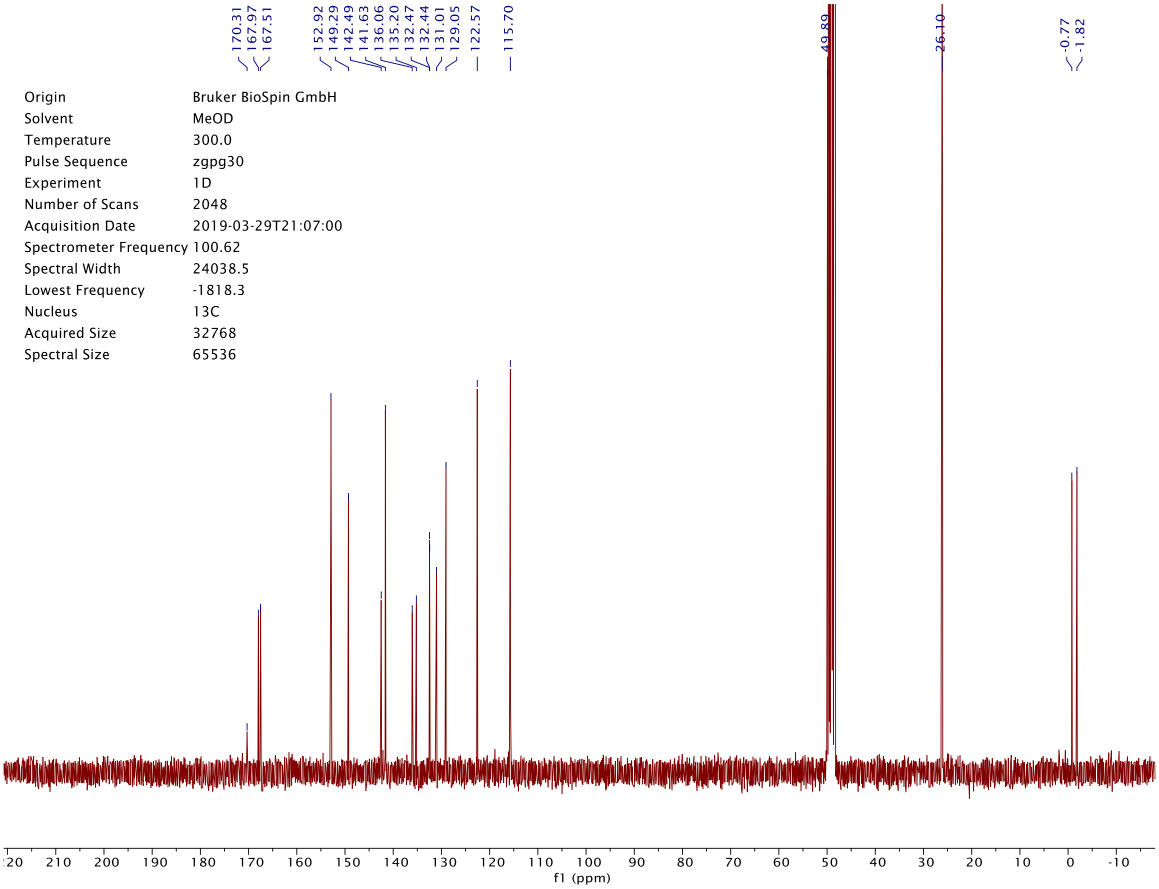

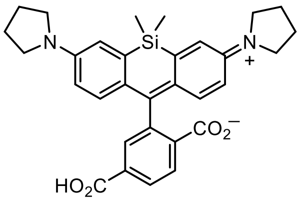
N
Si
N
+
$CO_2^-$
$HO_2C$

**S18**

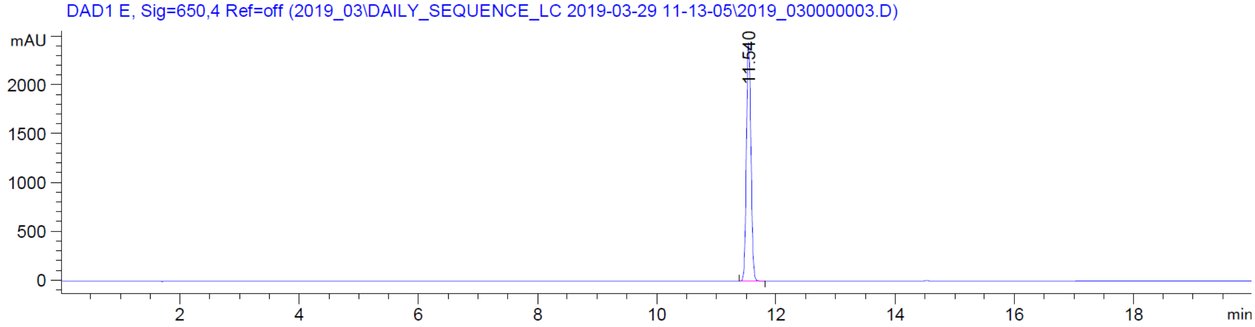
DAD1 E, Sig=650,4 Ref=off (2019_03\DAILY_SEQUENCE_LC 2019-03-29 11-13-05\2019_030000003.D)
mAU
2000
1500
1000
500
0
11.540
2
4
6
8
10
12
14
16
18
min

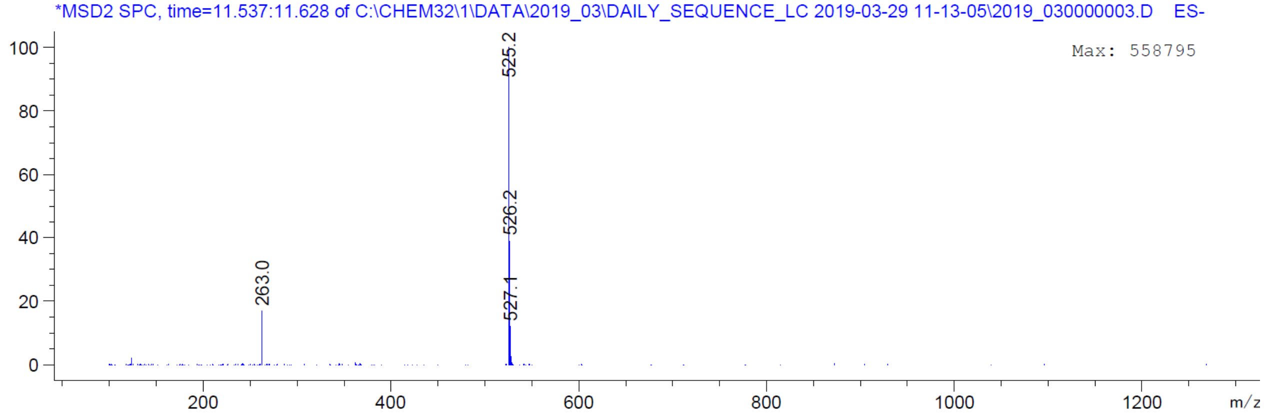
*MSD2 SPC, time=11.537:11.628 of C:\CHEM32\1\DATA\2019_03\DAILY_SEQUENCE_LC 2019-03-29 11-13-05\2019_030000003.D ES-
Max: 558795
100
80
60
40
20
0
525.2
526.2
527.1
263.0
200
400
600
800
1000
1200
m/z

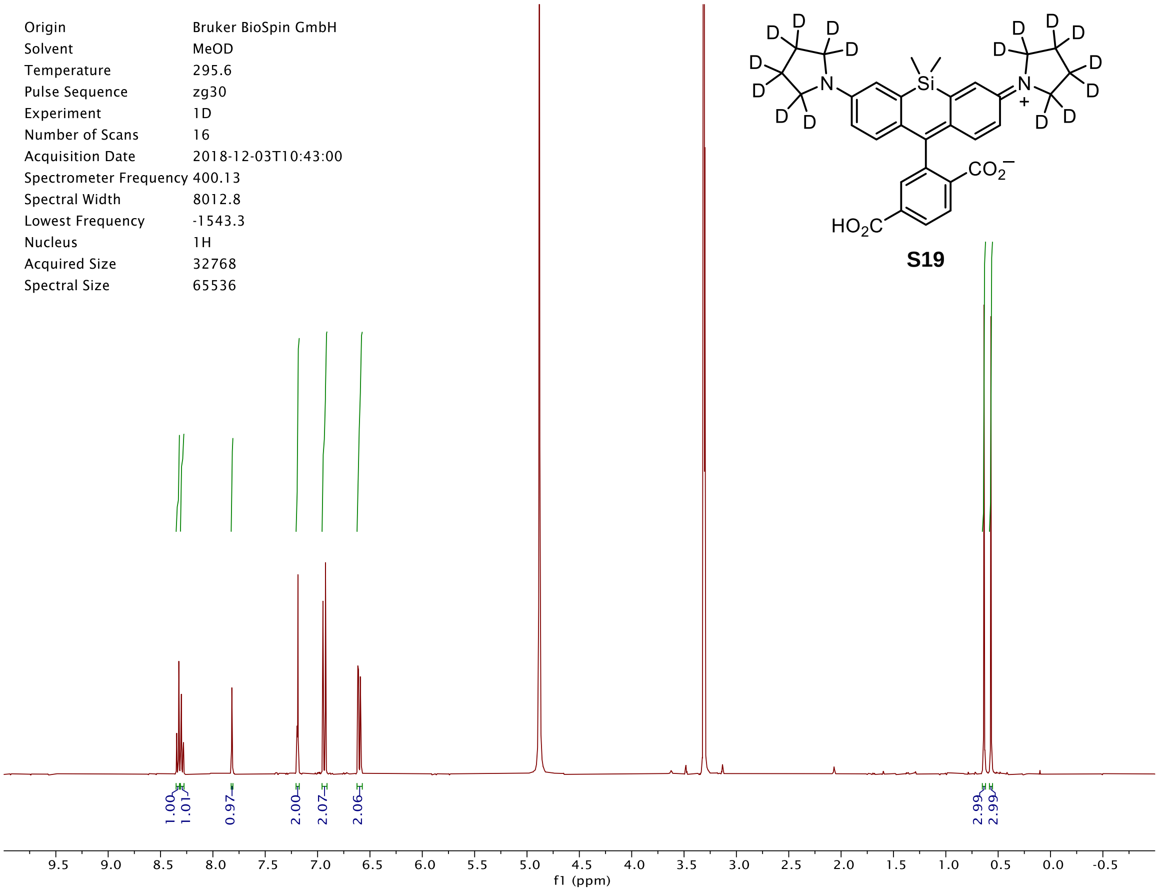
Origin Bruker BioSpin GmbH
Solvent MeOD
Temperature 295.6
Pulse Sequence zg30
Experiment 1D
Number of Scans 16
Acquisition Date 2018-12-03T10:43:00
Spectrometer Frequency 400.13
Spectral Width 8012.8
Lowest Frequency -1543.3
Nucleus 1H
Acquired Size 32768
Spectral Size 65536
1.00
1.01
0.97
2.00
2.07
2.06
2.99
2.99
9.5 9.0 8.5 8.0 7.5 7.0 6.5 6.0 5.5 5.0 4.5 4.0 3.5 3.0 2.5 2.0 1.5 1.0 0.5 0.0 -0.5
f1 (ppm)
D
Si
N
N
+
CO2−
HO2C

**S19**

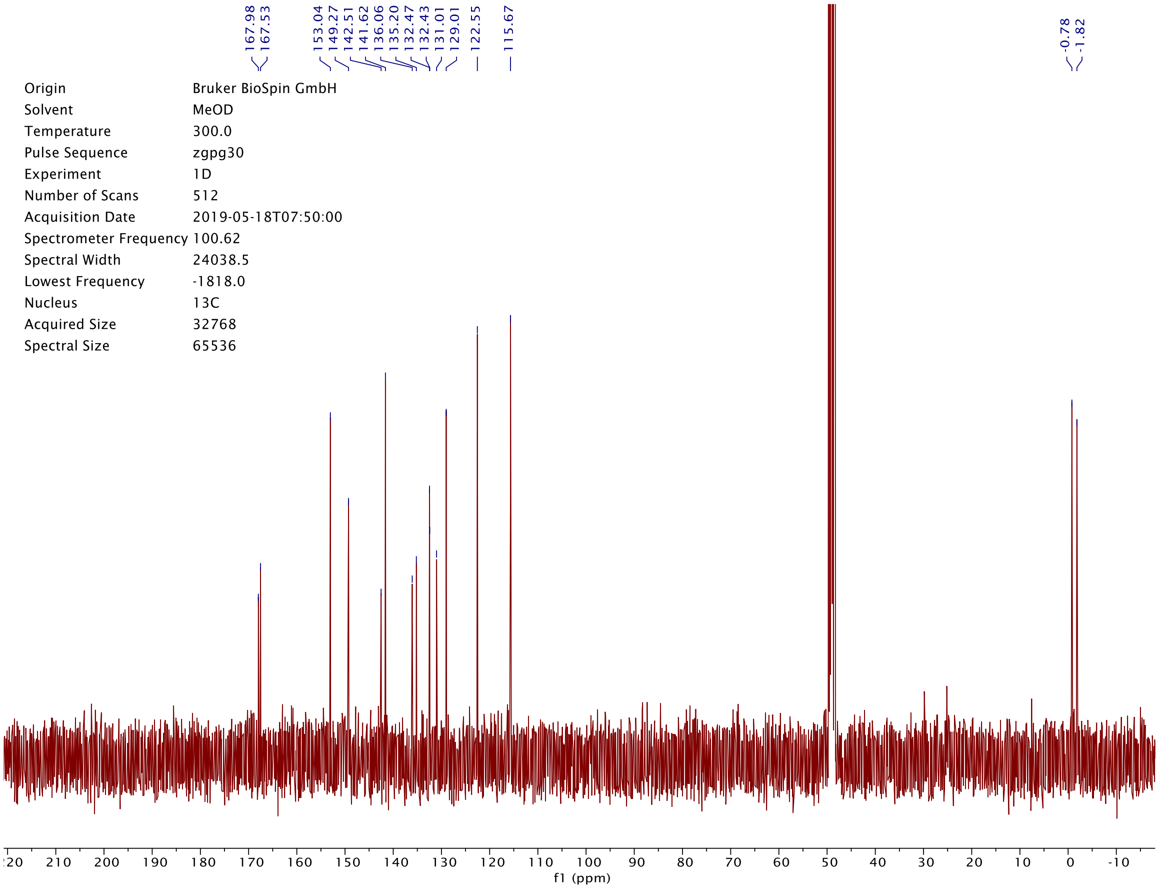
167.98
167.53
153.04
149.27
142.51
141.62
136.06
135.20
132.47
132.43
131.01
129.01
122.55
115.67
-0.78
-1.82
Origin Bruker BioSpin GmbH
Solvent MeOD
Temperature 300.0
Pulse Sequence zgpg30
Experiment 1D
Number of Scans 512
Acquisition Date 2019-05-18T07:50:00
Spectrometer Frequency 100.62
Spectral Width 24038.5
Lowest Frequency -1818.0
Nucleus 13C
Acquired Size 32768
Spectral Size 65536
210 200 190 180 170 160 150 140 130 120 110 100 90 80 70 60 50 40 30 20 10 0 -10
f1 (ppm)

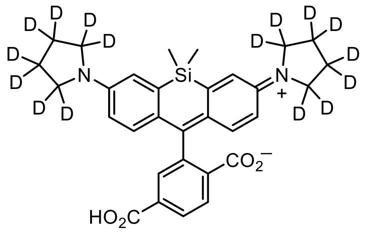

**S19**

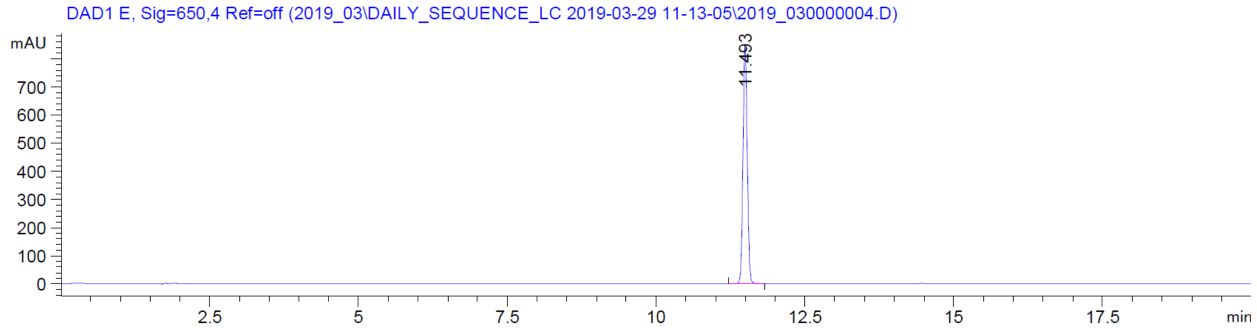
DAD1 E, Sig=650,4 Ref=off (2019_03\DAILY_SEQUENCE_LC 2019-03-29 11-13-05\2019_030000004.D)
mAU
11.493
min

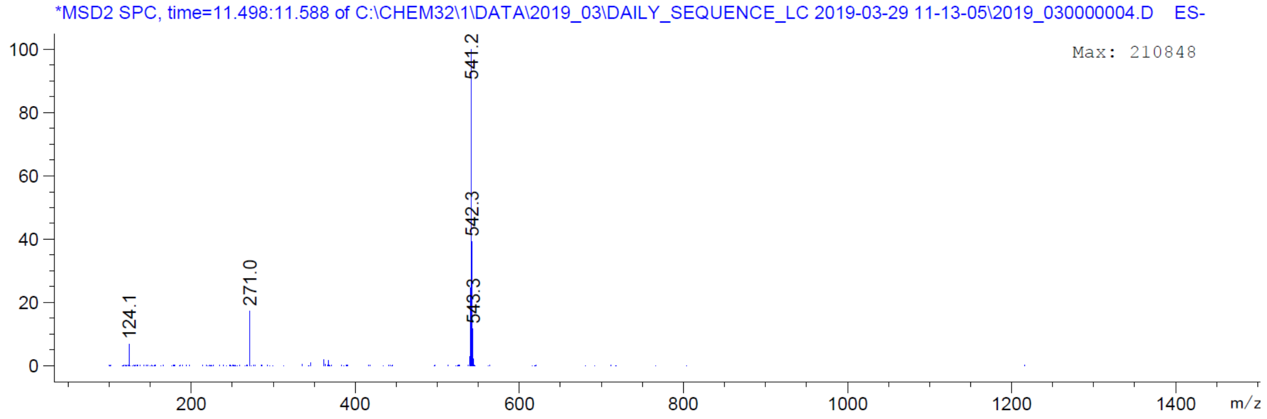
*MSD2 SPC, time=11.498:11.588 of C:\CHEM32\1\DATA\2019_03\DAILY_SEQUENCE_LC 2019-03-29 11-13-05\2019_030000004.D ES-
Max: 210848
541.2
542.3
543.3
271.0
124.1
m/z

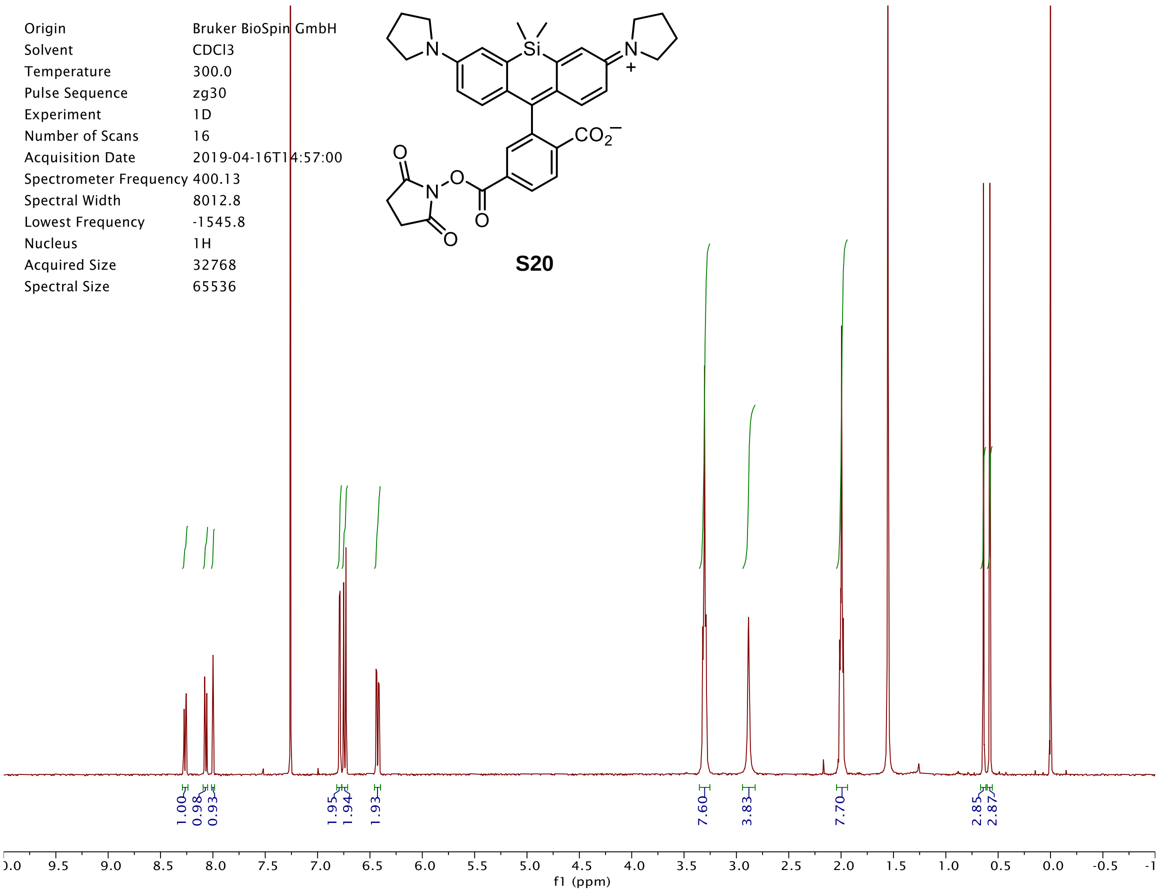
Origin Bruker BioSpin GmbH
Solvent CDCl3
Temperature 300.0
Pulse Sequence zg30
Experiment 1D
Number of Scans 16
Acquisition Date 2019-04-16T14:57:00
Spectrometer Frequency 400.13
Spectral Width 8012.8
Lowest Frequency -1545.8
Nucleus 1H
Acquired Size 32768
Spectral Size 65536
S20
1.00
0.98
0.93
1.95
1.94
1.93
7.60
3.83
7.70
2.85
2.87
f1 (ppm)

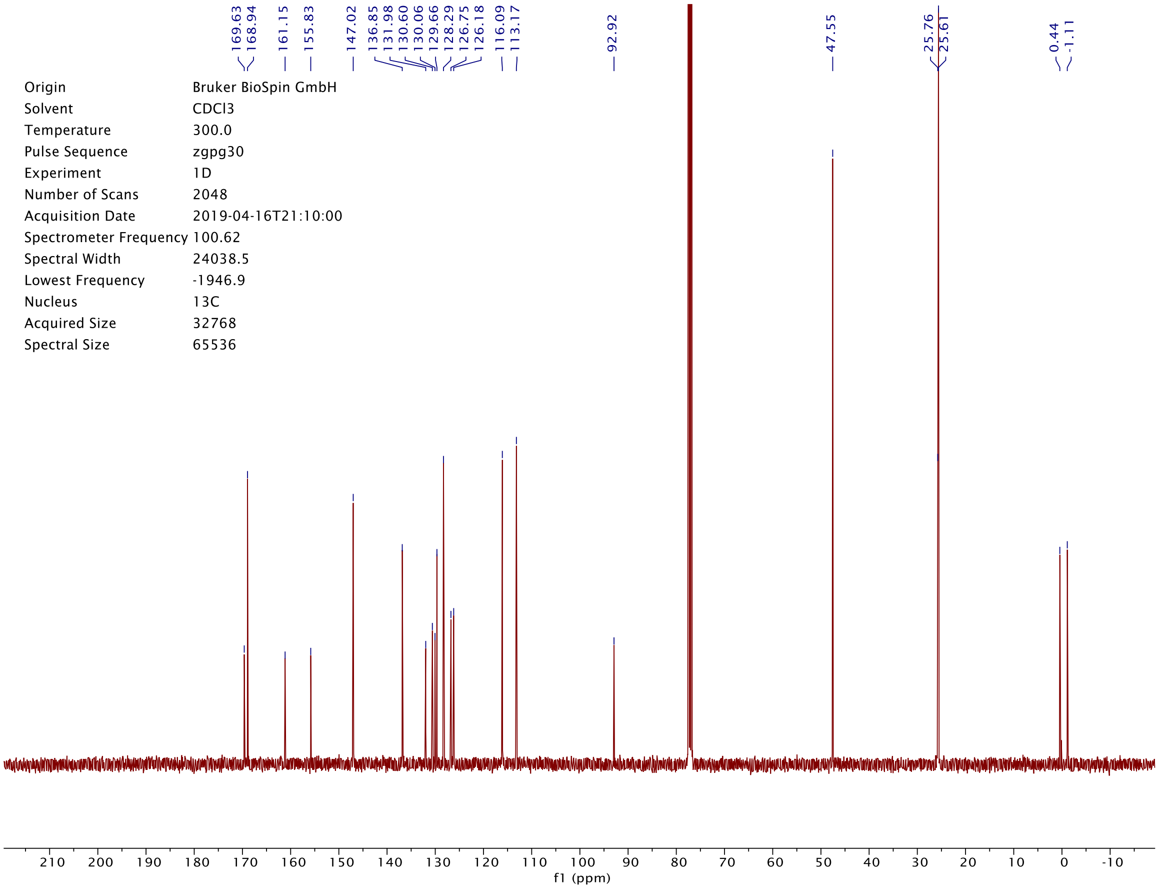
169.63
168.94
161.15
155.83
147.02
136.85
131.98
130.60
130.06
129.66
128.29
126.75
126.18
116.09
113.17
92.92
47.55
25.76
25.61
0.44
-1.11
Origin Bruker BioSpin GmbH
Solvent CDCl3
Temperature 300.0
Pulse Sequence zgpg30
Experiment 1D
Number of Scans 2048
Acquisition Date 2019-04-16T21:10:00
Spectrometer Frequency 100.62
Spectral Width 24038.5
Lowest Frequency -1946.9
Nucleus 13C
Acquired Size 32768
Spectral Size 65536
f1 (ppm)

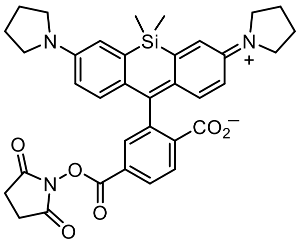
N
Si
N
+
CO2−
O
N
O
O
O
O

**S20**

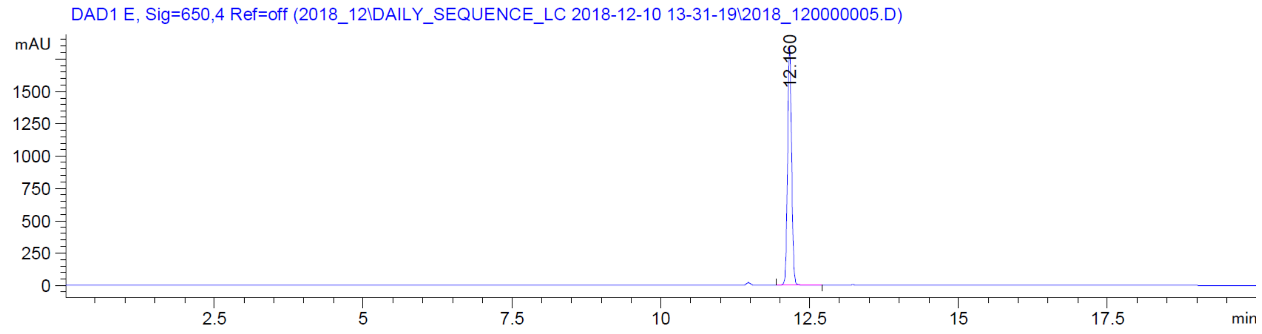
DAD1 E, Sig=650,4 Ref=off (2018_12\DAILY_SEQUENCE_LC 2018-12-10 13-31-19\2018_120000005.D)
mAU
1500
1250
1000
750
500
250
0
12.160
2.5
5
7.5
10
12.5
15
17.5
min

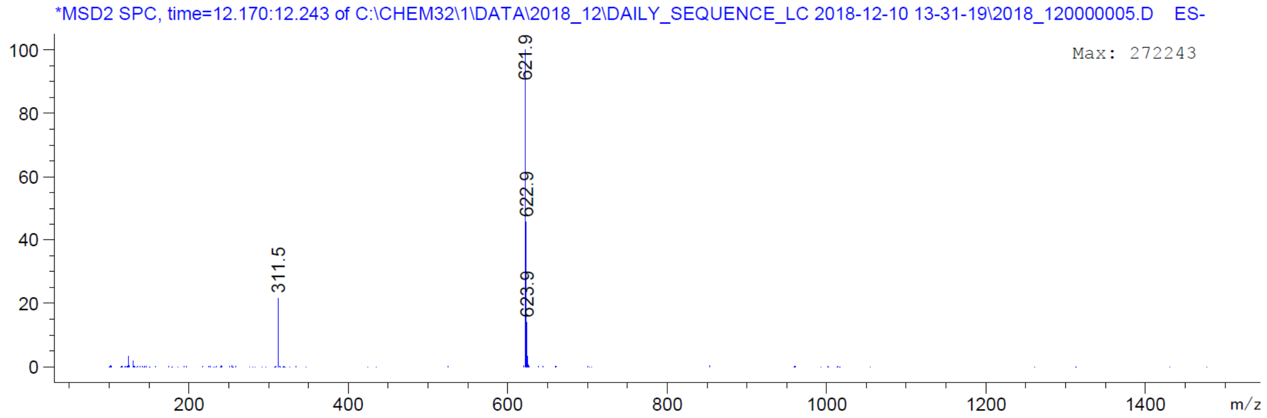
*MSD2 SPC, time=12.170:12.243 of C:\CHEM32\1\DATA\2018_12\DAILY_SEQUENCE_LC 2018-12-10 13-31-19\2018_120000005.D ES-
Max: 272243
100
80
60
40
20
0
621.9
622.9
623.9
311.5
200
400
600
800
1000
1200
1400
m/z

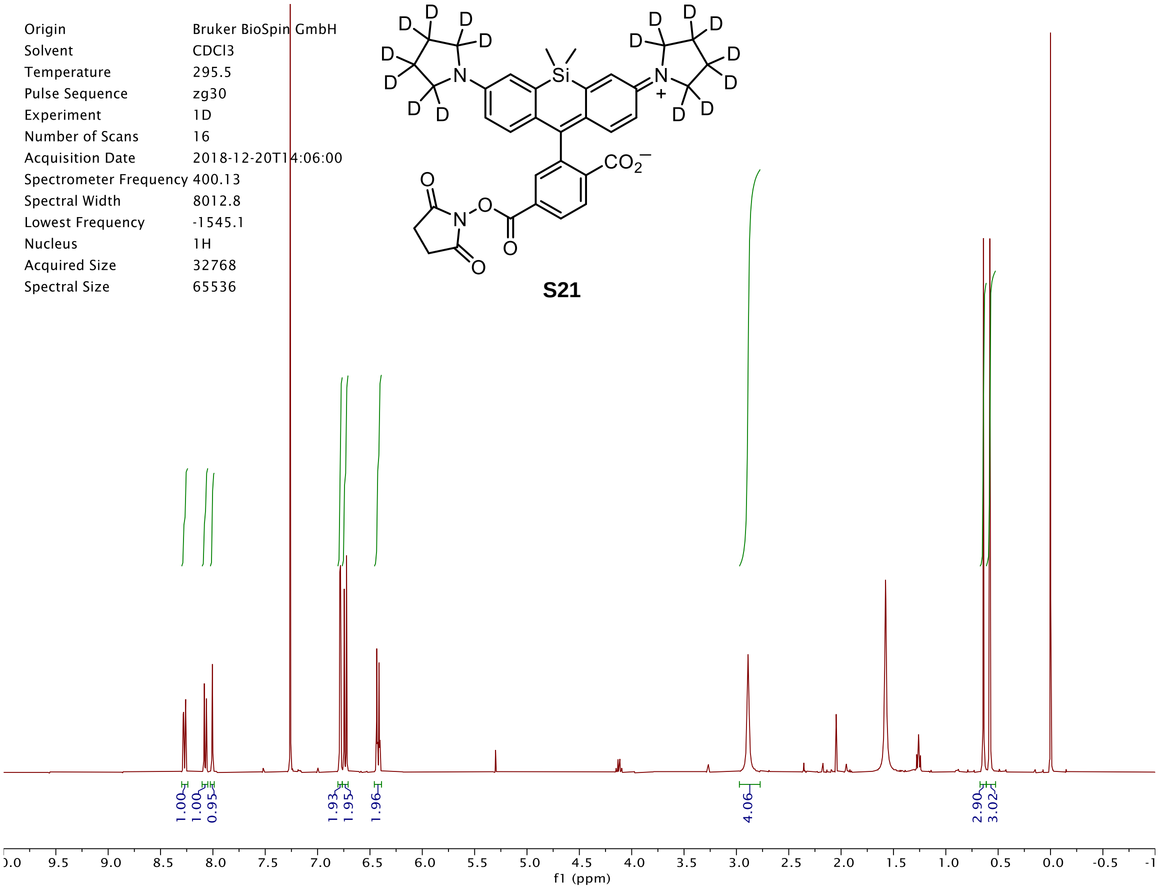
Origin Bruker BioSpin GmbH
Solvent CDCl3
Temperature 295.5
Pulse Sequence zg30
Experiment 1D
Number of Scans 16
Acquisition Date 2018-12-20T14:06:00
Spectrometer Frequency 400.13
Spectral Width 8012.8
Lowest Frequency -1545.1
Nucleus 1H
Acquired Size 32768
Spectral Size 65536
S21
1.00 1.00 0.95 1.93 1.95 1.96 4.06 2.90 3.02
f1 (ppm)

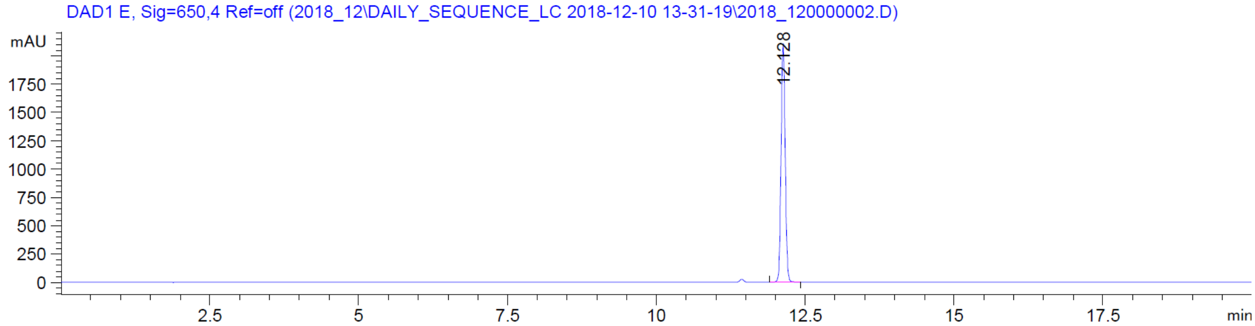
DAD1 E, Sig=650,4 Ref=off (2018_12\DAILY_SEQUENCE_LC 2018-12-10 13-31-19\2018_120000002.D)
mAU
12.128
min

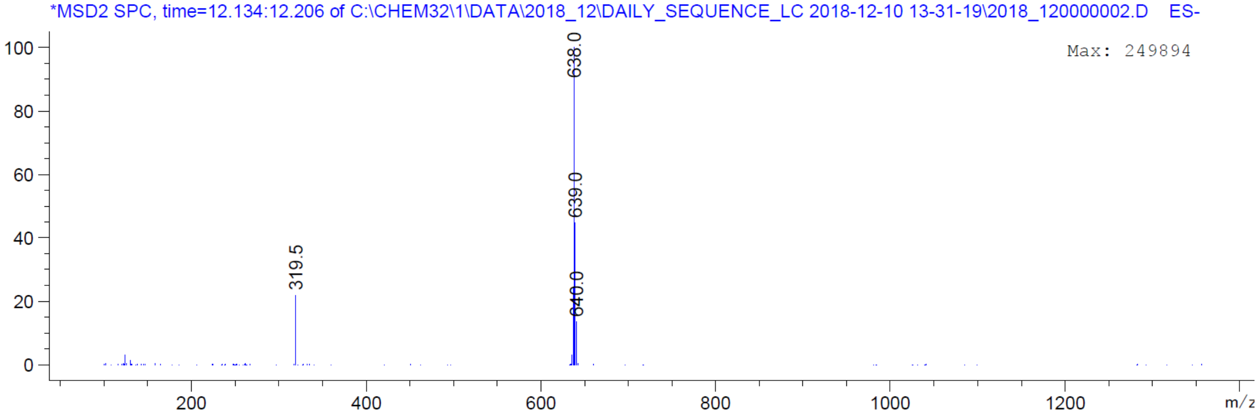
*MSD2 SPC, time=12.134:12.206 of C:\CHEM32\1\DATA\2018_12\DAILY_SEQUENCE_LC 2018-12-10 13-31-19\2018_120000002.D ES-
Max: 249894
638.0
639.0
640.0
319.5
m/z

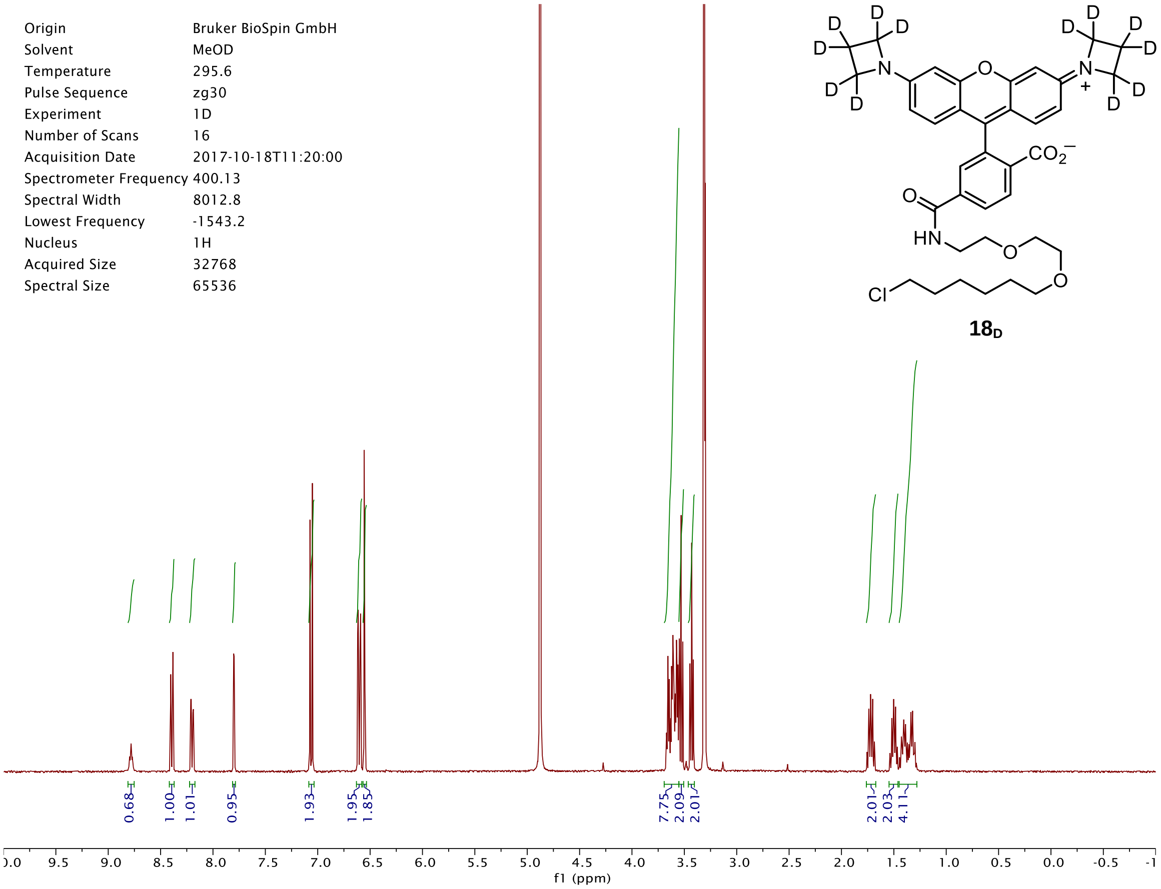
Origin Bruker BioSpin GmbH
Solvent MeOD
Temperature 295.6
Pulse Sequence zg30
Experiment 1D
Number of Scans 16
Acquisition Date 2017-10-18T11:20:00
Spectrometer Frequency 400.13
Spectral Width 8012.8
Lowest Frequency -1543.2
Nucleus 1H
Acquired Size 32768
Spectral Size 65536
18D
0.68
1.00
1.01
0.95
1.93
1.95
1.85
7.75
2.09
2.01
2.01
2.03
4.11
f1 (ppm)

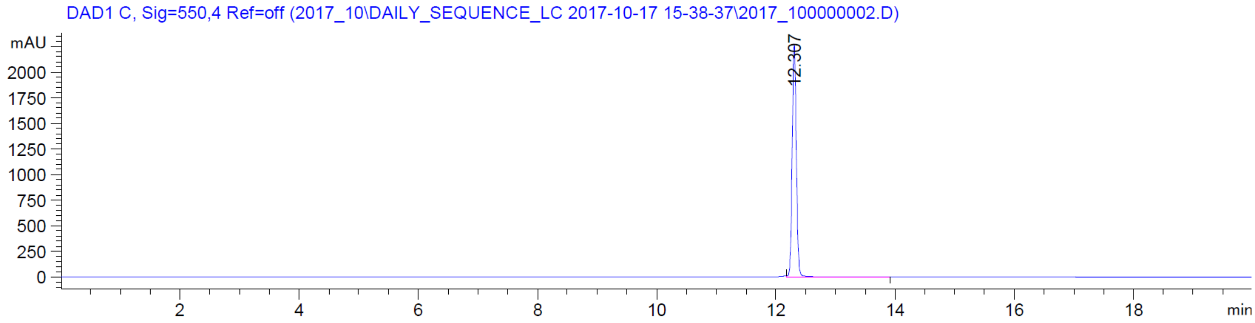
DAD1 C, Sig=550,4 Ref=off (2017_10\DAILY_SEQUENCE_LC 2017-10-17 15-38-37\2017_100000002.D)
mAU
12.307
min

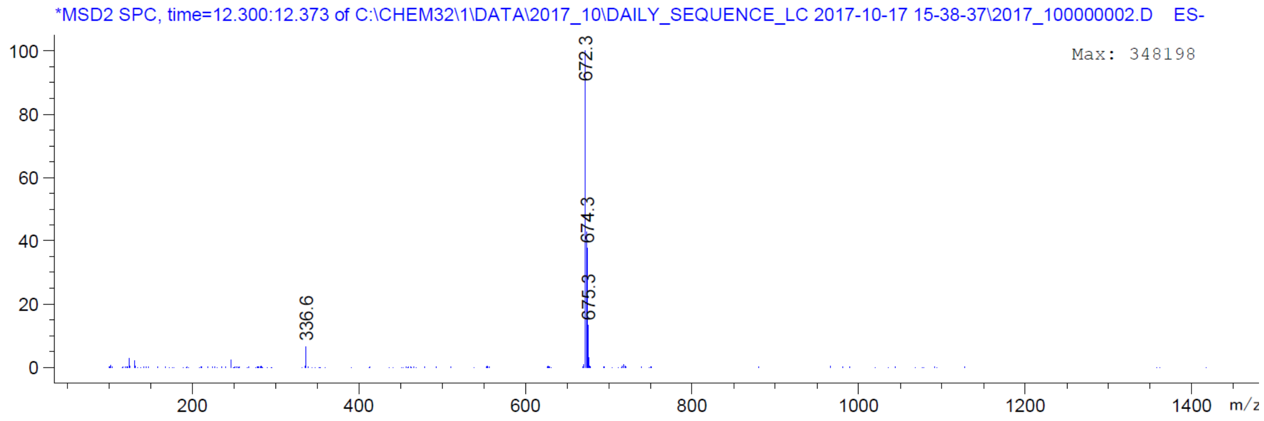
*MSD2 SPC, time=12.300:12.373 of C:\CHEM32\1\DATA\2017_10\DAILY_SEQUENCE_LC 2017-10-17 15-38-37\2017_100000002.D ES-
Max: 348198
672.3
674.3
675.3
336.6
m/z

| | |
|---|---|
| Origin | Bruker BioSpin GmbH |
| Solvent | MeOD |
| Temperature | 300.0 |
| Pulse Sequence | zg30 |
| Experiment | 1D |
| Number of Scans | 16 |
| Acquisition Date | 2017-05-04T11:15:00 |
| Spectrometer Frequency | 400.13 |
| Spectral Width | 8012.8 |
| Lowest Frequency | -1543.1 |
| Nucleus | 1H |
| Acquired Size | 32768 |
| Spectral Size | 65536 |

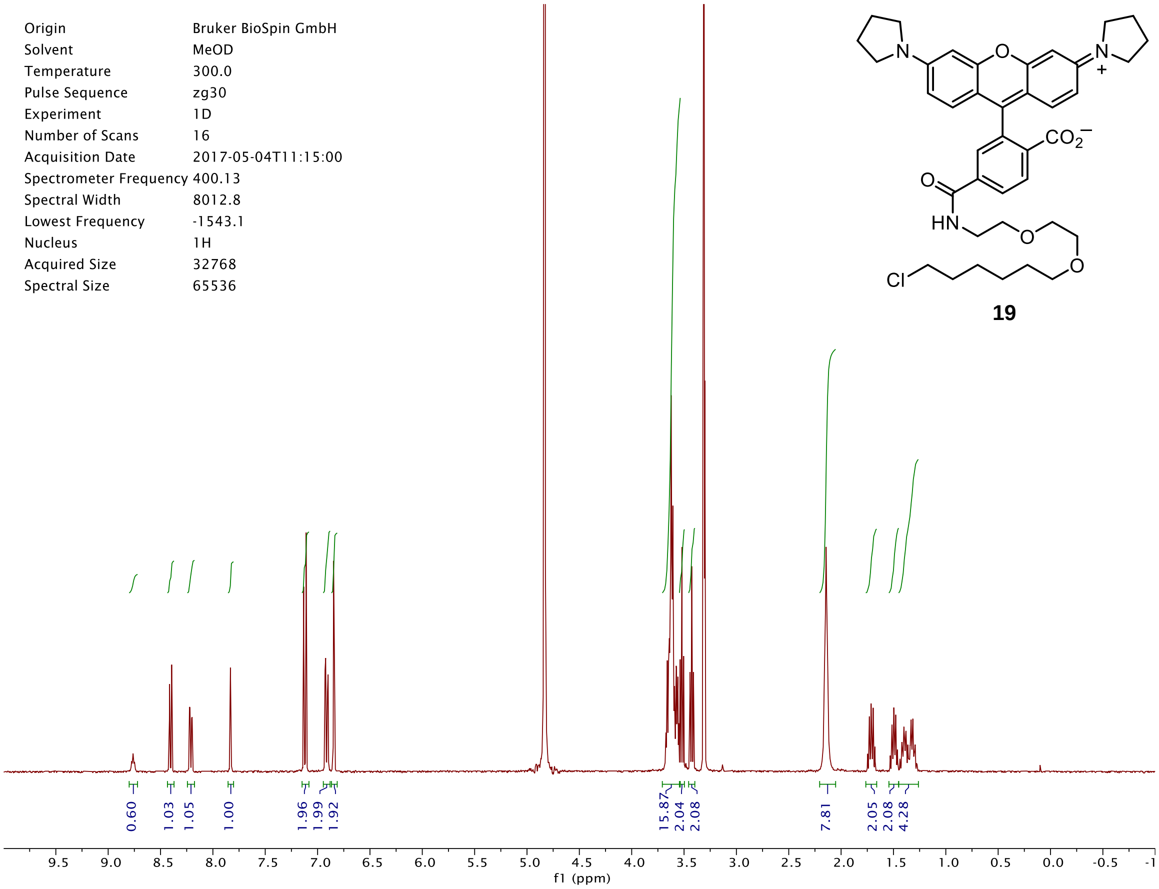


DAD1 C, Sig=550,4 Ref=off (2017_04\DAILY_SEQUENCE_LC 2017-05-02 11-11-33\2017_040000003.D)

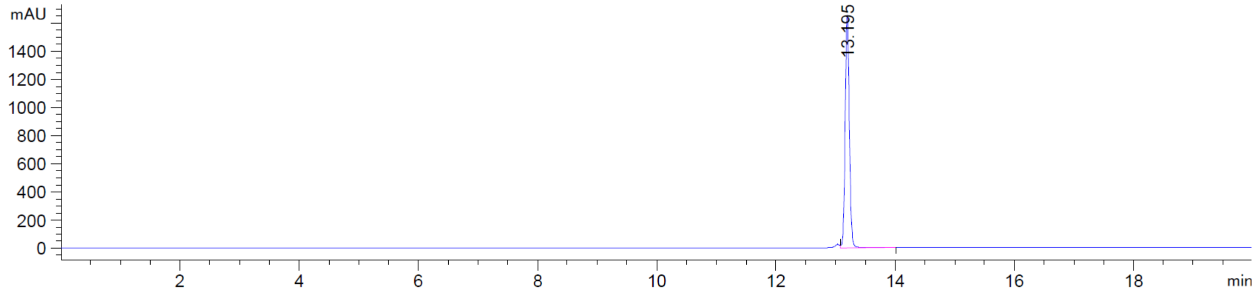


*MSD2 SPC, time=13.191:13.282 of C:\CHEM32\1\DATA\2017_04\DAILY_SEQUENCE_LC 2017-05-02 11-11-33\2017_040000003.D ES-

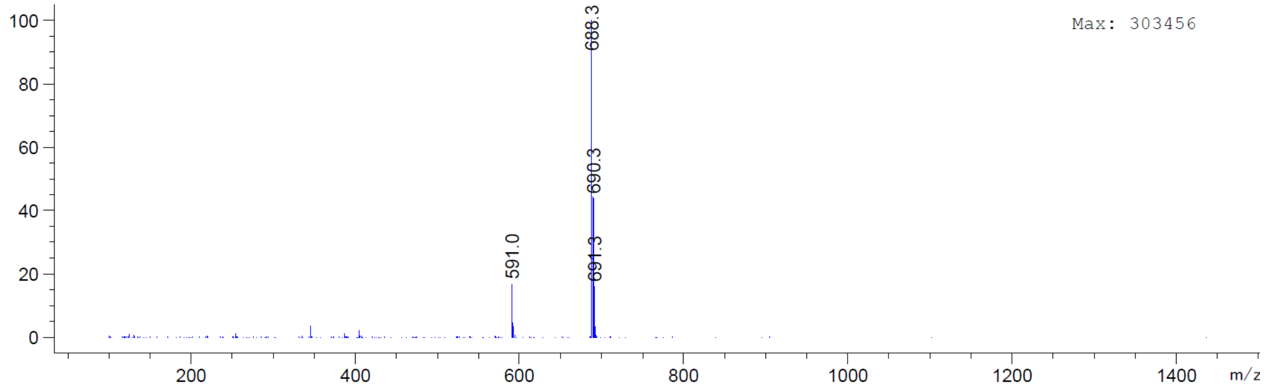

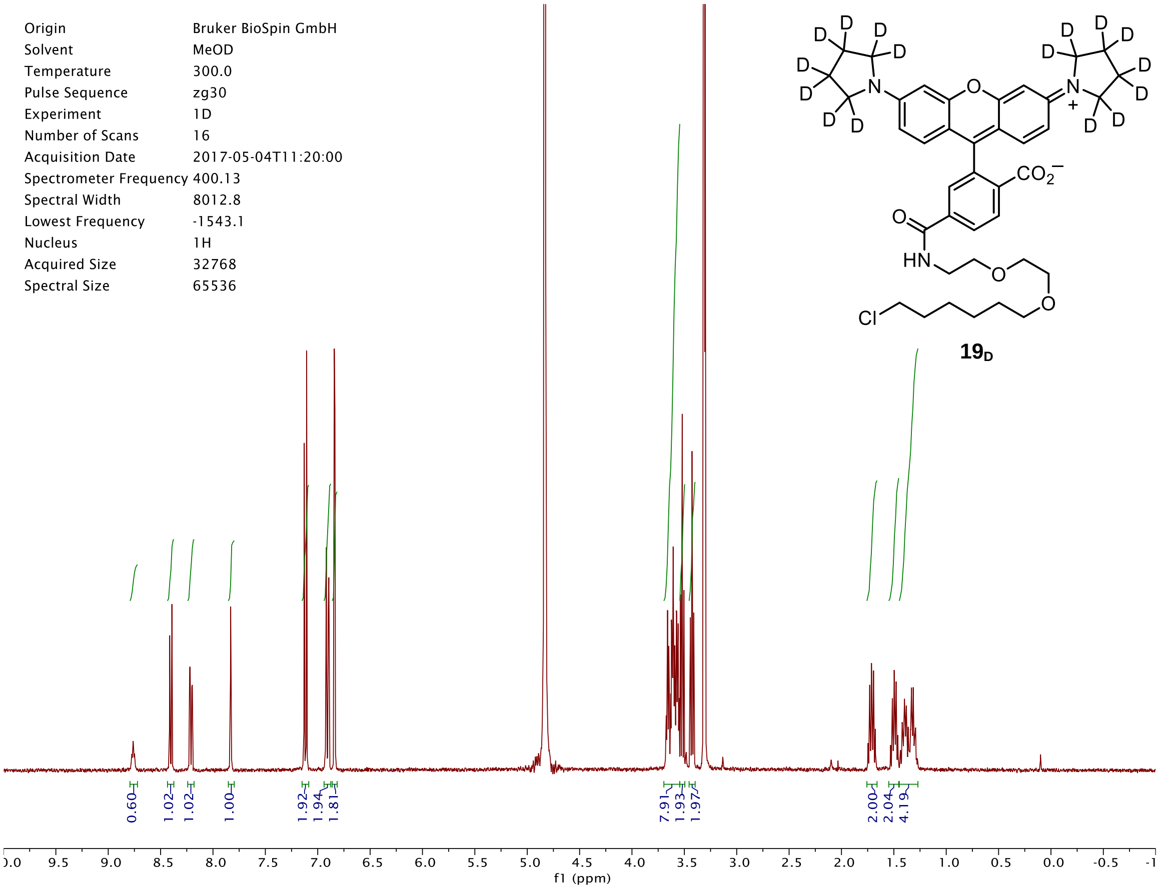

Origin Bruker BioSpin GmbH
Solvent MeOD
Temperature 300.0
Pulse Sequence zg30
Experiment 1D
Number of Scans 16
Acquisition Date 2017-05-04T11:20:00
Spectrometer Frequency 400.13
Spectral Width 8012.8
Lowest Frequency -1543.1
Nucleus 1H
Acquired Size 32768
Spectral Size 65536
$19_D$
0.60
1.02
1.02
1.00
1.92
1.94
1.81
7.91
1.93
1.97
2.00
2.04
4.19
f1 (ppm)


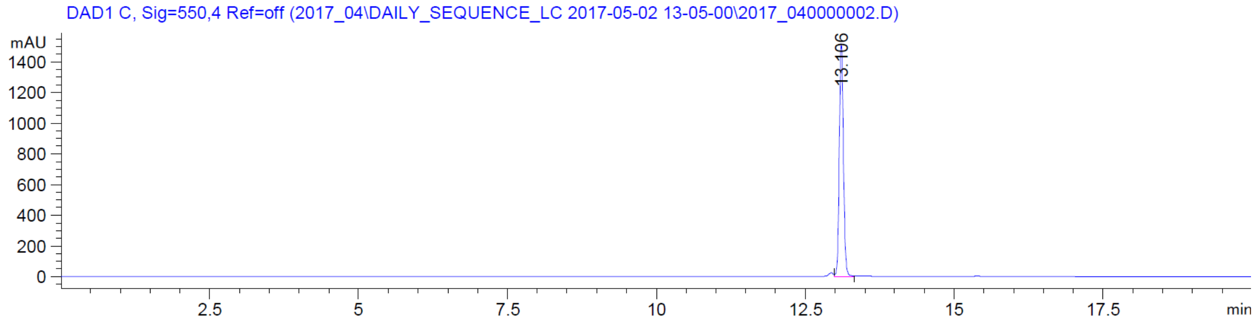

DAD1 C, Sig=550,4 Ref=off (2017_04\DAILY_SEQUENCE_LC 2017-05-02 13-05-00\2017_040000002.D)
mAU
13.106
min


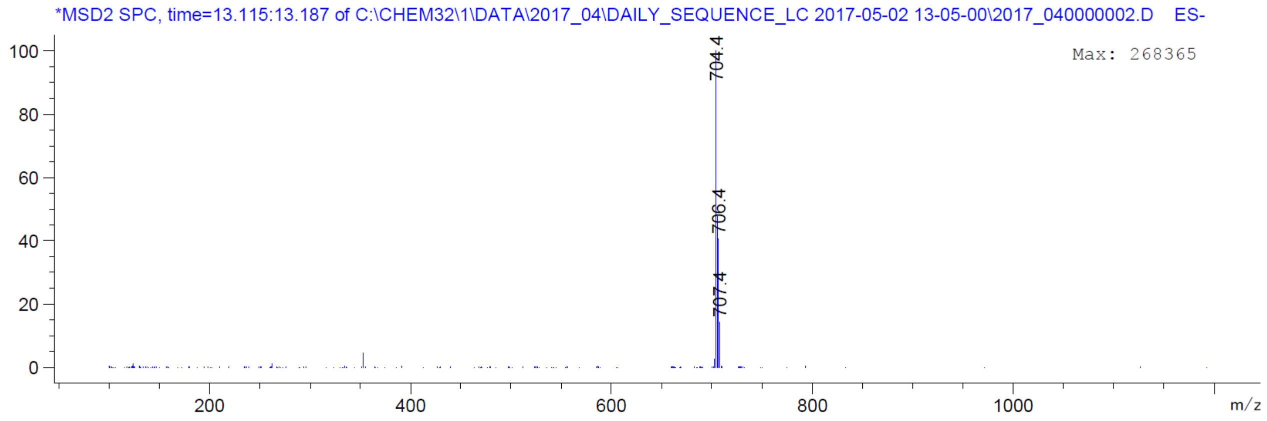

*MSD2 SPC, time=13.115:13.187 of C:\CHEM32\1\DATA\2017_04\DAILY_SEQUENCE_LC 2017-05-02 13-05-00\2017_040000002.D ES-
Max: 268365
704.4
706.4
707.4
m/z

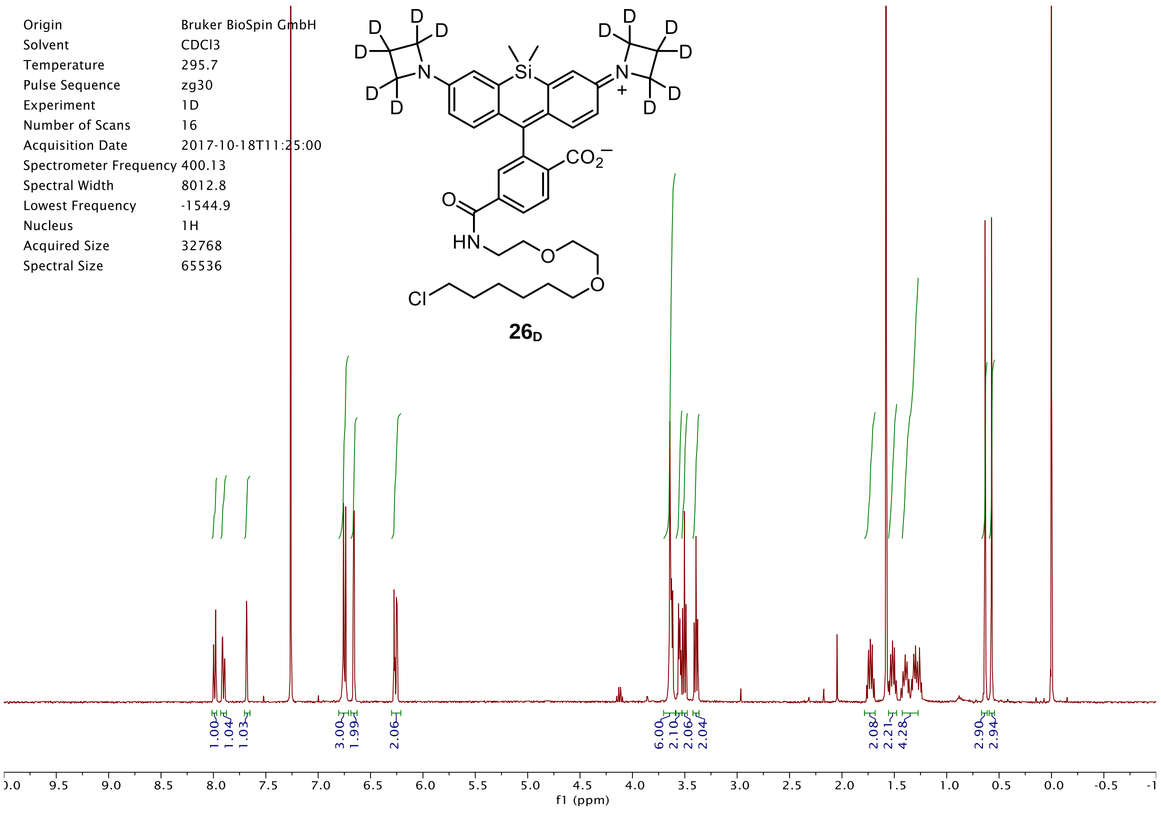
Origin Bruker BioSpin GmbH
Solvent CDCl3
Temperature 295.7
Pulse Sequence zg30
Experiment 1D
Number of Scans 16
Acquisition Date 2017-10-18T11:25:00
Spectrometer Frequency 400.13
Spectral Width 8012.8
Lowest Frequency -1544.9
Nucleus 1H
Acquired Size 32768
Spectral Size 65536
$26_D$
1.00 1.04 1.03 3.00 1.99 2.06 6.00 2.10 2.06 2.04 2.08 2.21 4.28 2.90 2.94
f1 (ppm)

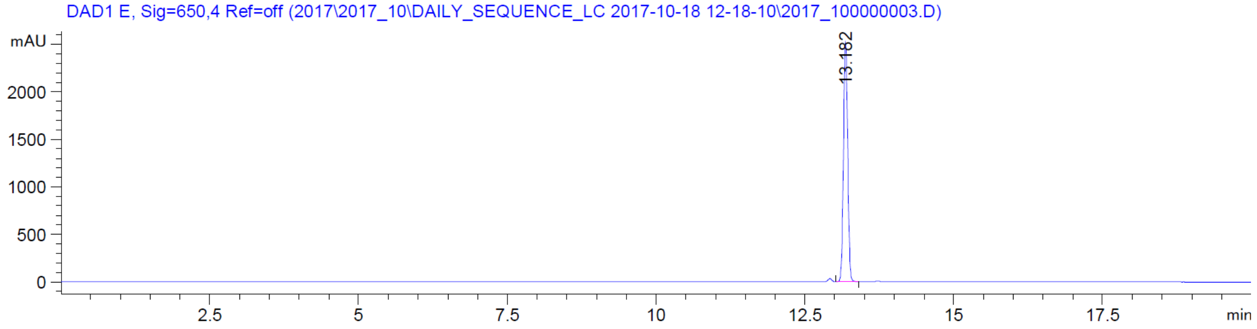
DAD1 E, Sig=650,4 Ref=off (2017\2017_10\DAILY_SEQUENCE_LC 2017-10-18 12-18-10\2017_100000003.D)
mAU
13.182
min

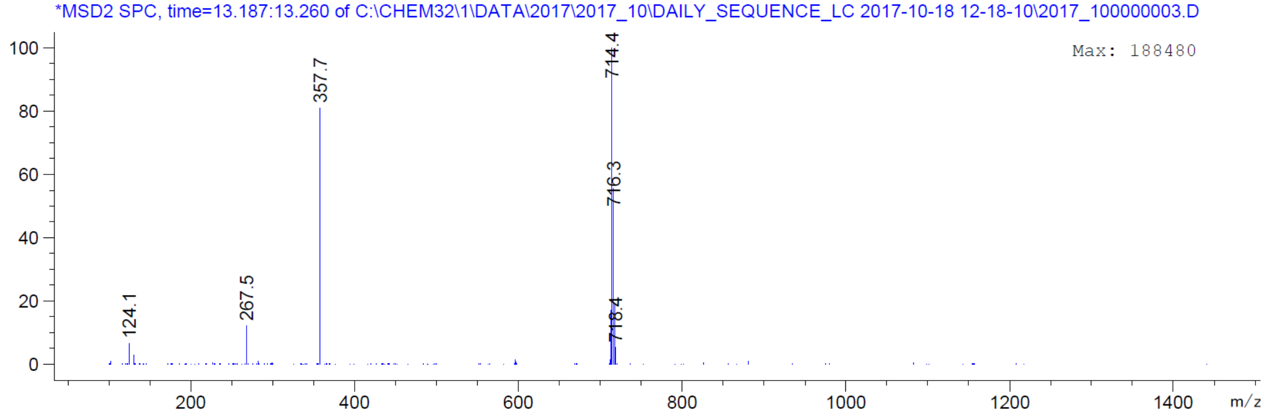
*MSD2 SPC, time=13.187:13.260 of C:\CHEM32\1\DATA\2017\2017_10\DAILY_SEQUENCE_LC 2017-10-18 12-18-10\2017_100000003.D
Max: 188480
124.1
267.5
357.7
714.4
716.3
718.4
m/z

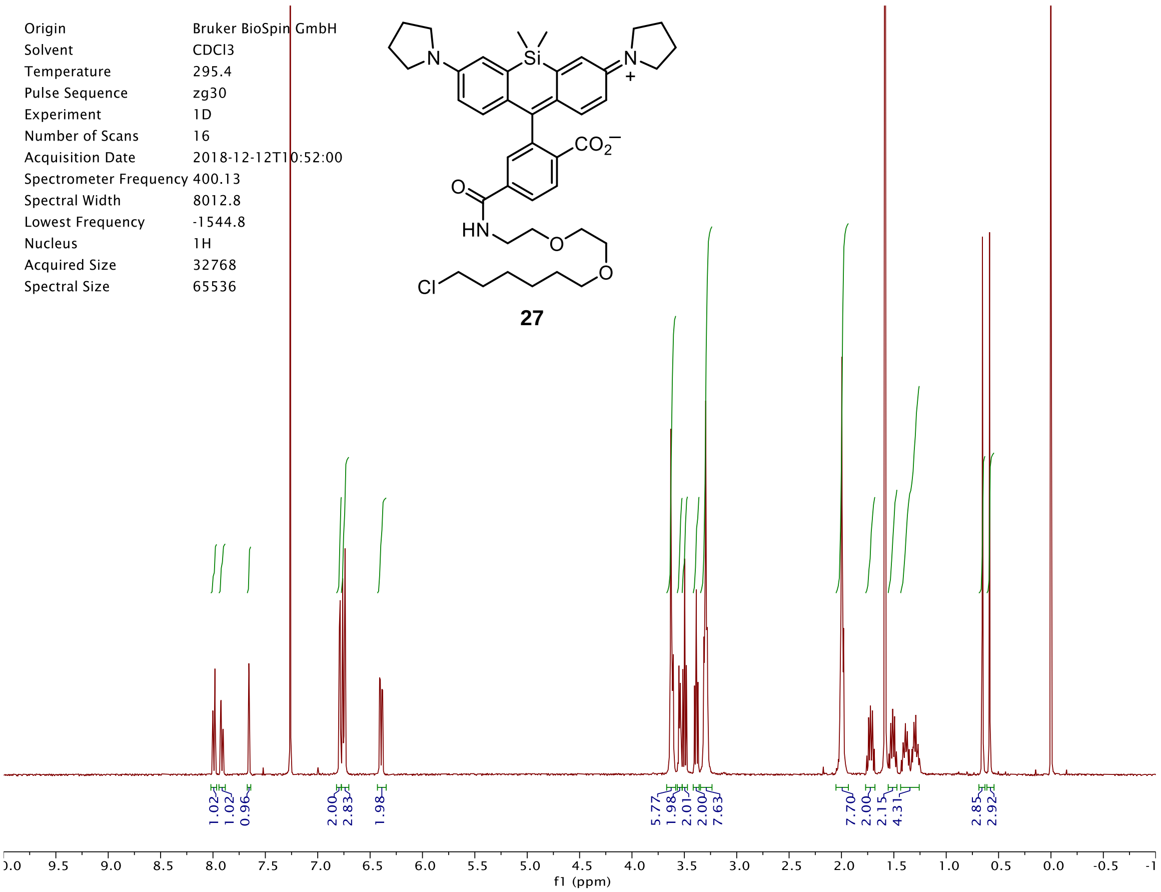
Origin Bruker BioSpin GmbH
Solvent CDCl3
Temperature 295.4
Pulse Sequence zg30
Experiment 1D
Number of Scans 16
Acquisition Date 2018-12-12T10:52:00
Spectrometer Frequency 400.13
Spectral Width 8012.8
Lowest Frequency -1544.8
Nucleus 1H
Acquired Size 32768
Spectral Size 65536
27
1.02
1.02
0.96
2.00
2.83
1.98
5.77
1.98
2.01
2.00
7.63
7.70
2.00
2.15
4.31
2.85
2.92
f1 (ppm)

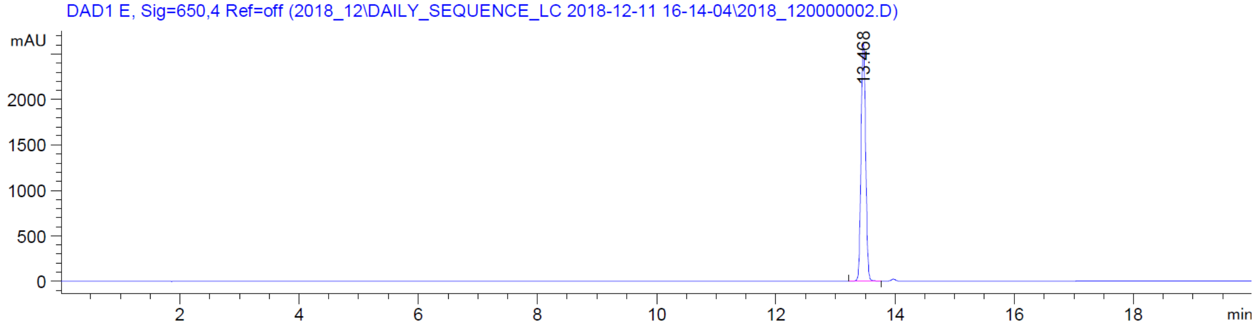
DAD1 E, Sig=650,4 Ref=off (2018_12\DAILY_SEQUENCE_LC 2018-12-11 16-14-04\2018_120000002.D)
mAU
13.468
min

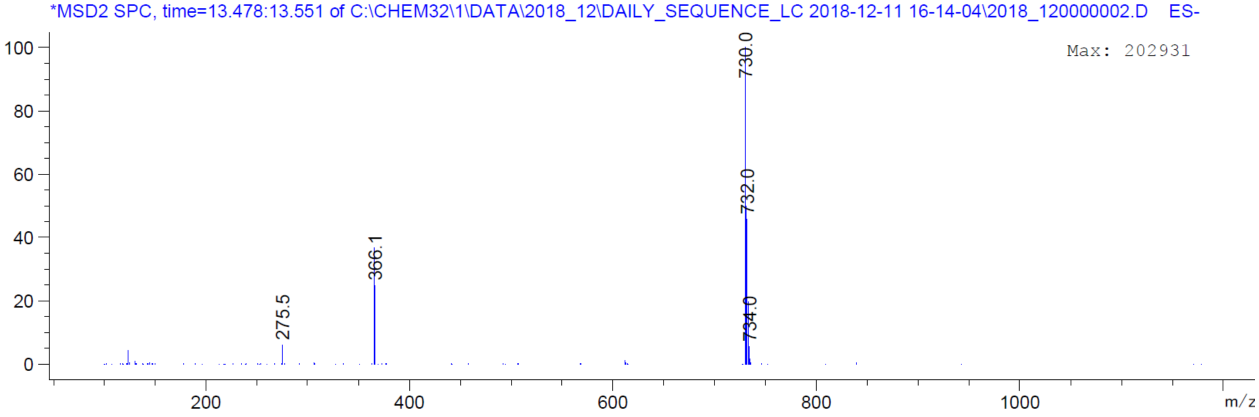
*MSD2 SPC, time=13.478:13.551 of C:\CHEM32\1\DATA\2018_12\DAILY_SEQUENCE_LC 2018-12-11 16-14-04\2018_120000002.D ES-
Max: 202931
730.0
732.0
734.0
366.1
275.5
m/z

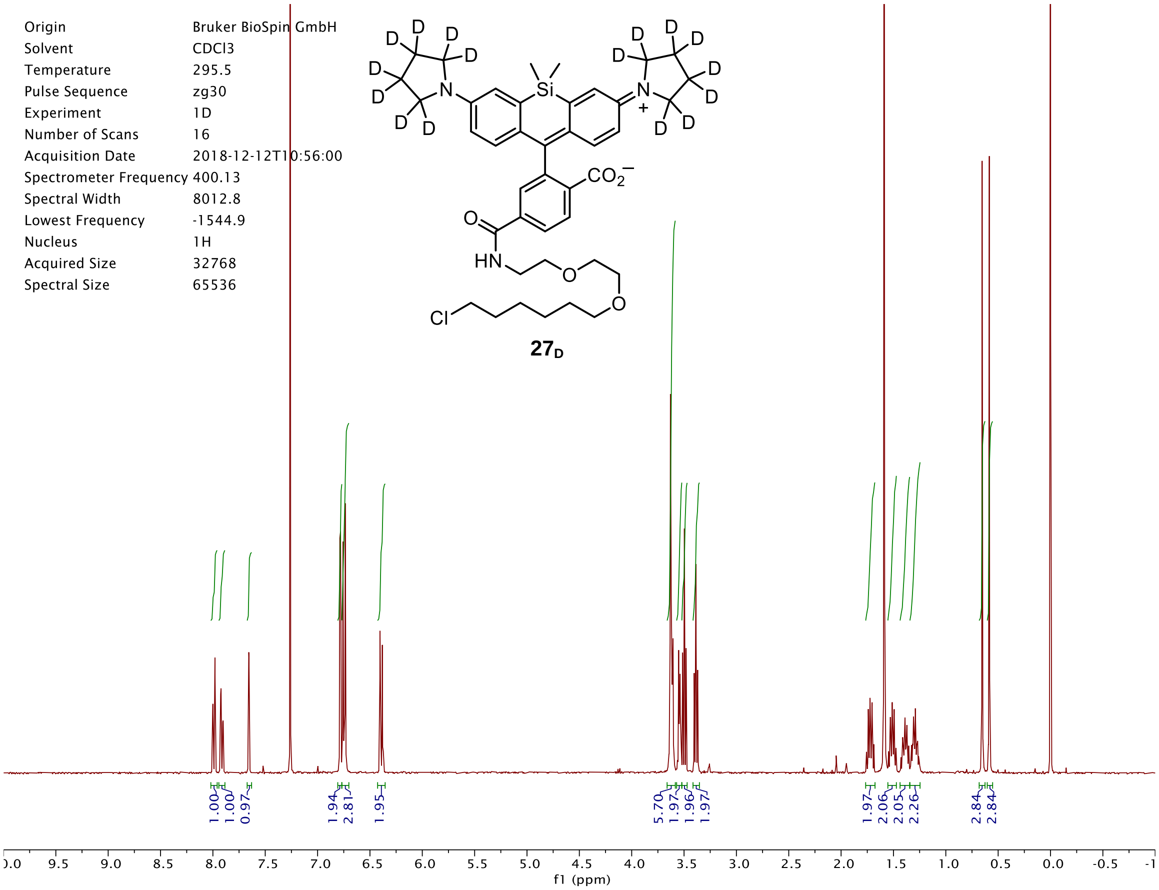

Origin Bruker BioSpin GmbH
Solvent CDCl3
Temperature 295.5
Pulse Sequence zg30
Experiment 1D
Number of Scans 16
Acquisition Date 2018-12-12T10:56:00
Spectrometer Frequency 400.13
Spectral Width 8012.8
Lowest Frequency -1544.9
Nucleus 1H
Acquired Size 32768
Spectral Size 65536
$27_D$
1.00 1.00 0.97 1.94 2.81 1.95 5.70 1.97 1.96 1.97 1.97 2.06 2.05 2.26 2.84 2.84
f1 (ppm)


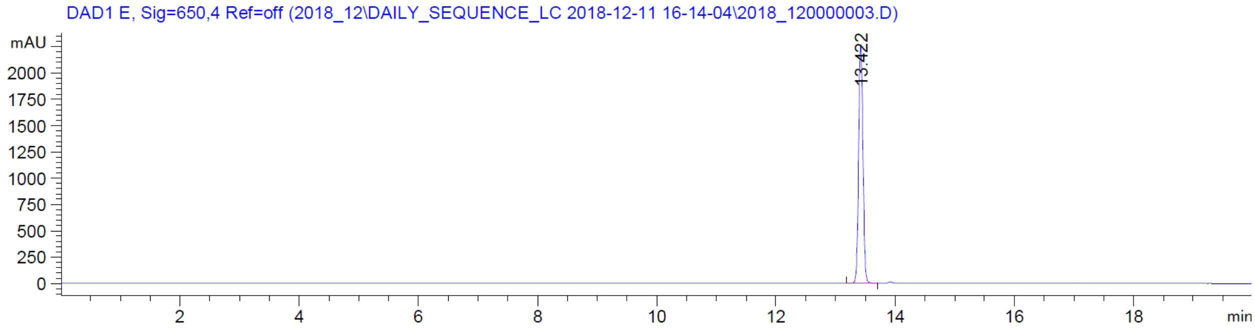

DAD1 E, Sig=650,4 Ref=off (2018_12\DAILY_SEQUENCE_LC 2018-12-11 16-14-04\2018_120000003.D)
mAU
13.422
min


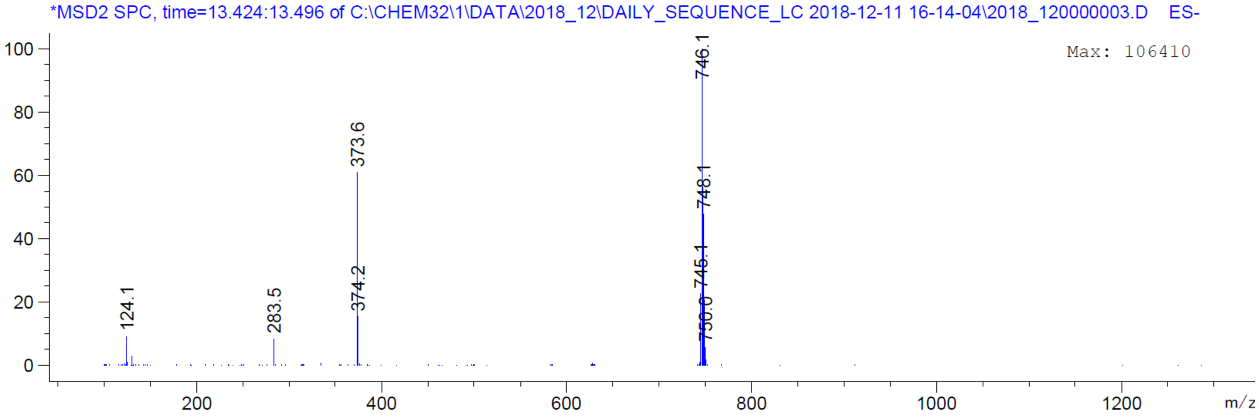

*MSD2 SPC, time=13.424:13.496 of C:\CHEM32\1\DATA\2018_12\DAILY_SEQUENCE_LC 2018-12-11 16-14-04\2018_120000003.D ES-
Max: 106410
746.1
748.1
745.1
750.0
373.6
374.2
283.5
124.1
m/z
